# Differential gene regulation in DAPT-treated Hydra reveals molecular pathways dependent on Notch signalling during interstitial cell differentiation and formation of the oral-aboral axis in *Hydra*

**DOI:** 10.1101/2021.02.09.430419

**Authors:** Jasmin Moneer, Stefan Siebert, Stefan Krebs, Jack Cazet, Andrea Prexl, Qin Pan, Celina Juliano, Angelika Böttger

**Author notes:** Corresponding author: Angelika Böttger, Ludwig-Maximilians University Munich, Biocentre, Großhaderner Str. 2, 82152 Planegg-Martinsried, Germany.

## Abstract

The Notch pathway is highly conserved and essential for animal development. We investigated the function of Notch-signalling in *Hydra* by using the presenilin inhibitor DAPT, which efficiently blocks propagation of Notch-signals. In *Hydra*, DAPT treatment prevents differentiation of proliferating nematocyte progenitor cells into mature nematocytes. Moreover, it causes defects in the *Hydra* head by compromising the head organizer. In order to understand the molecular mechanisms by which the Notch pathway regulates these processes we performed RNAseq to identify genes that are differentially regulated in response to 48 hours of DAPT-exposure. This revealed downregulation of 624 genes and upregulation of 207 genes. To identify candidate direct regulators of Notch-signalling, we also profiled gene expression changes that occur during restoration of Notch-activity 3 and 6 hours after DAPT-removal. We then analysed gene expression patterns of these Notch-responsive genes in untreated animals by interrogating the available single cell sequencing data set for untreated animals and found that almost half of the Notch responsive genes were specifically expressed in nematocytes and nematocyte progenitors. This confirms the critical role for Notch-signalling in nematocyte development. Promoter analyses and gene expression profiling after DAPT-removal suggested an indirect role for Notch in regulating a *POU*-transcription factor, which is critical for nematogenesis. In support of a role for Notch-signalling in head organizer formation, we identified several head organizer genes in the Notch regulated gene data set, including *Cngsc*, a homologue of *goosecoid,* a gene associated with the Spemann organizer, and the Wnt pathway genes *Sp5, Tcf* and *Wnt-7.* Finally, the expression levels of the tentacle patterning genes *HyAlx* and *Sp5* rapidly recovered after DAPT removal. Given that these genes possess Notch-responsive RBPJ transcription factor binding sites in their regulatory regions, these genes are likely directly targeted by Notch signalling. In summary, our data provide a comprehensive picture of the molecular pathways regulated by Notch signalling in interstitial cell differentiation and formation of the oral-aboral axis in *Hydra*.

## Introduction

Notch-signalling facilitates cell fate decisions and pattern formation by inducing terminal differentiation, mediating lateral inhibition, boundary formation and synchronization of developmental processes in animals. Well-studied examples of Notch-regulated processes include the differentiation of the wing margin and the specification of neurons from neuroectoderm in *Drosophila* embryos and somite formation during vertebrate development (Liao and Oates, 2017; Siebel and Lendahl, 2017). The core components of the Notch pathway include the Notch receptor, the DSL-ligands (Delta/ Serrate/ Lag-2), and RBPJ (**r**ecombining **b**inding **p**rotein **s**uppressor of hairless) transcription factors (also called CSL for **C**BF1 in mammals, **S**u(H) in *Drosophila* and **L**ag-1 in *Caenorhabditis*), (Andersson et al., 2011). Both, the DSL ligands and Notch receptors are transmembrane proteins, therefore signalling occurs between directly adjacent cells. Interactions of DSL-ligands with Notch receptors result in cleavage of the Notch receptor by presenilin followed by nuclear translocation of the intracellular domain of Notch (NICD) (reviewed in (Mumm and Kopan, 2000)). NICD works as a transcriptional co-activator of CSL factors. Only a few direct target genes of Notch-signalling have been identified so far (Wang et al., 2015). Targets of Notch-signalling are activated or repressed in different cell types depending on the composition of transcriptional complexes induced by Notch activity and the epigenetic status at the respective loci. A primary and evolutionary conserved target of Notch is the Hey-Hes family of transcriptional repressors. Other context dependent direct target genes of Notch signalling that have been identified include *c-myc, cyclin D1* and *MEK5c* in tumor cells (reviewed in (Borggrefe and Oswald, 2009)). In hematopoietic cells, *GATA3*, the master regulator for T-cell development and several Hox-genes are direct Notch targets (Fang et al., 2007).

To reveal the ancestral core regulatory network directed by the highly conserved Notch signalling pathway, we have focused on a cnidarian, the fresh water polyp *Hydra*. As a sister to bilaterian animals, cnidarians hold an informative phylogenetic position. Moreover, *Hydra* provides the unique opportunity to obtain an animal-wide picture of Notch-target genes with cell-type resolution due to the recently available single cell expression map (Siebert et al., 2019).

*Hydra* polyps have a simple body structure, representing a tube with an oral head structure and an aboral foot. The head consists of the hypostome, with a central mouth opening surrounded by a crest of tentacles. The foot consists of a peduncle, terminating in the basal disc. The body column of the polyp is composed of two epithelial monolayers, termed ectoderm and endoderm, separated by an acellular extracellular matrix, the mesoglea. Ecto- and endoderm are self-renewing epithelial cell lineages. A third cell lineage, the interstitial cells, resides in interstitial spaces of both epithelia (David and Campbell, 1972; David and Gierer, 1974). It is supported by self-renewing multipotent stem cells that provide a steady supply of neurons, gland cells, and nematocytes. Nematocytes are cnidarian specific sensory cells, which harbour the nematocyst or cnidocyst used for capturing prey. Epithelial cells divide along the entire body column of the polyps (Holstein et al., 1991) leading to the displacement of cells towards the oral and aboral ends, and into asexually produced buds. Cells arriving at the base of tentacles or at the basal disc cease cell division and induce differentiation into tentacle or basal disc cells. Buds develop into new polyps and are then released from the parent polyp. Sexual reproduction occurs when interstitial lineage derived germ cells develop into egg and sperm cells (Bosch and David, 1986). Ectodermal tentacle cells are battery cells, where each cell harbours several mature nematocytes. Older cells are shed at the tips of the tentacles and the foot. Due to continual cell divisions, almost all *Hydra* cells are replaced approximately every 20 days (Otto and Campbell, 1977). Therefore, the homeostatic animal is in a constant state of development requiring the presence of signalling for patterning the body axis and direct cell fate specification (Steele, 2002).

The *Hydra* Notch pathway components include the receptor HvNotch, the ligand HyJagged and the CSL-homolog, HvSu(H). The basic mechanisms of Notch signalling are conserved in *Hydra* including regulated intramembrane proteolysis (RIP) through presenilin, followed by nuclear translocation of the intracellular domain of Notch (NICD) (reviewed in (Mumm and Kopan, 2000)). Moreover, the promoter of the *Hydra* HES-family member *HyHes* can be activated by HvNotch-NICD indicating *HyHes* as a direct target of Notch signalling (Käsbauer et al., 2007; Münder et al., 2010; Prexl et al., 2011).

The presenilin inhibitor DAPT efficiently blocks nuclear translocation of NICD and phenocopies Notch loss-of-function mutations in *Drosophila* and zebrafish (Geling et al., 2002; Micchelli et al., 2003). DAPT treatment similarly inhibits NICD translocation in *Hydra*, which results in four strong effects. (1) DAPT blocks post-mitotic differentiation in the nematoblast and germ cell lineages. Early differentiating nematocytes are genetically specified by the expression of the achaete-scute homolog *CnASH* (Grens et al., 1995; Lindgens et al., 2004) and morphologically by the presence of a post-Golgi vacuole as an element of capsule development. This cell state disappears in DAPT treated animals. (2) DAPT blocks post-mitotic differentiation of female germ cells causing proliferating germ cell precursors to form tumor-like growths (Alexandrova et al., 2005; Käsbauer et al., 2007). (3) DAPT impairs boundary formation at both, parent-bud and body column-tentacle boundaries such that typically sharp gene expressed borders margins at these structures become diffuse. At the parent-bud boundary this mis-expression of the *Hydra* FGF-R-homolog *kringelche*n leads to failure of bud foot formation and detachment (Münder et al., 2010; Sudhop et al., 2004). At the base of tentacles, *HyAlx* expression, which demarcates the tentacle boundaries (Smith et al., 2000), becomes diffuse and we observe malformations of the head structure (Münder et al., 2013). (4) DAPT inhibits *Hydra* head regeneration and regenerating tissue is not able to re-establish an oral organiser as evidenced by lack of Wnt-3 expression. This leads to failure in developing a properly patterned head with hypostome and evenly spaced tentacles (Münder et al., 2013).

To gain a better understanding of the underlying molecular causes of the Notch inhibition phenotypes, we aimed to identify the transcriptional target genes of Notch-signalling. We identified 831 genes that were differentially expressed in response to 48 hours of DAPT treatment; 75% of these were down-regulated. Single-cell expression data were used to uncover the gene expression patterns at cell-state resolution for the Notch-responsive genes. We found that Notch responsive genes are expressed in cell states such as differentiating nematocytes and oral cell types, which was consistent with the DAPT-induced phenotypes. To identify potential direct targets of Notch signalling, we also profiled gene expression changes that occur immediately after DAPT removal. Investigating the expression dynamics of Notch responsive genes and performing motif enrichment analysis enabled us to predict likely direct targets of Notch-signalling in *Hydra*.

## Results

### 1. Differential gene expression analysis reveals Notch-responsive genes

To identify targets of Notch signalling in *Hydra*, we elucidated transcriptional changes that occur in response to DAPT treatment. We expected that sustained DAPT treatment would result in the mis-regulation of both, direct and indirect Notch-targets. We furthermore predicted that direct targets would return to control expression levels after DAPT removal more quickly than indirect targets. We therefore profiled gene expression changes immediately after 48 hrs of sustained DAPT treatment (0 hrs time point) to identify all Notch-affected genes. In addition, we profiled gene expression 3 and 6 hrs after DAPT removal to monitor the recovery of these Notch-affected genes.

To characterise the three and six hours post-DAPT treatment time points we used RT-qPCR to profile the expression levels of two genes: 1) *HyHES*, which is a known direct Notch target, (Münder et al., 2010)) and 2) *CnASH*, which is expressed in post-mitotic differentiating nematoblasts (Lindgens et al., 2004), a cell state that is lost in response to DAPT-treatment (Käsbauer et al., 2007). Loss of *CnASH* expression is a secondary (or indirect) effect of Notch inhibition and re-establishment of *CnASH* expression will only occur after DAPT removal once nematogenesis is restored.

As expected, both *HyHES* and *CnASH* were down-regulated after 48 hrs of DAPT treatment. *HyHES* expression returned to normal levels 3 hrs after inhibitor removal, whereas *CnASH* expression was still reduced at 6 hrs (Fig. S1). RNA-seq was therefore performed on tissue samples collected after 48h DAPT treatment (0hrs) and at time points 3 hrs and 6 hrs after DAPT removal, since these intervals appeared sufficient to distinguish between direct and indirect Notch-targets. The workflow for this experiment is illustrated in Fig. 1.

**Figure 1.**
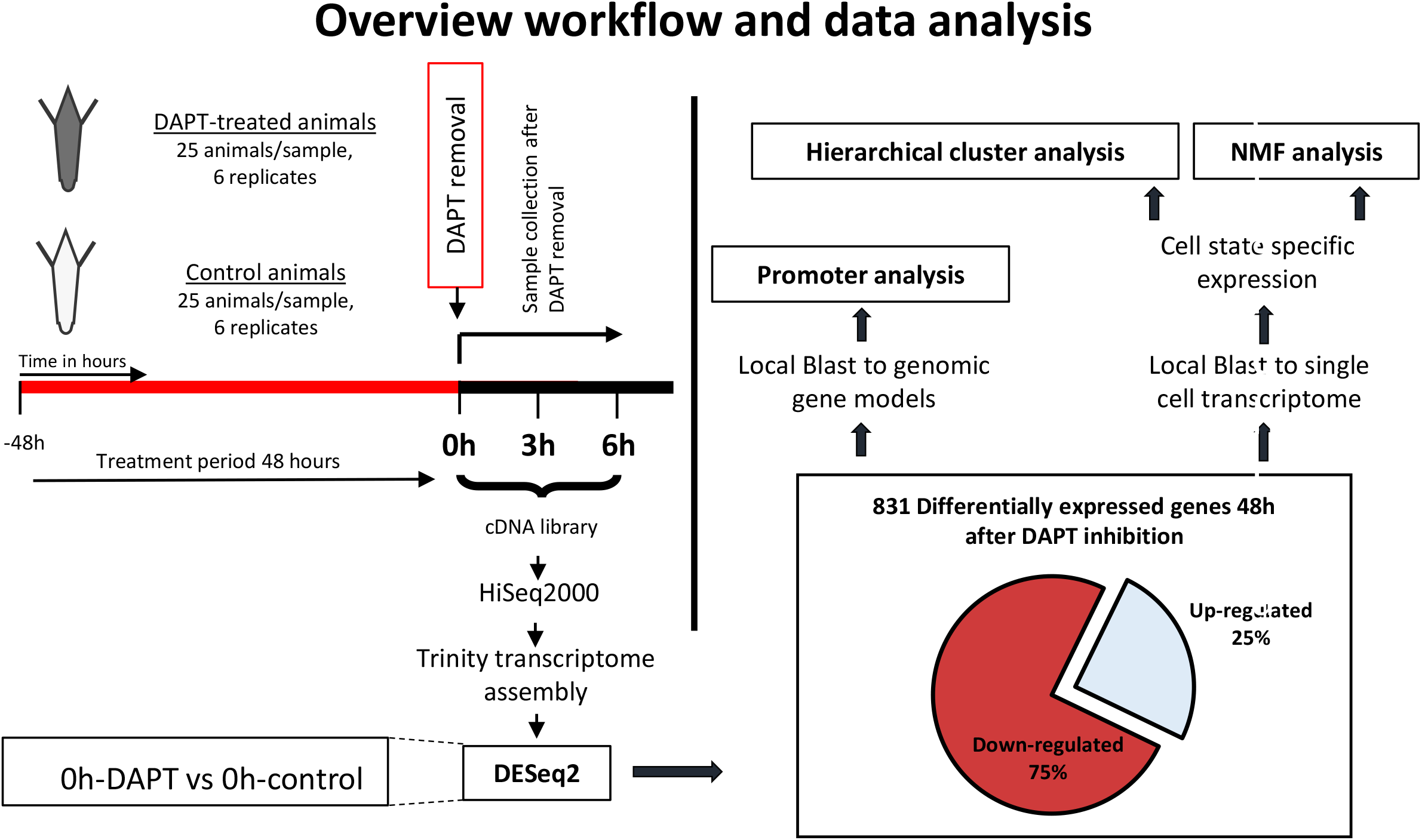
Overview of the experimental and analysis workflow. *Hydra* polyps were treated with either DAPT or DMSO (control) for 48hours. Thereafter, total RNA for sequencing was collected at three time points. The sample 0h was taken immediately after 48hours of DAPT treatment. This is also the time point at which DAPT was removed from the samples and total RNA was collected 3 and 6 hours after DAPT removal. Six biological replicates for each treatment were collected and processed at the same time point. Pairwise differential gene expression analysis was performed between DAPT- and DMSO treated samples for each of the three collection time points. This analysis revealed 831 Notch responsive genes (NR-genes) after 48h of DAPT treatment (0h). For these genes we characterized the expression at time points 3h and 6h. For 666 NR-genes single cell expression data from homeostatic polyps was available (Siebert et al., 2019) and used to elucidate expression pattern and cell state specific expression using hierarchical cluster- and NMF analysis. Additionally, motif enrichment was performed for the set of NR-genes.

Genes that were differentially expressed after 48 hours of DAPT treatment (time point 0 hrs) were referred to as Notch responsive genes (NR-genes). Of the 831 NR-genes identified, 624 were down-regulated (75%) and 207 were up-regulated (25%), (Fig. 2A). Clustering NR-genes according to their fold changes (Fig. 2B) at the three time points after DAPT removal (0 hrs, 3 hrs, and 6 hrs) revealed a group of genes with re-established expression levels at 3hrs, including the confirmed Notch target *HyHES*. A second group of genes showed re-established expression by 6 hrs and a third group of genes was still differentially expressed at 6 hrs, including *CnASH*. 45 genes, including *CnGSC*, were differentially expressed at time points 0 hrs and 6 hrs, but not at 3 hrs (Fig. 2A, “Other”). In all categories, both up- and down-regulated genes were present. Overall, these data reveal changes in gene expression caused by inhibition of the Notch pathway, and uncover which changes are rapidly reversed upon relief of this inhibition. This allowed us to explore the cell type specific effects of DAPT treatment and identify possible direct targets of Notch signalling (full list of NR-genes in Table S1).

**Figure 2.**
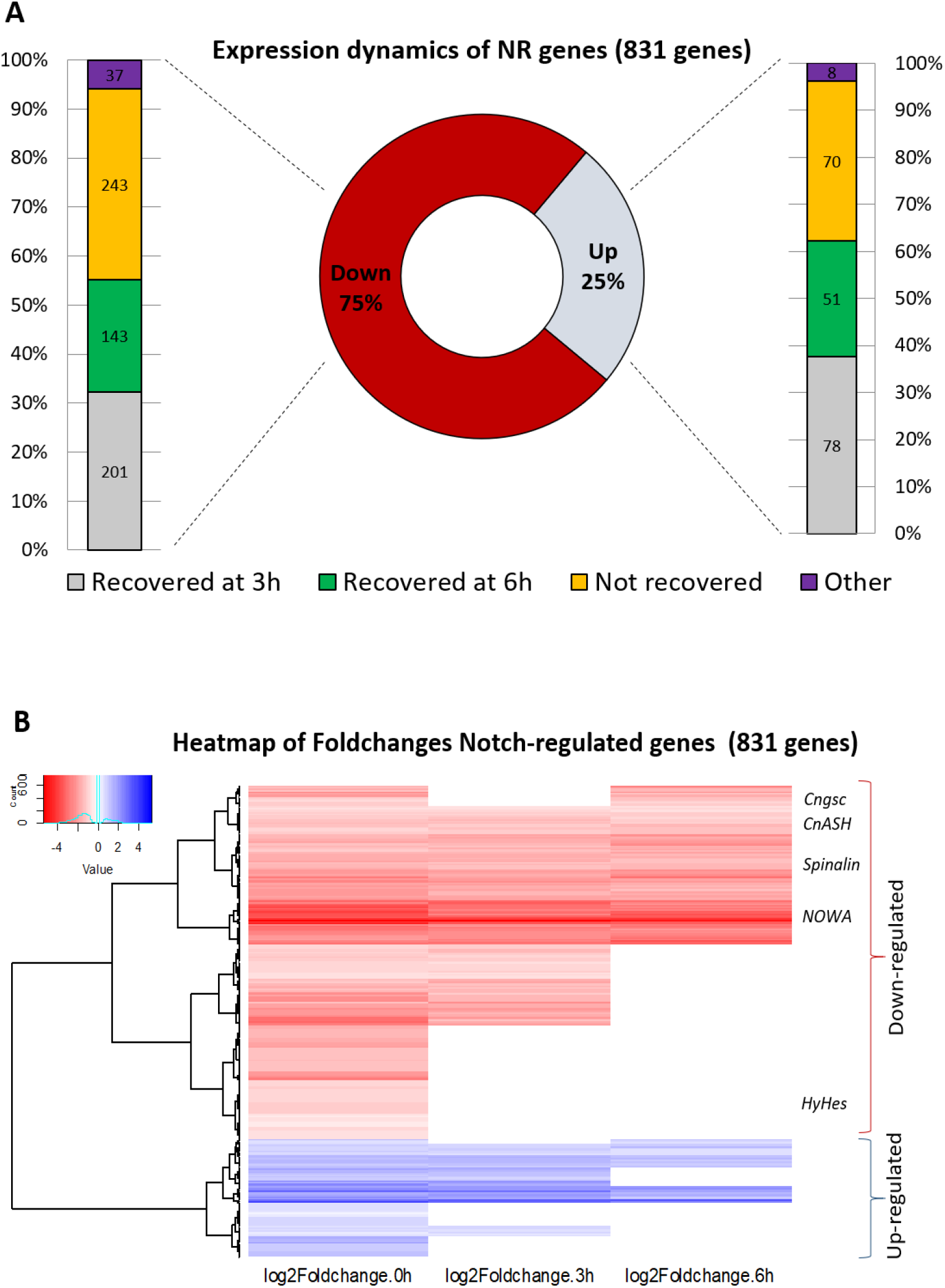
Differential expression of Notch-responsive (NR) genes post DAPT-treatment. A) Expression dynamics of differentially expressed NR-genes and time points for the recovery of original expression levels. For both up-, and down-regulated genes, about 35% recover original expression within the first 3 hours (“Recovered at 3h”), about 25 % recover within 6 hours (“Recovered at 6h”), and about 30-40% do not recover original expression within the time course of the experiment (“Not recovered”). The remaining genes (about 5%, “Other”) behave irregularly, e.g. recovered after 3 hrs, deregulated again after 6 hrs. B) Heatmap highlighting expression differences of all 831 NR-genes. Clustering of NR-genes by their log2Foldchange for each time point revealed up-regulated (blue) and down-regulated (red) genes. No value (white background) means the gene was not differentially expressed at that particular time point and thus had control expression levels. We identified sets of genes that recover their expression by 3h (e.g. *HyHes,* differentially expressed at 0h, thereafter back to control expression level), genes that recover expression by 6h, and genes that do not recover expression during the course of the experiment, i.e. at 6h after DAPT removal (e.g. the post-mitotic nematocyte gene markers *CnASH*, *NOWA* and *Spinalin*). A fourth set includes genes that are differentially expressed at 0h and 6h, but not at the 3h time point (e.g. *CnGSC*).

### 2. Single-cell expression data demonstrate nematogenesis and epithelial expression of Notch-responsive genes

Next, we elucidated NR-gene expression patterns by exploring *Hydra* single-cell expression data, which were available for 666 (80%) NR-genes (Fig. 1). We defined cell state and spatial expression of NR-genes on the basis of published cell state annotations (Siebert et al., 2019). Hierarchical cluster analysis revealed groups of genes expressed in specific cell states (Fig. 3): nematoblasts/nematocytes (red, blue, violet and yellow clusters, 315 genes), ectodermal epithelial cells including battery cells (black cluster, 90 genes), endodermal epithelial cells including tentacle cells (grey cluster, 80 genes), and genes more ubiquitously expressed across cell states (cyan cluster, 79 genes). An additional small subset comprising 15% of NR-genes included genes with restricted expression in several distinct cell states such as specific neurons, gland cells, germline cells or ectodermal basal disk cells (green cluster, 102 genes). The majority of NR-genes fell into two broad categories: 1) 47% of NR-genes are expressed in nematoblasts and nematocysts, and 2) 20% of NR-genes are expressed in epithelial cells (black and grey cluster, Fig. 3).

**Figure 3.**
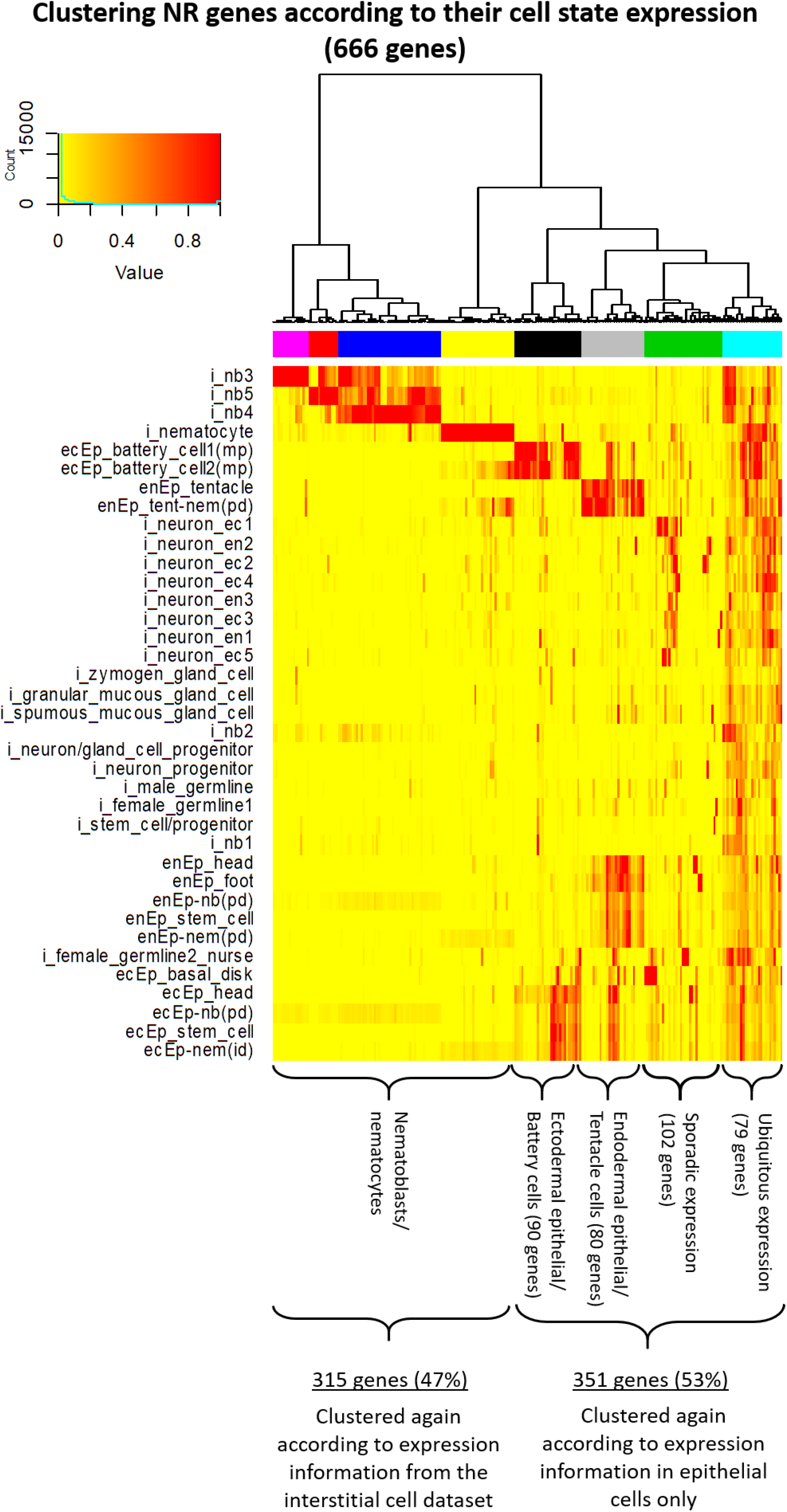
NR-genes expression in homeostatic polyps based on single-cell expression data. Expression data and cell state annotations were retrieved from Siebert et al. (2019). Hierarchical clustering was performed for 666 NR-genes using average expression values for each annotated cell state. This revealed expression in nematoblast/nematocyte-specific genes (violet, red, blue and yellow cluster), ectodermal epithelial cell genes including battery cell genes (black), endodermal epithelial cell genes including tentacle genes (grey), genes ubiquitously expressed across a wide range of cell states (cyan) and genes with a sporadic expression (green). Nematoblast/nematocyte genes constituted 47% of the NR genes. i: cell of the interstitial lineage, nb: nematoblast, ecEp: ectodermal epithelial cell, enEP: endodermal epithelial cell.

In addition, we performed non-negative matrix factorisation (NMF) on the NR-gene set as an unbiased means to uncover modules of co-expressed genes (metagene) and identified 23 metagenes. We then visualized metagene expression on the tSNE (t-distributed stochastic neighbour embedding) representation of selected clusterings from Siebert et al. (2019) (Fig. S2, metagenes). The NMF analysis identified cell state specific modules that were consistent with the hierarchical clustering results (Fig. S2 metagenes and Fig. 3). Interestingly, a single metagene was found to be expressed in female germ line cells suggesting Notch function during female gametogenesis (supplementary metagenes, “IC derivatives - female nurse cells”, A).

#### 2.1. Nematoblast and nematocyte expression of NR-genes

The largest fraction of NR-genes have nematoblast or nematocyte specific expression. In *Hydra*, this lineage comprises four types of nematocytes, each of which harbors a single capsule (or nematocyst) of the atrichous isorhiza-, holotrichous isorhiza-, stenotele- or desmoneme- type. Nematocytes develop from interstitial stem cells via a proliferative amplification phase with incomplete cytokinesis that results in the formation of nests of 4, 8, 16, and 32 nematoblasts. The cells in these nests undergo a final mitosis and start capsule morphogenesis, a process that can be divided into five stages: (1) formation of a growing capsule primordium from a large post-Golgi vacuole, (2) growing of a tubule elongation of the capsule, (3) invagination of the tubule into the capsule, (4) formation of spines inside the invaginated tubule and (5) hardening of the capsule wall. Nests with mature nematocytes break up and single nematocytes then get incorporated into battery cells of the tentacles or into epithelial cells of the body column (David and Gierer, 1974; Engel et al., 2002).

As Notch inhibition by DAPT treatment results in a severe block of nematocyte differentiation, which occurs coincident with or immediately after mitotic exit of differentiating nematoblasts (Käsbauer et al., 2007) we sought to identify the exact differentiation step that was affected. We therefore performed hierarchical clustering for NR-genes with expression in nematoblast or nematocyte cell states (Fig. 3, violet, red, blue and yellow clusters) using the *Hydra* single-cell data (Siebert et al., 2019). The single-cell data revealed four distinct nematocyte differentiation trajectories and gene expression state changes were identified along these trajectories, from stem cells to differentiated nematocytes. Moreover, the single cell analysis revealed 8 distinct nematoblast stages along these four trajectories (nb1 through nb8). Two of these trajectories were annotated as desmoneme and stenotele differentiation, based on marker gene expression (Siebert et al., 2019). In the present study, the clustering of NR-genes expressed during nematogenesis revealed that the majority of those are strongly expressed in cell states nb4, 5, 6, 7, 8 and in differentiated nematocytes (nem), with no or much lower expression in the earlier cell states SC.nb, nb1, nb2 and nb3 (Fig. 4A). This was also observed by plotting the expression of the NR-gene modules onto the single cell tSNE representation (Fig. S2, metagenes) which indicates expression in all three nematocyte types including desmonemes (Fig. S2 “Desmonemes” A, B, C), stenoteles (Fig. S2 “Stenoteles” A, B) and isorhizas (Fig. S2 “Isorhizas” A, B).

**Figure 4.**
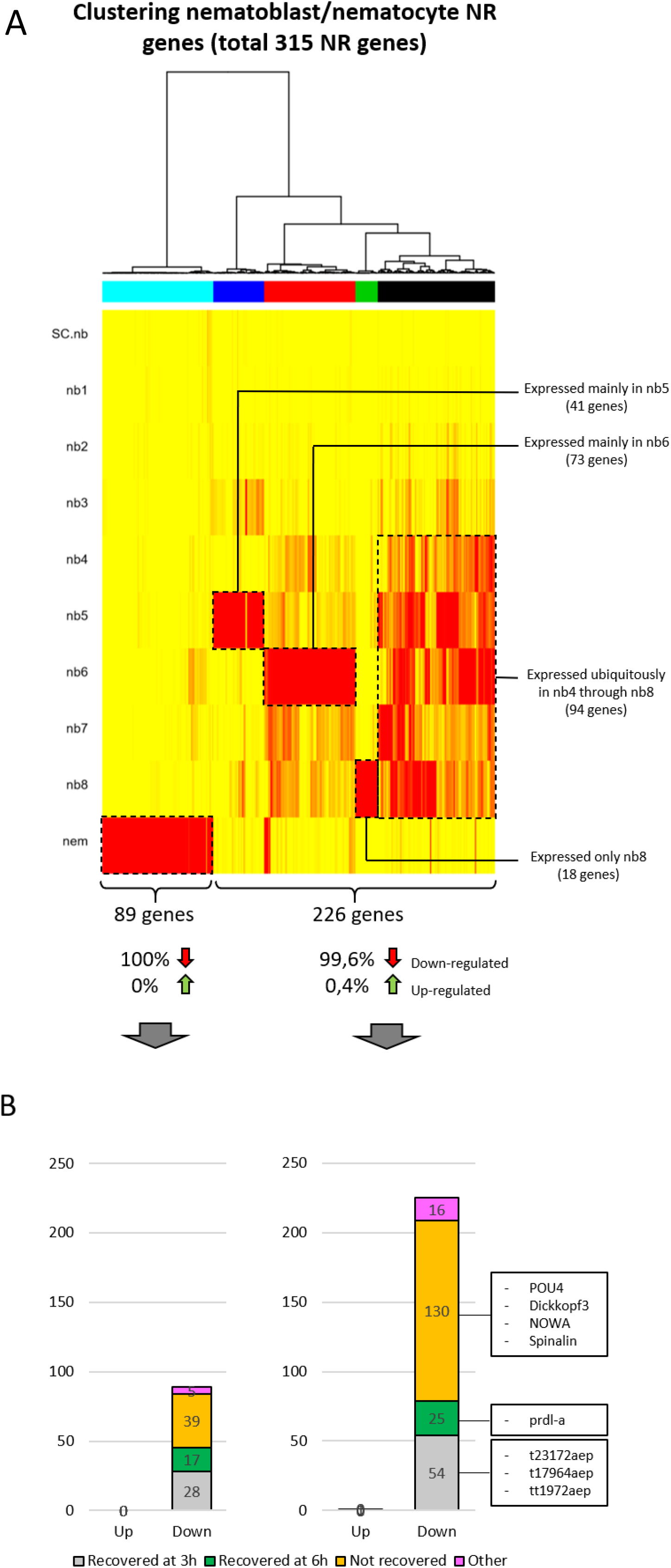
Hierarchical clustering of NR-genes expressed in the nematocyte lineage. A) NR-genes expressed in cells of the nematocyte lineage were clustered separately to reveal their expression in the differentiation states of nematogenesis. This revealed a set of genes only expressed in mature nematocytes (cyan cluster), genes mainly expressed in cell state nb5 (blue), genes mainly expressed in nb6 (red), genes mainly expressed in nb8 (green) and genes expressed ubiquitously in stages nb4 through nb8 (black). Almost all of these genes were down-regulated upon DAPT treatment. B) The majority of both mature nematocyte genes and nematoblast genes did not recover their expression 6 hours after DAPT removal. This includes POU4, Dickkopf3, NOWA and Spinalin.

To identify the point in the trajectories in which differentiating nematoblasts transition from proliferating to post-mitotic nematoblasts, we looked at the expression profiles of two genes that mark proliferating nematoblasts: (1) proliferating cell nuclear antigen *PCNA* (t10355aep) and (2) the Zn-finger transcription factor gene zic/odd-paired homolog *Hydra-zic (Hyzic)* (t13359aep; (Lindgens et al., 2004)). *PCNA* expression is seen in states SC.nb, nb1 and nb2 classifying them as proliferating nematoblasts (Fig. 5A). Nb1 and nb2 express HyZic, confirming that HyZic expression is restricted to proliferative nematoblast states as had been shown before (Lindgens et al., 2004). The absence of PCNA-expression in cell states nb3 and nb4 suggests that these are the earliest post-mitotic nematoblasts producing the nematocyst spine and inner wall protein spinalin (Käsbauer et al., 2007; Koch et al., 1998) and *spinalin* expression is clearly seen in these cells (Fig. 5A). Expression of the early differentiation marker genes *NOWA* and *CnASH* become detectable in differentiation states nb5 through nb8 when nematocyst capsules are formed (Fig. 5A and 5C).

**Figure 5.**
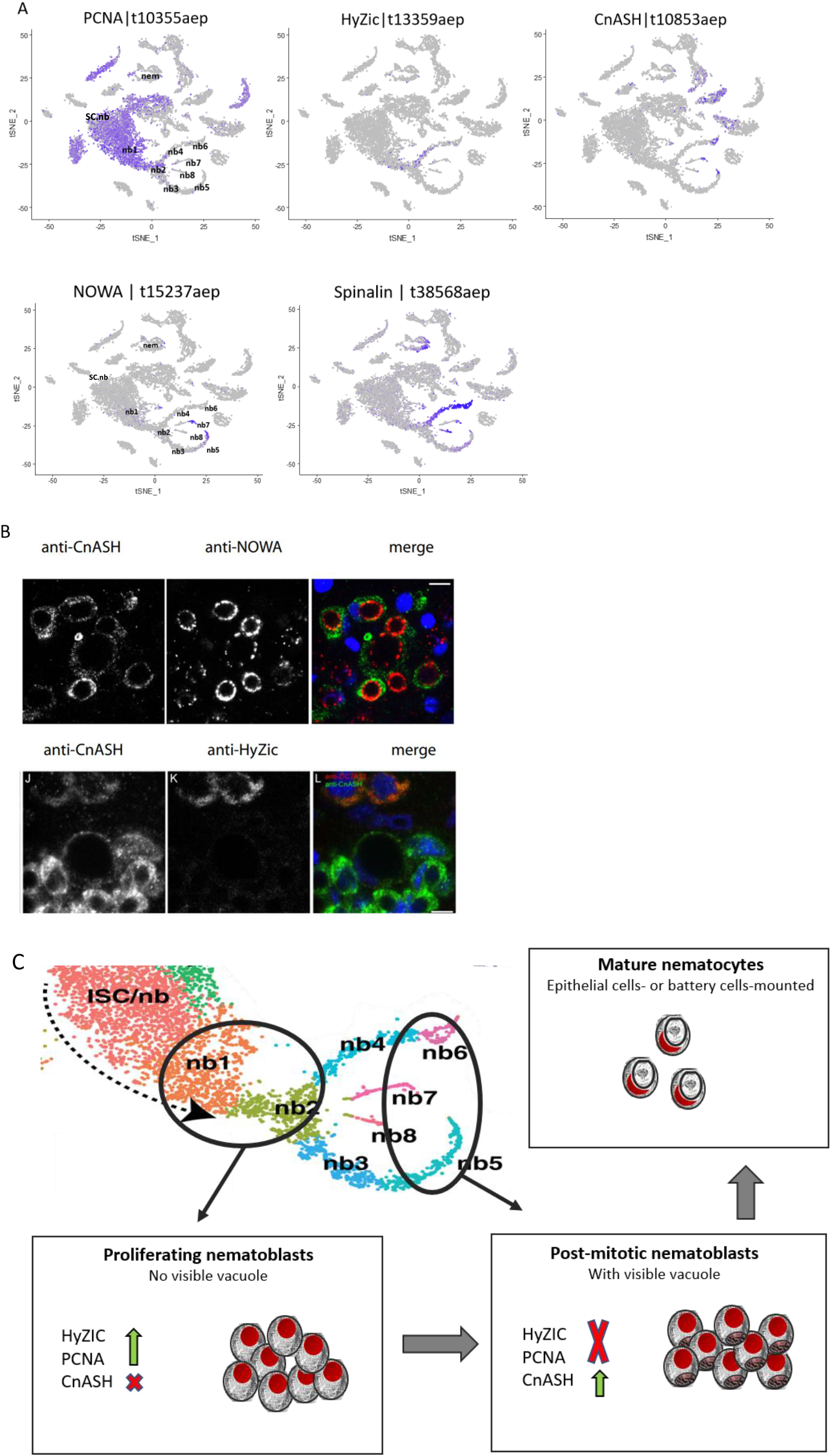
Homeostatic expression of nematoblast marker genes and proteins. A) t-SNE representation showing the interstitial cell state expression of *HyZic, PCNA, CnASH, NOWA* and *spinalin*. Cluster labels are provided for cell states of the nematoblast lineage according to (Siebert et al. 2019) nb: nematoblast, nem: nematocyte, SC: stemcell. Blue dots indicate cells expressing the respective genes. *PCNA* is expressed in proliferating cells - nematoblast cell states nb1 and nb2. *HyZIC* is mainly expressed in nb2. This is in accordance with previously published work indicating *HyZIC* expression in proliferating nematoblasts. *CnASH* is expressed in nb4, nb5, 6, 7, 8, representing post-mitotic nematoblasts lacking PCNA expression. This is in complete agreement with previous work (Lindgens et al 2004). *NOWA* encoding an outer capsule protein, is expressed in nb5 and nb7, *spinalin* encoding a protein occurring inside the capsule, is expressed in nb 5, 6, 7 and 8 and in mature nematocytes (nem), all representing post-mitotic nematoblast stages. B) Laser confocal microscopic sections of co-immunofluorescence staining with anti-HyZIC, anti-CnASH and anti-NOWA antibodies, in merged images DNA stain DAPI (blue), CnASH (green), HyZIC (red), NOWA (red). Anti-NOWA antibody delineates capsules (upper panel, middle image and red in merged). Co-staining with anti-CnASH antibody indicates signal in cytoplasm of capsule containing cells (upper panel, left hand image and green in merged). Capsule containing CnASH positive cells (lower panel, left hand side and merged image gren) are not stained with anti-HyZIC antibody (lower panel, middle image and merged image red). *Scale bars 20 μm.* C) Schematic summary of gene expression in the nematoblast lineage indicating a differentiation pathway from interstitial stem cell precursors (ISC/nb) via proliferating PCNA and HyZIC expressing amplifying nematoblast precursors (nb1, nb2) via post-mitotic nematoblasts not expressing PCNA (nb3, nb4) to capsule forming CnASH expressing nematoblasts (nb5, 6, 7, 8).

As further evidence that *HyZic* and *CnASH* mark mitotic and post-mitotic stages of nematogenesis respectively, using immunofluorescence we show that CnASH protein is detected in the cytoplasm of nematoblasts that contain vacuoles, which were visualised with anti-NOWA antibody (Engel et al., 2002). By contrast, HyZic protein was detected in the cytoplasm of nematoblasts without visible vacuoles and not found in CnASH positive nematoblasts (Fig. 5B).

Of the 315 NR-genes that are expressed in nematoblasts or nematocytes, 314 were down-regulated upon Notch-inhibition (Fig. 4A). These down-regulated genes include many genes expressed in developing nematocytes such as *POU4* (t11335aep), *Prdl-a* (t21636aep; (Gauchat et al., 2004), *HyDickkopf 3* (t20111aep; similar to *HyDKK3* (Fedders et al., 2004), *CnASH* (t10853aep, (Grens et al., 1995), Fig. 4B and 5B), *NOWA* (t15237aep; (Engel et al., 2002)) and *Spinalin* (t38568aep; (Koch et al., 1998), this gene has now 3 NCBI entries and encodes a longer protein as initially described, see Fig. S3);(Fig. 4B). Using double in situ hybridization to detect *POU4* and *HyZic* transcripts, we found mutually exclusive expression of these two genes in nematoblasts, which demonstrates that *POU4* is expressed in post-mitotic nematoblasts (Fig. 6A). Using in situ hybridization, we also confirmed that HyZic-positive nematoblasts were not affected by DAPT-treatment, whereas CnASH- and POU4-positive nematoblasts were lost (Fig. 6B and C).

**Figure 6.**
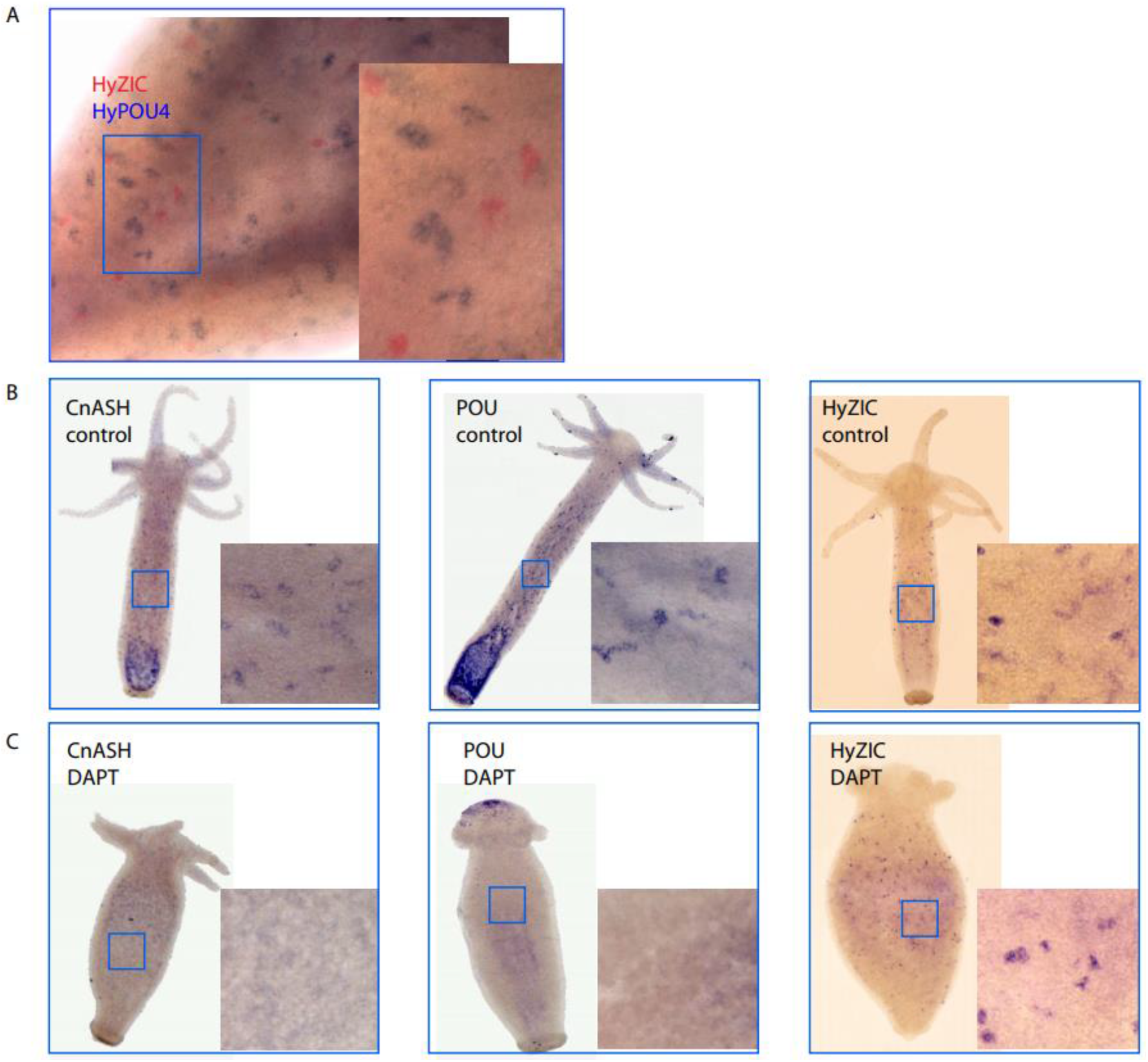
In situ hybridization for nematocyte marker genes. A) Double in situ-hybridisation for expression of *HyZIC (red signals)* and *HyPOU (blue signals).* B) Whole mount in situ hybridization for *HyZic, CnASH and POU4* in *Hydra* polyps treated for 48 hrs with 1 % DMSO for control (-DAPT). C) Whole mount *in situ hybridization for HyZic, CnASH* and *POU4* in *Hydra* polyps treated for 48 hrs with DAPT. Scale bars whole animal pictures 200μm, zoomed pictures 40μm.

More than 50% of the nematoblast-specific NR-genes remained down-regulated and did not recover their normal expression level within 6 hrs after the Notch-inhibitor was removed (Fig. 4B). This suggests that down regulation of nematogenesis genes reflects the loss of cell states and is mainly an indirect effect of the block in this process caused by Notch inhibition. By contrast, some nematocyte specific putative transcription factors did recover their expression levels quickly after DAPT removal. These included a possible class I member of the HMG-Box superfamily, similar to SOXB3 from *Hydractinia Echinata* (t23172aep, XP_012555836.1, alignments Fig. S3), a protein with a C-terminal bZIP-Jun-domain (t17964aep, alignment Fig. S3) and a predicted forkhead box protein I1c-like (t1972aep, alignment in Fig. S3) (Fig. 4B). Given the rapid recovery of their expression after DAPT removal, these genes may be directly targeted by Notch signalling and possibly play a major role in driving nematogenesis.

#### 2.2 Epithelial expression of Notch regulated genes

About 25% of NR-genes had enriched expression in epithelial cells (Fig. 3, black and grey cluster) while the remaining non-nematoblast NR-genes showed either sporadic or ubiquitous expression (Fig. 3, cyan and green clusters).

Since previous Notch inhibition studies demonstrated severe malformations of the *Hydra* head structure (Münder et al., 2013), we aimed to elucidate the effect of DAPT treatment on epithelial body column cells and their derivatives (e.g. specialized head and foot cells). Hierarchical clustering using single-cell data for epithelial cells revealed genes that were expressed 1) in all endodermal and ectodermal epithelial cell types along the oral-aboral axis (Fig. 7, grey cluster), 2) mainly in ectodermal epithelial cells (Fig. 7, cyan cluster) and 3) mainly in endodermal epithelial cells (Fig. 7, green cluster). The majority of these epithelial genes from the grey, cyan and green clusters were up-regulated in response to DAPT treatment (Fig. 7). These included 36 genes associated with ER, Golgi- and endosomal proteins, for example involved in glycosylation like the oligosaccharyl transferase DAD1(t14233aep|DAD1), a negative regulator of cell death (Roboti and High, 2012). Some were involved in redox-regulation and unfolded protein response and some were chaperones (Tables S1, S2). Moreover, membrane proteins, including twelve G-protein coupled receptors, caspase D (t7281aep; (Lasi et al., 2010) and the homolog of the ubiquitin-ligase and Notch-modulator mind bomb were also up-regulated (t3105aep). Thus, many of the up-regulated epithelial genes seem to be involved in stress responses to DAPT-treatment. In contrast, *Sp5*, which is involved in *Hydra* head patterning (t29291aep; (Vogg et al., 2019) was down-regulated (Fig. 8A).

**Figure 7.**
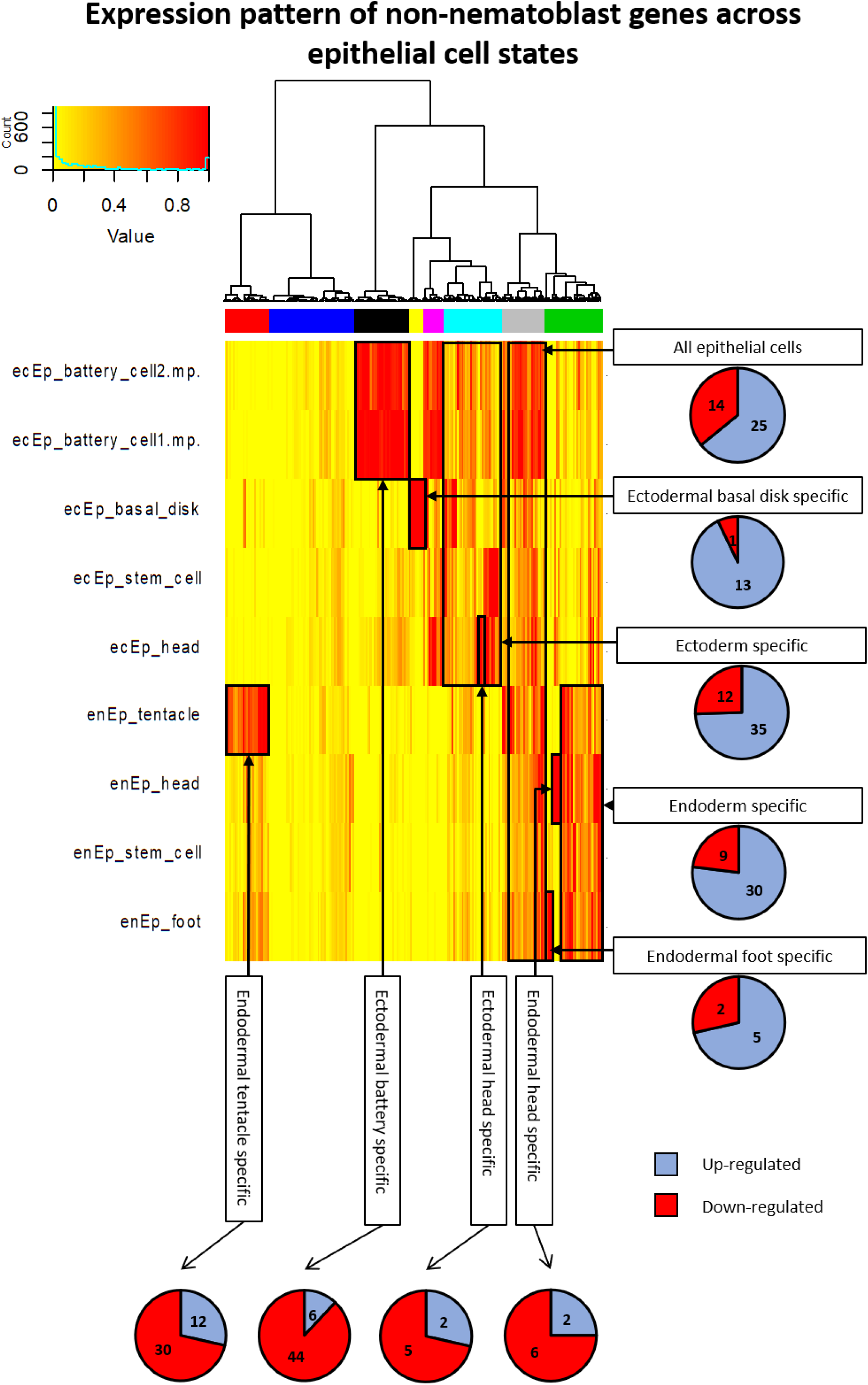
NR gene subset with expression in epithelial cells. Non-nematoblast NR genes were clustered separately to determine their expression in epithelial cell states. This revealed sets of genes that are most strongly expressed in endodermal tentacle cells (red cluster), ectodermal basal disc cells (yellow), ectodermal battery cells (black), body column ectoderm cells (cyan), body column endoderm cells (green) and all epithelial cells (grey). The analysis also revealed smaller gene sets expressed in endodermal foot cells, endodermal head cells or ectodermal head cells. Tentacle, battery and head specific genes were mainly down-regulated upon DAPT treatment whereas the genes in the remaining clusters were mainly up-regulated.

**Figure 8.**
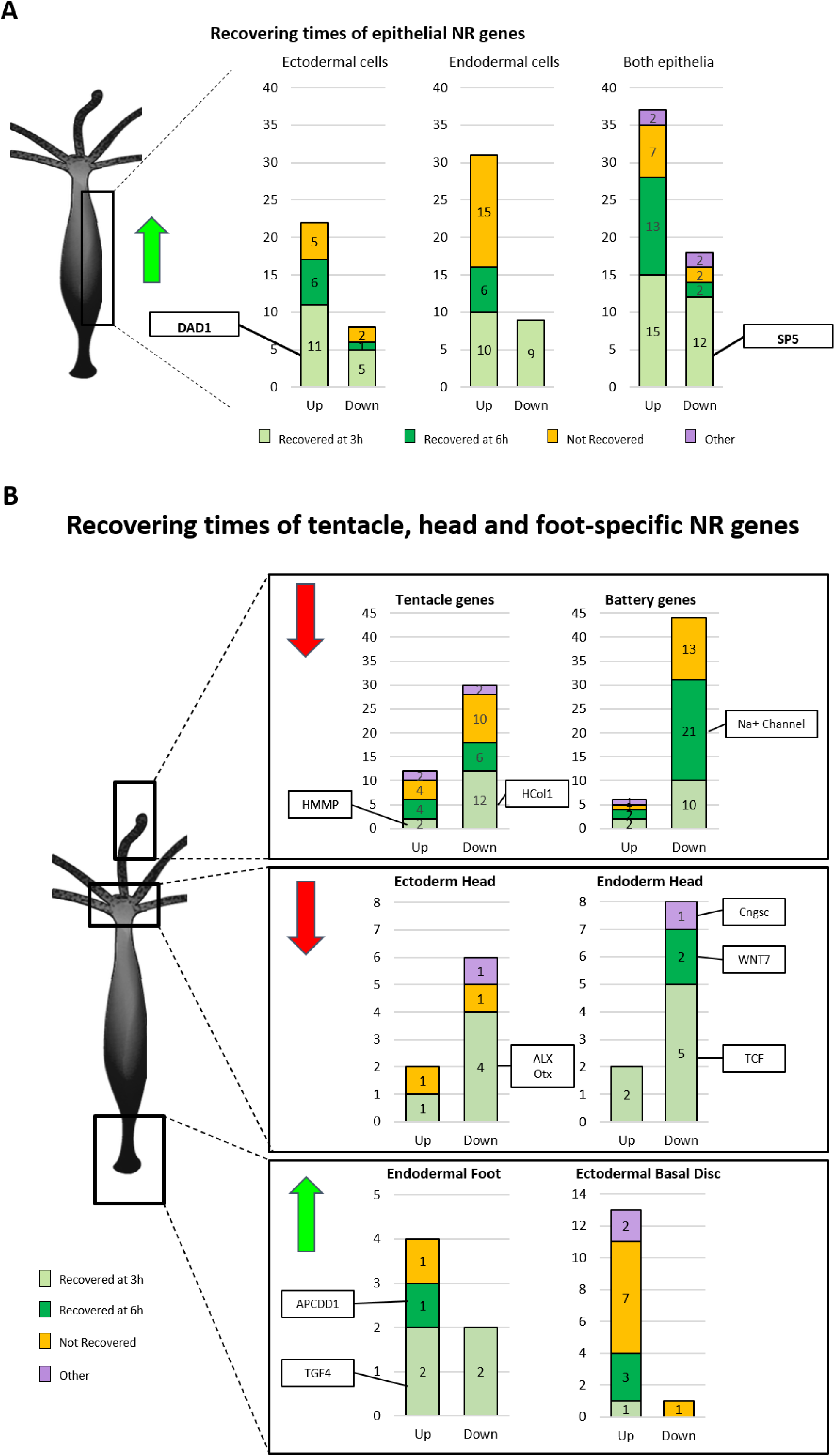
Recovery time of NR-genes. A) The majority of epithelial NR-genes were up-regulated upon DAPT treatment. 50% of the up-regulated ectodermal-specific NR-genes recover expression within the first 3 hours post DAPT removal and include the apoptosis-involved gene DAD1. SP5 on the other hand, which is expressed in both epithelia, is down-regulated and recovers expression also within the first 3 hours. B) Head-specific NR-genes, including tentacle, battery and ectodermal and endodermal head genes, are mostly down-regulated upon DAPT treatment. In contrast, foot-specific genes, including endodermal foot genes and basal disc genes are mostly up-regulated. Many of these genes play a predominant role in patterning.

Two sets of genes comprised tentacle genes expressed in endodermal tentacle cells (Fig. 7, red cluster) and in ectodermal battery cells (Fig. 7, black cluster). In both sets, the majority of genes were down-regulated upon Notch-inhibition (88% of battery cell genes and 71% of endodermal tentacle genes). These included a gene encoding a Na+ channel in battery cells (t18364aep; (Golubovic et al., 2007)) and the collagen gene Hcol1 (t14477; (Deutzmann et al., 2000)) in endodermal tentacle cells (Fig. 8B). The endodermal matrix metalloprotease gene HMMP was upregulated (t16424aep; (Leontovich et al., 2000)). Both extracellular matrix genes, HMMP and Hcol1, recovered their expression levels within 3h (Fig. 8B).

Furthermore, small sets of NR-genes were specifically expressed in 1) ectodermal head cells, 2) endodermal head cells, and 3) endodermal foot cells (Fig. 7, included in cyan and green clusters). Another NR gene cluster is expressed in ectodermal basal disc cells (Fig. 7 yellow cluster). These expression patterns also surfaced on tSNE plots after NMF analysis (see supplementary metagenes, “Ectoderm specific - battery” A, “Ectoderm specific - basal disk” A, “Endoderm specific - tentacle cells” A, “Ecto.Endo – foot” A).

The NR-genes expressed in endodermal and ectodermal head cells were largely down-regulated and several of these have known functions in head patterning. Of note, *HyALX* (t16456aep) (Smith et al., 2000) is expressed at tentacle boundaries and previous work demonstrated that HvNotch is needed to maintain this expression pattern (Münder et al., 2013). Furthermore, several potential head organizer genes including *Wnt7* (t28874aep; (Lengfeld et al., 2009)), the transcription factor gene *TCF* (t11826aep; (Hobmayer et al., 2000))), an Otx-related homeodomain protein (t33622aep), an FGF-homolog (t8338aep; annotation confirmed by Monika Hassel, Marburg, Germany) and *CnGSC* (t1216aep, (Broun et al., 1999)), were among this down-regulated set of head-specific genes (Fig. 8B). Of those, *HyALX*, *CNGSC, Wnt 7, FGF* and *HyTCF* recover their normal expression levels within 3 hrs making these genes candidates for direct targets of Notch signalling. The organizer gene *CNGSC* was also downregulated and recovered expression after 3 hrs. However, it was then downregulated again at 6 hrs. This unusual expression behaviour might indicate the presence of an inhibitory feedback mechanism responding to Notch signalling.

By contrast, the NR-genes that are specifically expressed in endodermal foot cells and in ectodermal basal disc cells are largely up-regulated in response to Notch inhibition. This includes *TGF-4* (t25624aep; (Watanabe et al., 2014)) and a predicted secreted Wnt-inhibitor APCDD1 (t11061aep). Thus, Notch inhibition by DAPT resulted in reciprocal regulation of foot and head genes in *Hydra*, with genes normally expressed at the oral end being down-regulated and genes normally expressed at the aboral end being up-regulated (Fig. 8B).

### 3 Promoter analysis of NR-genes reveals likely direct targets of Notch signalling

The differential gene expression analysis revealed sets of genes that showed shared behaviour after Notch-inhibition and re-activation by DAPT removal. This suggests shared regulation and we performed a motif enrichment analysis to uncover respective regulatory elements in genes with similar expression dynamics. This analysis was done for the following gene sets: 1) down-regulated only at 0h, 2) down-regulated at 0 and 3h, 3) down-regulated at 0, 3 and 6h, 4) up-regulated only at 0h, 5) up-regulated at 0 and 3h, and 6) up-regulated at 0,3, and 6h. Regions of open chromatin, as identified by previously published ATAC-seq data, within 5kb upstream of each gene were considered in the enrichment analysis (see Fig. 9 and methods for details) (Siebert et al., 2019).

**Figure 9.**
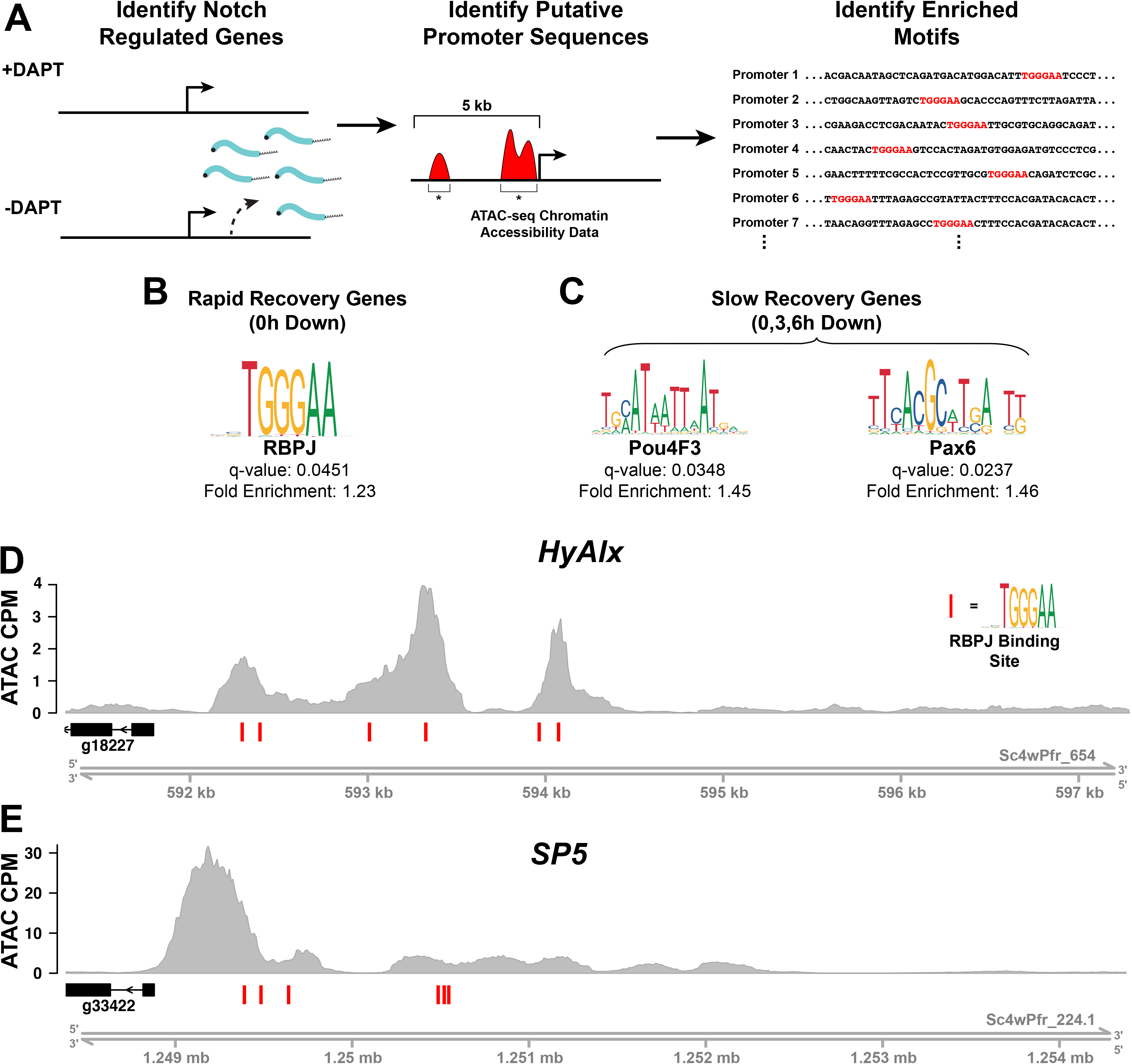
Motif enrichment analysis of NR gene promoter regions. A) Workflow of motif enrichment analysis. Putative promoter regions were identified using a previously published ATAC-seq dataset generated using whole wild-type *Hydra* (Siebert et al., 2019). NR gene promoter regions were defined as ATAC-seq peaks that fell within 5kb upstream of the transcription start site of an NR gene. Using HOMER, NR gene promoters were compared against control peaks that were not associated with NR genes to identify significantly enriched (FDR ≤ 0.05) transcription factor binding motifs. B) The Notch/RBPJ binding motif was significantly enriched in the putative promoters of genes that were downregulated upon DAPT treatment and recovered rapidly following inhibitor removal. C) Pou and Pax transcription factor binding motifs were significantly enriched in the putative promoters of genes that were downregulated upon DAPT treatment and did not recover their expression over the course of the RNA-seq experiment. Plots of normalized ATAC-seq read density in the 5kb upstream of D) HyAlx and E) SP5 demonstrate the presence of predicted RBPJ binding sites in the putative promoters of NR genes. Red bars indicate predicted instances of Notch binding motifs.

The group of genes that were downregulated in response to DAPT treatment and then recovered normal expression by 3 hrs are the best candidates for direct targets of Notch signalling. If genes are direct targets of Notch signalling, we would expect to find RBPJ-binding sites (Kopan and Ilagan, 2009). In line with our prediction, the RBPJ-motif (Bailey and Posakony, 1995) was enriched in NR-genes of this group (Table S3, Fig. 9B). Among the 21 genes with RBPJ binding sites in their regulatory region, *HyAlx* (t16456aep) and *Sp5* (t29291aep), with six RBPJ motifs, are the top candidates for direct targets of Notch signalling (Fig. 9D, E, Table S5). We also identified the transcription factors pituitary homeobox 1-like (specifically expressed in head cells of the endoderm, t5275aep) and a homeobox protein of the OTX-family (t33622aep). Putative RBPJ-motifs are additionally present in genes encoding potential membrane or extracellular proteins, including a foot specific secreted frizzled-related protein, a potential regulator of Wnt-signalling (t15331aep, annotation provided by Bert Hobmayer, Innsbruck, Austria, personal communication).

In addition to the RBPJ binding site, we found enrichment of further transcription factor binding motifs belonging to 10 transcription factor families. Homeobox transcription factors were the most abundant motifs identified and several different HMG, Forkhead and bHLH motifs were also found (Table S3). Interestingly, this corresponds with the downregulation of transcription factors that potentially bind to these domains, e.g. HyHES (bHLH), Jun (bZIP), FoxP1 (Forkhead), HyAlx, (homeobox, t16456aep), OTX-related (homeobox, t33622aep) and PITX-related factors (homeobox, t5275aep) and three SOX-related proteins (HMG-boxes, t23837aep, t23172aep, t5528aep) (see Tables S1, S2, S3, S4 and alignment and phylogeny in Fig. S3). Therefore, these data reveal the possible components of a gene regulatory network influenced by Notch signalling.

For the group of genes down-regulated at all three time points, six enriched motifs were found, most notably POU- and PAX-binding motifs (Fig. 9C, Table S3). The *POU*-gene has previously been implicated in nematocyte differentiation and was found enriched in genes expressed at late stages of nematogenesis (Siebert et al., 2019). *HyPOU4TF-2 like* was down-regulated by Notch-inhibition at 0 and 3 hrs (Table S4). The three predicted *Hydra-Pax*-genes (t9974aep, t6559aep and t11467aep, see Table S1 and Fig. S3) were not amongst the NR-genes.

In the group of genes that were up-regulated at 0h, but recovered by 3h, only the IRF9 binding motif (interferon regulatory factor) was found enriched. For genes that were up-regulated at 0 and 3h, but recovered expression at 6 hours, many bZIP-factor binding motifs were enriched.

## Discussion

Inhibiting Notch-signalling induces a block in nematocyte differentiation and disrupts head patterning in *Hydra* (Münder et al., 2013). In this study we have identified Notch-regulated (NR) genes by analysing RNAseq-data obtained at different timepoints after treatment of *Hydra* polyps with DAPT. Exploration of the *Hydra* single-cell gene expression atlas (Siebert et al., 2019) revealed sets of genes that were expressed in cell states consistent with observed inhibition phenotypes. Moreover, in many NR-genes we detected binding sites for DNA-binding protein RBPJ - the principle effector of Notch signalling.

Unexpectedly, we detected up-regulation of genes encoding heat shock proteins and proteins involved in apoptosis (Table S2), hinting at a stress response of the animals to the treatment. Promoter analysis of those up-regulated genes revealed enrichment of a Trp-cluster motif (IRF9, Table S3). This motif is targeted by the interferon regulatory factor as part of a stress and anti-viral defence pathway in mammals (Jefferies, 2019). Its occurrence and function in *Hydra* genes should be elucidated in the future.

Strikingly, almost half of the Notch responsive genes were expressed in cells of the nematocyte lineage, 99% of those were down-regulated and expressed in post-mitotic nematoblast stages. This reflected Notch-regulation of a gene module specific to post-mitotic nematoblasts that are in the process of capsule formation. In this group we find most genes that have been previously shown to encode structural capsule proteins, such as minicollagens (Engel et al., 2001), spinalin (Koch et al., 1998), NOWA (Engel et al., 2002), nematogalectin (Zhang et al., 2019), N-col15 (Adamczyk et al., 2008), nematocilin (Hwang et al., 2008) and others. Moreover, we find transcription factors like HyPOU (Siebert et al., 2019) and CnASH (Grens et al., 1995). This is consistent with Notch-inhibition blocking differentiation and the initiation of a transcription program for capsule formation. We propose that HyPOU is involved in executing this program but that it is not a direct Notch-target since expression is not re-established within 3 hrs. In accordance with this hypothesis, motif enrichment analysis identified POU4F-DNA-binding motifs in the putative regulatory region of genes that did not recover from DAPT within 6 hrs. Similarly, *CnASH* expression does not recover within 6 hrs, confirming it as an indirect or secondary target (Fig. S1). Genes that are expressed in nematoblast precursors, including *HyZIC* (Lindgens et al., 2004) and the *HydraPax-2A* homolog (t9974aep) were not found affected by DAPT.

Genes that are potentially directly targeted by the Notch intracellular domain (NICD) not only should recover their expression level quickly when DAPT-treatment is removed and NICD is allowed to enter the nucleus but should also contain RBDJ-sites in their promoter regions. Such binding motifs have been detected in a number of nematocyte specific genes with unknown function (Table S5). These genes do not encode transcription factors suggesting that NICD directly activates the nematocyte differentiation gene complex. Future studies will reveal their role during nematoblast differentiation and also whether they can account for the missing differentiation cue that is directly blocked with DAPT.

However, as an alternative explanation, failure to carry out the nematoblast differentiation program in our experiments could be caused by missing patterning signals from the *Hydra* head. This hypothesis is suggested because of the strong head phenotypes that we had previously described after DAPT-inhibition. The first observable phenotype after 48 hrs of Notch-inhibition was a substantial shortening of the tentacles. Moreover, transplantation experiments with GFP-labelled body column tissue indicated that during the time of Notch-inhibition cells did not cross the boundary between body column and tentacles (Münder et al., 2013). A “neck” like structure appeared underneath the tentacle zone, where cells have ceased proliferating, but also did not differentiate into battery cells. In this study, we reveal a cluster of down-regulated head-specific genes amongst the genes that are dysregulated in response to Notch inhibition.

Of particular interest is the *aristaless*-related gene *HyALX,* which has six potential RBPJ-sites in its putative regulatory region. Our study strongly suggests that Notch-signalling directly activates *HyALX* expression. *HyALX* has previously been proposed to instruct the specification of tentacle tissue (Smith et al., 2000) and we suggest that *HyAlx* could play this key role in directing tentacle fate by activating genes with homeobox transcription factor binding motifs. In support of this, we found the homeobox motif enriched in the NR-genes down-regulated by DAPT that recover their expression quickly after DAPT removal. *HyALX* is expressed in evenly spaced rings at the body column-tentacle boundaries. After release of DAPT-inhibition it rapidly recovers expression levels, yet it does not recover a regular expression pattern but becomes expressed in irregular rings, in the extreme case in only one ring surrounding the whole animal (Münder et al., 2013). Assuming that NICD acts as a direct activator of *HyALX*, this indicates that Notch-signalling is resumed in the wrong places. Therefore, a feedback mechanism can be suggested, where the Notch-signalling pattern depends on the head organizer, which in turn is co-instructed by Notch-signalling.

The potential *Hydra* head organizer gene *CnGSC* (Broun et al., 1999) is down-regulated by Notch-inhibition and does not recover its activity after 6 hrs. Furthermore, *Wnt 7, TCF* and *Sp5,* genes implicated in the canonical Wnt-signalling pathway (Broun et al., 1999; Lengfeld et al., 2009; Vogg et al., 2019) are also down-regulated. In contrast, we found a small cluster with foot- and peduncle genes that were up-regulated, including the BMP-pathway gene *TGF-4* (Watanabe et al., 2014). Together, these data may guide the uncovering of molecular pathways responsible for the irregularly shaped heads that develop in polyps after a 48 hrs period of DAPT treatment (Münder et al., 2013). They also confirm a role of Notch-signalling in establishing and maintaining the *Hydra* head organizer, which was previously discovered in transplantation experiments, where the organizer capacity of regenerating *Hydra* head tissue had been inhibited by DAPT (Münder et al., 2013).

## Conclusion

This study suggests target genes of Notch-signalling in *Hydra*, and provides a resource for the investigation of molecular mechanisms by which HvNotch affects patterning, maintenance of the head organizer and post-mitotic nematocyte differentiation. The expression of the only direct HvNotch target gene, for which experimental evidence is available, *HyHes*, was also found amongst NR-genes, which quickly recovered original expression levels after DAPT removal. We have identified *HyAlx* and *Sp5* as prime candidates for further direct HvNotch-targets involved in head patterning due to their quick recovery after DAPT relief and the presence of RBPJ-sites in their promoter regions. An obvious candidate for a direct HvNotch-target gene, for instance a transcription factor gene, expressed in early differentiating nematoblast states and quickly recovering from DAPT treatment has not surfaced. Thus, the impact of HvNotch on regulation of this differentiation step may be conveyed through signals from epithelial cells, or by inducing expression of genes encoding proteins other than transcription factors, for instance genes that are required to form the post-Golgi vacuole. It has also to be considered that many genes with yet unknown functions are amongst potential direct Notch-targets in nematoblasts.

## Methods

### Hydra culture

Animals of the strain *Hydra vulgaris* (Basel) were grown in *Hydra* medium (0.1 mM KCl, 1 mM NaCl, 0.1 mM MgSO_4_, 1 mM Tris and 1 mM CaCl_2_) at 18°C and fed regularly with freshly hatched *Artemia nauplii.*

### DAPT-treatment

Regularly fed animals were starved for 24 hours and incubated in either 20μM DAPT/1% DMSO in *Hydra* medium or only 1% DMSO in *Hydra* medium (control sample) for 48 hours. DAPT and DMSO were renewed every 12 hours. Animals were collected and total RNA was isolated at three different time points: directly at the end of 48 hours (0h), 3 hours after DAPT removal (3h) and 6 hours after DAPT removal (6h). After 48 hours incubation, DAPT was removed and replaced with 1% DMSO in *Hydra* medium for the samples 3h and 6h. About 25 animals were collected per sample. Three or six biological replicates were analyzed using qPCR or RNA-seq respectively.

### qPCR

Total RNA was extracted from whole animals using the RNeasy Mini kit Plus (Qiagen) according to the manufacturer’s protocol. RNA quality and quantity were assessed using an Agilent Bioanalyzer. RNA with a RIN value of at least 8 was used for cDNA synthesis using the iScript cDNA synthesis kit (BioRad) according to manufacturer’s protocol.

Primers for qPCR were tested before to ensure they amplified the correct fragment from cDNA. Gene specific primer pairs which yielded one melt peak and a linear standard curve were used for qPCR quantification.

cDNA was diluted 1:25 to ensure the used concentration was within the standard curve of the primers. Real-time PCR with SYBR green detection was performed using an CFX96 Touch Real-Time PCR Detection System (Biorad). Each measurement was performed in three technical replicates. The genes *RPL13, SDH, EF1α, GAPDH* and *Actin* served as housekeeping genes and were used for normalization. The samples were analyzed on a 96 well plate in a CFX96 Touch Real-Time PCR Detection System from BioRadRelative expression was calculated as 2^-(*dCt*(test sample) - *dCt*(reference sample)). The relative expression of the control samples (DMSO treated animals) was set to 1 and the relative expression of the other samples was normalized according to the control.

### Immunohistochemistry

Animals were briefly (1-2 min) relaxed in 2% Urethane/*Hydra* medium and fixed immediately after in 2% Paraformaldehyde/*Hydra* medium for (1h). Animals were washed with PBS, permeabilized with 0.5% Triton-X-100/PBS (15 min) and blocked with 0.1% Triton-X-100/1% BSA/PBS (20 min). Primary antibodies were applied overnight at 4 °C. After a PBS-wash, animals were incubated with secondary antibodies (2h), washed again with PBS, counterstained for DNA with DAPI (Sigma, 1 μg/ml) and mounted on slides in Vectashield mounting medium (Axxora).

### Whole-mount in situ hybridisation

RNA in situ hybridization experiments were carried out as previously described (Grens et al., 1995) using digoxigenin labelled RNA probes (Roche) and substrates NBT/BCIP or BM Purple (Roche).

### RNA-seq

RNA-seq libraries were prepared for six biological replicates for each experimental condition. cDNA libraries were synthesized from total RNA using the strand specific SENSE mRNA-Seq Library Prep Kit V2 for Illumina (Lexogen) and the Purification Module with Magnetic Beads (Lexogen). The samples were multiplexed and sequenced on three lanes on Illumina Hiseq2000 with a 100bp paired end sequencing strategy. Downstream analyses were performed using the Galaxy platform and within R (RStudio Team (2016); version 1.1.463) (Rcode provided in supplement). Illumina adapters and polyA sequences were trimmed and splice leader sequences (Stover and Steele, 2001) were removed from both forward and reverse reads. The tool “fastqfilter” was used to ensure the paired nature of the filtered dataset, to filter out reads with a quality score lower than 20 and to exclude reads with a read length shorter than 30b. Reads that contained Ns were also removed from the dataset.

### De novo transcriptome assembly

All forward and all reverse reads from all sequencing libraries were concatenated. The two resulting files were then used as input to the Trinity (version 2.8.4, (Grabherr et al., 2011)) *de novo* transcriptome assembler. The assembly was run with the following parameters: - strand-specific library, *in silico* normalisation, -min_contig_length 300, - min_kmer_cov 1, no genome guided mode and no Jaccard Clip options. The resulting reference transcriptome resulted in 62.419 transcripts (43.481 genes) with an average transcript length of 1008b. Transcripts that belonged to the same gene were joined to form SuperTranscripts (tool “Generate SuperTranscripts from a Trinity assembly”, Galaxy version 2.8.4, Davidson et al., 2017), which were then used for local Blast search. These were treated as genes models in downstream analyses (see below).

### Mapping reads to transcriptome

The processed reads of the 36 RNA-seq libraries (timepoints 0, 3 and 6 h, DAPT and control samples, six replicates) were separately mapped to the novo assembled transcriptome reference, within the Galaxy platform. The reads were mapped as strand-specific and with a maximum insert size of 800. RSEM (Li and Dewey, 2011) - with Bowtie2 as alignment method-was used as the abundance estimation method. The transcript alignment files of all samples and the gene_to_transcript_map were used as input to generate an expression matrix for all 43.481 assembled genes.

### Differential expression analysis

The raw counts were used as input for differential expression (DE) analysis by DESeq2 (version 1.18.1). Genes that were not detected in all 36 samples were excluded from this analysis. DE analysis was performed for each time point separately by comparing the DAPT treatment replicates with those from the control animals (0h DAPT vs. 0h DMSO, 3h DAPT vs. 3h DMSO and 6h DAPT vs. 6h DMSO). Differentially expressed genes at time point 0h were selected according to their P-adjusted value (Padj(FDR) < 0,01). We refer to this gene set as Notch regulated genes (NR-genes). For each of these NR-genes, we investigated whether differential expression was also identified at time points 3h and 6h, thereby applying the same cutoff for differential expression (Padj(FDR) < 0,01).

### Blast search

Several blast searches were performed to annotate NR-genes. The NCBI *Hydra vulgaris* protein database (on 2020.02.24) was interrogated using blastx. Sequences with no blast hit or a blast hit with an E-value > 10e-100 were blasted manually. Three types of manual blast searches were performed, NCBI blastn and blastx and NCBI smartBLAST (supplementary list of NR-genes). Sequences for which no blast hits were found in either blast search were denoted with “no blast hit”. This was also the case for sequences, for which a blast hit was found but with an E-value > 10e-20. For the genes that were blasted manually, the NCBI description and accession number was replaced by those of the blast hit with the highest E-value and query cover (for example if the manual blastn search yielded a better hit than the local blast to the NCBI protein database). The Pubmed accession number was added to the supplementary list for known Hydra genes. Uniprot was used to search for information about the function and the compartment of the identified sequences, these were denoted as “unclear” in case it was unclear or unknown. Multiple alignments were performed for genes with a similar TrinityID and genes with similar/same NCBI description.

### Cell state analysis

To make use of the available Hydra single cell data we first identified NR-genes within the single cell transcriptome reference using blastn (Siebert et al., 2019) (<u>**Transcriptome Shotgun Assembly project: GHHG01000000).**</u> Duplicated hits were removed by keeping the alignments with the highest blast score. Existing Seurat data objects were used to retrieve expression data and cell state annotations (Siebert et al., 2019) (Dryad https://doi.org/10.5061/dryad.v5r6077). For hierarchical clustering approaches average cluster expression was calculated for each cell state (Seurat_2.3.4::AverageExpression). Seurat objectes were then subsetted to the Notch DR-gene set and expression was scaled from 0 to 1. Hierarchical clustering was performed using functions stats::dist(“euclidian”) and stats::hclust(“ward.D”). A heatmap (gplots_3.0.0::heatmap.2) with the scaled average expression was generated.

### NMF analysis

Normalized expression information was extracted from the whole transcriptome Seurat object for each DE gene with an AEP reference and used for non-negative matrix factorization (NMF) analysis. This analysis was performed as described by Siebert and colleagues (Siebert et al., 2019).

### Motif enrichment analysis

To identify putative promoter regions of NR genes, we used a previously published ATAC-seq dataset generated from whole wild-type *Hydra* (Siebert et al., 2019) to locate regions of accessible chromatin (i.e. peaks) within 5 kb upstream of NR gene transcription start sites. We then grouped these NR promoter regions based on the expression dynamics of their putative target genes in our DAPT-treated RNA-seq time course. A total of six sets of NR genes were considered for downstream motif enrichment analyses: 1) genes that were downregulated but recovered by 3 hours post-treatment, 2) genes that were downregulated but recovered by 6 hours post-treatment, 3) genes that were downregulated and remained downregulated at 6 hours post-treatment, 4) genes that were upregulated but recovered by 3 hours post-treatment, 5) genes that were upregulated but recovered by 6 hours post-treatment, and 6) genes that were upregulated and remained upregulated at 6 hours post-treatment.

For our motif enrichment analysis, we used a curated set of known transcription factor binding motifs provided by the JASPAR database (Fornes et al., 2020). Specifically, we used position weight matrices from the non-redundant vertebrate, insect, nematode, and urochordate JASPAR datasets. JASPAR-formatted position weight matrices were converted to HOMER-formatted motifs using the HOMER parseJasparMatrix function. HOMER-formatted motifs require the specification of a score threshold that is used for identifying true motif hits in a query sequence. No such score threshold is included in JASPAR-formatted motifs, so we manually set the threshold to be 40% of the maximum possible score (i.e. the score that would be received by a sequence that perfectly matches the canonical binding sequence) for each motif.

We then used this custom set of HOMER motifs to identify transcription factor binding motifs that were significantly enriched in each of the six abovementioned NR peak sets. We did this by comparing the NR peak sets to non-NR peaks using a binomial enrichment test as implemented in the HOMER findMotifsGenome function. Motif enrichment results were then filtered using a false discovery rate threshold of ≤ 0.05.

We found that our raw HOMER results included numerous enriched motifs with highly similar sequences. To simplify these results, we sought to identify and remove redundant motifs from the results tables. To accomplish this, we first generated a matrix of pairwise similarity scores for all motifs in our custom motif set using the HOMER compareMotifs function. These similarity scores were then used to perform hierarchical clustering to identify groups of highly similar motifs. We then reduced the redundancy of our enrichment results by including only the most significantly enriched motif from each motif cluster in the final results table.

To identify putative RBPJ binding sites in NR promoter regions, we used the HOMER scanMotifGenomeWide function to find sequences that matched the RBPJ and Su(H) binding motifs (JASPAR matrix IDs MA1116.1 and MA0085.1 respectively). In addition, we also made use of a custom Su(H) motif based on a previously reported description of the Su(H) consensus binding site (Bailey and Posakony, 1995). The custom HOMER Su(H) motif was generated using the HOMER seq2profile function; the score threshold was set to be 40% of the maximum possible score.

Plots of ATAC-seq read density and predicted RBPJ binding sites were generated using the R Gviz package (Hahne and Ivanek, 2016). ATAC-seq reads from individual biological replicates were pooled before generating read density plots.

## Supporting information

Supplemental Table 1-NR-genes

supplemental Figure S2 metagenes

RCode

## Acknowledgements

The Galaxy platform of Blum’s group at the Gencentre Munich was used for read processing. This work was funded by DFG-grant BO1748-12-1.

---

**Figure S1**. *RT/qPCR measurement of HyHes and HyCnASH upon DAPT washout*. Animals were treated with 20 μM DAPT/1% DMSO or with 1% DMSO as a control for 48 hours. Thereafter, DAPT was removed and RT/ qPCR were performed at 0h, 3h and 6h. Control animals were treated with 1% DMSO. The relative expression was calculated according to the Cq values. The relative expression of the control animals was set to 1.

**Figure S2 Metagenes**. *Metagenes.* Non-negative matrix factorization (NMF) was performed in order to identify groups (metagenes) of genes with similar expression patterns. Normalized cell state expression information for the 666 NR-genes with an AEP reference was extracted from the whole-transcriptome dataset (Siebert et al., 2019) and used as input for the NMF analysis. This analysis yielded 23 Metagenes, which were divided into 10 groups, according to the cell types in which the metagenes were expressed. The overall expression of each metagene is displayed on the tSNE plots of all four datasets: whole-transcriptome, interstitial cell lineage, ectodermal cell lineage and endodermal cell lineage (Siebert et al., 2019).

**Figure S3**. Annotation of selected NR-genes by sequence comparison and phylogenetic analyses.

**Table S1**. *Full list of all 831 NR-genes*. “Trinity.ID” is the name that was given to the genes assembled by Trinity. These are the genes, that were used for the differential gene expression analysis. The SuperTranscript sequence of each gene is given in column “Sequence.SuperTranscript”. “AEP.ID” and “AEP.annotation” are the AEP references and their annotation as characterized by Siebert et al. (2019). “Gene.Model” corresponds to the published gene models at the Hydra 2.0 Web Portal (arusha.nhgri.nih.gov/hydra). “NA” means no AEP reference and/or gene model reference was found for the corresponding Trinity-gene. The log2Foldchanges at the time points 0h, 3h and 6h are given in columns E (“log2Foldchange.0h”), F (“log2Foldchange.3h”) and G (“log2Foldchange.6h”) respectively. For known and published genes, the Pubmed accession number is given in column “Pubmed”. Clusters according to the performed hierarchical clustering indicating the main cell states of gene expression are given in column “Cluster.cell.state”. The “NCBI.Accession” number with the corresponding “NCBI.e.value” and “NCBI.Description” was given for the Trinity genes, for which a blast hit was found. NR-genes were manually analysed for predicted subcellular localization and function of encoded proteins as indicated in N and O. In case of human homologs predictions were based on information from GeneCards https://www.genecards.org. For *Hydra* genes with previously published information references and comments are provided in P. Colored blocks are used to separate genes expressed in different cell types visually, red letters indicate up-regulated genes, black letters indicate downregulated genes.

**Table S2**. *Summary of functional annotations of NR-genes*: 1, 2: Numbers NR-genes according to expression patterns of AEP-references and manually assigned subcellular localisation and function of predicted proteins; Selected gene-IDs are shown and referenced in last lane of table. For some genes additional alignments and phylogenetic comparisons are provided in Figure S3. 3: Overview of numbers of NR-genes with indicated cell state expression.

**Table S3**. *Enriched motifs in NR-genes regulatory regions*. Results of promoter analysis for NR-genes down-regulated at 0h, down-regulated at 0h and 3h and down-regulated at 0h, 3h and 6h. NR-genes up-regulated at 0h, up-regulated at 0h and 3h and up-regulated at 0h, 3h and 6h. The Motif ID column indicates the JASPAR database motif name and matrix ID; the Motif Sequence column indicates the motif consensus sequence, the FDR column indicates the adjusted p-value (i.e. False Discovery Rate) statistic for the HOMER binomial enrichment test; the Frequency in NR Peaks column indicates the percentage of putative NR promoter regions that contained at least one instance of the target motif, with the total number of peaks considered provided in the column header; the Frequency in non-NR peaks indicates the percentage of control peak regions (i.e. all peaks that were not in the NR peak list) that contained at least one instance of the target motif, with the total number of peaks considered provided in the column header; the Fold Enrichment column indicates the ratio in the frequency of motif occurrences in NR peaks when compared to non-NR peaks; the Binding TF Class column indicates the class of DNA binding domain that binds the target motif; and the Potential NR Regulators column indicates NR transcription factors that could plausibly be regulating the target motif based on their expression dynamics during the DAPT treatment time course. All motifs with an enrichment FDR > 0.05 were discarded.

**Table S4**. *Expression dynamics of NR transcription factors*. Transcription factors that showed significant changes in response to DAPT treatment were grouped into five categories: downregulated but recovered by 3 hours post-treatment, downregulated but recovered by 6 hours post-treatment, downregulated and remained downregulated at 6 hours post-treatment, upregulated but recovered by 3 hours post-treatment, and upregulated but recovered by 6 hours post-treatment. We did not identify any transcription factors that were upregulated and remained upregulated by 6 hours post-treatment. The Trinity ID column indicates the name given to the transcripts that were assembled for this study using Trinity assembly software, the AEP ID column indicates the transcript’s corresponding name in the aepLRv2 transcriptome (Siebert et al., 2019), the Gene Model column indicates transcript’s corresponding gene model name in the *Hydra* 2.0 genome (arusha.nhgri.nih.gov/hydra), the Log2FC columns provide log2 fold changes for each respective time point (0, 3, or 6 hours post-treatment) relative to DMSO-treated controls (non-significant differentials are reported as 0), the Pubmed Hit column provides the transcript’s corresponding accession number in Pubmed, the NCBI Accession and NCBI Description columns provide information on the transcript’s best BLAST hit when searching the NCBI NR database, the Short Name column provides an abbreviated transcript annotation, and the DNA Binding Domain column indicates the type of DNA binding domain identified in the protein sequence using PFAM (annotations generated as part of the *Hydra* 2.0 genome annotation project; arusha.nhgri.nih.gov/hydra).

**Table S5**. *RPPJ-motif frequency in NR-genes:* RBPJ-motif frequencies are given for NR-genes with a minimum of 3 such motifs. AEP-IDs, minimum distance from the transcriptional start, protein annotations or identified conserved domains and expression patterns are listed. Colours: brown: genes are expressed predominantly in epithelia cells of the head and tentacles; blue: genes expressed in foot; green: genes expressed in nematoblasts/nematocytes and white: others

## Notes

### Competing Interest Statement

The authors have declared no competing interest.

