## Supplementary figures and images for "Differential gene regulation in DAPT-treated Hydra reveals molecular pathways dependent on Notch signalling during interstitial cell differentiation and formation of the oral-aboral axis in *Hydra*"

### supplemental Figure S1

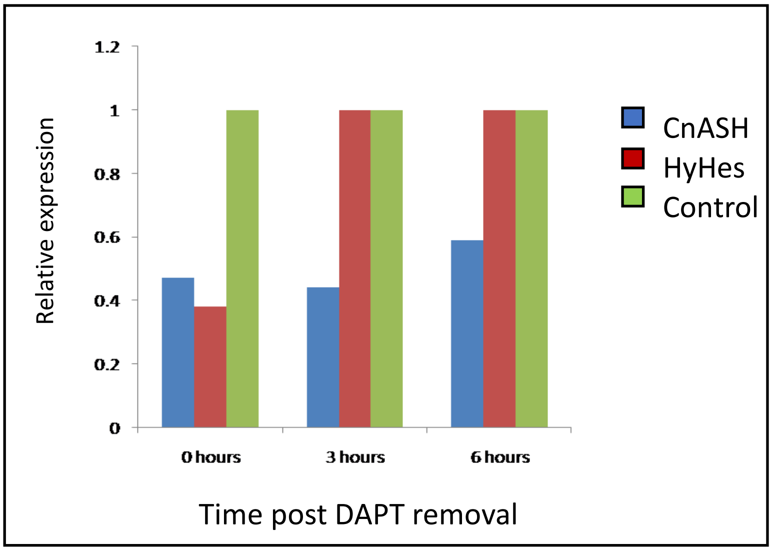
