## supplemental Figure S2 metagenes for "Differential gene regulation in DAPT-treated Hydra reveals molecular pathways dependent on Notch signalling during interstitial cell differentiation and formation of the oral-aboral axis in *Hydra*"

### Nematoblast progenitors

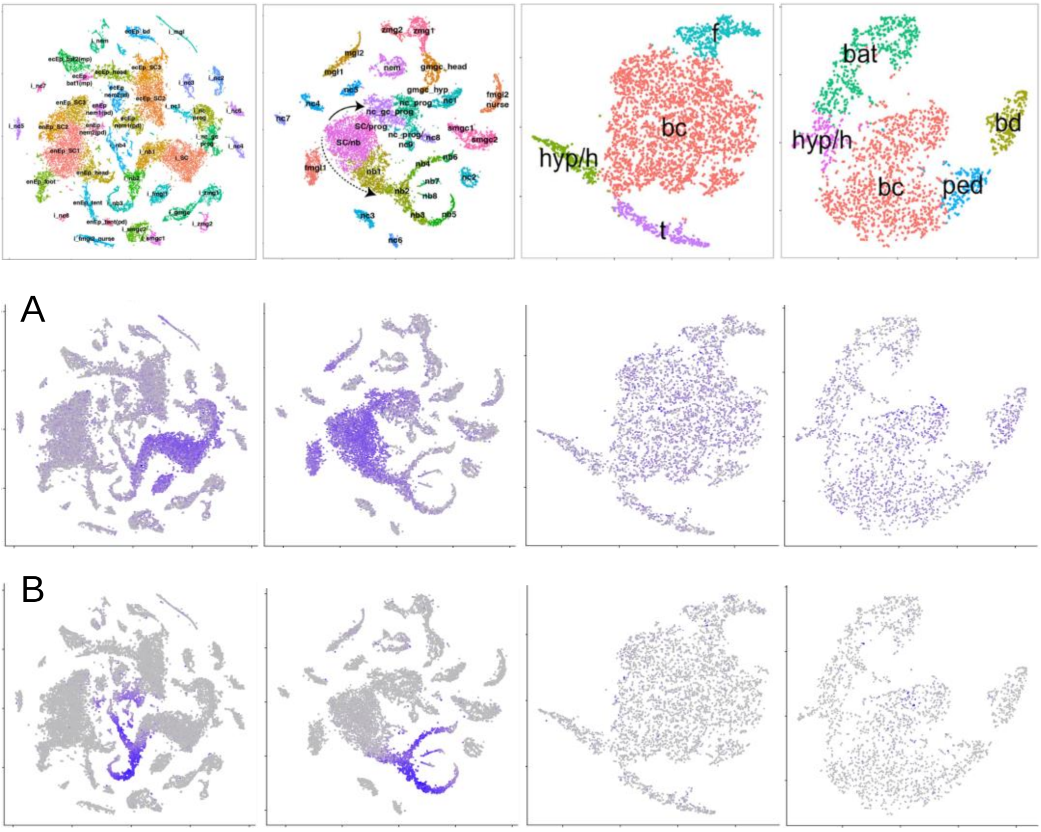

### Desmonemes

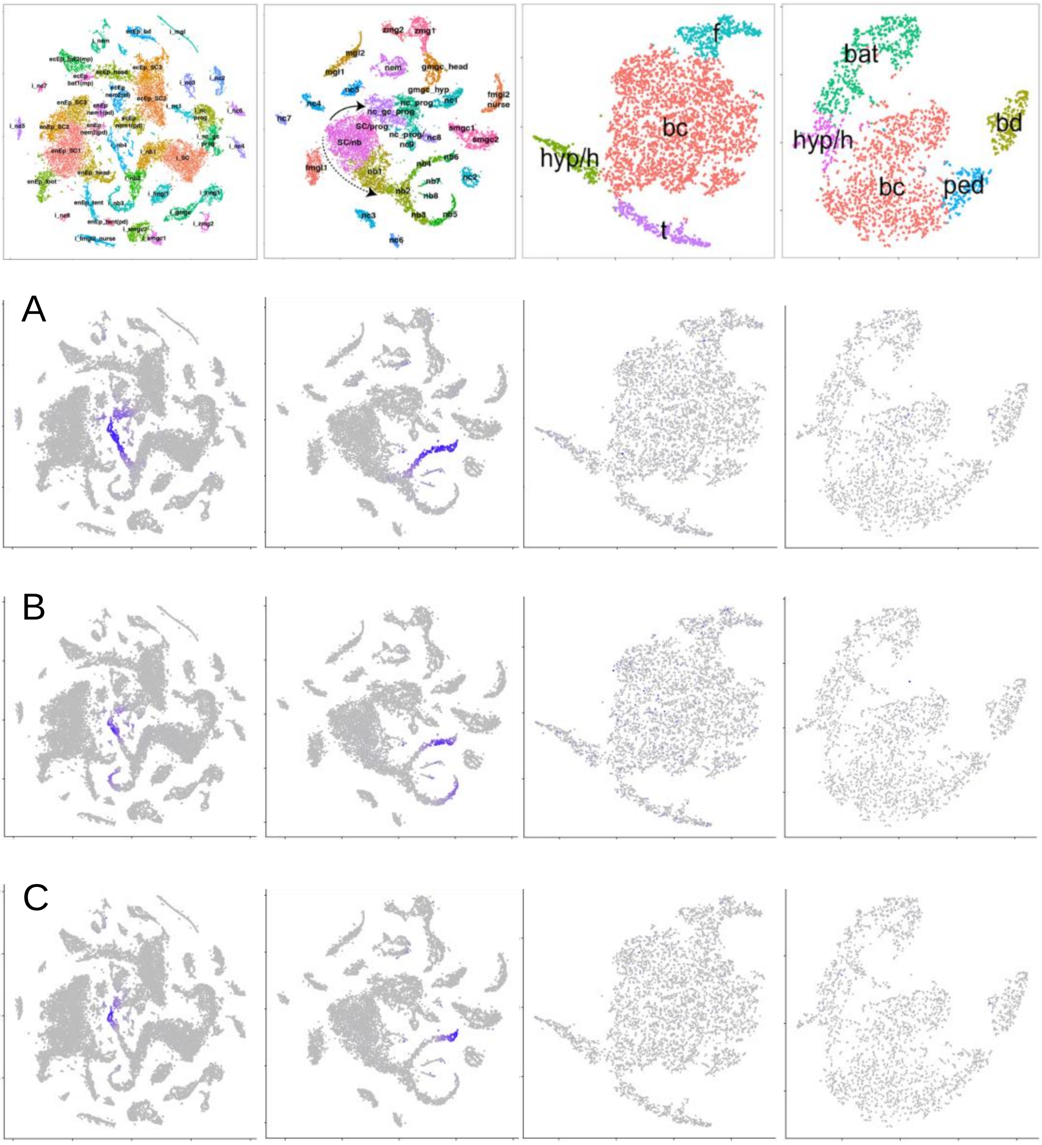

Stenoteles

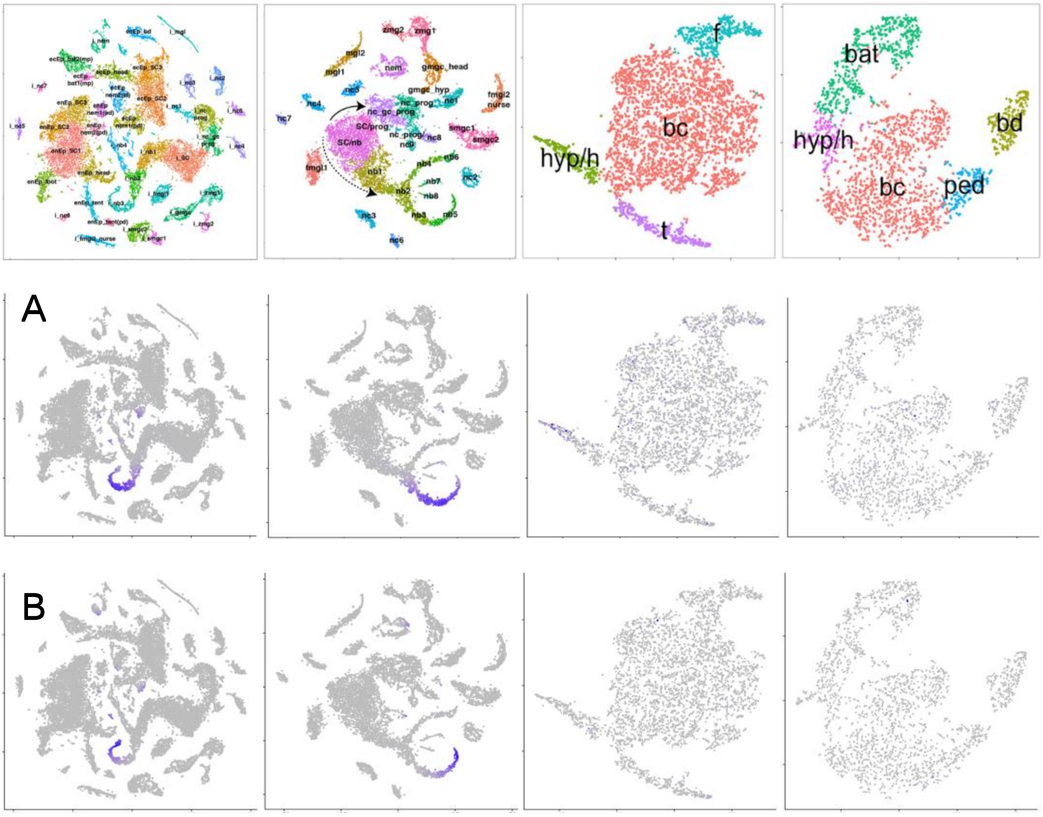

zmg2
zmg1
mgl2
mgl1
nem
gmgc_head
gmgc_hyp
fmgl2
nurse
nc5
nc4
nc7
nc1
SC/prog
SC/nb
smgc1
smgc2
fmgl1
nb1
nb2
nb3
nb4
nb5
nb6
nb7
nb8
nc2
nc3
nc6
f
bc
hyp/h
t
bat
hyp/h
bd
bc
ped
A
B

Isorhizas

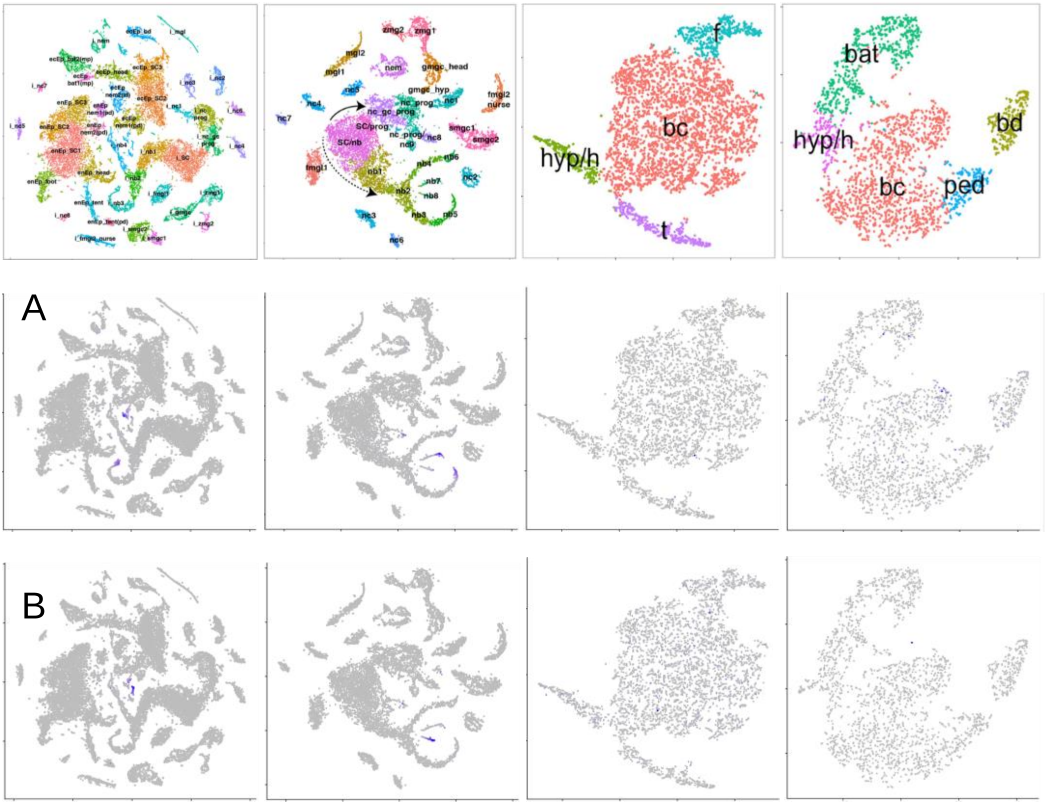

#### Mature nematocytes

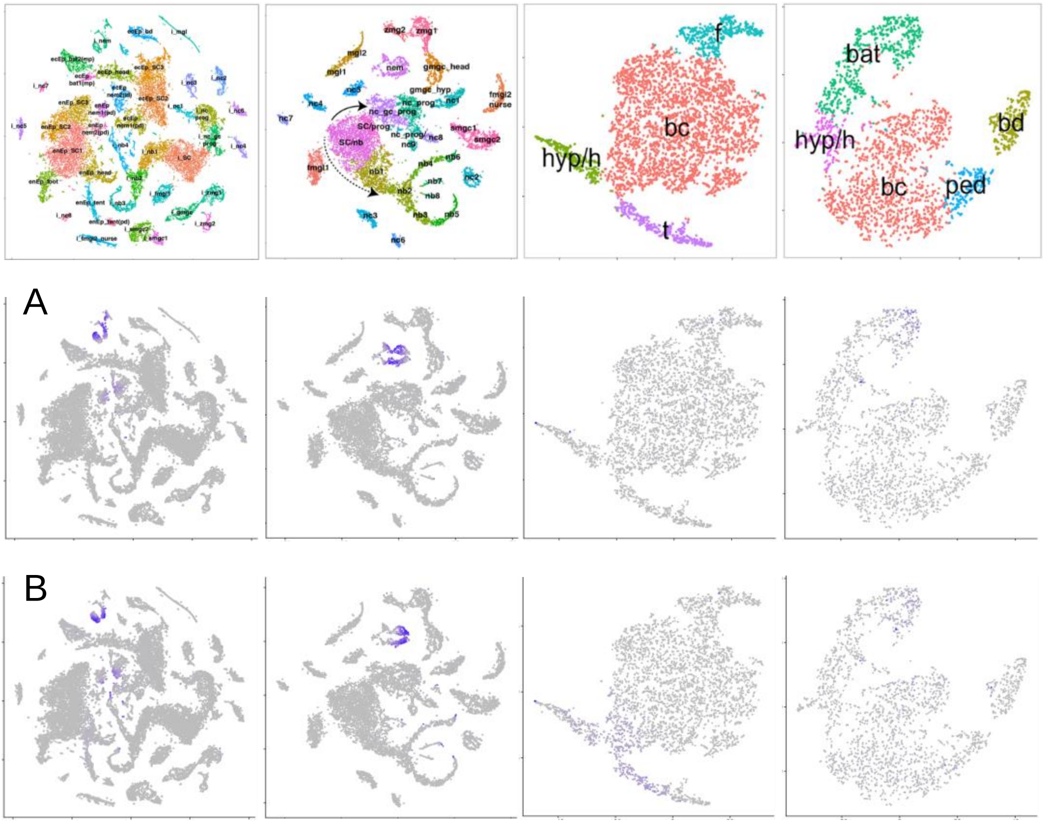

Ectoderm specific - body column

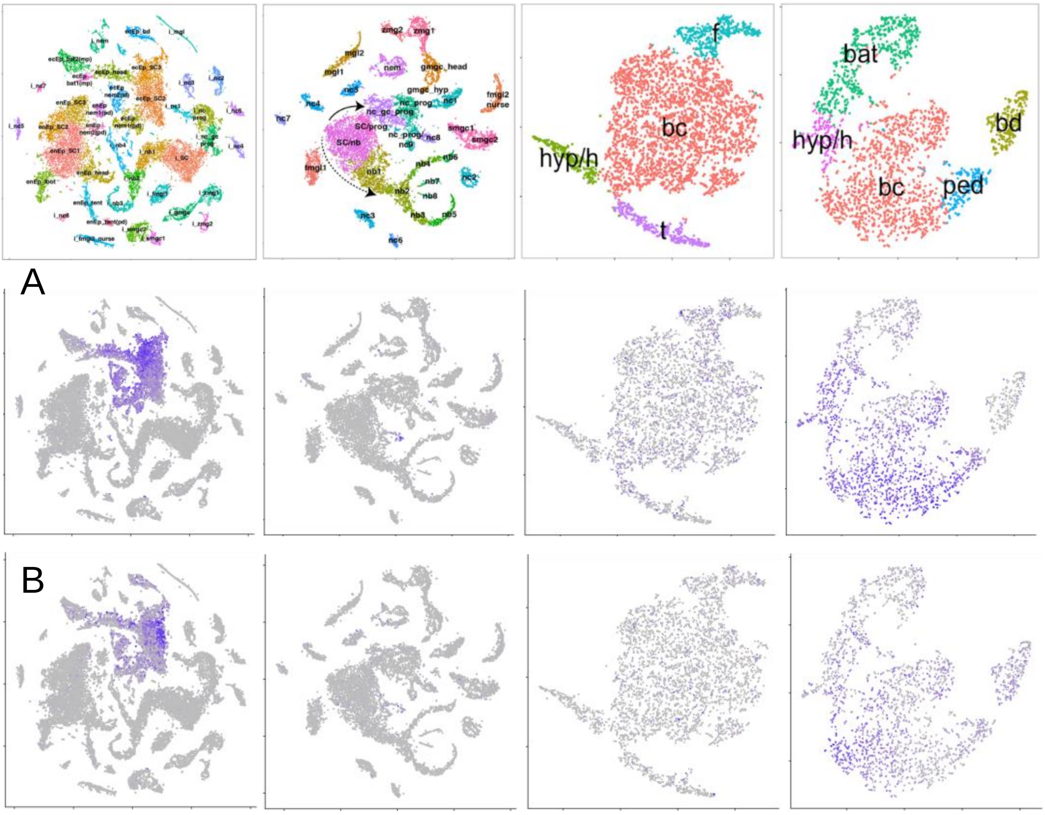


Ectoderm specific - battery cells

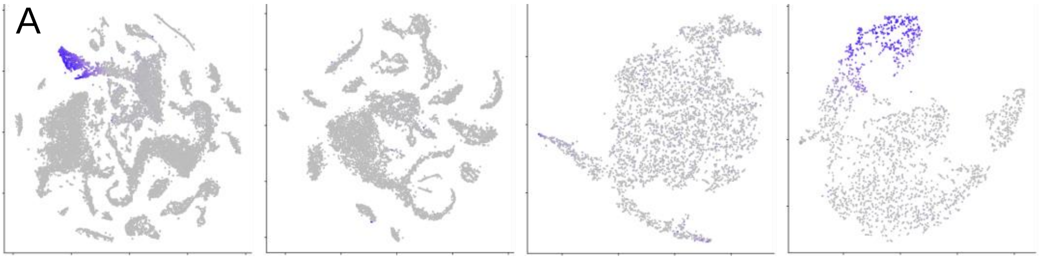


Ectoderm specific - basal disk

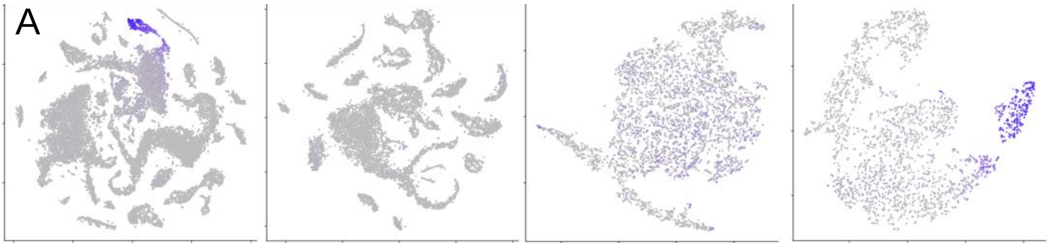

### Endoderm specific - body column

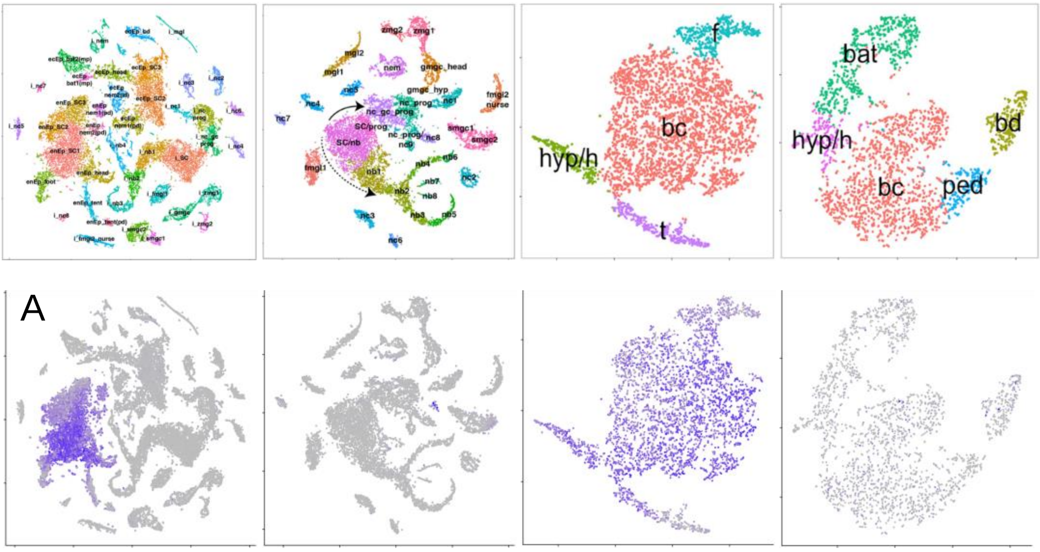


### Endoderm specific - tentacle

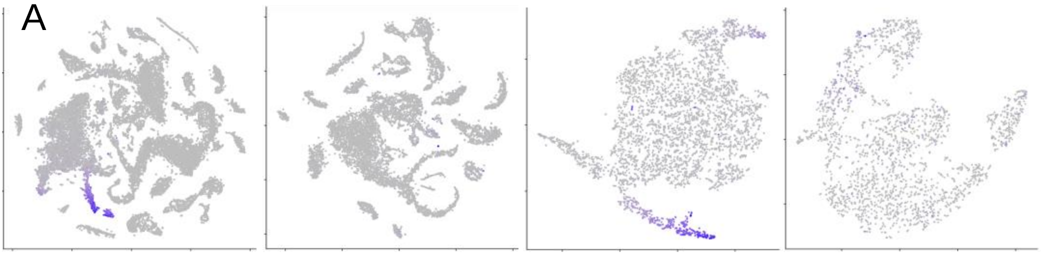

### Battery + Nematocytes

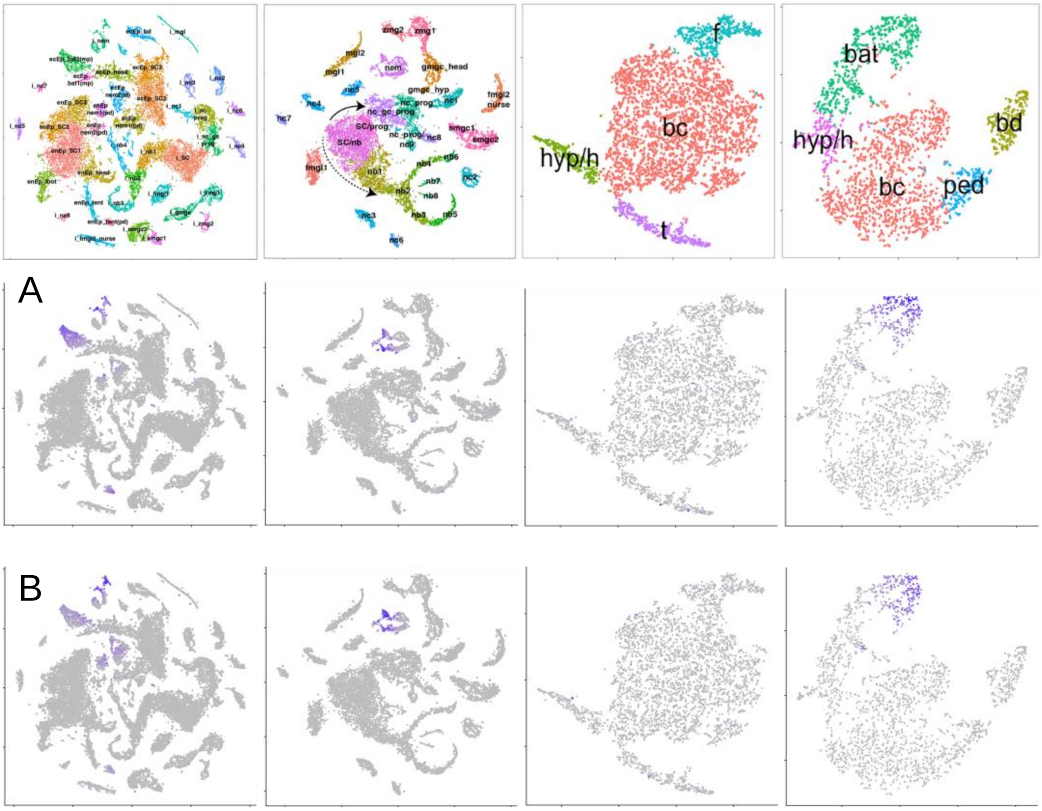

#### Ecto.Endo - foot

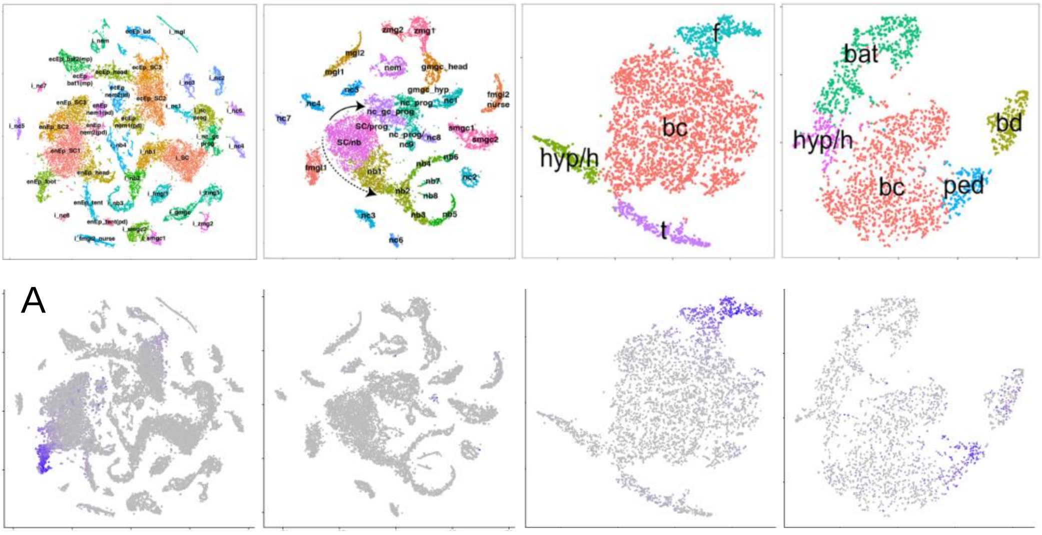


#### Ecto.Endo - head

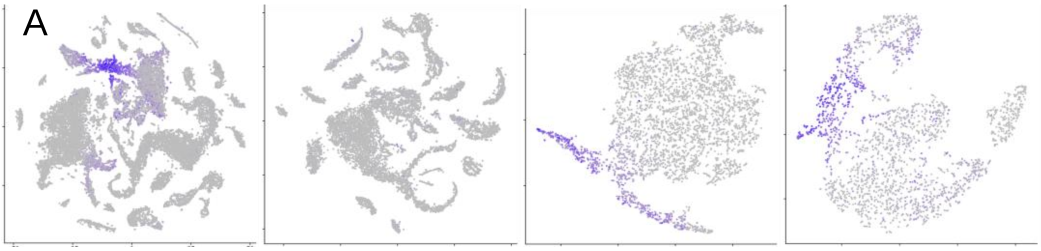

### IC derivatives - gland cells

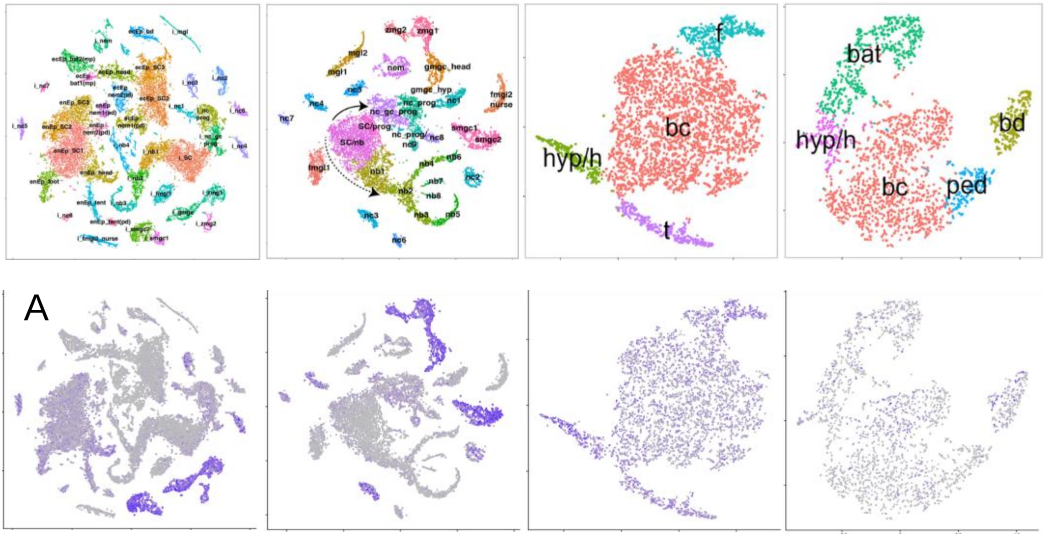


### IC derivatives - female nurse cells

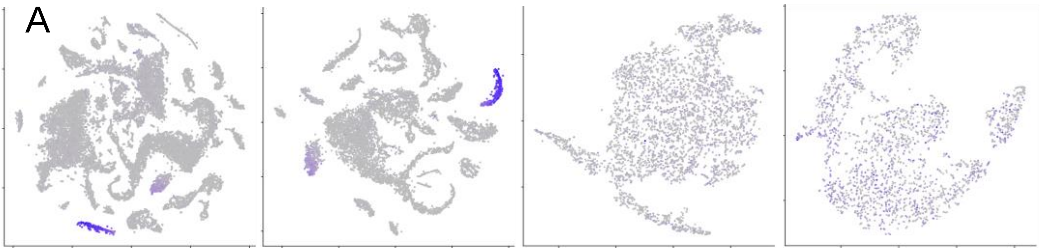
