## supplemental Table S2 for "Differential gene regulation in DAPT-treated Hydra reveals molecular pathways dependent on Notch signalling during interstitial cell differentiation and formation of the oral-aboral axis in *Hydra*"

### 1 UP-REGULATED NR-GENES:

207 genes, 166 with AEP reference, 41 have no AEP reference

The 41 Up-regulated NR-genes for which no AEP-annotation was found also lack expression information.

#### 1.1 UP-REGULATED NR-GENES WITH NO AEP REFERENCE (41 GENES)

| Predicted localisation<br>*According to gene cards of homolog or functional assignment* | Number of genes | Predicted function<br>*for majority* | Selected gene-IDs description and references |
|---|---|---|---|
| ECM | 7 | lectins<br>antistasin family-metalloprotease | PPOD, lectin (Böttger et al., 2012; Thomsen and Bosch, 2006) |
| ER | 1 | Cytochrome P450 | |
| Nucleus | 2 | splicing, histone-modification | |
| Plasma membrane | 2 | GPCRs | |
| Not known | 3 | disulfide cross-linking, associated with phospholipids | |
| Uncharacterised | 10 | | |
| No blast hit | 16 | | |

For 166 Up-regulated NR-genes an AEP annotation and thus expression information was found. These genes are listed in the following according to the cell states in which they are expressed.

#### 1.2 UP-REGULATED NR-GENES, EXPRESSED IN ECTODERMAL BASAL DISK (13 GENES)

| Predicted localisation<br>*According to gene cards of homolog or functional assignment* | Number of genes | Predicted function<br>*for majority* | Selected gene-IDs description and references |
|---|---|---|---|
| ECM | 3 | lectin<br>protease | PPOD2, lectin (Böttger et al., 2012; Thomsen and Bosch, 2006) |
| Not known | 4 | Redox-regulation, proteases, signalling | |
| Uncharacterised | 5 | | |

| | |
|---|---|
| No blast hit | 1 |

#### 1.3 Up-regulated NR-genes, expressed in Ectodermal Battery Cells (6 genes)

| Predicted localisation<br>*According to gene cards of homolog or functional assignment* | Number of genes | Predicted function<br>*for majority* | Selected gene-IDs description and references |
|---|---|---|---|
| ECM | 1 | lectin C-type | |
| Membrane, cytosol | 1 | Redox-regulation | |
| Plasma membrane | 1 | Transporter | |
| Nucleus | 1 | | |
| Uncharacterised | 2 | | |

#### 1.4 Table 1.4 Up-regulated NR-genes, expressed in Ectodermal Battery Cells AND Ectodermal Head Cells (6 genes)

| Predicted localisation<br>*According to gene cards of homolog or functional assignment* | Number of genes | Predicted function<br>*for majority* | Selected gene-IDs description and references |
|---|---|---|---|
| ECM | 2 | | |
| Nucleus | 1 | Transcriptional regulation | (t5966aep ); Kayak-like |
| Plasma membrane | 2 | Signalling (GPCR) | |
| Uncharacterized | 1 | | |

#### 1.5 Up-regulated NR-genes, expressed in Ectodermal Head Cells AND in Ectodermal Stem Cells (15 genes)

| Predicted localisation<br>*According to gene cards of homolog or functional assignment* | Number of genes | Predicted function<br>*for majority* | Selected gene-IDs description and references |
|---|---|---|---|
| Cytoskeleton | 1 | actin | |
| ECM | 1 | | |
| Plasma membrane | 1 | GPCR | |
| Not known | 3 | Redox-regulation, di-sulfide cross-linking, metabolism | |

| | |
|---|---|
| Uncharacterized | 2 |
| No blast hit | 7 |

#### 1.6 Up-regulated NR-genes, expressed in Ectodermal Epithelial Cells (22 genes)

| Predicted localisation<br>*According to gene cards of homolog or functional assignment* | Number of genes | Predicted function<br>*for majority* | Selected gene-IDs description and references |
|---|---|---|---|
| Cytosol | 1 | signalling | |
| ECM | 1 | lectin | |
| Endosome | 1 | regulates gamma secretase | (t12805aep); predicted Transmembrane emp24 domain-containing protein 2 |
| ER, Golgi | 9 | Vesicle traffic, apoptosis, redox-regulation, UPF, chaperone, glycosylation | (t5431aep); predicted canopy homolog (FGF-regulation) |
| Nucleus, cytosol | 1 | signalling | |
| Not known | 2 | Redox-regulation | |
| Uncharacterized | 3 | | |
| No blast hit | 4 | | |

#### 1.7 Up-regulated NR-genes, expressed in Endodermal Foot Cells (4 genes)

| Predicted localisation<br>*According to gene cards of homolog or functional assignment* | Number of genes | Predicted function<br>*for majority* | Selected gene-IDs description and references |
|---|---|---|---|
| ER | 1 | lipid associated | |
| Plasma membrane | 1 | signalling | |
| ECM, secreted | 1 | TGF-beta | (t25624aep), HydraTGF3; (Watanabe et al., 2014) |
| Unclear | 1 | Wnt-inhibition | (t11061aep) APCDD1-like, Figure S3 |

#### 1.8 Up-regulated NR-genes, expressed in Endodermal Head Cells (2 genes)

| Predicted localisation<br>*According to gene cards of homolog or functional assignment* | Number of genes | Predicted function<br>*for majority* | Selected gene-IDs description and references |
|---|---|---|---|
| Plasma membrane | 1 | GPCR | |
| No blast hit | 1 | | |

#### 1.9 Up-regulated NR-genes, expressed in Endodermal Tentacle Cells (12 genes)

| Predicted localisation<br>*According to gene cards of homolog or functional assignment* | Number of genes | Predicted function<br>*for majority* | Selected gene-IDs description and references |
|---|---|---|---|
| ECM | 3 | collagen modifier, spondin, matrixmetalloprotease | (t16424aep), HMMP, (Leontovich et al., 2000) |
| ER, Golgi | 2 | EGF regulator, PDI (protein disulfide-isomerase) | |
| Plasma membrane | 2 | GPCR | |
| Uncharacterized | 4 | | |
| No blast hit | 1 | | |

#### 1.10 Up-regulated NR-genes, expressed in Endodermal Epithelial Cells (31 genes)

| Predicted localisation<br>*According to gene cards of homolog or functional assignment* | Number of genes | Predicted function<br>*for majority* | Selected gene-IDs description and references |
|---|---|---|---|
| Cytosol, membrane | 3 | Rab-like G-protein, apoptosis | ACY95433.1 Caspase D; (Lasi et al., 2010) |
| Endosome, lysosome | 1 | Lipid binding | |
| ER | 4 | Protein folding, calcium binding, redox-regulation | |
| Plasma membrane | 7 | GPCRs, protease, carrier | |
| Not known | 4 | leucine rich, Zinc finger, USP | |
| Uncharacterized | 11 | | |
| No blast hit | 1 | | |

#### 1.11 Up-regulated NR-genes, expressed in Female Nurse Cells (3 genes)

| Predicted localisation *According to gene cards of homolog or functional assignment* | Number of genes | Predicted function *for majority* | Selected gene-IDs description and references |
|---|---|---|---|
| Nucleus, cytoplasm | 2 | RNA-processing, signalling | |
| Uncharacterised | 1 | | |

#### 1.12 Up-regulated NR-genes, expressed in both Epithelia (37 genes)

| Predicted localisation *According to gene cards of homolog or functional assignment* | Number of genes | Predicted function *for majority* | Selected gene-IDs description and references |
|---|---|---|---|
| ECM | 2 | Redox-regulation (cuticle hardening), IGF-binding domain containing | |
| ER, Golgi, endosome | 13 | PDI, sugar transport, glycosylation, hypoxia, HSP/chaperone/stress induced | (t3105aep); predicted protein Mind bomb (Notch signalling), Böttger unpublished |
| Plasma membrane | 2 | GPCR, transporter | |
| Nucleus | 4 | cell cycle, splicing, rRNA processing, transcriptional regulation | (t38458aep); CBF-beta domain |
| Vesicles | 1 | SNARE | |
| Not known | 2 | transposase | |
| Uncharacterised | 7 | | |
| No blast hit | 6 | | |

#### 1.13 Up-regulated NR-genes, expressed in Mucous Gland Cells (1 gene)

| Predicted localisation *According to gene cards of homolog or functional assignment* | Number of genes | Predicted function *for majority* | Selected gene-IDs description and references |
|---|---|---|---|
| No blast hit | 1 | | |

##### 1.14 UP-REGULATED NR-GENES, EXPRESSED IN NEMATOBLASTS STAGE NB6 (1 GENE)

| Predicted localisation *According to gene cards of homolog or functional assignment* | Number of genes | Predicted function *for majority* | Selected gene-IDs description and references |
|---|---|---|---|
| Uncharacterized | 1 | | |

##### 1.15 UP-REGULATED NR-GENES, EXPRESSED IN NEURONS (5 GENES)

| Predicted localisation *According to gene cards of homolog or functional assignment* | Number of genes | Predicted function *for majority* | Selected gene-IDs description and references |
|---|---|---|---|
| Golgi | 1 | Glycosylation | |
| Plasma membrane | 1 | GPCR | |
| Not known | 1 | Protein modification | |
| Uncharacterised | 2 | | |

##### 1.16 UP-REGULATED NR-GENES, EXPRESSED IN NEURONS AND NEMATOBLASTS (4 GENES)

| Predicted localisation *According to gene cards of homolog or functional assignment* | Number of genes | Predicted function *for majority* | Selected gene-IDs description and references |
|---|---|---|---|
| ER, Golgi | 3 | unfolded protein response (ER stress), chaperone, secretion | |
| Cytosol | 1 | signalling phosphatase | |

##### 1.17 UP-REGULATED NR-GENES, SPORADIC EXPRESSION (4 GENES)

| Predicted localisation *According to gene cards of homolog or functional assignment* | Number of genes | Predicted function *for majority* | Selected gene-IDs description and references |
|---|---|---|---|
| Plasma membrane | 1 | GPCR | |
| Cytoskeleton | 1 | microtubule binding | |
| Uncharacterized | 1 | | |
| No blast hit | 1 | | |

### 2 DOWN-REGULATED NR-GENES:

624 genes, 500 with AEP reference, 124 have no AEP reference

The 124 Down-regulated NR-genes for which no AEP-annotation was found also lack expression information.

#### 2.1 DOWN-REGULATED NR-GENES WITH NO AEP REFERENCE (124 GENES)

| Predicted localisation *According to gene cards of homolog or functional assignment* | Number of genes | Predicted function *for majority* | Selected gene-IDs description and references |
|---|---|---|---|
| nematocyst | 6 | minicollagens, disulfide cross-linker. cnidoin | NM_001280961.1(Kurz et al., 1991); KR024036.1; cnidoin (Balasubramanian et al., 2012) |
| ECM, membrane and secreted | 4 | lectins, metalloprotease, secreted protein | |
| Cytoplasm | 2 | translation | |
| ER | 1 | chaperon | |
| Cytosol | 1 | translation | |
| ECM, secreted | 1 | secreted protein | |
| Mitochondrion | 1 | ATP-synthesis | |
| Ribosome | 13 | ribosomal protein | |
| Plasma membrane | 3 | Transporter, protease | |
| Cytoskeleton | 1 | microtubules | XP_002161507.1; predicted tubulin alpha |
| Not known | 12 | toxins, polyamine metabolism, chaperones, acid/basic homeostasis | |
| Uncharacterised | 21 | | |
| No blast hit | 61 | | |

For 500 Down-regulated NR-genes an AEP annotation and thus expression information was found. These genes are listed in the following according to the cell states in which they are expressed.

#### 2.2 Down-regulated NR-genes, expressed in Ectodermal Basal Disc and/or Endodermal Foot Cells (3 genes)

| Predicted localisation *According to gene cards of homolog or functional assignment* | Number of genes | Predicted function *for majority* | Selected gene-IDs description and references |
|---|---|---|---|
| Plasma membrane | 1 | Wnt-signalling | (t15331aep); secreted frizzled-related protein annotation provided by B. Hobmayer, Innsbruck |
| Not known | 1 | Redox-regulation | |
| No blast hit | 1 | | |

#### 2.3 Down-regulated NR-genes, expressed in Ectodermal Battery Cells (44 genes)

| Predicted localisation *According to gene cards of homolog or functional assignment* | Number of genes | Predicted function *for majority* | Selected gene-IDs description and references |
|---|---|---|---|
| ECM | 5 | lectins | |
| Cytosol | 4 | muscle contraction, chaperone, signalling (GTPase), $Ca^{2+}$ binding | (t9989aep); ras like protein |
| Cytoskeleton | 3 | actin binding, microtubules, actin | |
| ER, Golgi | 2 | transporter | |
| Nucleus | 1 | transcriptional regulation | (t17518aep); pred. protein couch potato-like |
| Plasma membrane, endosome | 10 | cell adhesion, ion channels, transporter | |
| Not known | 3 | signalling, ubiquitine ligase complex, methyltransferase | |
| Uncharacterized | 9 | | |
| No blast hit | 6 | | |

#### 2.4 DOWN-REGULATED NR-GENES, EXPRESSED IN ECTODERMAL BATTERY AND ECTODERMAL HEAD CELLS (13 GENES)

| Predicted localisation *According to gene cards of homolog or functional assignment* | Number of genes | Predicted function *for majority* | Selected gene-IDs description and references |
|---|---|---|---|
| Nucleus +cytosol and<br><br>Nucleus+plasma membrane | 4 | signalling<br><br>Wnt-regulation; planar cell polarity | (t21185aep); MAPKinase<br><br>(t19041aep) protein prickle-like (WntPCP regulator), Figure S3 |
| Nucleus | 2 | Transcriptional regulation, chromatin | (t16456aep ); HyALX (Smith et al., 2000), Figure S3 |
| ECM, secreted | 1 | FGF-signalling | (t8338aep); fibroblast growth factor 13-annotation as FGF-homolog was confirmed by Monika Hassel, Marburg, Germany |
| Plasma membrane | 2 | transporter | |
| Vesicles | 1 | metalloprotease | |
| Uncharacterized | 3 | | |

#### 2.5 DOWN-REGULATED NR-GENES, EXPRESSED IN ECTODERMAL HEAD CELLS (5 GENES)

| Predicted localisation *According to gene cards of homolog or functional assignment* | Number of genes | Predicted function *for majority* | Selected gene-IDs description and references |
|---|---|---|---|
| ECM, secreted | 2 | secreted proteins | (t20254aep), Hydra Ks1, (Weinziger et al., 1994)<br><br>(t10549aep ), (Khalturin et al., 2008) |
| Plasma membrane | 1 | GPCR | |
| Nucleus | 1 | Transcriptional regulation | (t33622aep); Otx1, (Reddy et al., 2019), Figure S3 |
| No blast hit | 1 | | |

#### 2.6 Down-regulated NR-genes, expressed in Ectodermal Head Cells AND Ectodermal Stem Cells (4 genes)

| Predicted localisation *According to gene cards of homolog or functional assignment* | Number of genes | Predicted function *for majority* | Selected gene-IDs description and references |
|---|---|---|---|
| Cytoplasm | 1 | ubiquitin ligase | |
| Plasma membrane | 1 | Gap junction | innexin 9a; (Takaku et al., 2014) |
| Uncharacterized | 2 | | |

#### 2.7 Down-regulated NR-genes, expressed in Ectodermal Epithelial Cells (8 genes)

| Predicted localisation *According to gene cards of homolog or functional assignment* | Number of genes | Predicted function *for majority* | Selected gene-IDs description and references |
|---|---|---|---|
| Nucleus | 1 | Transcriptional regulation | (t34112aep); Max dimerization protein 1, Figure S3 |
| Nucleus, cytosol | 1 | ubiquitin ligase | (t9865aep); fizzy-related protein homolog, APC adapter |
| Plasma membrane | 2 | Transporter, signalling | |
| Not known | 1 | | |
| Uncharacterized | 2 | | |
| No blast hit | 1 | | |

#### 2.8 Down-regulated NR-genes, expressed in Endodermal Head Cells (7 genes)

| Predicted localisation *According to gene cards of homolog or functional assignment* | Number of genes | Predicted function *for majority* | Selected gene-IDs description and references |
|---|---|---|---|
| Nucleus | 3 | Transcriptional regulation | (t5275aep) PITX, (t5528aep) Sox21B, Figure S3, ); (t1216aep) CnGSC (Broun et al., 1999) |
| ECM, secreted | 1 | Wnt signalling | Hydra Wnt7; (Lengfeld et al., 2009) |
| Uncharacterized | 3 | | |

#### 2.9 DOWN-REGULATED NR-GENES, EXPRESSED IN ENDODERMAL TENTACLE CELLS (30 GENES)

| Predicted localisation *According to gene cards of homolog or functional assignment* | Number of genes | Predicted function *for majority* | Selected gene-IDs description and references |
|---|---|---|---|
| cytosol | 3 | signalling (GTPase) | |
| ECM, secreted | 6 | lectin, metalloprotease, collagen, cysteine rich, redox-regulation | |
| ER | 2 | Cytochrom P450, Calcium release like IP3 | |
| Nucleus | 3 | Transcriptional regulation | (t22023aep), Nuclear factor related; (t26873aep), Krüppel-like |
| Plasma membrane | 3 | GPCR, aquaporin, enzyme | |
| Not known | 4 | hydrolases, methyltransferase, ribonuclease | |
| Uncharacterized | 6 | | |
| No blast hit | 3 | | |

#### 2.10 DOWN-REGULATED NR-GENES, EXPRESSED IN ENDODERMAL EPITHELIAL CELLS (9 GENES)

| Predicted localisation *According to gene cards of homolog or functional assignment* | Number of genes | Predicted function *for majority* | Selected gene-IDs description and references |
|---|---|---|---|
| Cytosol | 1 | signalling | (t1767aep); Ras-like protein |
| ER, endosome, lysosome | 1 | hydrolase | |
| Nucleus | 2 | transcriptional regulation | (t11826aep); TCF (Hobmayer et al., 2000) |
| Plasma membrane | 2 | GPCR | |
| Uncharacterized | 3 | | |

#### 2.11 Down-regulated NR-genes, expressed in Female Nurse Cells (4 genes)

| Predicted localisation *According to gene cards of homolog or functional assignment* | Number of genes | Predicted function *for majority* | Selected gene-IDs description and references |
|---|---|---|---|
| Plasma membrane | 1 | GPCR | |
| Nucleus | 2 | tanscriptional regulation, RNA binding | (t16018aep) IRX6; see Figure S3, (t22296aep), Boule-like |
| Unclear | 1 | | |

#### 2.12 Down-regulated NR-genes, expressed in Both Epithelia (18 genes)

| Predicted localisation *According to gene cards of homolog or functional assignment* | Number of genes | Predicted function *for majority* | Selected gene-IDs description and references |
|---|---|---|---|
| Cytosol | 2 | metabolism, protein methylation | |
| ECM | 1 | apoptosis (hydrolase) | |
| Endosome, lysosome | 1 | thioredoxin family | |
| ER, Golgi | 1 | vesicle trafficking | |
| Nucleus | 4 | transcriptional regulation, Ubiquitin-ligase, mi-RNA-ubiquitination, | (t29291aep) Sp5(Vogg et al., 2019) |
| Plasma membrane (ER) | 3 | ion pump, GPCR, metabolism | |
| Not known | 3 | acid/basic homeostasis, poly-gamma-glutamate synthesis | |
| Uncharacterized | 2 | | |
| No blast hit | 1 | | |

#### 2.13 DOWN-REGULATED NR-GENES, EXPRESSED IN MATURE NEMATOCYTES (89 GENES)

| Predicted localisation<br>*According to gene cards of homolog or functional assignment* | Number of genes | Predicted function<br>*for majority* | Selected gene-IDs description and references |
|---|---|---|---|
| Cytosol | 7 | signalling, calcium binding, enzyme, protease inhibitor | (t21185aep), pred. MAPK |
| Cytoskeleton | 9 | microtubules, nuclear lamina, actin binding, actin, cilia | (t14035aep), pred. tubulin alpha chain |
| ECM | 4 | Laminin subunits, tetraspanin family | |
| ER, Golgi (secretory granules) | 3 | transporter, RNA localisation | (t18510aep) pred. protein bicaudal C homolog 1-like |
| Nucleus | 4 | transcriptional regulation | (t26616aep); pred. transcription factor Ovo-like 1, (t26993aep); pred. Zinc finger protein Fez 2 like |
| Plasma membrane | 14 | aquaporin, gap junction, signalling | (t25400aep); HyTRAF4, TNF -receptor adaptor; (Steichele 2021);<br>(t21368aep) innexin (Takaku et al., 2014) |
| Non-coding RNA | 1 | | |
| Not known | 9 | glutathione synthesis, chaperone, protease, phospholipid-signaling, Redox-regulation, acid/basic homeostasis | |
| Uncharacterized | 30 | | |
| No blast hit | 8 | | |

#### 2.14 Down-regulated NR-genes, expressed in Mucous Gland Cells (3 genes)

| Predicted localisation *According to gene cards of homolog or functional assignment* | Number of genes | Predicted function *for majority* | Selected gene-IDs description and references |
|---|---|---|---|
| ER, Golgi | 1 | protease (APP processing) | (t14222aep); pred. beta-secretase |
| No blast hit | 2 | | |

#### 2.15 Down-regulated NR-genes, expressed in Nematoblasts, stages Nb4 through Nb8 (94 genes)

| Predicted localisation *According to gene cards of homolog or functional assignment* | Number of genes | Predicted function *for majority* | Selected gene-IDs description and references |
|---|---|---|---|
| Cytoskeleton | 1 | Cilia | (t13479); nematogalectin (Hwang et al., 2008; Hwang et al., 2010) |
| Cytosol | 2 | enzyme, signalling | |
| ECM | 9 | mucous, collagen, cell wall assembly, lectin | |
| ER, Golgi, endosome, lysosome | 5 | lectin binding, glycolipid hydroxylation, proteases, protein folding | |
| Nematocyst | 9 | Nematocyte discharge<br>Nematocyst wall | (t22808); nematoblast specific nb042 and nb035 (Milde et al., 2009); (t20081aep); cnidoin (Beckmann et al., 2015) (t15237); NOWA; (Engel et al., 2002) |
| Nucleus | 4 | transcriptional regulation; DNA damage | (t21636aep); Prdl-b; (Gauchat et al., 1998); (t17964aep); HyJun, see Figure S3 |
| Plasma membrane (vesicles) | 11 | neurotransmitter uptake, transporter, protease, glutathione catabolic processing, ion pump, gap junction, GPCR | (t8479aep); innexin, (Takaku et al., 2014) |

| | | | |
|---|---|---|---|
| Not known | 9 | oxidation, disulfide cross-linking, glutatione hydrolase, Wnt-inhibitor | (20111aep); dickkopf 3 related (Fedders et al., 2004), Fig. S3 |
| Uncharacterized | 40 | | |
| No blast hit | 4 | | |

#### 2.16 Down-regulated NR-genes, expressed in Nematoblasts, stage Nb5 mainly (41 genes)

| Predicted localisation *According to gene cards of homolog or functional assignment* | Number of genes | Predicted function *for majority* | Selected gene-IDs description and references |
|---|---|---|---|
| Cytosol | 1 | signalling (tyrkinase) | |
| ECM | 2 | ECM turnover, metalloproteinase | |
| ER | 1 | signalling ($Ca^{2+}$) | |
| Plasma membrane | 2 | signalling (phospholipid)<br>GPCR | (t731aep); pred. secreted phospholipase A2 |
| Nematocyst | 1 | | (t35089aep) nematoblast specific protein nb042; (Milde et al., 2009) |
| Nucleus | 3 | transcriptional regulation, splicing, RNA binding | (t9145aep); forkhead box protein I1c-like; Fig. S3 |
| Not known | 4 | glycine rich protein, , proteases | |
| Uncharacterized | 26 | | |
| No blast hit | 1 | | |

#### 2.17 Down-regulated NR-genes, expressed in Nematoblasts, stage Nb6 mainly (72 genes)

| Predicted localisation<br>*According to gene cards of homolog or functional assignment* | Number of genes | Predicted function<br>*for majority* | Selected gene-IDs description and references |
|---|---|---|---|
| Cytoskeleton | 3 | Cilia | (t13479aep); (t13480aep); nematogalectins (Hwang et al., 2010) |
| Cytosol | 1 | protein modification | |
| ECM | 8 | cell wall assembly, nematoblast specific, lectins, mucous, protease | (t18591aep) minicollagen (Adamczyk et al., 2008; Khalturin et al., 2008) |
| ER, Golgi, vesicles | 6 | Proteases (neuropeptide processing), fucosidase, secretion | |
| lysosome-, endosome menbrane, nucleus | 1 | apoptosis | |
| Membrane vesicles | 1 | transporter (glutamate) | |
| Nematoblast | 3 | Nematoblast specific | (t28154aep); N-Col-4, minicollagen; (Kurz et al., 1991); (t9099aep); nematoblast specific protein nb012a(Milde et al., 2009) |
| Nucleus | 7 | transcriptional regulation | t23837aep Sox-21-B-like, t23172aep Sox 14 like; Fig. S3 (t11335aep) HyPOU4, (Siebert et al., 2019); (t24975aep) homeobox protein cut-like; Fig. S3; t12948aep); Forkhead box protein N1, Fig. S3 |
| Plasma membrane | 5 | aquaporin, proton channel, transporter, signalling (hormone processing) | |
| Unclear | 8 | disulfide cross-linking, actin binding, glutathion synthesis | |
| Uncharacterized | 20 | | |

| | |
|---|---|
| No blast hit | 9 |

#### 2.18 Down-regulated NR-genes, expressed in Nematoblasts, stage Nb8 mainly (18 genes)

| Predicted localisation *According to gene cards of homolog or functional assignment* | Number of genes | Predicted function *for majority* | Selected gene-IDs description and references |
|---|---|---|---|
| nematocyst | 1 | minicollagen | (t29423aep), (Kurz et al., 1991) |
| Plasma membrane | 1 | Signalling | |
| Not known | 1 | acid/basic homeostasis | |
| Uncharacterized | 12 | | |
| No blast hit | 3 | | |

#### 2.19 Down-regulated NR-genes, expressed in Neurons (26 genes)

| Predicted localisation *According to gene cards of homolog or functional assignment* | Number of genes | Predicted function *for majority* | Selected gene-IDs description and references |
|---|---|---|---|
| Cytosol | 1 | Actin | |
| Cytoskeleton | 1 | Microtubules | |
| ECM | 2 | lectin receptor, tetraspanin family | |
| ER, Golgi | 1 | Transporter | |
| Nucleus (cytosol) | 3 | transcriptional regulation | (t3617aep) HyHes (Münder et al., 2010) (expressed in neurons and in endodermal head cells |
| Plasma membrane | 6 | GPCR, RTK-signaling, transporter, ion channel, | |
| Not known | 2 | Exonuclease, enzyme | |
| Uncharacterized | 7 | | |
| No blast hit | 3 | | |

#### 2.20 DOWN-REGULATED NR-GENES, EXPRESSED IN NEURONS AND NEMATOBLASTS (2 GENES)

| Predicted localisation *According to gene cards of homolog or functional assignment* | Number of genes | Predicted function *for majority* | Selected gene-IDs description and references |
|---|---|---|---|
| Cytosol | 1 | Cell cycle/ cell death | |
| Nucleus | 1 | transcriptional regulation | (19720aep) forkhead box protein P1-B, Fig. S3 |

#### 2.21 DOWN-REGULATED NR-GENES, SPORADIC EXPRESSION (10 GENES)

| Predicted localisation *According to gene cards of homolog or functional assignment* | Number of genes | Predicted function *for majority* | Selected gene-IDs description and references |
|---|---|---|---|
| Cytoskeleton | 1 | actin motor | |
| Cytosol | 1 | metabolism | |
| ER | 1 | signalling (ceramid) | |
| Nucleus | 2 | transcriptional regulation, chromatin modification | (t10853aep) CnASH (Grens et al., 1995) |
| Not known | 1 | | |
| No blast hit | 2 | | |

### 3 OVERVIEW

| | **Up-regulated NR-genes** | **Down-regulated NR-genes** |
|---|---|---|
| NR-genes with no AEP reference | 41 | 124 |
| NR-genes with an AEP reference | 166 | 500 |
| **Cell states**<br>Only for NR-genes with an AEP reference | | |
| Ectodermal Basal disk | 13 | 3 |
| Ectodermal battery | 6 | 44 |
| Ectodermal battery and head | 6 | 13 |
| Ectodermal head | 0 | 5 |
| Ectodermal head and stem cells | 15 | 4 |
| Ectodermal epithelia | 22 | 8 |
| Endodermal head | 2 | 7 |
| Endodermal tentacle | 12 | 30 |
| Endodermal epithelia | 31 | 9 |
| Endodermal foot | 4 | 0 |
| Female nurse cells | 3 | 4 |
| Both epithelia | 37 | 18 |
| Mature nematocytes | 0 | 89 |
| Gland cells | 1 | 3 |
| Nematoblasts Nb4 through Nb8 | 0 | 94 |
| Nematoblasts Nb5 | 0 | 41 |
| Nematoblasts Nb6 | 1 | 72 |
| Nematoblasts Nb8 | 0 | 18 |
| Neurons | 5 | 26 |
| Neurons and nematoblasts | 4 | 2 |
| Sporadic | 4 | 10 |
