## supplemental Table S3 for "Differential gene regulation in DAPT-treated Hydra reveals molecular pathways dependent on Notch signalling during interstitial cell differentiation and formation of the oral-aboral axis in *Hydra*"

### Motifs enriched in the promoters of downregulated genes that recovered by 3 hours post-treatment

| Motif ID | Motif Sequence | FDR | Frequency in NR Peaks (n= 173) | Frequency in non-NR Peaks (n= 69267) | Fold Enrichment | Binding TF Class | Potential NR Regulators |
|---|---|---|---|---|---|---|---|
| ind/MA0228.1 | YTAATTA | 0.0164 | 88.44% | 75.77% | 1.1672 | Homeo domain factors | CnAlx (t16456aep); Smith 2000, Fig. S3<br>OTX1B (t33622aep); Fig. S3<br>PITX1 (t5275aep); pred.<br>aristaless-like (t16456aep); pred. |
| TBX1/MA0805.1 | AGGTGTGA | 0.0164 | 42.77% | 28.26% | 1.5134 | T-Box factors | |
| ETV5/MA0765.1 | ACCGGAAGTN | 0.0183 | 62.43% | 48.04% | 1.2995 | Tryptophan cluster factors | |
| FOXP2/MA0593.1 | AWGTAAACARA | 0.0183 | 89.02% | 77.89% | 1.1429 | Fork head / winged helix factors | FoxP1 (t19720aep) |
| CUX1/MA0754.1 | TAATCGATAH | 0.0183 | 80.92% | 68.34% | 1.1841 | Homeo domain factors | CnAlx (t16456aep); Smith 2000, Fig. S3<br>OTX1B (t33622aep); Fig. S3<br>PITX1 (t5275aep); pred.<br>aristaless-like (t16456aep); pred. |
| MEF2A/MA0052.3 | KCTAWAAATAGA | 0.0219 | 71.68% | 58.42% | 1.2270 | MADS box factors | |
| SOX10/MA0442.2 | NDAACAAAGVN | 0.0219 | 94.22% | 85.45% | 1.1026 | High-mobility group (HMG) domain factors | Sox14(t23172aep); Fig. S3<br>TF7-like 2 (t11826aep), pred. |
| D/MA0445.1 | TCCATTGTTBT | 0.0219 | 91.33% | 81.53% | 1.1202 | High-mobility group (HMG) domain factors | Sox14(t23172aep); Fig. S3<br>TF7-like 2 (t11826aep), pred. |
| Hoxd8/MA0910.1 | TAADTAATTAATRGCTW | 0.0283 | 90.75% | 81.40% | 1.1149 | Homeo domain factors | |
| Ahr::Arnt/MA0006.1 | YGCGTG | 0.0401 | 73.41% | 61.81% | 1.1877 | Basic helix-loop-helix factors (bHLH) | HyHES (t3617aep); Münder 2013<br>TFE3 (t22195aep)<br>Mad-protein (t34122aep) |
| PAX7/MA0680.1 | TAATCGATTA | 0.0401 | 50.29% | 38.31% | 1.3127 | Paired box factors | |
| cad/MA0216.2 | RGCCATAAAAM | 0.0401 | 90.17% | 81.46% | 1.1069 | Homeo domain factors | CnAlx (t16456aep); Smith 2000, Fig. S3<br>OTX1B (t33622aep); Fig. S3<br>PITX1 (t5275aep); pred.<br>aristaless-like (t16456aep); pred. |
| OTX1/MA0711.1 | YTAATCCG | 0.0401 | 95.38% | 88.36% | 1.0794 | Homeo domain factors | CnAlx (t16456aep); Smith 2000, Fig. S3<br>OTX1B (t33622aep); Fig. S3<br>PITX1 (t5275aep); pred.<br>aristaless-like (t16456aep); pred. |
| OLIG1/MA0826.1 | AACATATGKT | 0.0449 | 30.06% | 20.33% | 1.4786 | Basic helix-loop-helix factors (bHLH) | HyHES (t3617aep); Münder 2013<br>TFE3 (t22195aep)<br>Mad-protein (t34122aep) |

| | | | | | | | |
|---|---|---|---|---|---|---|---|
| NFATC3/MA0625.1 | WTTTTCCATT | 0.0449 | 98.84% | 93.98% | 1.0517 | Rel homology region (RHR) factors | |
| TEAD3/MA0808.1 | ACATTCCA | 0.0449 | 55.49% | 44.09% | 1.2586 | TEA domain factors | |
| Deaf1/MA0185.1 | TTCGKS | 0.0449 | 78.61% | 68.27% | 1.1515 | SAND domain factors | |
| HNF1B/MA0153.2 | GTTAATNATTAAY | 0.0449 | 61.27% | 49.99% | 1.2256 | Homeo domain factors | CnAlx (t16456aep); Smith 2000, Fig. S3<br>OTX1B (t33622aep); Fig. S3<br>PITX1 (t5275aep); pred.<br>aristaless-like (t16456aep); pred. |
| NR4A2/MA0160.1 | AAGGTCAC | 0.0449 | 79.77% | 69.94% | 1.1405 | Nuclear receptors with C4 zinc fingers | |
| RBPJ/MA1116.1 | BSTGGGAANN | 0.0451 | 57.80% | 46.86% | 1.2335 | Rel homology region (RHR) factors | |

### Motifs enriched in the promoters of downregulated genes that recovered by 6 hours post-treatment

| Motif ID | Motif Sequence | FDR | Frequency in NR Peaks (n=98) | Frequency in non-NR Peaks (n=71913) | Fold Enrichment | Binding TF Class | Potential NR Regulators |
|---|---|---|---|---|---|---|---|
| FOXP1/MA0481.2 | NDGTAAACAGDN | 0.0489 | 98.98% | 88.24% | 1.1217 | Fork head / winged helix factors | FOXI1 (t9145aep); Fig. S3 |
| NKX3-2/MA0122.2 | RCCACTTAA | 0.0489 | 93.88% | 79.75% | 1.1772 | Homeo domain factors | |

### Motifs enriched in the promoters of downregulated genes that remained downregulated by 6 hours post-treatment

| Motif ID | Motif Sequence | FDR | Frequency in NR Peaks (n=214) | Frequency in non-NR Peaks (n=73172) | Fold Enrichment | Binding TF Class | Potential NR Regulators |
|---|---|---|---|---|---|---|---|
| ZIC4/MA0751.1 | GACCCCCCGCTGYGH | 0.0149 | 7.01% | 1.92% | 3.6510 | C2H2 zinc finger factors | Zinc finger protein 26 like (t11591aep); pred. |
| Mafb/MA0117.2 | AAADTGCTGACD | 0.0237 | 68.69% | 55.64% | 1.2345 | Basic leucine zipper factors (bZIP) | |
| lin-14/MA0261.1 | GAACAC | 0.0237 | 87.85% | 77.40% | 1.1350 | Unknown | |
| Pax6/MA0069.1 | TTCACGCWTGANTT | 0.0237 | 37.38% | 25.59% | 1.4607 | Paired box factors | |
| POU4F3/MA0791.1 | ATGMATAATTAATGAG | 0.0348 | 82.71% | 72.22% | 1.1453 | Homeo domain factors | IRX6-2 (t16018aep)<br>Prdl-b (t21636aep); Gauchat 1998 |
| Deaf1/MA0185.1 | TTCGKS | 0.0425 | 77.10% | 66.21% | 1.1645 | SAND domain factors | |

### Motifs enriched in the promoters of upregulated genes that recovered by 3 hours post-treatment

| Motif ID | Motif Sequence | FDR | Frequency in NR Peaks (n=76) | Frequency in non-NR Peaks (n=68746) | Fold Enrichment | Binding TF Class | Potential NR Regulators |
|---|---|---|---|---|---|---|---|
| IRF9/MA0653.1 | AACGAAACCGAAACT | 0.0131 | 7.89% | 0.71% | 11.1127 | Tryptophan cluster factors | |

#### Motifs enriched in the promoters of upregulated genes that recovered by 6 hours post-treatment

| Motif ID | Motif Sequence | FDR | Frequency in NR Peaks (n=53) | Frequency in non-NR Peaks (n=71510) | Fold Enrichment | Binding TF Class | Potential NR Regulators |
|---|---|---|---|---|---|---|---|
| Atf1/MA0604.1 | RTGACGTA | 0.0143 | 90.57% | 65.01% | 1.3932 | Basic leucine zipper factors (bZIP) | |
| Vsx2/MA0180.1 | KTTAATTAG | 0.0143 | 83.02% | 58.02% | 1.4309 | Homeo domain factors | |
| GMEB2/MA0862.1 | TTACGTAA | 0.0143 | 90.57% | 67.86% | 1.3347 | SAND domain factors | |
| EcR::usp/MA0534.1 | VAGTTCATTGAMCTT | 0.0143 | 45.28% | 22.33% | 2.0278 | Nuclear receptors with C4 zinc fingers | |
| Crx/MA0467.1 | AAGRGGATTAG | 0.0212 | 69.81% | 46.17% | 1.5120 | Homeo domain factors | |
| Creb3l2/MA0608.1 | GCCACGTGT | 0.0212 | 26.42% | 9.77% | 2.7042 | Basic leucine zipper factors (bZIP) | |
| HOXC13/MA0907.1 | KCTCGTAAAAH | 0.0451 | 77.36% | 56.16% | 1.3775 | Homeo domain factors | |
| PAX7/MA0680.1 | TAATCGATTA | 0.0459 | 60.38% | 38.87% | 1.5534 | Paired box factors | |

#### Motifs enriched in the promoters of upregulated genes that remained upregulated at 6 hours post-treatment

| Motif ID | Motif Sequence | FDR | Frequency in NR Peaks (n=78) | Frequency in non-NR Peaks (n=72007) | Fold Enrichment | Binding TF Class | Potential NR Regulators |
|---|---|---|---|---|---|---|---|
| Crem/MA0609.1 | KATGACGTAA | 0.0091 | 64.10% | 39.77% | 1.6118 | Basic leucine zipper factors (bZIP) | |
