## Supplemental Table S4 for "Differential gene regulation in DAPT-treated Hydra reveals molecular pathways dependent on Notch signalling during interstitial cell differentiation and formation of the oral-aboral axis in *Hydra*"

### Downregulated transcription factors that recovered by 3 hours post-treatment

| Trinity ID | AEP ID | Gene Model | log2FC 0h | log2FC 3h | Log2FC 6h | Pubmed Hit | NCBI Accession | NCBI Description | Short Name | DNA Binding Domain |
|---|---|---|---|---|---|---|---|---|---|---|
| TRINITY_DN7608_c0_g1 | t22195aep | Sc4wPfr_552.g10536.t1 | -0.326 | 0 | 0 | NA | CDG71838.1 | Hydra vulgaris Microphthalmia-associated transcription factor [Hydra vulgaris] | TFE3 | bHLH |
| TRINITY_DN10125_c0_g1 | t26873aep | Sc4wPfr_1909.g11470.t1 | -0.641 | 0 | 0 | NA | XP_012558352.1 | PREDICTED: zinc finger protein 37-like isoform X1 [Hydra vulgaris] | ZSC31 | zf-C2H2 |
| TRINITY_DN14709_c0_g1 | t5528aep | Sc4wPfr_59.2.g12567.t1 | -0.925 | 0 | 0 | NA | XP_002154370.1 | PREDICTED: transcription factor Sox-19a-like [Hydra vulgaris] | HySox19a | HMG box |
| TRINITY_DN5359_c0_g1 | t23172aep | Sc4wPfr_297.g13156.t1 | -1.507 | 0 | 0 | NA | XP_012555836.1 | PREDICTED: uncharacterized protein LOC101236863 [Hydra vulgaris], similar to SoxB3 Hydractinia echinata | Sox14 | HMG box |
| TRINITY_DN4294_c0_g1 | t20709aep | Sc4wPfr_237.2.g16165.t1 | -1.072 | 0 | 0 | NA | CDG67849.1 | Hydra vulgaris Zinc finger protein ZIC 5, partial [Hydra vulgaris] | Zic5-like | zf-C2H2 |
| TRINITY_DN37877_c0_g1 | t26616aep | Sc4wPfr_14.g1768.t2 | -1.427 | 0 | 0 | NA | CDG72115.1 | Hydra vulgaris Putative transcription factor Ovo-like 1, partial [Hydra vulgaris] | NA | zf-C2H2 |
| TRINITY_DN5649_c0_g1 | t16456aep | Sc4wPfr_654.g18227.t1 | -1.160 | 0 | 0 | NA | AAG03082.1 | aristaless-like protein [Hydra vulgaris] | CnAlx (Smith, 2000) | Homeobox |
| TRINITY_DN3178_c1_g1 | t17964aep | Sc4wPfr_68.g2328.t1 | -1.605 | 0 | 0 | NA | XP_012567188.1 | PREDICTED: uncharacterized protein LOC105851038 [Hydra vulgaris] | Jun | bZIP |

| | | | | | | | | | | |
|---|---|---|---|---|---|---|---|---|---|---|
| TRINITY_DN859_c0_g1 | t11826aep | Sc4wPfr_319.g27364.t1 | -0.397 | 0 | 0 | NA | CDG67153.1 | Hydra vulgaris Transcription factor 7-like 2 [Hydra vulgaris] | TF7-like 2 | HMG box |
| TRINITY_DN1402_c1_g1 | t34122aep | Sc4wPfr_215.1.g29578.t1 | -1.010 | 0 | 0 | NA | CDG70360.1 | Hydra vulgaris Max dimerization protein 1 [Hydra vulgaris] | Mad-protein | bHLH |
| TRINITY_DN5643_c0_g1 | t19720aep | Sc4wPfr_396.g3075.t3 | -0.481 | 0 | 0 | NA | XM_012710796.1 | PREDICTED: Hydra vulgaris forkhead box protein P1-B-like (LOC100202406), transcript variant X3, mRNA | FoxP1 | Forkhead |
| TRINITY_DN7386_c0_g1 | t3617aep | Sc4wPfr_338.1.g31632.t1 | -1.036 | 0 | 0 | NA | XM_004207957.2 | Hydra vulgaris Transcription factor HES-2 [Hydra vulgaris] | HyHes (Münder 2013) | bHLH |
| TRINITY_DN2411_c0_g1 | t29291aep | Sc4wPfr_224.1.g33422.t1 | -1.048 | 0 | 0 | AXP19710.1 | CDG69495.1 | Hydra vulgaris Transcription factor Sp5, partial [Hydra vulgaris] | Sp5 (Vogg, 2019) | zf-C2H2 |
| TRINITY_DN19967_c0_g1 | t33622aep | Sc4wPfr_224.1.g33440.t1 | -0.938 | 0 | 0 | NA | QCF59210.1 | homeobox transcription factor Otx1 [Hydra vulgaris] | OTX1B | Homeobox |
| TRINITY_DN14675_c0_g1 | t5275aep | Sc4wPfr_390.g5621.t1 | -1.705 | 0 | 0 | NA | XP_002164986.2 | PREDICTED: pituitary homeobox 1-like [Hydra vulgaris] | PITX1 | Homeobox |

### Downregulated transcription factors that recovered by 6 hours post-treatment

| Trinity ID | AEP ID | Gene Model | log2FC 0h | log2FC 3h | Log2FC 6h | Pubmed Hit | NCBI Accession | NCBI Description | Short Name | DNA Binding Domain |
|---|---|---|---|---|---|---|---|---|---|---|
| TRINITY_DN18625_c0_g1 | t9145aep | Sc4wPfr_802.g11764.t1 | -2.224 | -2.070 | 0.000 | NA | XP_004207988.1 | PREDICTED: forkhead box protein I1c-like [Hydra vulgaris] | FOXI1 | Forkhead |
| TRINITY_DN3947_c0_g1 | t26993aep | Sc4wPfr_326.g15655.t1 | -0.932 | -1.103 | 0.000 | NA | CDG68553.1 | Hydra vulgaris Fez family zinc finger protein 2, partial [Hydra vulgaris] | FEZ2 | zf-C2H2 |
| TRINITY_DN5602_c0_g1 | t23837aep | Sc4wPfr_362.g23666.t1 | -1.659 | -1.409 | 0.000 | NA | XP_012563508.1 | PREDICTED: transcription factor Sox-21-B-like [Hydra vulgaris] | SOX21B | HMG_box |

| | | | | | | | | | | |
|---|---|---|---|---|---|---|---|---|---|---|
| TRINITY_DN6990_c0_g1 | t11335aep | Sc4wPfr_287.g9045.t1 | -1.255 | -0.753 | 0.000 | NA | XP_002158636.1 | PREDICTED: POU domain, class 4, transcription factor 2-like isoform X2 [Hydra vulgaris] | HyPOU4TF-2 (Siebert, 2019) | Pou |

### Downregulated transcription factors that remained downregulated by 6 hours post-treatment

| Trinity ID | AEP ID | Gene Model | log2FC 0h | log2FC 3h | Log2FC 6h | Pubmed Hit | NCBI Accession | NCBI Description | Short Name | DNA Binding Domain |
|---|---|---|---|---|---|---|---|---|---|---|
| TRINITY_DN3014_c0_g1 | t16018aep | Sc4wPfr_439.g20769.t1 | -0.816 | -0.567 | -0.735 | NA | CDG67528.1 | Hydra vulgaris Iroquois-class homeodomain protein IRX-2 [Hydra vulgaris] | IRX6-2 | Homeobox |
| TRINITY_DN1167_c0_g1 | t12948aep | Sc4wPfr_546.g25835.t1 | -3.863 | -3.103 | -3.221 | NA | CDG72033.1 | Hydra vulgaris Forkhead box protein N1 [Hydra vulgaris] | FoxN4 | Forkhead |
| TRINITY_DN20727_c0_g1 | t11591aep | Sc4wPfr_319.g27294.t1 | -2.000 | -2.061 | -1.572 | NA | XP_002157355.1 | PREDICTED: zinc finger protein 26-like [Hydra vulgaris] | Zinc finger protein 26 like | zf-C2H2 |
| TRINITY_DN2447_c0_g1 | t21636aep | Sc4wPfr_372.g27997.t1 | -3.035 | -2.765 | -1.856 | CAA75669 | XP_002168027.1 | prdl-b protein, partial [Hydra vulgaris] | Prdl-b (Gauchat, 1998) | Homeobox |
| TRINITY_DN10070_c0_g1 | t10853aep | Sc4wPfr_147.g8607.t1 | -1.060 | -1.212 | -0.852 | NP_001296673.1 | NM_001309744.1 | PREDICTED: achaete-scute homolog 1a-like [Hydra vulgaris] | achaete-scute 1a-like | HLH |

### Upregulated transcription factors that recovered by 3 hours post-treatment

| Trinity ID | AEP ID | Gene Model | log2FC 0h | log2FC 3h | Log2FC 6h | Pubmed Hit | NCBI Accession | NCBI Description | Short Name | DNA Binding Domain |
|---|---|---|---|---|---|---|---|---|---|---|
| TRINITY_DN1855_c0_g1 | t5966aep | Sc4wPfr_224.1.g33377.t3 | 1.192 | 0 | 0 | NA | XP_012561111.1 | PREDICTED: transcription factor kayak-like [Hydra vulgaris] | NA | bZIP |

### Upregulated transcription factors that recovered by 6 hours post-treatment

| Trinity ID | AEP ID | Gene Model | log2FC 0h | log2FC 3h | Log2FC 6h | Pubmed Hit | NCBI Accession | NCBI Description | Short Name | DNA Binding Domain |
|---|---|---|---|---|---|---|---|---|---|---|
| TRINITY_DN39755_c0_g1 | t14593aep | Sc4wPfr_547.1.g24983.t1 | 0.937 | 0.789 | 0 | NA | XP_004205480.1 | PREDICTED: zinc finger BED domain-containing protein 4-like [Hydra vulgaris] | NA | zf-BED |
