## Supplemental Table S5 for "Differential gene regulation in DAPT-treated Hydra reveals molecular pathways dependent on Notch signalling during interstitial cell differentiation and formation of the oral-aboral axis in *Hydra*"

| AEP-ID | Minimum distance from TS | Identity/conserved domains | RPBJ-frequency | Expression pattern |
|---|---|---|---|---|
| t16456aep | 100 | CnAlx (Smith2000) | 6 | EC battery, EC head |
| t2316aep | 18 | Putative sialic acid acetylcholine esterase | | EN tentacle |
| t29291aep | 26 | Sp5 (Vogg 2019) | 6 | EC, strong in head |
| t15331aep | 204 | Secreted frizzled-related protein; Bert Hobmayer, Innsbruck, Austria, personal communication | 5 | EN foot |
| t34122aep | 19 | Max-dimerisation domain, bHLH | 5 | EC |
| t18488aep | 47 | Shk-domain, pred. toxin | 5 | Nb 4 through nb 8 |
| t19736aep | 50 | Ubiquitin ligase with RING and SH3 | 4 | EC, EN |
| t5275aep | 21 | PITX1, homeobox, Fig. S3 | 4 | EN head |
| t17828aep | 18 | aminopeptidase | 4 | Nb 4 through nb 8 |
| t20080aep | 98 | GPCR | 4 | Mature nematocytes |
| t14454aep | 187 | Small secreted protein with glycine repeats | 3 | Nb 6 |
| t11622aep | 354 | uncharacterised | 3 | EN |
| t20709aep | 13 | ZIC5, Zf C2H2-domain | 3 | neurons |
| t28505aep | 286 | Uncharacterised with collagen binding sites | 3 | Nb 5 |
| t25509aep | 84 | uncharacterised | 3 | Mature nematocytes |
| t25463aep | 83 | uncharacterised | 3 | Mature nematocytes |
| t26247aep | 173 | Solute carrier family | 3 | Nb 4 through n b8 |
| t6693aep | 14 | Spry-domain and SOCS-box protein | 3 | Nb 6 |
| t23166aep | 1470 | uncharacterised | 3 | Mature nematocytes |
| t25163aep | 8 | Helix rich domain | 3 | Nb 4 through nb |
| t33622aep | 114 | OTX1B like, homeobox, Fig. S3 | 3 | EC head |
