## Supplementary material for "Differential gene regulation in DAPT-treated Hydra reveals molecular pathways dependent on Notch signalling during interstitial cell differentiation and formation of the oral-aboral axis in *Hydra*": RCode

### Supplementary, R-Code used for data analysis

*Load libraries*

```
library(dplyr)
library(gplots)
library(tibble)
library(Seurat)
library(stringr)
library(rlang)
```

DESeq2 was performed on the mapping gene raw counts for each time point (0h, 3h, 6h) seperately, thereby comparing DAPT-treated animals with control animals. A cutoff of Padj <= 0,01 was applied. The genes that were differentially expressed (DE) at 0h were selected and merged with their foldchanges at the time points 3h and 6h. This resulted in 831 DE-genes (which we call Notch-regulated genes (NR-genes)), which were then clustered according to their log2foldchanges at the three time points.

*Prepare the list with all 831 NR-genes and their log2foldchanges*

```
# Load the DESeq2 outputs for each time point
DE.0h <- read.table("DESeq2/DE.0h.txt", header = T, sep = "\t", row.names = NULL)
DE.3h <- read.table("DESeq2/DE.3h.txt", header = T, sep = "\t", row.names = NULL)
DE.6h <- read.table("DESeq2/DE.6h.txt", header = T, sep = "\t", row.names = NULL)

# Select only the Trinity gene ID and Log2Foldchange from each DESeq2 output file
# and change column names
DE.0h <- DE.0h %>% select(row.names, log2FoldChange)
colnames(DE.0h) <- c("Trinity.ID", "log2Foldchange.0h")
DE.3h <- DE.3h %>% select(row.names, log2FoldChange)
colnames(DE.3h) <- c("Trinity.ID", "log2Foldchange.3h")
DE.6h <- DE.6h %>% select(row.names, log2FoldChange)
colnames(DE.6h) <- c("Trinity.ID", "log2Foldchange.6h")

# Merge DE.0h with DE.3h and DE.6h
DE.genes <- merge(DE.0h, DE.3h, by = "Trinity.ID", all.x = TRUE)
DE.genes <- merge(DE.genes, DE.6h, by = "Trinity.ID", all.x = TRUE)

# Remove previous files
rm(DE.0h)
rm(DE.3h)
rm(DE.6h)
```

*Cluster the NR-genes according to their log2foldchanges and plot them in a heatmap*

```
# Convert the column Trinity.ID into rownames in order to obtain a numeric matrix
# then remove NA
DE.genes <- column_to_rownames(DE.genes, var = "Trinity.ID")
DE.genes[is.na(DE.genes)] = 0

# Calculate the distances and build the clusters
DE_dist <- dist(DE.genes, method = "euclidian")
DE_hclust <- hclust(DE_dist, method = "ward.D")

# Define the colors and breaks for the heatmap
my_palette <- colorRampPalette(c("red", "white", "blue")) (n=51)
breaks <- seq(from=min(DE.genes), to=abs(min(DE.genes)), length.out = 52)

# Plot and save the Heatmap
pdf("Heatmap foldchanges NR genes.pdf")
heatmap.2(as.matrix(DE.genes), trace = "none", cexCol=1, cexRow = 0.75, dendrogram = "row", Colv = F,
        symkey=F, scale = "none", col=my_palette, labRow = FALSE, srtCol=0, adjCol=c(0.5,0),
        breaks=breaks, Rowv=as.dendrogram(DE_hclust), main = "HM foldchanges NR-genes")
dev.off()

rm(DE_hclust)
rm(DE_dist)
```

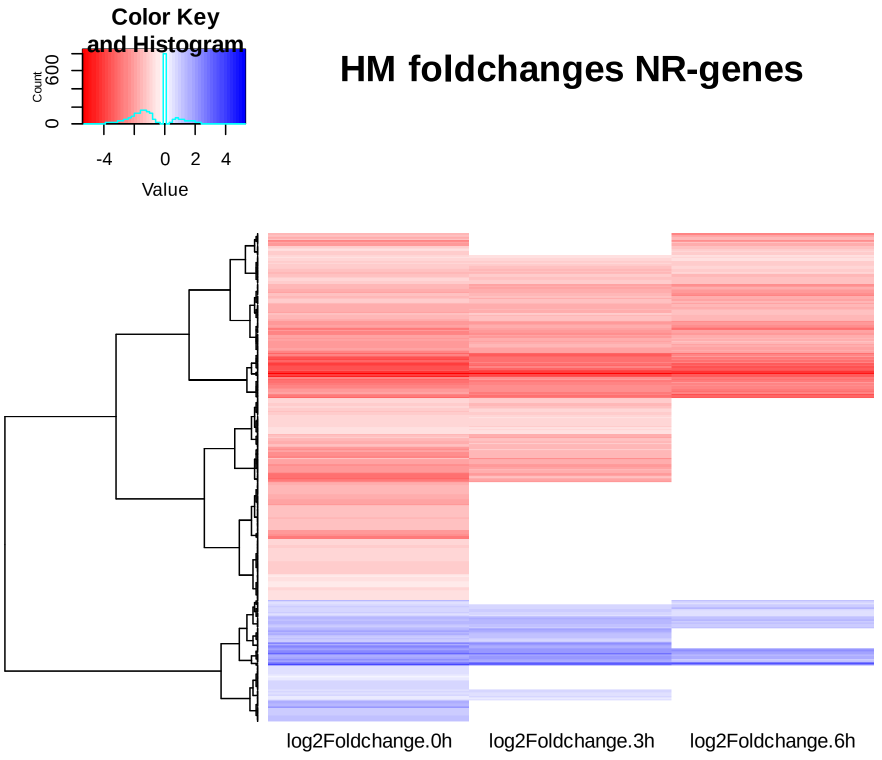

The heatmap shows two main clusters, one includes the down-regulated genes and the other the up-regulated genes. Within each cluster there are four sub-clusters.

*To get the number of genes and the genes within each group*

```
# Filter according to the log2foldchanges at the three time points
# d=down, u=up
# 0=differentially expressed at 0h, normal at 3h
# 03=differentially expressed at 0h and 3h, normal at 6h
# 036=differentially expressed at 0h, 3h and 6h
# 06=differentially expressed at 0h and 6h, normal at 3h
d0 <- subset(DE.genes, DE.genes$log2Foldchange.0h<0 & DE.genes$log2Foldchange.3h == 0 &
DE.genes$log2Foldchange.6h == 0)
rm(d0)
d03 <- subset(DE.genes, DE.genes$log2Foldchange.0h<0 & DE.genes$log2Foldchange.3h != 0 &
DE.genes$log2Foldchange.6h == 0)
rm(d03)
d036 <- subset(DE.genes, DE.genes$log2Foldchange.0h<0 & DE.genes$log2Foldchange.3h != 0 &
DE.genes$log2Foldchange.6h != 0)
rm(d036)
d06 <- subset(DE.genes, DE.genes$log2Foldchange.0h<0 & DE.genes$log2Foldchange.3h == 0 &
DE.genes$log2Foldchange.6h != 0)
rm(d06)
u0 <- subset(DE.genes, DE.genes$log2Foldchange.0h>0 & DE.genes$log2Foldchange.3h == 0 &
DE.genes$log2Foldchange.6h == 0)
rm(u0)
u03 <- subset(DE.genes, DE.genes$log2Foldchange.0h>0 & DE.genes$log2Foldchange.3h != 0 &
DE.genes$log2Foldchange.6h == 0)
rm(u03)
u036 <- subset(DE.genes, DE.genes$log2Foldchange.0h>0 & DE.genes$log2Foldchange.3h != 0 &
DE.genes$log2Foldchange.6h != 0)
rm(u036)
u06 <- subset(DE.genes, DE.genes$log2Foldchange.0h>0 & DE.genes$log2Foldchange.3h == 0 &
DE.genes$log2Foldchange.6h != 0)
rm(u06)
```

All 831 NR-genes were then blasted to the AEP transcriptome (Siebert et al., 2019) using local nt blast. For 666 NR genes (80%) an AEP reference was found. Data from Hydra single cell expression analysis (Siebert et al., 2019) were used to get insights into the spatial expression and cell state information of these NR genes.

*Prepare the file for the local blast by adding the SuperTranscript sequence to each Trinity.gene*

```
# Convert row names into a column
DE.genes <- rownames_to_column(DE.genes, var = "Trinity.ID")

# Load file with Trinity sequences and add the SuperTranscript sequences to DE.genes
sequences <- read.table("Trinity.sequences/Trinity.sequence.txt", header = TRUE)
DE.genes <- merge(DE.genes, sequences, by = "Trinity.ID")
rm(sequences)
```

*Load the results from the local blast and add them to DE.genes*

```
# Load result local blast DE.genes-to-AEP
DEG.AEP <- read.table("Blast/Blast.DEG.AEP.txt", header = TRUE)

# Select only the Trinity.ID (V1) and the AEP.ID (V2)
DEG.AEP <- DEG.AEP %>% select(V1, V2)
colnames(DEG.AEP) <- c("Trinity.ID", "AEP.ID")

# Add AEP.ID to DE.genes
DE.genes <- merge(DE.genes, DEG.AEP, by = "Trinity.ID", all.x = TRUE)
rm(DEG.AEP)
```

The NR-genes were all annotated manually by NCBI Blast search. Therefore, the SuperTranscript sequence was used.

*Load the annotations and add them to DE.genes*

```
# Add annotation to DE.genes
Annotation <- read.table("Annotation/Manual annotation.txt", header = TRUE, sep = "\t")
DE.genes <- merge(DE.genes, Annotation, by = "Trinity.ID", all.x = TRUE)
rm(Annotation)
```

The recent single cell analysis (Siebert et al., 2019) yielded multiple Seurat objects including the whole transcriptome dataset, containing all clustered cell states and the interstitial Seurat object, containing only derivatives of the interstitial cell lineage. Expression information for NR-genes was isolated from both mentioned Seurat objects.

*Isolate expression information from the whole-transcriptome Seurat object*

```
# Load the Seurat object for the whole transcriptome-mapped dataset
# and calculate the average expression
S <- readRDS("Seurat/wt_Seurat.rds")
S <- AverageExpression(S)
S <- as.data.frame(S)

# Remove the cluster "doublets" from the Seurat object
S$db <- NULL

# Convert rownames into column and split AEP annotation from AEP ID
DEG.S <- rownames_to_column(as.data.frame(S), var = "AEP.annotation")
ID <- str_split_fixed(DEG.S$AEP.annotation, "\\|", 2)
colnames(ID) <- c("AEP.ID", "Annotation")
ID <- as.data.frame(ID)
rownames(DEG.S) <- ID$AEP.ID
rm(ID)
rm(S)

# Isolate the NR-genes for which an AEP reference was found
DE.genes.AEP <- DE.genes %>% filter(AEP.ID != "NA")
DEG <- as.character(DE.genes.AEP$AEP.ID)

# Select the expression information from the Seurat object for these NR-genes and remove NA
DE.genes.S <- DEG.S[DEG,]
DE.genes.S <- data.matrix(DE.genes.S[,2:38], rownames.force = NA)
DE.genes.S[is.nan(DE.genes.S)] = 0

# Average expression was next scaled from 0 to 1
DE.genes.S <- t(apply(DE.genes.S,1,function(x)(x-min(x))/(max(x)-min(x))))
DE.genes.S[is.na(DE.genes.S)] = 0
rm(DEG)
rm(DEG.S)
rm(DE.genes.AEP)
```

*NR-genes with an AEP reference were next clustered according to their whole transcriptome cell state expression information*

```
# Calculate the distances and build the clusters
DEG.dist <- dist(DE.genes.S, method = "euclidian")
DEG.hclust <- hclust(DEG.dist, method = "ward.D")

# Plot the clusters, select the main 8 clusters, and save plot
pdf("Cluster analysis DE.genes.S.wt.pdf")
plot(DEG.hclust, labels = F, main = "Clustering NR Notch genes.wt", hang = F)
rect.hclust(DEG.hclust, k = 8)
dev.off()
```

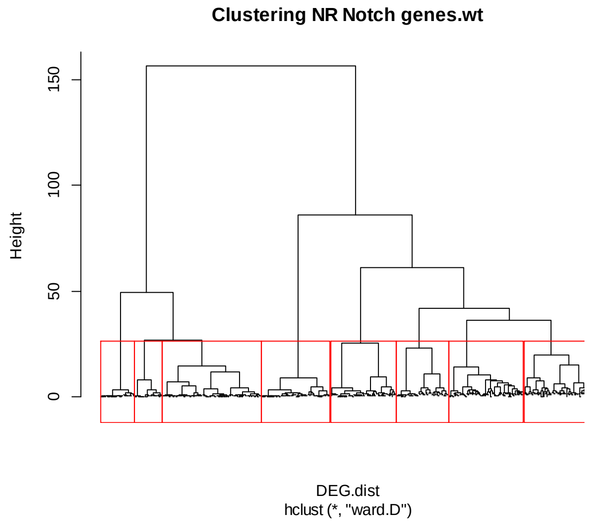


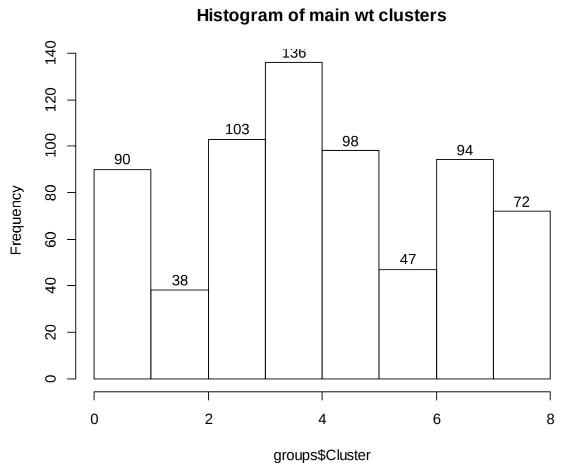


*The genes were saved according to their cluster*

```
# Save the genes according to the clusters, which are named groups
groups <- cutree(DEG.hclust, k = 8)
write.table(groups, "groupClusters.DE.genes.wt.txt", sep = “\t”)
groups <- as.data.frame(groups)
colnames(groups) <- "Cluster"
groups <- rownames_to_column(groups, var = "AEP.ID")

# Add the group number to DE.genes, the clusters will then be named manually
DE.genes <- merge(DE.genes, groups, by = "AEP.ID", all.x = TRUE)

# Draw and save a histogram with the numbers of genes in each group
pdf("Histogram 8 main clusters DE.genes.wt.pdf")
hist(groups$Cluster, labels = TRUE, breaks = c(0,1,2,3,4,5,6,7,8), main = "Histogram of main wt clusters")
dev.off()
```

*Draw a heatmap with the clustered NR-genes*

```
# Draw and save a heatmap with the clustered NR genes
pdf("Heatmap NR genes_wt, dist eucl., clust ward.d.pdf", bg = "white")
heatmap.2(as.matrix(DE.genes.S), trace = "none", cexCol=1, margins = c(13.5,1), dendrogram = "row",
        labRow=FALSE, symkey=F, scale = "none", col=my_palette, srtCol=90, adjCol=c(1,0),
        breaks=breaks, Rowv=as.dendrogram(DEG.hclust),
        RowSideColors = as.character(groups$Cluster), main = "clustering NR genes.wt")
dev.off()

rm(DE_hclust)
rm(DE_dist)
rm(groups)
rm(DE.genes.S)
```

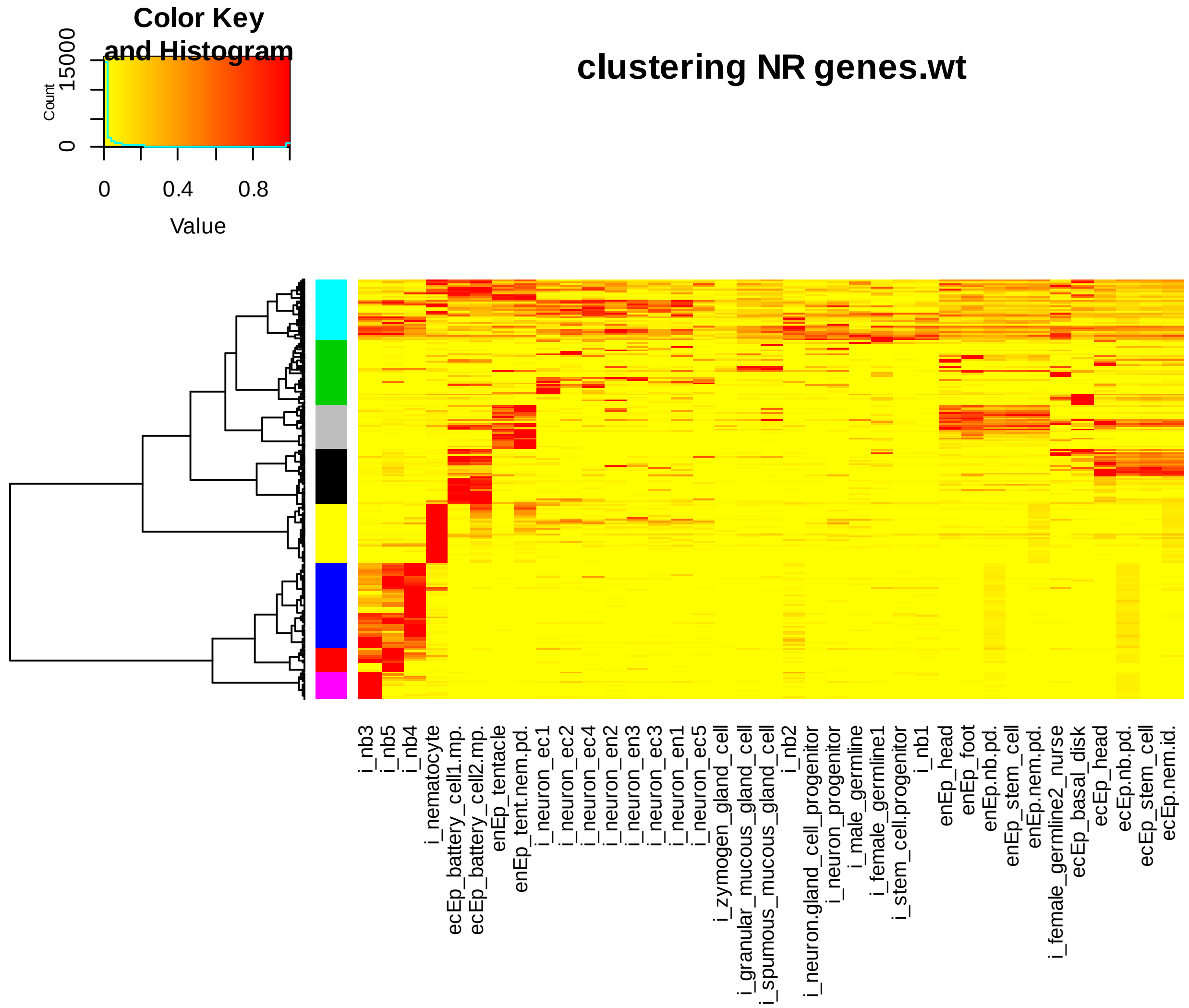

Next, genes in nb and nematocyte clusters were isolated and clustered again according to their cell state expression information from the interstitial cell dataset (IC-Seurat).
(the R-Code in the next steps is very similar to the code that is described above)

```
# Load the Seurat object for the interstitial cell-mapped dataset
# and calculate the average expression
S <- readRDS("Seurat/IC_Seurat.rds")
S <- AverageExpression(S)
S <- as.data.frame(S)

# Save the Seurat as a matrix and use it in order to save loading time
write.table(S, "S_IC_averageExp.txt")
S.IC <- read.table("Files/S_IC_averageExp.txt", header = TRUE, sep = "\t")
rm(S)

# Isolate nematoblast/nematocyte genes from the NR-genes with an AEP reference.
# (clusters 2, 4, 6 and 7 from wt (whole-transcriptome) Heatmap)
DE.genes.AEP <- DE.genes %>% filter(AEP.ID != "NA")
DE.nb <- DE.genes.AEP %>% filter(Cluster == "2" | Cluster == "4" | Cluster == "6" | Cluster == "7")

# Average expression was next scaled from 0 to 1
DEG <- as.character(DE.nb$AEP.ID)
DE.nb.S <- S.IC[DEG,]
DE.nb.S <- data.matrix(DE.nb.S, rownames.force = NA)
DE.nb.S[is.nan(DE.nb.S)] = 0
DE.nb.S <- t(apply(DE.nb.S,1,function(x)(x-min(x))/(max(x)-min(x))))
DE.nb.S[is.na(DE.nb.S)] = 0
rm(DEG)
rm(DE.genes.AEP)
rm(DE.nb)
rm(S.IC)

# Clustering Notch regulated nematoblast/nematocyte genes
DEG.dist <- dist(DE.nb.S, method = "euclidian")
DEG.hclust <- hclust(DEG.dist, method = "ward.D")

pdf("Cluster analysis DE.nb.S.IC.pdf")
plot(DEG.hclust, labels = F, main = "Clustering NR nematoblast genes.IC", hang = F)
rect.hclust(DEG.hclust, k = 5)
dev.off()

# Save the genes according to the clusters, which are named groups
groups <- cutree(DEG.hclust, k = 5)
groups <- as.data.frame(groups)
colnames(groups) <- "Cluster.nb"
groups <- rownames_to_column(groups, var = "AEP.ID")
```

```
# Add the cluster number to DE.genes, the clusters will then be named manually
DE.genes <- merge(DE.genes, groups, by = "AEP.ID", all.x = TRUE)
write.table(DE.genes, "DE.genes.txt", sep = "\t")

# Draw a histogram with the numbers of genes in each group
pdf("Histogram 5 main clusters DE.nb.IC.pdf")
hist(groups$Cluster.nb, labels = TRUE, breaks = c(0,1,2,3,4,5), main = "Histogram of main IC clusters")
dev.off()

# Draw a heatmap with the clustered NR nb genes
pdf("Heatmap NR nb genes_IC, dist eucl., clust ward.d.pdf", bg = "white")
heatmap.2(as.matrix(DE.nb.S), trace = "none", cexCol=1, margins = c(13.5,1), dendrogram = "row",
        labRow=FALSE, symkey=F, scale = "none", col=my_palette, srtCol=90, adjCol=c(1,0),
        breaks=breaks, Rowv=as.dendrogram(DEG.hclust),
        RowSideColors = as.character(groups$Cluster.nb), main = "clustering NR nb genes.IC")
dev.off()
```

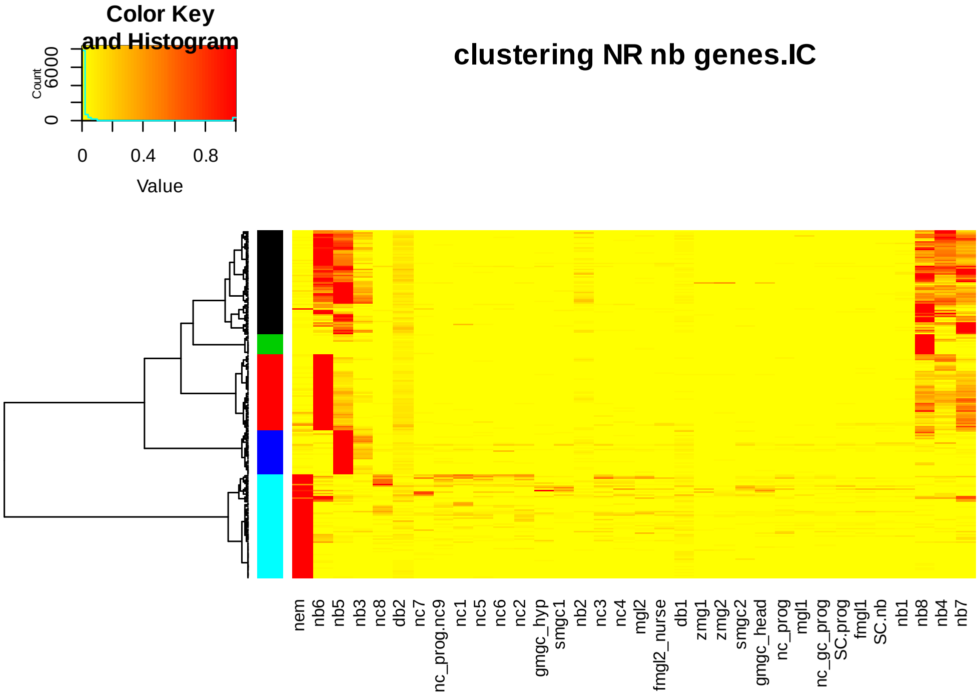


The genes are mainly, if not only expressed in nematoblasts and nematocytes, therefore, a new heatmap was made with only nematoblast/nematocyte cell states.

```
DE.genes$Cluster.nb <- NULL
DE.nb.S <- as.data.frame(DE.nb.S)
DE.nb.S <- rownames_to_column(DE.nb.S, var = "AEP.ID")

DE.nb.S <- DE.nb.S %>% select(SC.nb, nb1, nb2, nb3, nb4, nb5, nb6, nb7, nb8, nem)
```

```
DEG.dist <- dist(DE.nb.S, method = "euclidian")
DEG.hclust <- hclust(DEG.dist, method = "ward.D")

pdf("Cluster analysis DE.nb.S.IC.pdf")
plot(DEG.hclust, labels = F, main = "Clustering NR nematoblast genes.IC", hang = F)
rect.hclust(DEG.hclust, k = 5)
dev.off()

groups <- cutree(DEG.hclust, k = 5)
groups <- as.data.frame(groups)
colnames(groups) <- "Cluster.nb"
groups <- rownames_to_column(groups, var = "AEP.ID")

DE.genes$Cluster.nb <- NULL
DE.genes <- merge(DE.genes, groups, by = "AEP.ID", all.x = TRUE)
write.table(DE.genes, "DE.genes.txt", sep = "\t")

pdf("Histogram 5 main clusters DE.nb.IC.pdf")
hist(groups$Cluster.nb, labels = TRUE, breaks = c(0,1,2,3,4,5), main = "Histogram of main IC clusters")
dev.off()

pdf("Heatmap NR nb genes_IC, dist eucl., clust ward.d.pdf", bg = "white")
heatmap.2(as.matrix(DE.nb.S), trace = "none", cexCol=1, margins = c(13.5,1), dendrogram = "row",
        labRow=FALSE, symkey=F, scale = "none", col=my_palette, srtCol=90, adjCol=c(1,0),
        breaks=breaks, Rowv=as.dendrogram(DEG.hclust),
        RowSideColors = as.character(groups$Cluster.nb), main = "clustering NR nb genes.IC")
dev.off()
```

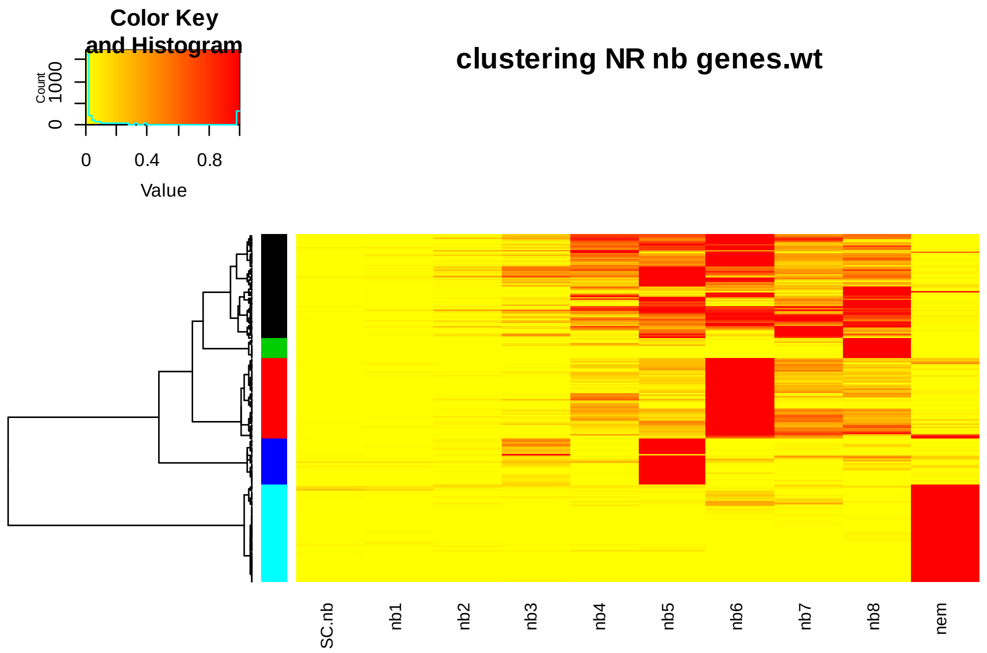

*Isolate non-nematoblast genes from the wt-heatmap and cluster again according to their expression in only epithelial cells*

```
# Load the whole-transcriptome Seurat object
wt.S <- read.table("Files/DE.genes.wt.scaledAverExp.txt", header = TRUE, sep = "\t")

# Isolate the genes in the black, grey, green and cyan cluster, representing cluster 1, 3, 5 and 8
DE.epi <- DE.genes %>% filter(Cluster == 1 | Cluster == 3 | Cluster == 5 | Cluster == 8)
DE.epi <- as.character(DE.epi$AEP.ID)

# Isolate the expression pattern of epithelial genes from the whole transcriptome Seurat
DE.epi.S <- wt.S[DE.epi,]

# Select only the epithelial cell states
DE.epi.S <- DE.epi.S %>% select(enEp_stem_cell, ecEp_stem_cell, enEp_head, ecEp_head, enEp_foot,
ecEp_battery_cell2.mp., enEp_tentacle, ecEp_basal_disk, ecEp_battery_cell1.mp.)

# Cluster epithelial genes according to their expression pattern
DEG.dist <- dist(DE.epi.S, method = "euclidian")
DEG.hclust <- hclust(DEG.dist, method = "ward.D")

# Cluster NR epithelial genes according to their cell state information and select the main 8 clusters
pdf("Cluster analysis DE.epi.S.wt.pdf")
plot(DEG.hclust, labels = F, main = "Clustering NR epithelial genes.wt", hang = F)
rect.hclust(DEG.hclust, k = 8)
dev.off()

# Save the genes according to the clusters, which are named groups
groups <- cutree(DEG.hclust, k = 8)
groups <- as.data.frame(groups)
colnames(groups) <- "Cluster.epi"
groups <- rownames_to_column(groups, var = "AEP.ID")

# Add the cluster number to DE.genes, the clusters will then be named manually
DE.genes <- merge(DE.genes, groups, by = "AEP.ID", all.x = TRUE)
write.table(DE.genes, "DE.genes.txt", sep = "\t")

# Draw a histogram with the numbers of genes in each group
pdf("Histogram 8 main clusters DE.epi.wt.pdf")
hist(groups$Cluster.epi, labels = TRUE, breaks = c(0,1,2,3,4,5,6,7,8), main = "Histogram of main epi.wt
clusters")
dev.off()
```

```
# Draw a heatmap with the clustered NR epi genes
pdf("Heatmap NR epi genes_wt, dist eucl., clust ward.d.pdf", bg = "white")
heatmap.2(as.matrix(DE.epi.S), trace = "none", cexCol=1, margins = c(13.5,1), dendrogram = "both",
        labRow=FALSE, symkey=F, scale = "none", col=my_palette, srtCol=90, adjCol=c(1,0),
        breaks=breaks, Rowv=as.dendrogram(DEG.hclust),
        RowSideColors = as.character(groups$Cluster.epi), main = "clustering NR epi genes.wt")
dev.off()
```

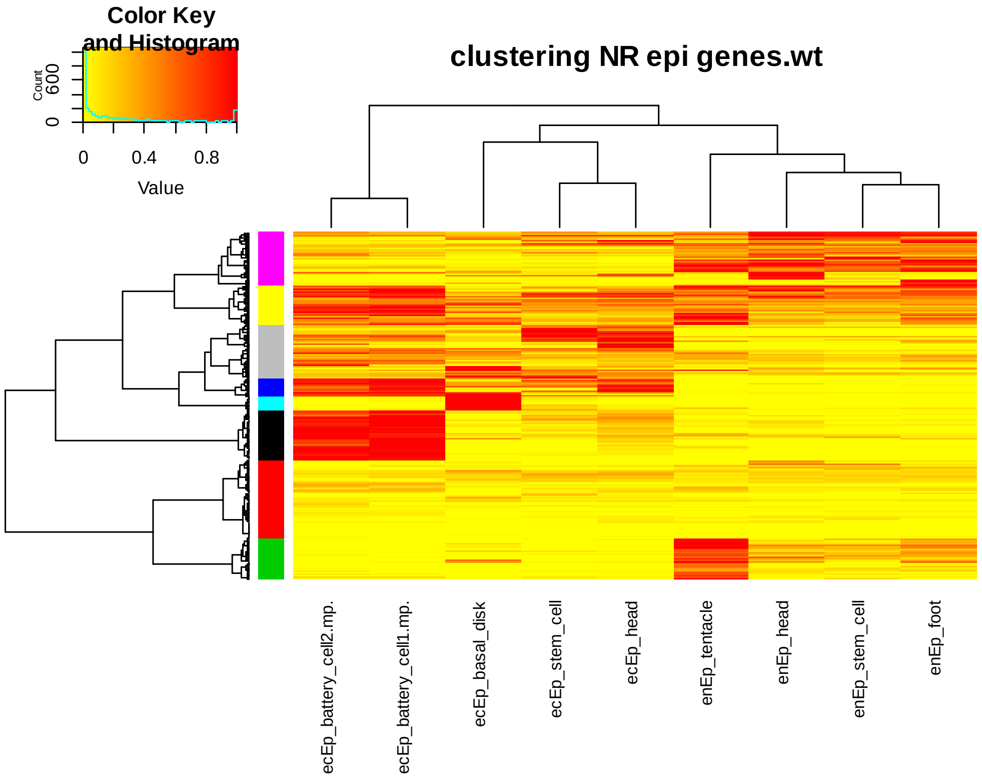


From the epithelial heatmap, genes in the grey and violet clusters were isolated and clustered separately in order to extract the genes that are specifically expressed in ectodermal head-, endodermal head-, and endodermal foot cells. Genes in the red cluster don’t seem to show expression in neither nematoblast cells nor in epithelial cells. Therefore, these genes were isolated and clustered again according to their expression pattern in IC derivatives, except nematoblasts and nematocytes. The same R Code as described above was applied. A complete list of all NR-genes and the cells states in which they are mainly expressed is found in the supplementary “DE.genes”.

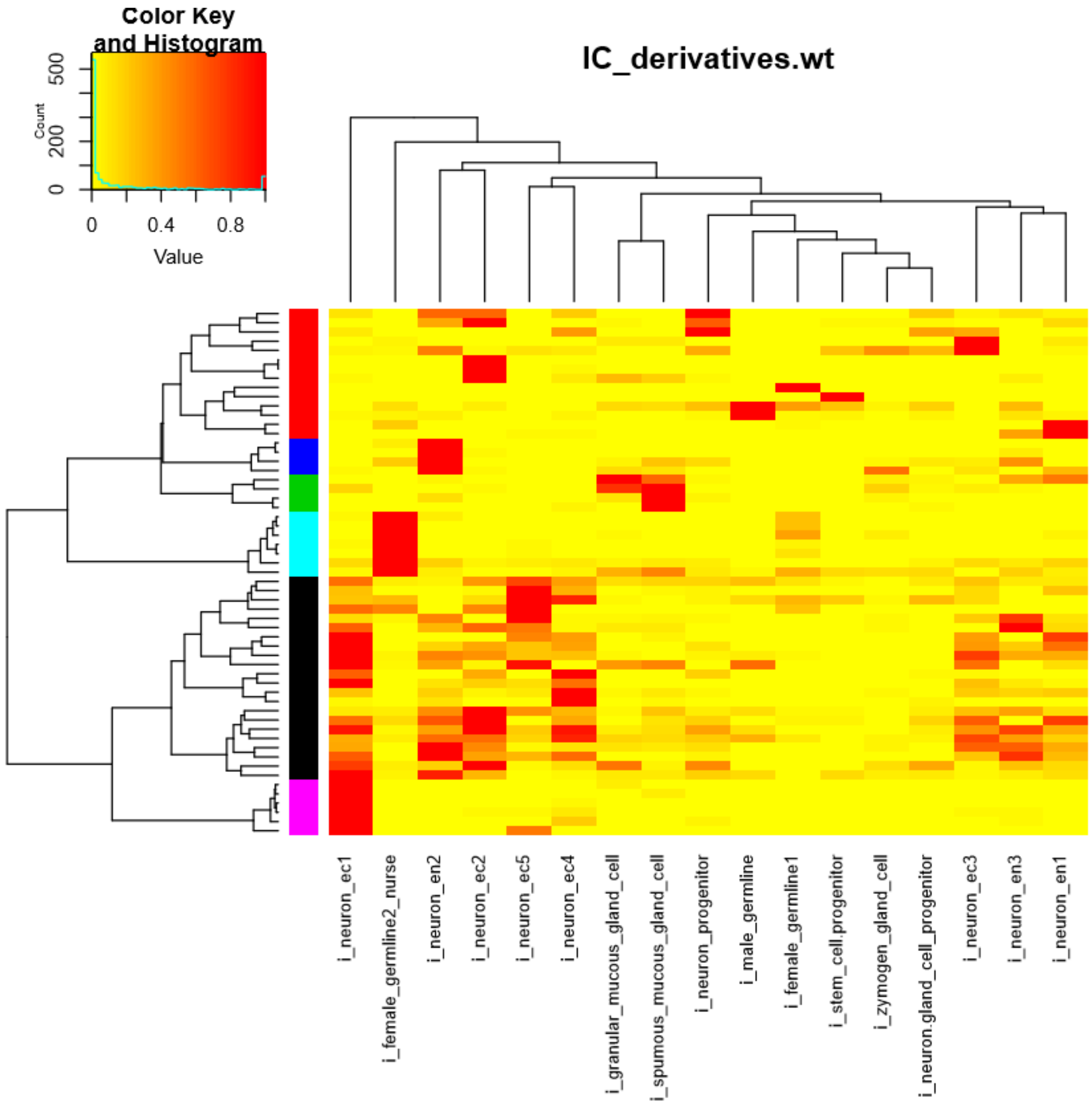
Color Key
and Histogram
Count
500
200
0
0
0.4
0.8
Value
IC_derivatives.wt
i_neuron_ec1
i_female_germline2_nurse
i_neuron_en2
i_neuron_ec2
i_neuron_ec5
i_neuron_ec4
i_granular_mucous_gland_cell
i_spumous_mucous_gland_cell
i_neuron_progenitor
i_male_germline
i_female_germline1
i_stem_cell.progenitor
i_zymogen_gland_cell
i_neuron.gland_cell_progenitor
i_neuron_ec3
i_neuron_en3
i_neuron_en1
