## supplemental Figure S3 for "Differential gene regulation in DAPT-treated Hydra reveals molecular pathways dependent on Notch signalling during interstitial cell differentiation and formation of the oral-aboral axis in *Hydra*"

**Selected Alignments of protein sequences of NR genes**

**1.) Dickkopf**

**t22117aep|DKK3_MOUSE**
TRINITY_DN37863_c0_g1 ORF10 (**HyDKK3|ORF10**)

HyDKK protein References:
1.Hydra Vulgaris Dickkopf 1/2/4-A Protein, (**HYDVU-DKKGuder**)
2.Hydra Magnipapillata Dickkopf 1/2/4-A Protein, (**HYMAG-DKKGuder**)
Reference: An ancient Wnt-Dickkopf antagonism in Hydra. (Guder et al., 2006)
3.Hydra vulgaris Dickkopf-3 related protein, (**HyDKK3-Fedders**)
Reference: A Dickkopf-3-related gene is expressed in differentiating nematocytes in the basal metazoan Hydra. (Fedders et al., 2004)

Alignments:

```
HYDVU-DKKGuder    ~~~~~~~~~~~~~~~~~~~~~~~~~~~~~~~~~~~~~~~~~~~~~~~~~~~~~~~~~~~~    -
HYMAG-DKKGuder    ~~~~~~~~~~~~~~~~~~~~~~~~~~~~~~~~~~~~~~~~~~~~~~~~~~~~~~~~~~~~    -
HyDKK3|ORF10      MYDPHTHIMFYILSTLLIGALVKTGLATDGEF--TLKKDTDESGSMKVKKMIAPGYYSQE   58
HyDKK3-Fedders    ~~~~~MSKLFIIYTFCIFVA-----YAEDKKVPKPVAPSLEKR-GDIQPRMLAPGYYSQE   49

                                                            CRD1 domain
                  CRD1 domain

HYDVU-DKKGuder    ~~~~~~~~~~MRFLAVLLVVAAFVAFSEAESCKKDADCKNGCCVNFLLTKQCNSYVKEGA   50
HYMAG-DKKGuder    ~~~~~~~~~~MRFLAVFLVVAAFVAFSEAGRCNKDADCENGCCVNYLLTKQCNSYVKEGE   50
HyDKK3|ORF10      CNVHKPCPDPTKYCHMFLCVD---CLKENVACTQNGQCCPGTECTY---GRCKKGSSKGA  112
HyDKK3-Fedders    CNAHKACPE-KKYCHLFLCVH---CLKENVACTQNGQCCEG-QCTY---GRCKAGVSEGQ  101
                             45  6fL V       E   C  1  C  G    5     C   v eG

HYDVU-DKKGuder    LCGFRDKFA----------C----------------------------------------   60
HYMAG-DKKGuder    LCGFRDKFA----------C----------------------------------------   60
HyDKK3|ORF10      AGTFCDRQKDCVGP--DLCCVREPAINPVISICKPALDEHQTCGPYNQFRTVYIGGTVQP  170
HyDKK3-Fedders    PGTFCDRHEDCAGEGKAACCVREPAINPHISICKPPLAENMVCGPINFFRNVYVGAQVQK  161
                     F D4            C
```

```
HYDVU-DKKGuder    ---GCEPGLECVKVRGTLTGMVRRCVDNSGSGSLY   92
HYMAG-DKKGuder    ---GCEPGLECVKVRGTLTGMVRRCVDNSGSGSLY   92
HyDKK3|ORF10      VCGPCKQGLVCKQV--GIFGVHEICLPSGGKGKK~  202
HyDKK3-Fedders    ACGPCKQALICKQV--GLFGIHEICMKEDDKKK~~  192
                      C  gL C  V   6 G6   C6   g g
```

Vertebrate Dickkopf molecules consist of two cysteine-rich domains (CRD1 and CRD2), which are separated by a spacer region, diagnostic for grouping of Dkk proteins. CRD2 is necessary and sufficient to repress canonical Wnt signaling by competing with the Wnt-Frizzled complex for binding to the Lrp5/Lrp6 receptor. CRD1 is thought to have a modulating function on CRD2.

HyDKK1-Guder protein only has the CRD2 domain (blue boxes).

HyDKK3|ORF10 and HyDKK-Fedders have CRD2 (blue box) and CRD1 domains (red box).

---

### 2.) Spinalin

**t38568aep**
TRINITY_DN2600_c0_g1ORF15 (**Spin|ORF15**)

**AAC39121.1** spinalin [Hydra vulgaris](Koch et al., 1998)
**ACM79874.1** nematoblast-specific protein nb054-sv9, partial [Hydra vulgaris] (Milde et al., 2009)
**XP_012553808.1** PREDICTED: midasin [Hydra vulgaris]

Alignment of AAC39121.1, ACM79874.1, t38568aep, XP_012553808.1 and Spin|ORF15:

```
t38568aep       MVIAQAALVLLLVAVDARPWGPGCADGSYGYGGCGHHQANGYGGAHHAAGCCNGQAHGGH   60
Spin|ORF15      ------------------------------------------------------------   -
AAC39121.1      MVIAQAALVLLLVAVDARPWGPGCADGSYGYGGCGHHQANGYGGAHHAAGCCNGLAHGGH   60
XP_012553808.1  MVIAQAALVLLLVAVDARPWGPGCADGSYGYGGCGHHQANGYGGAHHAAGCCNGLAHGGH   60
ACM79874.1      ------------------------------------------------------------   -


t38568aep       HGGAYGQAAHHAGGYGGAYGGHHEDHGEDVKVETHGVHHIASHGGIHHEHGGLRGGHHGA  120
Spin|ORF15      --------------------------MVESIMNTAVLEEDIMEHIIHHEHGGLRGGHHGA   34
AAC39121.1      HGGAYGQAAHHAGGYGGAYGGHHEDHGEDVKVETHGVHHIASHGGIHHEHGGLRGGHHGA  120
XP_012553808.1  HGGAYGQAAHHAGGYGGAYGGHHEDHGEDVKVETHGVHHIASHGGIHHEHGGLRGGHHGA  120
ACM79874.1      ------------------------------------------------------------   -


t38568aep       YGGHGAHEAGNFGHVHGGHYGLGGNHYGGSDYGHGHHGGYVHEPCHTPCHTVGGCGGCGG  180
Spin|ORF15      YGGHGAHEAGNFGHVHGGHYGLGGNHYGGSDYGHGHHGGYVHEPCHTPCHTVGGCGGCGG   94
AAC39121.1      YGGHGAHEAGNFGHVHGGHYGLGGNHYGGSDYGHGHHGGYVHEPCHTPCHTVGGCGGCGG  180
XP_012553808.1  YGGHGAHEAGNFGHVHGGHYGLGGNHYGGSDYGHGHHGGYVHEPCHTPCHTVGGCGGCGG  180
ACM79874.1      ------------------------------------------------------------   -

t38568aep       YGHLGGVGGYGAGLGG---VGGLHVYKKETVKGKKVQQVTKAEEENEDDDDDESEDAEAE  237
Spin|ORF15      YGHLGGVGGYGTGLGGVGGVGGLHVYKKETVKGKKVQQVTKAEEENEDDDDDESEDAEAE  154
AAC39121.1      YGHLGGVGGYGTGLGGVGGVGGLHVYKKETVKGKKVQQVTKAEEENEDDDDDESEDAEAE  240
XP_012553808.1  YGHLGGVGGYGTGLGGVGGVGGLHVYKKETVKGKKVQQVTKAEEENEDDDDDESEDAEAE  240
ACM79874.1      ------------------------------------------------------------   -
```

```
t38568aep        SGNSSDEDMPNGAKKSTLVKEKKNEEEKNDKEKPEINMMINSEKKKEDNLPENKSSEKIN  297
Spin|ORF15       SGNSSDEDMPNGAKKSTLVKEKKNGEEKNDKEKPEINMMMNSEKKKEDNLPENKSSEKIN  214
AAC39121.1       SGNSSDEDMPNGGD----------------------------------------------  254
XP_012553808.1   SGNSSDEDMPNGAKKSTLVKEKKNGEEKNDKEKPEINMMMNSEKKKEDNLPENKSSEKIN  300
ACM79874.1       ------EDMPNGAKKSTLVKEKKNGEEKNDKEKPEINMMMNSEKKKEDNLPENKSSEKIN   54
                       EDMPNG

t38568aep        DEKKVVDTKNSSDVKSVDDVSKVEQSKSGDVKVADSKEDVKIKEVIKAEGIKDREDKKIQ  357
Spin|ORF15       DEKKVGDTKNSSDVKSVDDVSKVEQLKSGDAKVVDNKEDVKLKEGVKAEGIKDKEDAKIQ  274
AAC39121.1       ------------------------------------------------------------    -
XP_012553808.1   DEKKVGDTKNSSDVKSVDDVSKVEQLKSGDAKVVDNKEDVKLKEGVKAEGIKDKEDAKIQ  360
ACM79874.1       DEKKVGDTKNSSDVKSVDDVSKVEQLKSGDAKVVDNKEDVKLKEGVKAEGIKDKEDAKIQ  114

t38568aep        DSEEKTIQKTQNKTNQDAGTEKVQSGENKKITAEKDAAEPKTKEDIKLRGAQKAYHKAKE  417
Spin|ORF15       DSEEKTVQKTQNKTNQGAGTEKVQSAENKKITAEKDVAEPKTKEDIKLRGAQKAYHKAKE  334
AAC39121.1       ------------------------------------------------------------    -
XP_012553808.1   DSEEKTVQKTQNKTNQDAGTEKVQSAENKKITAEKDVAEPKTKEDIKLRGAQKAYHKAKE  420
ACM79874.1       DSEEKTVQKTQNKTNQDAGTEKVQSAENKKITAEKDVAEPKTKEDIKLRGAQKAYHKAKE  174

t38568aep        AHKKDAKKFENTLS--EQPKAEAKEENGQVAEKAENHDTATVLGNPRPSSFSESKHNNKT  475
Spin|ORF15       AHKKDAKKFENTLSVNEQPKAEAKEENGQLAEKAENHDTATVLGNPRPSSFSESKHNNKT  394
AAC39121.1       ------------------------------------------------------------    -
XP_012553808.1   AHKKDAKKFENTLSVNEQPKAEAKEENGQLAEKAENHDTATVLGNPRPSSFSESKHNNKT  480
ACM79874.1       AHKKDAKKFENTLSVNEQPKAEAKEENGQLAEKAENHDTATVLGNPRPSSFSESKHNNKT  234

t38568aep        AVGDPEAKKYLSTANAAIDVAAEAVVDAETLESGHKENAVDKGGMKYLKAASDAVAAAHE  535
Spin|ORF15       TVGDPEAKKYLSTANAAIDVAAEAVVDAETLESGHKENAVDKGGMKYLKAASDAVAAAHE  454
AAC39121.1       ------------------------------------------------------------    -
XP_012553808.1   TVGDPEAKKYLSTANAAIDVAAEAVVDAETLESGHKENAVDKGGMKYLKAASDAVAAAHE  540
ACM79874.1       TVGDPEAKKYLSTANAAIDVAAEAVVDAETLESGHKENAVDKGGMKYLKAASDAVAAAHE  294
```

```
t38568aep        ALSRAAELGGFGSLPAPNFKPLEKGEKDTTLAKKYTQAASEAVNAAEKALDNLAAIGYPD  595
Spin|ORF15       ALSRAAELGGFGSLPAPNFKPLEKGEKDTTLAKKYTQAASEAVNAAEKALDNLAAIGYPD  514
AAC39121.1       ------------------------------------------------------------    -
XP_012553808.1   ALSRAAELGGFGSLPAPNFKPLEKGEKDTTLAKKYTQAASEAVNAAEKALDNLAAIGYPD  600
ACM79874.1       ALSRAAELGGFGSLPAPNFKPLEKGEKDTTLAKKYTQAASEAVNAAEKALDNLAAIGYPD  354


t38568aep        VIPPVNEAKSQPTNENNAVKKHDVNFILNKEDGSNEDAESGSGVSETNTENAQVKSIIEA  655
Spin|ORF15       VIPPVNEAKSQPTNENNAVSD---------------------------------------  535
AAC39121.1       ------------------------------------------------------------    -
XP_012553808.1   VIPPVNEAKSQPTNENNAVKKHDVNFILNKEES-NEDAES--GSGETNTENAQVKSIIEA  657
ACM79874.1       VIPPVNEAKSQPTNENNAVKKHDVNFILNKEES-NEDAES--GSGETNTENAQVKSIIEA  411


t38568aep        PSKRDQQGEFIEVANKAVDEAHEALMNAVRKLPNKRSV  693
Spin|ORF15       --------------------------------------    -
AAC39121.1       --------------------------------------    -
XP_012553808.1   PSKRDRQGEFIEVANKAVDEAHEALMNAVRKLPNKRSV  695
ACM79874.1       PSKRDRQGEFIEVANKAVDEAHEALMNAVRKLPN----  445
```

XP_012553808.1 is the most complete sequence. AAC39121.1 and ACM79874.1 are both part of this sequence-indicating that the spinalin sequence published by Koch et al 1998 (Koch et al., 1998)only covers the 254 aminao acids of a 695 amino acids long protein. The nematoblast-specific protein nb054-sv9, published by Milde et al 2008 (Milde et al., 2009) as a taxonomically restricted gene overlaps with this sequence by 8 amino acids only and covers almost the whole C-terminal region. SpinORF15 is 535 amino acids long, the first 20 amino acids (italic) are not part of the protein. It covers the middle part and overlaps with both, AAC39121.1 and ACM79874.1.

---

#### 3.) Sox-family of transcription factors-comparison of Hydra and Hydractinia sequences

**t5528aep|SX21B_DANRE**
>XP_002154370.1 PREDICTED: transcription factor Sox-19a-like [Hydra vulgaris]
TRINITY_DN14709_c0_g1 -ORF6 (**HySox19a|ORF6**)
**t23837aep**
>XP_012563508.1 PREDICTED: transcription factor Sox-21-B-like [Hydra vulgaris]
TRINITY_DN5602_c0_g1 -ORF6 (**HySox21b|ORF6**)
**t23172aep|SOX14_MOUSE**
>XP_012555836.1 PREDICTED: uncharacterized protein LOC101236863 [Hydra vulgaris]
TRINITY_DN5359_c0_g1 (**HySox14|ORF7**)

Hydra Sox2, Sox4, Sox8:
Reference: Punctuated emergences of genetic and phenotypic innovations in eumetazoan, bilaterian, euteleostome, and hominidae ancestors. (Wenger et al., 2016)

Hydractinia echinata Sox22, Sox23, Sox24, Sox25, SoxB1, SoxB2, SoxB3
Reference: An Evolutionarily Conserved SoxB-Hdac2 Crosstalk Regulates Neurogenesis in a Cnidarian. (Flici et al., 2017)

Alignment of Sox genes from Hydra Vulgaris and Hydractinia echinata

```
HySox14|ORF7    ------------------------------------------------------------     -
HYDEC--SoxB3    ------------------------------------------------------------     -
Hydra--Sox8     ---------------------------------------------MMAASETNSNFATHV    15
Hydra--Sox4     -----------------------------------------------MELDFGLDY----     9
HYDEC--Sox25    ------------------------------------------------------------     -
HySox21b|ORF6   MMRDSSSPYMPSN---DVPALTKMINDSG-EDDIDLIKSEPRDVQNVLHTDANSNYHTLQ    56
HYDEC--Sox23    MLRTTASPYLPSEAQEDVPALAKLVNLSSSEETSQSIKTEAHDVPSLVQVDANSNYPPSQ    60
Hydra--Sox2     MA-------------------TA---------------------INVIQHPSDANNDTHF    20
HYDEC--SoxB1    MT-------------------TTA-------EVISQTATAT---INVNHIKPDNNYPTHH    31
HySox19a|ORF6   -------------------------------------------MNAMVTSQAKPEFMT--    15
HYDEC--SoxB2    -------------------------------------------MNNMVAQVK-QDVLTHI    16
HYDEC--Sox22    MYEKS-----------NIPKVEPV-------EMVSNPMTSLNPHSQILAQQQQQHNGSGS    42
HYDEC--Sox24    -----------------------------------------------MSQQEQANNEQHI    13
```

```
HySox14|ORF7     ------------------------------------------------------------    -
HYDEC--SoxB3     ---------------------------------------------------------MEN    3
Hydra--Sox8      IQNNNITHAVSQEQRKGEGYGSTTISLSEAAQCLKNLGEEGILCDDALLRLTMSSTSEHG   75
Hydra--Sox4      ---------------------------FMSGALPTPEPQDGDIKHLNP--LELNIDFQLG   40
HYDEC--Sox25     --------------------------MEALHDVQLTEDK-ET-----------LTMESNR   22
HySox21b|ORF6    SNHL--------------------NTNLQNHHSPIQNNH-------N------AALSSQS   83
HYDEC--Sox23     PPPP-----------------SHQTSHQSSHHSPQPRSQ-------Q------S--GQSL   88
Hydra--Sox2      KQVVGY--------------NTSQ-----SMLVPQN----LNLSNRV------SNSLVSD   51
HYDEC--SoxB1     --ILPM--------------TMSATSLSPTMQTSNI----SHLMSQE------AQSYAHD   65
HySox19a|ORF6    PHTTQ----------------------LGLHQSQSPAPT-----------------VAPT   36
HYDEC--SoxB2     PQLTQ-------------------T--AQLQQVPQQPPRSSP-----------SV-PATT   43
HYDEC--Sox22     PSPTLM--------------SMPQQTNLSSGQT---------------------TLNPSQ   67
HYDEC--Sox24     --INAQ--------------NITANTNVNNGQVQSNGPFDATVASVD------NVEAKTD   51
```

HMG boxes A and B DNA-binding domains

```
HySox14|ORF7     MASDLDKPKGHIKRPMNSFMVWSRMERKRISEANPKMHNSEISKQLGTSWKMLSEEDRAP   60
HYDEC--SoxB3     NELEIEVKKGHIKRPMNSFMVWSRLERKKISEANPKMHNSEISKRLGASWKLLTEEERKP   63
Hydra--Sox8      ETEFLDTKNSRVKRPMNSFMVWAQTARKKLAEKYPHLHNAHLSKMLGKLWKMLSPDEKQP  135
Hydra--Sox4      CGNKKKSSMDHVKRPMNAFMVWSQIERKKMADIYPDMHNAEISRRLGKRWKLLSDADRRP  100
HYDEC--Sox25     KPKEERTHNTKVKRPMNAFMLWSKQKRREISRKDPSLHNAQISKLLGEEWKVLDVDAKLP   82
HySox21b|ORF6    PHASVEEDDDHVKRPMNAFMVWSRTERRKLALKYPNMLNCEISKLLGAEWSRMGEEEKSP  143
HYDEC--Sox23     GQGQPEEDDDHVKRPMNAFMVWSRTERRKLALKYPNMLNCEISKLLGAEWSRMNEEEKSP  148
Hydra--Sox2      HETSKQEDFDKVKRPMNSFMVWSREKRRRLAHENPKMHNSEISKRLGAEWKVLTEDEKAP  111
HYDEC--SoxB1     SDPSKLDDPERVKRPMNSFMVWSREKRRKLAQENPKMHNSEISKRLGAEWKVLTDEEKAP  125
HySox19a|ORF6    SPNPLNGDMSHVKRPMNAFMVWSRGKRRQMAQDNPRMHNSEISKRLGAEWKCLTQQEKQP   96
HYDEC--SoxB2     TLHGVNMDPNHVKRPMNAFMVWSRGKRRQMAQENPRMHNSEISKRLGAEWKCLTQVEKQP  103
HYDEC--Sox22     TNGDVAKKPDHVKRPMNAFMVWSREKRRKMAQINPRMHNSEISKILGAEWKRMGEIEKQP  127
HYDEC--Sox24     ESQKKTDEPQKIKRPMNPFMIFGCEKRRKLAQVHPRMHNSEISKILGAEWKRMSEYEKAP  111
                            6KRPMN FM65s    R4 6a   P 6hN  6S4 LG  Wk 6   e4 P
```

```
HySox14|ORF7     YAEEAKRLRDLHMSEYPDYKYRPKRKPKASSVNQEKL--PSIA-PKRP------------  105
HYDEC--SoxB3     YAEEAKRLRELHMEEHPEYKYRPKRKPKVGLISQEKI--PLPSQANRP------------  109
Hydra--Sox8      YVLEASRLDKLHKDEHPEYKYRPRRRPKGLKRGYS-----TPTMIVSPTTTSKPYTIPSN  190
Hydra--Sox4      FVIRSEKLREEHMRRYPDYKYRPKKKAKELAAKELAAK-ELKNFSKV-----------CE  148
HYDEC--Sox25     YVEESQKLMMKHKKEHPDYRYKPRQNKLSKNMKTF-STYTPPVLI-----KG-------N  129
HySox21b|ORF6    YIQESKRLRTIHSQKYPDYSYKPRRRKRKVTQNQ--PTYPSINF----------------  185
HYDEC--Sox23     YIQESKRLRTIHSQKYPDYSYKPRRRKRKVTQSQ--TVYPPINF----------------  190
Hydra--Sox2      FVFEAKRLRAEHMKSHPDYKYRPRRKAKTSSKKNE-QKMP---HVITT--DGKQIFMPTQ  165
HYDEC--SoxB1     FVYEAKRLRAEHMKSHPDYKYRPRRKTKSLSKKTE-NKIAMPTAVIGA--DGKQIFMSAQ  182
HySox19a|ORF6    FIDEAKRLRAVHIQEHPDYKYKPKRRKQKTTKKDIYTPYPSIG-----------------  139
HYDEC--SoxB2     FIDEAKRLRAVHIQEHPDYKYKPKRRKPKQLKKDLYPSYPNMN-----------------  146
HYDEC--Sox22     YVEEAKRLQTQHSIEYPNYKYKPRRRKPKTMIKKDKMGFPYAT-------DP--------  172
HYDEC--Sox24     YVQEAKRLKEQHSIEYPNYKFKANRRKPRQQVKKERPTFPYAA-------DL--------  156
                 5  e k4L   H    P Y 54p

HySox14|ORF7     --------SS----IPLADYTQSRS------VP-------------TAVPVYTGFHRRTP  134
HYDEC--SoxB3     --------TA----MSIPEYSQQRS------G---------------AVSMYASYHHIPA  136
Hydra--Sox8      WARIVHAPENENLISPRAEA-----------AQIVYATDGM---QYYAVQKGSGF-----  231
Hydra--Sox4      YNI-VTLTNGRNDDWDKSKYTNNVV---------------C--CENN---KAPPN---KT  184
HYDEC--Sox25     MPYAPGHFPRY-----RPYDQHYLS------RQDVCPVPGCYECIMYERSYERKYHYCKP  178
HySox21b|ORF6    -PSAGNAMQ--NYQFPSSAYQKFS--------SDAFKMGGFGPDTGYQLPMHDPN--KAA  232
HYDEC--Sox23     -PSGATALP--PYQFSGSGYGKMP--------SDAFKMPGFA-DNGYQVPLQDPN--KA-  235
Hydra--Sox2      YTSGYAITGGINY--PFNGYLSALV----------------NGHEAYASNTSPSA-----  202
HYDEC--SoxB1     YPY--AIASGINYPVPFNGYMSALV----------------NGHEAYPAG---AT-----  216
HySox19a|ORF6    ----QGMVPNIDSKYASIGYQPTLSYGMS---SDMYNK--LNGGYGYQTTISTGY----P  186
HYDEC--SoxB2     ----AGMMPGIDPKYGSMAYQQSMGYGISSISPDMYSK--MSPGYGYPAAISPGY---HP  197
HYDEC--Sox22     ----NGIPTGMKFPYPSPFPQDSMY-------APIYQM--PG--------APPGY-----  206
HYDEC--Sox24     ----NAIPTAMKFPYPSPFFQDSMY-------GSMYPA--MAGS------SHPAY-----  192

HySox14|ORF7     -----EYR-----------PHVGPENVTTVVYPHSRY-YRTDSPEKVSHTHSPLDHYHGR  177
HYDEC--SoxB3     HQYVGHPR-----------VVNLPEGVTTIHYPSSRYHYRSHSPVEKRRSRSPLDRDFRR  185
Hydra--Sox8      --------TPTMAHHHGNSVIVSPTNQ-PI-----TLVQSTSAP------TSNYPAYVTV  271
Hydra--Sox4      FVTISRQVGQIGSL---SNITVSPQKC--IDIPPSTLSPPSVVPD---CCGTPPDEYNSE  236
HYDEC--Sox25     FTY-----APSMAS---NCQCNRPR---------VNTPEEKRSPK---AFDVE-----SL  213
```

```
HySox21b|ORF6   LSY---LNSATLGL---SNDMK--------SYYNLT---SPAAN----MFSPATSQYSGL  271
HYDEC--Sox23    LNY---LN--AMAL---PADMK--------SYYNMT---AAGSN----MFSPATSNYSSL  272
Hydra--Sox2     -VY-GFPHGATIAV---TSTISQSQ------LPSSS---SSSVT----AATGLSAQYSVV  244
HYDEC--SoxB1    -MY-GIPPGTAIAV---ASLPQSTT----------S---SSATS----VTPVTAAQYSVV  254
HySox19a|ORF6   LMYSNYSVGPSMVG---SHSQSSPTGA-HQSYPSSTITSQIGTPV---ITDSTYR-----  234
HYDEC--SoxB2    VMYSNYSLSSM---------TPSPTAG-GRGYTTVTMTSPNGTPV---MTDSNANTY---  241
HYDEC--Sox22    AMYADYG-SPAIAR---QAMVP------------AQVRHSPGAT----TTASS-------  239
HYDEC--Sox24    SLYSGGDYSSALAR---QAMVT------------GSVVRGAQPPS---TTGSASAEYYPL  234
                  y

HySox14|ORF7    --SPPIRIYHRSPPSPYD-SREHYSRSRENQRESLG-IR----PRSRSPVTRYYSYDEDI  229
HYDEC--SoxB3    --HSPELIYHPISPQEHHSFRGDYIKSTREYRRSVSPVAPHNKPQSNSPKREK-------  236
Hydra--Sox8     ----------PMP-----------ATSTAIFHPIAIPVQGVHTPAF--VLRSSYPG-SNI  307
Hydra--Sox4     S----FYNYDSFSGK---------NNSNSLYHEDMDLLINFAR---------FTDG----  270
HYDEC--Sox25    IKKTNCHVTKVENE----------VKRKT-------------------------------  232
HySox21b|ORF6   SLSGNN-QTNL------------YSNGSNYLPPLFKDHQ---------------------  297
HYDEC--Sox23    SLGGPSSQNSL------------YSSQSNYMPPLFKDHQ---------------------  299
Hydra--Sox2     HSGTPTYMYSPL-GF-------PYLNGAA-----MGTYGGQP------------------  273
HYDEC--SoxB1    P-GGTTYMYSPFSSF-------PYLNGAT-----MSAYSAQA------------------  283
HySox19a|ORF6   -----------VSTS-------DYINSKNYFSNMNSPYSPVDS-----------AA--TS  263
HYDEC--SoxB2    -----------RPAT-------DYMNSKSYYSNVTSQYSPVPA-----------AS-APQ  271
HYDEC--Sox22    -------VHDFYSSM-------AMQ------------TARITS-----------------  256
HYDEC--Sox24    NKASNAPINDYYSGV-------VSNKSSDYYSAMHSAAAAGTRT------ERTAPG-GEV  280

HySox14|ORF7    A---DPKSDYEDGDAK-YHQKLSAEKKRKGVDDLLGEKMRAKARAEQIKHWENDKGYNGY  285
HYDEC--SoxB3    ---------SPTEEDT-TERKNSQRNKRKGVDDLLGEKMKEQAREEQSRRSGDENFNSSF  286
Hydra--Sox8     LSPSAIPIAYHNIDHRT-----------------------IESHMQPNQHIKTESTNTIV  344
Hydra--Sox4     -----LPDLYTT------------------------------------------------  277
HYDEC--Sox25    ------------------------------------------------------------    -
HySox21b|ORF6   -----------------F------------------------------------------  298
HYDEC--Sox23    -----------------F------------------------------------------  300
Hydra--Sox2     ---------YSHALTL------------K-----PDTSSITVSTEKATSHLSM-----HH  302
HYDEC--SoxB1    ----AAAAAYPHTVTV------------K-----HEAAVTTTATEKPAEAHTVETAKATS  322
```

```
HySox19a|ORF6   ---THSQNRYPTTDEN----RNIVNQAHS-----VGNGMVSK--------LIQEHDV---   300
HYDEC--SoxB2    ---VHSQARYPSPDEN----RHSTQVS-------SNDIMLTK--------ANHESTT---   306
HYDEC--Sox22    ------AEMMPMT-----NDRYSSVVTQS-----NMPSFTTT----PTLHI---PSD---   290
HYDEC--Sox24    ---VNLAERYPAADNRTFENRYSTSFNGS-----QGSTIATYSTMQATSHLSTETPA---   329


HySox14|ORF7    D---EYTKYSPEFLSKSYVVPIPVMVHKRGSPHHMVHTN---KNEVCREECCIVPAVRYL   339
HYDEC--SoxB3    VRCNERRSPIAYRSSAPYVVPVPVVLQPRSSH------M---SPDVCYKECCMVPAVQYI   337
Hydra--Sox8     KTPDGET--------------IVIIQRPIDG--HVVPFQLREVPSGATNTITVVPQSQVI   388
Hydra--Sox4     ----------------------PEVSEMLTG--QWLEND---L--------GF-------   295
HYDEC--Sox25    ------------------------------------------------------------     -
HySox21b|ORF6   ------------------------------------------------------------     -
HYDEC--Sox23    ------------------------------------------------------------     -
Hydra--Sox2     DAVRSTI--------------APSVYYPIG---IYPTAV---DH--NC-NLTYDPKIPYA   339
HYDEC--SoxB1    TTTTATA--------------LPSHYYPIGG--MYPTAV---AVDQNSNITAYDPKTQYA   363
HySox19a|ORF6   ----QSN--------------FPDNAVN--R--NWSLNF---SSV---------------   320
HYDEC--SoxB2    ----PPT--------------YSSENTT--R--HWPQTT---QQSDLSRPVAYVP-----   336
HYDEC--Sox22    ----SIA--------------YSQMYAQ--R--H--------------------------   302
HYDEC--Sox24    ----AYS--------------YSSLYDQ--R--H--------------------------   341


HySox14|ORF7    PHSYHHHHNYHDIRAY--------------ENHPKHCFCSECKRISPK----------SK   375
HYDEC--SoxB3    PSGMEYHKNYIPRHNMRTH--------SSHNSTSKRCRCSDCEEGAHRQFYEYKSYPRSP   389
Hydra--Sox8     QQHIVQTPNNQE----------------------------------------------TK   402
Hydra--Sox4     ------------------------------------------------------------     -
HYDEC--Sox25    ------------------------------------------------------------     -
HySox21b|ORF6   ------------------------------------------------------------     -
HYDEC--Sox23    ------------------------------------------------------------     -
Hydra--Sox2     TMTLLEQPHGNVKREQTRE-GTTRNISDDHQ---------SRRVN-------------TP   376
HYDEC--SoxB1    SMALIGQAKRVEMQGTNPHPAVVITMPEHHSPSPARSMNTSVQQN-------------SP   410
HySox19a|ORF6   ------------------------------------------------------------     -
HYDEC--SoxB2    -VLL--------------------------------------------------------   339
HYDEC--Sox22    ------------------------------------------------------------     -
HYDEC--Sox24    ------------------------------------------------------------     -
```

```
HySox14|ORF7    RYSTVQSYSVDECTPHQNMIEKKLSMTEREISKNETSDDSDSSVKINSNVKIEGKALEQS  435
HYDEC--SoxB3    KYEHEVKEEVAECTNECDDSTS---------SRPPRSK--DRSLSP--------------  424
Hydra--Sox8     I------L----------------------------------------------------  404
Hydra--Sox4     ------------------------------------------------------------    -
HYDEC--Sox25    ------------------------------------------------------------    -
HySox21b|ORF6   ------------------------------------------------------------    -
HYDEC--Sox23    ------------------------------------------------------------    -
Hydra--Sox2     T------YSISSNTATQ-------------------------------------------  387
HYDEC--SoxB1    V------LAVSE------------------------------------------------  416
HySox19a|ORF6   ------------------------------------------------------------    -
HYDEC--SoxB2    ------------------------------------------------------------    -
HYDEC--Sox22    ------------------------------------------------------------    -
HYDEC--Sox24    ------------------------------------------------------------    -


HySox14|ORF7    NNQIKAM  442
HYDEC--SoxB3    -------    -
Hydra--Sox8     -------    -
Hydra--Sox4     -------    -
HYDEC--Sox25    -------    -
HySox21b|ORF6   -------    -
HYDEC--Sox23    -------    -
Hydra--Sox2     -------    -
HYDEC--SoxB1    -------    -
HySox19a|ORF6   -------    -
HYDEC--SoxB2    -------    -
HYDEC--Sox22    -------    -
HYDEC--Sox24    -------    -
```

All proteins in the alignment contain a single SOX-TCF_HMG-box and are class I members of the HMG-box superfamily of DNA-binding proteins. Other members of the family include SRY and its homologs in insects and vertebrates, and transcription factor-like proteins, TCF-1, -3, -4, and LEF-1.

Phylogenetic tree for Hydra Vulgaris and Hydractinia echinata Sox genes:

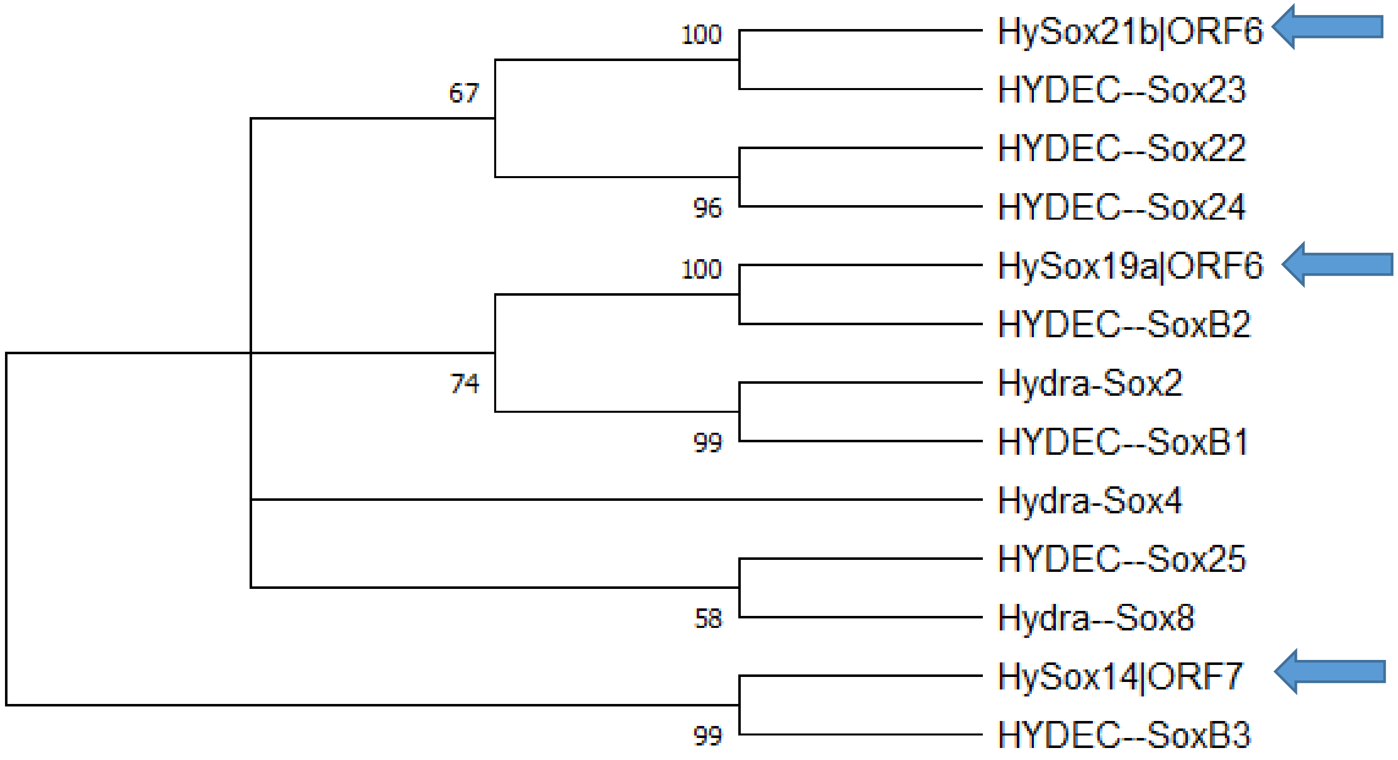

Phylogenetic tree for Hydra Vulgaris, Hydractinia echinata Sox genes and all Human Sox genes:

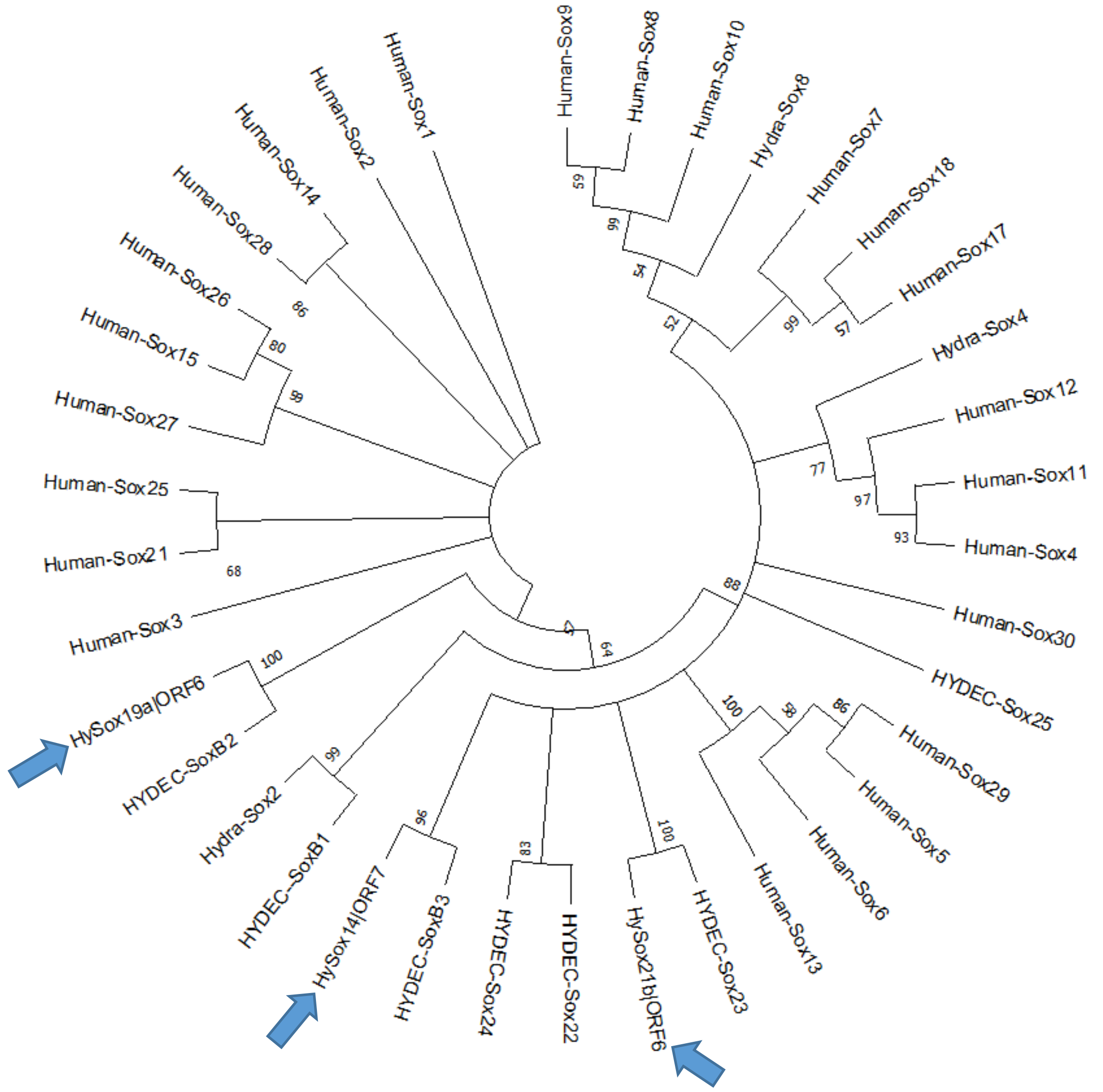

**4.) Otx**

**t33622aep|**OTX1B_DANRE
TRINITY_DN19967_c0_g1 (**HyOtx1|ORF4**)

**QCF59210.1** homeobox transcription factor Otx1 [Hydra vulgaris], (**HyOTX1Reddy**)
Reference: Molecular signature of an ancient organizer regulated by Wnt/β-catenin signalling during primary body axis patterning in *Hydra*. (Reddy et al., 2019)

Alignment of TRINITY_DN19967_c0_g1-ORF4 (**HyOtx1|ORF4**) and QCF59210.1 (**HyOTX1Reddy**) and OTX proteins from other animals:

```
CaeelTtx-1    MSLTSSSAPSDSIVELIQSQNLSGSGSGTGGNTGSTTMNSGNFIPTPSLTGTGASQSSGS   60
HyOtx1|ORF4   ------------------------------------------------------------   -
HyOTX1Reddy   ------------------------------------------------------------   -
DromeOC-E     ------------------------------------------------------------   -
DanreOTX-1    ------------------------------------------------------------   -
HumanOTX-1    ------------------------------------------------------------   -
MouseOTX-1    ------------------------------------------------------------   -


CaeelTtx-1    ASSGNFPMSYIPNVSSATTVAAANMSAYFNQKSAYPTSH-LGFPS--NVGSHSFLQSNMY  117
HyOtx1|ORF4   -----------------------------------------------MSWSYS-PTPPFP   12
HyOTX1Reddy   -----------------------------------------------MSWSYS-PTPPFP   12
DromeOC-E     -----------------------MAAGFLKSGDLGPHPHSYGGPHPHHSVPHGPLPPGMP   37
DanreOTX-1    ------------------------MMSYL-------------------------KQPPYA   11
HumanOTX-1    ------------------------MMSYL-------------------------KQPPYG   11
MouseOTX-1    ------------------------MMSYL-------------------------KQPPYG   11
                                                                    p
```

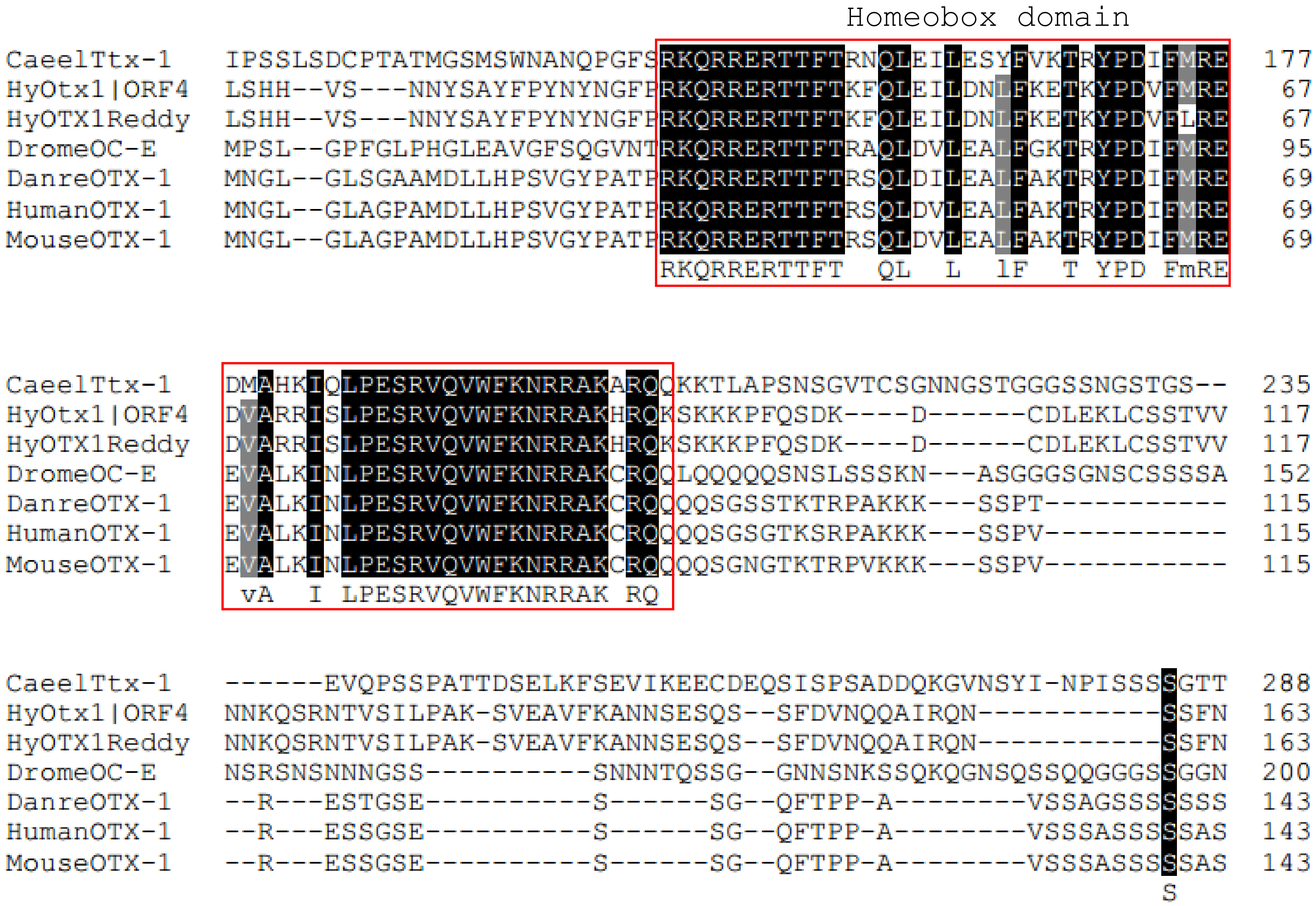
Homeobox domain
CaeelTtx-1 IPSSLSDCPTATMGSMSWNANQPGFSRKQRRERTTFTRNQLEILESYFVKTRYPDIFMRE 177
HyOtx1|ORF4 LSHH--VS---NNYSAYFPYNYNGFPRKQRRERTTFTKFQLEILDNLFKETKYPDVFMRE 67
HyOTX1Reddy LSHH--VS---NNYSAYFPYNYNGFPRKQRRERTTFTKFQLEILDNLFKETKYPDVFLRE 67
DromeOC-E MPSL--GPFGLPHGLEAVGFSQGVNTRKQRRERTTFTRAQLDVLEALFGKTRYPDIFMRE 95
DanreOTX-1 MNGL--GLSGAAMDLLHPSVGYPATPRKQRRERTTFTRSQLDILEALFAKTRYPDIFMRE 69
HumanOTX-1 MNGL--GLAGPAMDLLHPSVGYPATPRKQRRERTTFTRSQLDVLEALFAKTRYPDIFMRE 69
MouseOTX-1 MNGL--GLAGPAMDLLHPSVGYPATPRKQRRERTTFTRSQLDVLEALFAKTRYPDIFMRE 69
RKQRRERTTFT QL L lF T YPD FmRE
CaeelTtx-1 DMAHKIQLPESRVQVWFKNRRAKARQQKKTLAPSNSGVTCSGNNGSTGGGSSNGSTGS-- 235
HyOtx1|ORF4 DVARRISLPESRVQVWFKNRRAKHRQKSKKKPFQSDK----D------CDLEKLCSSTVV 117
HyOTX1Reddy DVARRISLPESRVQVWFKNRRAKHRQKSKKKPFQSDK----D------CDLEKLCSSTVV 117
DromeOC-E EVALKINLPESRVQVWFKNRRAKCRQQLQQQQSNSLSSSKN---ASGGGSGNSCSSSSA 152
DanreOTX-1 EVALKINLPESRVQVWFKNRRAKCRQQQQSGSSTKTRPAKKK---SSPT----------- 115
HumanOTX-1 EVALKINLPESRVQVWFKNRRAKCRQQQQSGSGTKSRPAKKK---SSPV----------- 115
MouseOTX-1 EVALKINLPESRVQVWFKNRRAKCRQQQQSGNGTKTRPVKKK---SSPV----------- 115
vA I LPESRVQVWFKNRRAK RQ
CaeelTtx-1 ------EVQPSSPATTDSELKFSEVIKEECDEQSISPSADDQKGVNSYI-NPISSSSGTT 288
HyOtx1|ORF4 NNKQSRNTVSILPAK-SVEAVFKANNSESQS--SFDVNQQAIRQN-----------SSFN 163
HyOTX1Reddy NNKQSRNTVSILPAK-SVEAVFKANNSESQS--SFDVNQQAIRQN-----------SSFN 163
DromeOC-E NSRSNSNNNGSS----------SNNNTQSSG--GNNSNKSSQKQGNSQSSQQGGGSSGGN 200
DanreOTX-1 --R---ESTGSE----------S------SG--QFTPP-A--------VSSAGSSSSSSS 143
HumanOTX-1 --R---ESSGSE----------S------SG--QFTPP-A--------VSSSASSSSSAS 143
MouseOTX-1 --R---ESSGSE----------S------SG--QFTPP-A--------VSSSASSSSSAS 143
S

```
CaeelTtx-1    SYNGTSSFRPQAQYPYASYPAY---D-FQYSQSNTNSTNTYTLDGNSTWKFQMS------  338
HyOtx1|ORF4   SSNQQISSAALEK--SQTFPCYLPTSLSSYRP-------FLIEAHSSTYSSAQARSFYSN  214
HyOTX1Reddy   SSNQQISSAALEK--SQTFPCYLPTSLSSYRP-------FLIEAHSSTYSSAQARSFYSN  214
DromeOC-E     NSNNNSAAAAASA--AAAVAAA--------QS---------IKTHHSSFLSAAAAAASGG  241
DanreOTX-1    STNNT-------------GICS--------TS-------TSISTVSSIW-S---------  165
HumanOTX-1    SSSANPAAAAAAGLGGNPVAAA--------SS-------LSTPAASSIW-S---------  178
MouseOTX-1    SASANPAAAAAAGLGGNPVAAA--------SS-------LSTPTASSIW-S---------  178
              s                                             S   s

CaeelTtx-1    ------------------------------------------------------------    -
HyOtx1|ORF4   YAQAPVEYQSCVVNFNNSQAVST-------------------------------------  237
HyOTX1Reddy   YAQAPVEYQSCVVNFNNSQAVST-------------------------------------  237
DromeOC-E     TNQSAN-NNSNNNNQGNSTPNSSSSGGGGGSQAGGHLSAAAAAAALNVTAAHQNSSPLLP  300
DanreOTX-1    -------------------PA-I-------------------------SP--GSAPPSVS  178
HumanOTX-1    -------------------PASI-------------------------SP--GSAPASVS  192
MouseOTX-1    -------------------PASI-------------------------SP--GSAPTSVS  192

CaeelTtx-1    ------------------------------------------------------------    -
HyOtx1|ORF4   -----------------QWTNYGSSI----------------------------------  246
HyOTX1Reddy   -----------------QWTNYGSSI----------------------------------  246
DromeOC-E     TPATSVSPVSIVCKKEHLSGGYGSSVGGGGGGGGASSGGLNLGVGVGVGVGVGVGVSQDL  360
DanreOTX-1    LPEPV-APSNTSCMQRSVSG---TAS-----------STYPMPYNQTTGYSQGYPTP---  220
HumanOTX-1    VPEPLAAPSNTSCMQRSVAAGAATAA-----------ASYPMSYGQGGSYGQGYPTP---  238
MouseOTX-1    VPEPLAAPSNASCMQRSVAAGAATAA-----------ASYPMSYGQGGSYGQGYPAP---  238

CaeelTtx-1    ------------------------------------------------------------    -
HyOtx1|ORF4   ------------------------------------------------------------    -
HyOTX1Reddy   ------------------------------------------------------------    -
DromeOC-E     LRSPYDQLKDAGGDIGAGVHHHHSIYGSAAGSNPRLLQPGGNITPMDSSSSITTPSPPIT  420
DanreOTX-1    -SGSYFSGVDCGSYLA-PMHSHHH---------PH------QLSPMTASSMPTHPHHHIS  263
HumanOTX-1    -SSSYFGGVDCSSYLA-PMHSHHH---------PH------QLSPMAPSSMAGHHHHHPH  281
MouseOTX-1    -SSSYFGGVDCSSYLA-PMHSHHH---------PH------QLSPMAPSSMAGHHHHHPH  281
```

```
CaeelTtx-1    ------------------------------------------------------------   -
HyOtx1|ORF4   ------------------------------------------------------------   -
HyOTX1Reddy   ------------------------------------------------------------   -
DromeOC-E     PMSPQSAAAAAHAAQSAQSAHHSAAHSAAYMSNHDS-----------YNFWHNQYQQY-P   468
DanreOTX-1    QSS-----------GHHHHHHQAYSGTGLAFNSSDCLDYKEQT----ASSWKLNFNTTDC   309
HumanOTX-1    AHHPLSQSSGHHHHH-HHHHHQGYGGSGLAFNSADCLDYKEPGAAAASSAWKLNFNSPDC   340
MouseOTX-1    AHHPLSQSSGHHHHHHHHHHHQGYGGSGLAFNSADCLDYKEPAAAAASSAWKLNFNSPDC   341


CaeelTtx-1    ------------------------------------------------------------   -
HyOtx1|ORF4   ------------------------------------------------------------   -
HyOTX1Reddy   ------------------------------------------------------------   -
DromeOC-E     NNYAQAPSYYSQMEYFSNQNQVNYNMGHSGYTASNFGLSPSPSFTGTVSAQAFSQNSLDY   528
DanreOTX-1    LDYKDQASWRFQVL----------------------------------------------   323
HumanOTX-1    LDYKDQASWRFQV-----------------------------------------------   353
MouseOTX-1    LDYKDQASWRFQVL----------------------------------------------   355


CaeelTtx-1    -----------   -
HyOtx1|ORF4   -----------   -
HyOTX1Reddy   -----------   -
DromeOC-E     MSPQDKYANMV   539
DanreOTX-1    -----------   -
HumanOTX-1    -----------   -
MouseOTX-1    -----------   -
```

HyOtx1|ORF4 is identical with Reddy’s QCF59210.1.

Phylogenetic tree of OTX Proteins:

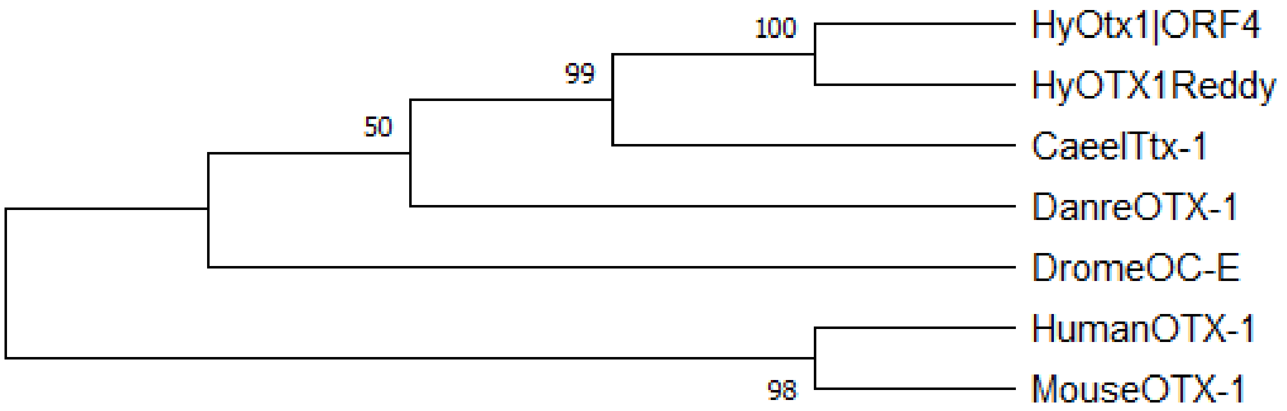


---

**5.) Alx**

**t16456aep|RX_RAT**
TRINITY_DN5649_c0_g1 (**HyALX|ORF4**)

**AAG03082.1** aristaless-like protein [Hydra vulgaris]
Reference: HyAlx, an aristaless-related gene, is involved in tentacle formation in Hydra;(Smith et al., 2000)

**XP_012557002.1** PREDICTED: homeobox protein cut-like isoform X1 [Hydra vulgaris]

Alignment of TRINITY_DN5649_c0_g1-ORF4 (**HyAlx|ORF4**), AAG03082.1 (**HyAlx-Smith**), XP_012557002.1 (**HyCut-like**) and Alx proteins from other animals:

```
HyCut-like      -----------MKTLALCTILLFALVVV--QSKPLSS-----ELPHSREFYAG-------   35
Drome-Hbn       ------------------------------------------------------------    -
HyALX|ORF4      ------------------------------------------------------------    -
HyAlx-Smith     ------------------------------------------------------------    -
STRPU-Alx       MKRKFEAPQIAMDSAVCCTPRTMDADNRWNASVPSPERHREAGMV-V--------RESNA   51
MouseAlx3       -----------MDPERCAPFSVGPAAGPY-------------------------------   18
Human-ALX       ----------------------------------------------------------ME    2
DanreAlx4       -----------MNAETCVSYCEMT--SMDSYYSPSAPQGRDHQANPFRTFQASDTKYSPA   47
MouseAlx4       -----------MNAETCVSYCESPAAAMDAYYSPVSQSR--EGSSPFRGFPGGD-KFGTT   46


HyCut-like      -------------YIKAITDNFDQLFESDRQTRSTKDEDESLKT-----------S----   67
Drome-Hbn       ------------------------MMTTTTS------------------------QHHQH   12
HyALX|ORF4      ------------------------------------------------------------    -
HyAlx-Smith     ------------------------------------------------------------    -
STRPU-Alx       FGAVAPATRGLAGMTGSVGKCNDMMMNSGS----WVDVSRMTDA-------RVKDNEQVF  100
MouseAlx3       -AAAGDEAPGP----------------QGTP-----DAAPHLHPA--PPRGPRLSRFPAC   54
Human-ALX       FLSEKFALKSPP------SKNSDFYMGAGGP---LEHVMETLDNESFYSKASAGKCVQAF   53
DanreAlx4       FLT--NKGQGYG------EKSGSP-FQQE---------CQSLD------ATAGEGTFN--   81
MouseAlx4       FLSAGAKGQGFG------DAKSRARYGAG-----QQDLAAPLES-----SSGARGSFNKF   90


HyCut-like      --------------HEVNEKPVHKVDKRAEISNQEDNNINKRDKENFKNTGESLKKRHIQ  113
Drome-Hbn       HPIM----PPAMRPAPVQESPVSRPRAVYSIDQILGNQHQ-------------------I   49
HyALX|ORF4      ----------------MIHKP--MAKMQFSIDMILGNSKS-------------------E   23
HyAlx-Smith     ----------------MIHKP--MAKMQFSIDMILGNSKS-------------------E   23
STRPU-Alx       TSLI--------------------------KTQVLDSS------------VEK------K  116
MouseAlx3       GPLEPYLPEPAKPP----AKYLQDLG----PGPVLNGGHF------YEGSAEA------E   94
Human-ALX       GPLPR----------AEHHVRLERT-----------------------------------   68
DanreAlx4       ----------------KYHLFMQRSS----CKTPPDSSKL------------Q------Q  103
MouseAlx4       QPQPPTPQPPPAPPAPPAHLYLQRGA----CKTPPDGSLK------------L------Q  128
```

```
HyCut-like      DKVDEDSDIQKRDTVPESNGENTANNLIAEKRNDIKNLTEQ----------TD-------  156
Drome-Hbn       KRSD--TPSEVLITHPHHGHPHHIHHLHSSNSNGSNHLSHQ-------QQQQHSQ-QQHH   99
HyALX|ORF4      EK----------------------------------------------------------   25
HyAlx-Smith     EK----------------------------------------------------------   25
STRPU-Alx       DKTER-SIH---IWRPALDGDE--------KETSKKDVDVEVSRAARIWRPNDPQ-PEAH  163
MouseAlx3       EKASK-AASFPQLPVDCRGGPR--------DG--PSNV----------------Q---AS  124
Human-ALX       --------------SPCQDSS---------VNYGITKV--E-------GQPLHTELNRAM   96
DanreAlx4       EN----SGHNGGL-IACYGKDS--------TGLTDSEL------------PQNSD-PAGM  137
MouseAlx4       EGS---GGHNAALQVPCYAKE---------SNLGEPEL------------PPDSE-PVGM  163

HyCut-like      -N------------DNIKTEFENSNQYDVRNDVDTDNNKIDADNNKVGLETNYISDKTTD  203
Drome-Hbn       SQQQQQQQQLQVQAKREDSPTNTDGGLDVDNDDELSSSLNN--GHDLS----DMERPRKV  153
HyALX|ORF4      -----QQNDVKL--Q---REFNQEKGINVSSTIRFNSPISSEIKNDEI----EGENRQKL   71
HyAlx-Smith     -----QQNDVKL--Q---REFNQEKGINVSSTIRFNSPISSEIKNDEI----EGENRQKL   71
STRPU-Alx       ST---TAT----TMTMGDQERNESSRYVDDFDSDGDDEHLDEHGSNAG----DRPTKRKQ  212
MouseAlx3       PG---PCLASL------------------------SVPLSPGLPDSME----LAKTKSKK  153
Human-ALX       DN----CNSLRMSPVKGMQE----------------KGELDELGDKCD----SNVSSSKK  132
DanreAlx4       DG---SYLSVKDSGVKSPQQ---------------ATSELASPLDKTE----GESNKGKK  175
MouseAlx4       DN---SYLSVKETGAKGPQD--------------RASAEIPSPLEKTD----SESNKGKK  202
                                                                          k

                                      Homeobox domain
HyCut-like      QQDDNSVKKYFVDDVDHASVNDEKTTNQESDYASGADPEDSNINEKKIPDLDFKENLNSQ  263
Drome-Hbn       RRSRTTFTTFQLHQ------------------------LERAFEKTQYPDVFTREDLAMR  189
HyALX|ORF4      RRNRTTFTTYQLHQ------------------------LERSFDKTQYPDVFTRENLALK  107
HyAlx-Smith     RRNRTTFTTYQLHQ------------------------LERSFDKTQYPDVFTRENLALK  107
STRPU-Alx       RRYRTTFTSYQLEE------------------------LERAFCKTHYPDVFTREELAMR  248
MouseAlx3       RRNRTTFSTFQLEE------------------------LEKVFQKTHYPDVYAREQLALR  189
Human-ALX       RRHRTTFTSLQLEE------------------------LEKVFQKTHYPDVYVREQLALR  168
DanreAlx4       RRNRTTFTSYQLEE------------------------LEKVFQKTHYPDVYAREQLALR  211
MouseAlx4       RRNRTTFTSYQLEE------------------------LEKVFQKTHYPDVYAREQLAMR  238
                rr rt3f   q6                          le  f kt yPD6  4E La
```

```
HyCut-like     NDKKEKSIRKKVEKKLPGKKLKNQKSKQIFQDWEEDENNAHLKSDGPITEQNLEDDEYSS  323
Drome-Hbn      LDLSEAR--------------------------------------------------VQV  199
HyALX|ORF4     LDLSEAR--------------------------------------------------VQV  117
HyAlx-Smith    LDLSEAR--------------------------------------------------VQV  117
STRPU-Alx      VDLTEAR--------------------------------------------------VQV  258
MouseAlx3      TDLTEAR--------------------------------------------------VQV  199
Human-ALX      TELTEAR--------------------------------------------------VQV  178
DanreAlx4      TDLTEAR--------------------------------------------------VQV  221
MouseAlx4      TDLTEAR--------------------------------------------------VQV  248
                dl Ear                                                   vqv

HyCut-like     SFSERKHLIDEPDNIMFQQRSFSPTLFGYRILRSPINPVGAEMAFTRFNSFSPINSFTPY  383
Drome-Hbn      WFQNRRAKWRKREKFMNQDKAGYLLP-----------EQ-GIPEFPLGIPLPP--HGLPG  245
HyALX|ORF4     WFQNRRAKWRKREK-LSFGNSPSNSVCSSAIEYHQDSQI-SIPEVPILEVTPIVKRSFIA  175
HyAlx-Smith    WFQNRRAKWRKREKVMQFGNSPSNSVCSSAIEYHQDSQI-SIPEVPILEVTPIVKRSFIA  176
STRPU-Alx      WFQNRRAKWRKREKLGLQPRLHPHPHFGP--------IL-G-------------------  290
MouseAlx3      WFQNRRAKWRKRERYGKMQEGRNP--FTT--------AY-DISVLPRTDSHP--------  240
Human-ALX      WFQNRRAKWRKRERYGQIQQAKSH--FAA--------TY-DISVLPRTDSYP--------  219
DanreAlx4      WFQNRRAKWRKRERFGQMQQVRTH--FST--------AY-EIPLLTRPENYA--------  262
MouseAlx4      WFQNRRAKWRKRERFGQMQQVRTH--FST--------AY-EIPLLTRAENYA--------  289
               wFqnR4akwrkre

HyCut-like     SAIAMQNYLFPTNSRDGMLIPVTRGIAREIITKRDGNNNNNNCCCQCCCPCCHHTCSHQD  443
Drome-Hbn      HPGSMQSEFWPPHFALHQHF-------------------NPAAAAAAGLL----------  276
HyALX|ORF4     NKQSCS----CSHITDNKH---------------------------LGYD----------  194
HyAlx-Smith    NKQSCS----CSHITDNKH---------------------------LGYD----------  195
STRPU-Alx      -----------GPISDSNLV-------------------SSHLCRLAC------------  308
MouseAlx3      ---QLQNSLWPSPGSGSPGG---------------------PCLMSPEGI----------  266
Human-ALX      ---QIQNNLWAGNASGGSVV---------------------TSCMLPRDT----------  245
DanreAlx4      ---QIQNPSWIGGSSAASPV---------------------PGCVVPCDS----------  288
MouseAlx4      ---QIQNPSWIGNNGAASPV---------------------PACVVPCDP----------  315
```

```
HyCut-like     CHDCCHCHHDVHH-H-D----CCHHHHVKHCHPTIHHTYHEHH--HDHCHPCHHHESHCH  495
Drome-Hbn      ---PQHLMAPHYKLPNFHT-LLSQYMGL---------SNLNGIFGAGAAAAAAAASAGYP  323
HyALX|ORF4     ---NSSLIKPTL--PHTCTARSCLYYGG---------KRHKNRFSDLRCS----HEENFN  236
HyAlx-Smith    ---NSSLIKPTL--PHTCTARSCLYYGG---------KRHKNRFSDLRCS----HEENFN  237
STRPU-Alx      -------IDR-AHRQ--------GLLTSPPPPPFV-------------------------  327
MouseAlx3      ---PSPCMSPYSHSHGN----VAGFMGVPASPA-----AHPGIYSIHGFPPALGG-HSFE  313
Human-ALX      ----SSCMTPYSHSPRTDSS----YTGFSNHQNQFSHVPLNNFFTDSLLTGATNG-HAFE  296
DanreAlx4      ---VTSCMPPHPHA----ASGVSDFLGVPSPGSHMGQTHMGSLFGSPGMGTGING---YD  338
MouseAlx4      ---VPACMSPHAHPPGSGASSVSDFLSVSGAGSHVGQTHMGSLFGAAGISPGLNG---YE  369
                        p

HyCut-like     -----HPCECKHVHHHEPCCHDCNHECHHDCDH---HDCCPFDCGHHEYGHLDDGCHNCL  547
Drome-Hbn      QNLSLHAGLSAMSQVSPPCSNSSPRESPKLVPHPTPPHATPPAGGNGGGLLTGGLISTA  383
HyALX|ORF4     ------KRTFHKIIYQEPCNRRYTDL----SDHTDH-------------SL-RYSYFH--  270
HyAlx-Smith    ------KRTFHKIIYQEPCNRRYTDL----SDHTDH-------------SL-RYSYFH--  271
STRPU-Alx      ------------------------------------------------------------    -
MouseAlx3      PSPDGDYKSPSLVS-----LRMKPKEPPGLLNWTT-------------------------  343
Human-ALX      TKPEFERRSSSIAV-----LRMKAKEHTANISWAM-------------------------  326
DanreAlx4      LNMDPDRKSSSIAA-----LRMKAKEHSAAISWAT-------------------------  368
MouseAlx4      MNGEPDRKTSSIAA-----LRMKAKEHSAAISWAT-------------------------  399

HyCut-like     NQHDYHGAYNCAPEHYCGGYQPHYQHSCGCNHGYQPAIHIIGNVDGTYGRHGGYLRRKR  607
Drome-Hbn      ---------AQSPNSA------------AGASSNASTPVSVVTKGED-------------  409
HyALX|ORF4     ------------------------------------------------------------    -
HyAlx-Smith    ------------------------------------------------------------    -
STRPU-Alx      ------------------------------------------------------------    -
MouseAlx3      ------------------------------------------------------------    -
Human-ALX      ------------------------------------------------------------    -
DanreAlx4      ------------------------------------------------------------    -
MouseAlx4      ------------------------------------------------------------    -
```

```
HyCut-like    GLKK  611
Drome-Hbn     ----    -
HyALX|ORF4    ----    -
HyAlx-Smith   ----    -
STRPU-Alx     ----    -
MouseAlx3     ----    -
Human-ALX     ----    -
DanreAlx4     ----    -
MouseAlx4     ----    -
```

HyALX|ORF4 is identical with HyALX-Smith.

Phylogenetic tree of ALX proteins:

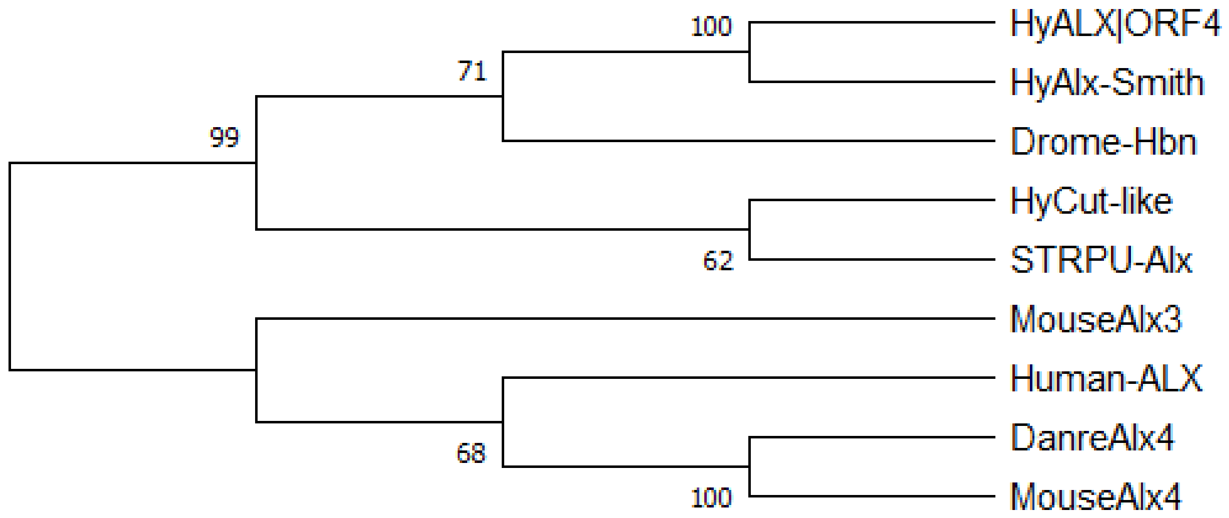


---

**6.)   JUN**

**t17964aep|JUN_AVIS1**
TRINITY_DN3178_c1_g1 (**HyJun|ORF8**)

Alignment of Jun proteins from Hydra and other animals:

```
HyJun|ORF8   MLSGTLPLNEKFLSMEYVLDTTFYSDDFLTRTLTT-SVL--Y-------------DKSQL   44
Caeel-Jun    ~~~~~~~~~~~~~~~~~~~~~~~~~~~~~~~~~~~~~~~~~~~~~~~~~~~~~~~~~~~~    -
Drome-Jun    ~~~~~~~~~~~~~~~~~~MKTPVSA-A--ANLSIQNAGSSGATAIQIIPKTEPVGEEGPM   39
Danre-Jun    ~~~~~~~~~~~~~~MSTKMETTFYDDSLNSAFSQHDSATFGYNH-KA--------LKHNM   37
Mouse-Jun    ~~~~~~~~~~~~~~MTAKMETTFYDDALNASFLQSESGAYGYSNPKI--------LKQSM   38
Human-Jun    ~~~~~~~~~~~~~~MTAKMETTFYDDALNASFLPSESGPYGYSNPKI--------LKQSM   38
                                 t


HyJun|ORF8   DLTF-------------------VENNNCLPSKRPQYESTVESPNLDFLKLGSPDLEQMF   85
Caeel-Jun    ~~~~~~~~~~~~~~~~~~~~~~~~~~~~~~~~~~~~~~~~~~~~~~~~~~~~~~~~~~~~    -
Drome-Jun    SLDFQSPNLNTSTPNPNKRPGSLDLNSKSAKNKRIFAPLVINSPDLSSKTVNTPDLEKIL   99
Danre-Jun    T-------LNLSDPA-----GNLK------PHLRAKASDILTSPDVGLLKLASPELERLI   79
Mouse-Jun    T-------LNLADPV-----GSLK------PHLRAKNSDLLTSPDVGLLKLASPELERLI   80
Human-Jun    T-------LNLADPV-----GSLK------PHLRAKNSDLLTSPDVGLLKLASPELERLI   80
                                              r        sp         p le


HyJun|ORF8   MNINENDSNFSSEDK--SDSSAFSLDDDFVSTENCFTDALQQLHDKQEILPTDFSGTNCN  143
Caeel-Jun    ~~~~~~~~~~~~~~~~~~~~~~~~~~~~MLNW---------GHHHNSYDEPSASSSSGSS   23
Drome-Jun    LSNN---LMQTPQPGKVFPTKAGPVTVEQLDFGRGFEEALHNLHTNSQAFPSANSAANSA  156
Danre-Jun    IQSSNGMITTTPTPTQFL--CPKNVTDEQEGFAEGFVRALAELHHQH--MPNVTSAPQTT  135
Mouse-Jun    IQSSNGHITTTPTPTQFL--CPKNVTDEQEGFAEGFVRALAELHSQNT-LPSVTSAAQPV  137
Human-Jun    IQSSNGHITTTPTPTQFL--CPKNVTDEQEGFAEGFVRALAELHSQNT-LPSVTSAAQPV  137
                                                f  al  lH      P   S
```

```
HyJun|ORF8   NRVGKENLHSKILQGKRKPISSPMNIQKLRTQNPMKQLYNPNQQRRNNFQHPCSHPPNKN   203
Caeel-Jun    SSSVA-------------------------------------------------------    28
Drome-Jun    A-----------------------------------------------------------   157
Danre-Jun    I-----------------------------------------------------------   136
Mouse-Jun    S-----------------------------------------------------------   138
Human-Jun    N-----------------------------------------------------------   138

HyJun|ORF8   QHRLNNLAVHHQNNISHLSISEQMALNNQLTMMNNMNQSINAHHMQQQHEIAFLTQAMGF   263
Caeel-Jun    ----AN-----------LSVSYN---------SDSRNQGCMGGG--------QYSGNIGG    56
Drome-Jun    ----NN-----------T-TAAA---------MTAVNNGISGG-----------------   175
Danre-Jun    --------------------NSS---------MAPV-SSIAGGA--------VYSSAMRA   158
Mouse-Jun    ----GA-----------GMVAPA---------VASV-AGAGGGG--------GYSASLHS   165
Human-Jun    ----GA-----------GMVAPA---------VASV-AGGSGSG--------GFSASLHS   165
                                                      g

HyJun|ORF8   QSDREPSLNYQNDRTMINSNGFQSQHNVPSQVPSQLQQKLLEHQLMQQQIGASYQNNNNN   323
Caeel-Jun    GGGG--------------------YGD---------------------------------    63
Drome-Jun    ---------------------TFTYTNMTE------------------------------   184
Danre-Jun    DPPVYADLNTFNPAISSSSAN-----PAAMSFPS--------------------------   187
Mouse-Jun    EPPVYANLSNFNPGALSSGGGAPSYGAAGLAFPSQPQQQ----------------QQPPQ   209
Human-Jun    EPPVYANLSNFNPGALSSGGGAPSYGAAGLAFPAQPQQQ----------------QQ---   206

                                   Basic-leucine zipper(bZIP) domain
HyJun|ORF8   TTHHVPQIDSING-----LGI-DEGTLKMLMENPHMVPVNLELQELIKRERKKLRNRVAS   377
Caeel-Jun    -----------------------------YSHIDPINMMALDDQEKKKLERKRARNRQAA    94
Drome-Jun    --------------GFSVIKDEP----VNQASSPTVNPIDMEAQEKIKLERKRQRNRVAA   226
Danre-Jun    ----APPQLPVQHPRLQALKEEPQTVPEMPGETPPLSPIDMESQERIKAERKRMRNRIAA   243
Mouse-Jun    PPHHLPQQIPVQHPRLQALKEEPQTVPEMPGETPPLSPIDMESQERIKAERKRMRNRIAA   269
Human-Jun    PPHHLPQQMPVQHPRLQALKEEPQTVPEMPGETPPLSPIDMESQERIKAERKRMRNRIAA   266
                                              p 6 p6 6e QE iK ERK4 RNR Aa
```

```
HyJun|ORF8   SKCRKRKLEREGRLEDKVKDLKEKNIELNAVANALKQQICDLKQRVMDHVGDGCQIVLPP  437
Caeel-Jun    TKCRQKKMDRIKELEEQVLHEKHRGQRLDAELLELNRALEHFRRTVEHHSGNGCPNNSIR  154
Drome-Jun    SKCRKRKLERISKLEDRVKVLKGENVDLASIVKNLKDHVAQLKQQVMEHIAAGCTVPPNS  286
Danre-Jun    SKCRKRKLERISRLEDKVKTLKSQNSELASTANMLREQVAQLKQKVMNHVNSGCQLMLTQ  303
Mouse-Jun    SKCRKRKLERIARLEEKVKTLKAQNSELASTANMLREQVAQLKQKVMNHVNSGCQLMLTQ  329
Human-Jun    SKCRKRKLERIARLEEKVKTLKAQNSELASTANMLREQVAQLKQKVMNHVNSGCQLMLTQ  326
             3KCRk4K6eRi  LE  Vk lK  n  L       L   6  l4q Vm H   GC

HyJun|ORF8   ~~~~~    -
Caeel-Jun    V~~~~  155
Drome-Jun    TDQ~~  289
Danre-Jun    QLQTF  308
Mouse-Jun    QLQTF  334
Human-Jun    QLQTF  331
```

TRINITY_DN3178_c1_g1-ORF8 contains a complete JUN-bZIP domain.

---

### 7.) Forkhead box proteins

**t9145aep|FOXI1_XENTR;**
TRINITY_DN18625_c0_g1 (**HyFoxI1c|ORF6**)
**t12948aep|FOXN4_DANRE;**
TRINITY_DN1167_c0_g1 (**HyFoxN1|ORF7**)
**t19720aep|FOXP1_XENLA;**
TRINITY_DN5643_c0_g1 (**HyFoxP1|ORF14**)

CDG72033.1 Hydra vulgaris Forkhead box protein N1 [Hydra vulgaris] (**HyFOXN1-Wenger**)
Reference: Punctuated emergences of genetic and phenotypic innovations in eumetazoan, bilaterian, euteleostome, and hominidae ancestors.(Wenger and Galliot, 2013)

Alignment of **HyFoxI1c|ORF6**, **HyFoxN1|ORF7**, **HyFoxP1|ORF14**, **HyFOXN1-Wenger** and other ortholog proteins:

```
HyFOXP1|ORF14     ------------------------------------------------------------   -
Drome--FOXP-d     ------------------------------------------------------------   -
Danre-FOXP1-B     MMQESGTEAA-NGTAHQNGAP-----PSVEGHREVRSKSTTPSSDITASDIINFQ--QHQ   52
Mouse--FOXP-1     MMQESGSETKSNGSAIQNGSSGGNHLLECGALRDTRSNGEAPAVDLGAADLAHVQQQQQQ   60
Human-FOXP1-a     MMQESGTETKSNGSAIQNGSGGSNHLLECGGLREGRSNGETPAVDIGAADLAHAQQQQQQ   60
Drome-Jumeau      ------------------------------------------------------------   -
Human--FOXN1      ------------------------------------------------------------   -
Mouse--FOXN1      ------------------------------------------------------------   -
Danre--FOXN4      ------------------------------------------------------------   -
HyFOXN1|ORF7      ------------------------------------------------------------   -
HyFOXN1-Wenger    ------------------------------------------------------------   -
HyFOXI1c|ORF6     ------------------------------------------------------------   -
Drome--fd102C     ------------------------------------------------------------   -
Caeel--fkh-10     ------------------------------------------------------------   -
Danre--FoxI1      ------------------------------------------------------------   -
Human-FoxI1-a     ------------------------------------------------------------   -
Mouse--FoxI1      ------------------------------------------------------------   -


HyFOXP1|ORF14     ---------------------------------------------MNEHSDDSGVDNDKS   15
Drome--FOXP-d     ------------------------------------------------------------   -
Danre-FOXP1-B     ALQVARQILLQQQ--------------------------------QQSSVHKSPKNNDKQ   80
Mouse--FOXP-1     ALQVARQLLLQQQQQQQQQQQQQQQQQQQQQQQQQQQQQQQQQQQQQQVSGLKSPKRNDKQ  120
Human-FOXP1-a     ALQVARQLLLQQQ------------------------------QQQQVSGLKSPKRNDKQ   90
Drome-Jumeau      ------------------------------------------------------------   -
Human--FOXN1      ------------------------------------------------------------   -
```

```
Mouse--FOXN1        ------------------------------------------------------------     -
Danre--FOXN4        ------------------------------------------------------------     -
HyFOXN1|ORF7        ------------------------------------------------------------     -
HyFOXN1-Wenger      ------------------------------------------------------------     -
HyFOXI1c|ORF6       ------------------------------------------------------------     -
Drome--fd102C       ------------------------------------------------------------     -
Caeel--fkh-10       ------------------------------------------------------------     -
Danre--FoxI1        ------------------------------------------------------------     -
Human-FoxI1-a       ------------------------------------------------------------     -
Mouse--FoxI1        ------------------------------------------------------------     -


HyFOXP1|ORF14       PLIEQLPQQNQKQNISVAD-SFKAVTPHDDAYICSDTVHSYHGKQTNYYKDEPKSFL-LL    73
Drome--FOXP-d       ------------------------------------------------------------     -
Danre-FOXP1-B       PATQ-VP-------VSVAMMTPQVITPQQMQQILQHQVL------------SPQQLQLLL   120
Mouse--FOXP-1       PALQ-VP-------VSVAMMTPQVITPQQMQQILQQQVL------------SPQQLQVLL   160
Human-FOXP1-a       PALQ-VP-------VSVAMMTPQVITPQQMQQILQQQVL------------SPQQLQVLL   130
Drome-Jumeau        ------------------------------------------------------------     -
Human--FOXN1        ------------------------------------------------------------     -
Mouse--FOXN1        ------------------------------------------------------------     -
Danre--FOXN4        ------------------------------------------------------------     -
HyFOXN1|ORF7        ------------------------------------------------------------     -
HyFOXN1-Wenger      ------------------------------------------------------------     -
HyFOXI1c|ORF6       ------------------------------------------------------------     -
Drome--fd102C       ------------------------------------------------------------     -
Caeel--fkh-10       ------------------------------------------------------------     -
Danre--FoxI1        ------------------------------------------------------------     -
Human-FoxI1-a       ------------------------------------------------------------     -
Mouse--FoxI1        ------------------------------------------------------------     -


HyFOXP1|ORF14       NNQPKGMVNCTRETRNFEKSGMDEHTVIHKLLRNGRCVWP-------SCDQAFNSRTDFI   126
Drome--FOXP-d       ------------------------------MHRIHDDEYSED-----AKESDFKSSIQKE    25
Danre-FOXP1-B       QQQQALMLQ--QQLQEFYKKQQEQL-HLQLIQQQHGSKQQS--KEVSAQQLAFQQQLLQV   175
Mouse--FOXP-1       QQQQALMLQ--QQLQEFYKKQQEQL-QLQLLQQQHAGKQPKEQ-QVATQQLAFQQQLLQM   216
```

```
Human-FOXP1-a    QQQQALMLQQ-QQLQEFYKKQQEQL-QLQLLQQQHAGKQPKEQQQVATQQLAFQQQLLQM  188
Drome-Jumeau     ------MF----ELEDY-SSGI------------HEGFFSKY-------ADAAGPSLDFY   30
Human--FOXN1     ------------------------------------------------------------    -
Mouse--FOXN1     ------------------------------------------------------------    -
Danre--FOXN4     ------------------------------------------------------------    -
HyFOXN1|ORF7     ------------------------------------------------------------    -
HyFOXN1-Wenger   ------------------------------------------------------------    -
HyFOXI1c|ORF6    ------------------------------------------------------------    -
Drome--fd102C    ------------------------------------------------------------    -
Caeel--fkh-10    ------------------------------------------------------------    -
Danre--FoxI1     ------------------------------------------------------------    -
Human-FoxI1-a    ------------------------------------------------------------    -
Mouse--FoxI1     ------------------------------------------------------------    -


HyFOXP1|ORF14    RHLDSYHVLDEK--G-----------AAQARVQGYVVRELEEKLSYEKSKLTAMLT-HLQ  172
Drome--FOXP-d    ISVKSRHQISIPDIC--SDAVKNNCFPPT----GFLNNSITFASHVVKCSSPASSIDESS   79
Danre-FOXP1-B    QQLQQQHLLSLQRQGLLSIQPNQ-TLPLHTLTQGMIPAELQQL---WKEVTNSHVKEENS  231
Mouse--FOXP-1    QQLQQQHLLSLQRQGLLTIQPGQPALPLQPLAQGMIPTELQQL---WKEVTSAHTAEETT  273
Human-FOXP1-a    QQLQQQHLLSLQRQGLLTIQPGQPALPLQPLAQGMIPTELQQL---WKEVTSAHTAEETT  245
Drome-Jumeau     VSDSMQEMLNVDIRAEIANVVG----SSSSDLTSSLDQTLEAISAINNNQSNGNSSQSAS   86
Human--FOXN1     -------MVSLP--------------PPQSDV------TLPGPTRLE-----G-------   21
Mouse--FOXN1     -------MVSLL--------------PPQSDV------TLPGSTRLE-----G-------   21
Danre--FOXN4     -------------------------------M------TV--QSKLH-----G-------    9
HyFOXN1|ORF7     ------------------------------------------------------------    -
HyFOXN1-Wenger   ------------------------------------------------------------    -
HyFOXI1c|ORF6    ------------------------------------------------------------    -
Drome--fd102C    ------------------------------------------------------------    -
Caeel--fkh-10    ------------------------------------------------------------    -
Danre--FoxI1     ------------------------------------------------------------    -
Human-FoxI1-a    ------------------------------------------------------------    -
Mouse--FoxI1     ------------------------------------------------------------    -
```

```
HyFOXP1|ORF14    FTN---EKGSILLQKTAHNLSPPPGRHSVQVTSHPSTS-----NGSGQCVMVPYPPPG-S  223
Drome--FOXP-d    TAAQQHESNPHMHIQGQHMMAPVPDLGFYNVPEFISEQEKLMFSDAERF--LRSKDNE-V  136
Danre-FOXP1-B    VTNNGH--------RGLDLSSPSP------VPLKNHNQHG---STNGQYI-SHSLK----  269
Mouse--FOXP-1    SSN--H--------SSLDLTSTCVSSSAPSKSSLIMNPHA---STNGQLS-VHTPK----  315
Human-FOXP1-a    GNN--H--------SSLDLTTTCVSSSAPSKTSLIMNPHA---STNGQLS-VHTPK----  287
Drome-Jumeau     Y----NANANFLTSSGLHA-SPTAKWMGSS---ANFWSNSDYYADLGACVNPISVMPLIN  138
Human--FOXN1     -----ERQGDLMQAPGLPG-SPAPQSK--H---AG-FSCSSFVSDGPPERTPSLP-PHSP   68
Mouse--FOXN1     -----EPQGDLMQAPGLPD-SPAPQNK--H---AN-FSCSSFVPDGPPERTPSLP-PHSP   68
Danre--FOXN4     -----R---------------------------GK-FKK-RFFRAGQQVPRPTLE-LSSV   34
HyFOXN1|ORF7     -----------------------------------------MFNQYQTSS--LSD-LDSI   16
HyFOXN1-Wenger   -----------------------------------------MFNQYQTSS--LSD-LDSI   16
HyFOXI1c|ORF6    ------------------------------------------------------------    -
Drome--fd102C    ------------------------------------------------------------    -
Caeel--fkh-10    ------------------------------------------------------------    -
Danre--FoxI1     ------------------------------------------------------------    -
Human-FoxI1-a    ------------------------------------------------------------    -
Mouse--FoxI1     ------------------------------------------------------------    -


HyFOXP1|ORF14    RSHIGSFDHSTDNEHTVIHKLLRNGRCV----W-----------------PSCDQAFNSR  262
Drome--FOXP-d    CNNDFSYMHDEFAMRKYYHPLFAHGICR----W-----------------PGCEMDLEDI  175
Danre-FOXP1-B    --REGST---LDDHSPHSHPLYGHGVCK----W-----------------PGCEAVFEDF  303
Mouse--FOXP-1    --RESL----SHEEHPHSHPLYGHGVCK----W-----------------PGCEAVCDDF  348
Human-FOXP1-a    --RESL----SHEEHPHSHPLYGHGVCK----W-----------------PGCEAVCEDF  320
Drome-Jumeau     STSAGMFSPKKNKTASSTQGRSGAVPSSPS---------------AERDQHKSHLTFSP-  182
Human--FOXN1     RI--ASPGPEQVQGHCPAGPGPGPFRLSPSDKYPGFGFEEAAASSPGRFLKGSHAPFHP-  125
Mouse--FOXN1     SI--ASPDPEQIQGHCTAGPGPGSFRLSPSEKYPGFGFEEGPAGSPGRFLKGNHMPFHP-  125
Danre--FOXN4     WLSKIFYNPEQHHNKQKMIESGITTRMSGIHENPGQSHHT--------------------   74
HyFOXN1|ORF7     SVHDIDFD------------LGSTNNCYGQFTY-GD-YLQ--------------------   42
HyFOXN1-Wenger   SVHDIDFD------------LGSTNNCYGQFTY-GD-YLQ--------------------   42
HyFOXI1c|ORF6    ------------------------------------------------------------    -
Drome--fd102C    -------------MAHS-------DICALQTTTADQD-----------------------   17
Caeel--fkh-10    ------------------------------------------------------------    -
Danre--FoxI1     -----MFLEGERIMNAFGQQPSS----QQTSPLQQQDILDMT--------VYCDSNFSM-   42
Human-FoxI1-a    -------------MSSFDLPAPSPPRCSPQFPSIGQEPPEMN--------LYYENFFH--   37
Mouse--FoxI1     -------------MSSFDLPAPSPPRCSPQFPSIGQEPPEMN--------LYYENFFH--   37
```

```
HyFOXP1|ORF14   TDFIRHLDSYHVLDEKG---------------------------------AAQARVQGYV  289
Drome--FOXP-d   TSFVKHLNTEHGLDDRS---------------------------------TAQARVQMQV  202
Danre-FOXP1-B   QSFLKHLNNEHALDDRS---------------------------------TAQCRVQMQV  330
Mouse--FOXP-1   PAFLKHLNSEHALDDRS---------------------------------TAQCRVQMQV  375
Human-FOXP1-a   QSFLKHLNSEHALDDRS---------------------------------TAQCRVQMQV  347
Drome-Jumeau    --AQMKVSAGSMRRDQVMAHIPKQISVVTGTGTTAPATMATNSVLQRRNSSAVDAVRKDL  240
Human--FOXN1    --YKRPFHEDVFPEAE--------------------TTLAL-------------------  144
Mouse--FOXN1    --YKRHFHEDIFSEAQ--------------------TAMAL-------------------  144
Danre--FOXN4    ------------------------------------------------------------    -
HyFOXN1|ORF7    ------------------------------------------------------------    -
HyFOXN1-Wenger  ------------------------------------------------------------    -
HyFOXI1c|ORF6   ------------------------------------------------------------    -
Drome--fd102C   -------QGTS-------------------------------------------------   21
Caeel--fkh-10   ------------------------------------------------------------    -
Danre--FoxI1    --YQQNLHHHH-------------------------------------------------   51
Human-FoxI1-a   ------------------------------------------------------------    -
Mouse--FoxI1    ------------------------------------------------------------    -


HyFOXP1|ORF14   VRELEEKLS-------------------------YEKSKLTAMLTHLQFTNEKGSILLQK  324
Drome--FOXP-d   VSQLESHLQ-------------------------KERDRLQAMMHHLYLSKQLLSPT---  234
Danre-FOXP1-B   VQQLELQLA-------------------------KDKERLQAMMTHLHVKSTEPKPT---  362
Mouse--FOXP-1   VQQLELQLA-------------------------KDKERLQAMMTHLHVKSTEPKAA---  407
Human-FOXP1-a   VQQLELQLA-------------------------KDKERLQAMMTHLHVKSTEPKAA---  379
Drome-Jumeau    VTELRKAQSSPVPNSLEELGKGKGSTLLNASVGATNTIKLAPGIGGLTFANSAAYQKLKQ  300
Human--FOXN1    -----KGHSFKTPGPLEAFEEIP------VDVA-EAEAFLP-GFSAEAWCNGLPYPSQEH  191
Mouse--FOXN1    -----DGHSFKTQGALEAFEEIP------VDMG-DAEAFLP-SFPAEAWCNKLPYPSQEH  191
Danre--FOXN4    --------------SAQDYRLLT------TDPS-QLKDELPGDLQSLSWLTSVDVPRLQQ  113
HyFOXN1|ORF7    --------------ELQELEQGR------TPMK-MDRHSLPRNLTNLHWLHNVGMQADQQ   81
HyFOXN1-Wenger  --------------ELQELEQGR------TPMK-MDRHSLPRNLTNLHWLHNVGMQADQQ   81
HyFOXI1c|ORF6   ------------------------------------------------------------    -
Drome--fd102C   ----------------------------------------------------------SR   23
Caeel--fkh-10   ------------------------------------------------------------    -
Danre--FoxI1    ----------------------------------------------------------HH   53
Human-FoxI1-a   ------------------------------------------------------------    -
Mouse--FoxI1    ------------------------------------------------------------    -
```

```
HyFOXP1|ORF14     TAHNLSPPPGRHSVQ-VTSHPSTSNGSGQCVMVPYPPPGSRSHIGSFDHST-------DN  376
Drome--FOXP-d     -KIDRKDVPGREGK--FCRSPLTVNSIGRPIRQTN-----------------------SP  268
Danre-FOXP1-B     -PQPLNLVSNVA----LSK---TAPAASPPLSLPQ-----------------------TP  391
Mouse--FOXP-1     -PQPLNLVSSVT----LSK---SASEAS-PQSLPH-----------------------TP  435
Human-FOXP1-a     -PQPLNLVSSVT----LSK---SASEAS-PQSLPH-----------------------TP  407
Drome-Jumeau      TS--LVKSPG----------------------GISPGAGSNMGLKREDSNKRGLQASTTP  336
Human--FOXN1      ---------------------------------------GPQVLGSE--------VKVKP  204
Mouse--FOXN1      ---------------------------------------NQILQGSE--------VKVKP  204
Danre--FOXN4      IG--GGRPDFTSSAQSSLLERQTAQLN--SMTVA-GGAGSAIHLQSE--------MQHSP  160
HyFOXN1|ORF7      QQ--IRQEPI-----SLMVDPNT------NMPVQFSANRSAIH-----------------  111
HyFOXN1-Wenger    QQ--IRQEPI-----SLMVDPNT------NMPVQFSANRSAIH-----------------  111
HyFOXI1c|ORF6     ------------------------------------------------------------    -
Drome--fd102C     DS--NVPKPSLPNIYPLG-TTRTSQ-----------AQSTTMTLEQY-----RLQLYNYA   64
Caeel--fkh-10     ------------------------------------------------------------    -
Danre--FoxI1      HH--HQRPPAHPSS----------------------------------------------   65
Human-FoxI1-a     PQ--GVPSPQRP-S----------------------------------------------   48
Mouse--FoxI1      PQ--GMPSPQRPTS----------------------------------------------   49


HyFOXP1|ORF14     EHLTSPLH-PYTRDGFSSPPPSTSPRRFPSHNIGHNNFIPHHKSQGPYHPPPQRYYPALH  435
Drome--FOXP-d     SPLNLPM----------------------------------------------VNST---  279
Danre-FOXP1-B     TTPTAPLT-PLSQT--------------------------------HSV----ITPTSLH  414
Mouse--FOXP-1     TTPTAPLT-PVTQG--------------------------------PSV----ITTTSMH  458
Human-FOXP1-a     TTPTAPLT-PVTQG--------------------------------PSV----ITTTSMH  430
Drome-Jumeau      KSIA--S----------AANS--------------------------PHHQMQSNYS---  355
Human--FOXN1      PVLE--S----GAGMFCYQPP------------------------------LQHMYC---  225
Mouse--FOXN1      QALD--S----GPGMYCYQPP------------------------------LQHMYC---  225
Danre--FOXN4      LAIN--SMPQFSPGFPCAASV--------------------------YQTAPQQVLT---  189
HyFOXN1|ORF7      ------AMQSQNITFMSLAQT----------------------------------HA---  128
HyFOXN1-Wenger    ------AMQSQNITFMSLAQT----------------------------------HA---  128
HyFOXI1c|ORF6     ------------------------------------------------------------    -
Drome--fd102C     LNIERLRCPQYGGTGGSGA-S-------------------------------APWLH---   89
Caeel--fkh-10     --MEEIL---------------------------------------------RQFL----    9
Danre--FoxI1      ----------YGLGEYSSPST-------------------------------NPYLW---   81
Human-FoxI1-a     ----------FEGGGEYGATP-------------------------------NPYLW---   64
Mouse--FoxI1      ----------FEGGGEYGTTP-------------------------------NPYLW---   65
```

```
HyFOXP1|ORF14    QAYPIPPKRESPNEIDRRSPKKEHITSPGLIGMNHERHSSEIHRSMLVEGHTTGSSHMQQ  495
Drome--FOXP-d    NL----------CSIKKRNHDKNTFSINGGL-----------------------------  300
Danre-FOXP1-B    SV----------GPIRRRYSDKYNMP----------------------------------  430
Mouse--FOXP-1    TV----------GPIRRRYSDKYNVP----------------------------------  474
Human-FOXP1-a    TV----------GPIRRRYSDKYNVP----------------------------------  446
Drome-Jumeau     ----LGS----PSSL-----SSSSASSPLGNVSNLVNIAN-----------NNTSGAGSG  391
Human--FOXN1     ----SSQ----PPF---------HQYSPGGGSYPIPYLGS-----------S---HYQYQ  254
Mouse--FOXN1     ----SSQ----PAF---------HQYSPGGGSYPVPYLGS-----------P---HYPYQ  254
Danre--FOXN4     ----FTQ-----AN---------QQCSPGGLYGN--YNSQ-----------N---LFPQP  215
HyFOXN1|ORF7     ----MSQ------Q---------QQQQQQNLYQRHDLHQQ-----------N---ILQHQ  155
HyFOXN1-Wenger   ----MSQ------Q---------Q-QQQQNLYQRHDLHQQ-----------N---ILQHQ  154
HyFOXI1c|ORF6    ------------------------------------------------------------    -
Drome--fd102C    ----LSP-----------------------------------------------------   92
Caeel--fkh-10    -----SS-----------------------------------------------------   11
Danre--FoxI1     ----MNS----PG------------------ITSTPYLSS-----------PNGGSYIQS  104
Human-FoxI1-a    ----FNG----PT------------------MTPPPYLPG-----------PNASPFLPQ   87
Mouse--FoxI1     ----LNG----PA------------------MTPPPYLPG-----------TNASPFLPQ   88

HyFOXP1|ORF14    HLKMNSVHDSSYHPVRRRGEAAAMVDIGQELRAKGKLFEDPDVRPPYTYASLIRQGVMDS  555
Drome--FOXP-d    ----------PYM------LERAGLDVQQEIHRNREFYKNADVRPPFTYASLIRQAIIDS  344
Danre-FOXP1-B    -------------------------IS-PDIVQNKEFYMNAEVRPPFTYASLIRQAILES  464
Mouse--FOXP-1    -------------------------ISSADIAQNQEFYKNAEVRPPFTYASLIRQAILES  509
Human-FOXP1-a    -------------------------ISSADIAQNQEFYKNAEVRPPFTYASLIRQAILES  481
Drome-Jumeau     LVKPL---Q---------------------QKVKLPPVGSPFPKPAYSYSCLIALALKNS  427
Human--FOXN1     RMAPQ---------------------------ASTDGHQPLFPKPIYSYSILIFMALKNS  287
Mouse--FOXN1     RIAPQ---------------------------ANAEGHQPLFPKPIYSYSILIFMALKNS  287
Danre--FOXN4     RI----------------------------TAHSQDLQPKCFPKPIYSYSCLIAMALKNS  247
HyFOXN1|ORF7     KYAHD---K---------------------NKENGSPGAKIFPKPVFSYSCLIAMALRNS  191
HyFOXN1-Wenger   KYAHD---K---------------------NKENGSPGAKIFPKPVFSYSCLIAMALRNS  190
HyFOXI1c|ORF6    ---------------------------------------------------MIAQAIMVS    9
Drome--fd102C    -YGHH---GSGSLPSVN--RMAALSTIS-LFPQTQRIFQPEEPKPQHSYIGLIAMAILSS  145
Caeel--fkh-10    --SNH---SLKDSP-------FPMIPLS-FDTSIMSPTECQQPKPQHSYIGLIAMAILSS   58
Danre--FoxI1     GFGSN---QRQFLPPPT--GF-GSADLG-WLSISSQQELFKMVRPPYSYSALIAMAIQNA  157
Human-FoxI1-a    AYGV----QRPLLPSVS--GL-GGSDLG-WLPIPSQEELMKLVRPPYSYSALIAMAIHGA  139
Mouse--FoxI1     AYGM----QRQLLP----------SDLG-WLPIPSQEELMKLVRPPYSYSALIAMAIHGA  133
                                                             p   y  6I  a6  s
```

### Fork head domain

```
HyFOXP1|ORF14   PNGELTLNEIYNWFMKNFAYFRKNTSTWKNAVRHNLSLHKCFVRKENF-------KGAVW  608
Drome--FOXP-d   PDKQLTLNEIYNWFQNTFCYFRRNAATWKNAVRHNLSLHKCFMRVENV-------KGAVW  397
Danre-FOXP1-B   PEKQLTLNEIYNWFTRMFAYFRRNAATWKNAVRHNLSLHKCFVRVENV-------KGAVW  517
Mouse--FOXP-1   PEKQLTLNEIYNWFTRMFAYFRRNAATWKNAVRHNLSLHKCFVRVENV-------KGAVW  562
Human-FOXP1-a   PEKQLTLNEIYNWFTRMFAYFRRNAATWKNAVRHNLSLHKCFVRVENV-------KGAVW  534
Drome-Jumeau    RAGSLPVSEIYSFLCQHFPYFENAPSGWKNSVRHNLSLNKCFEKIERPATNGNQRKGCRW  487
Human--FOXN1    KTGSLPVSEIYNFMTEHFPYFKTAPDGWKNSVRHNLSLNKCFEKVENKS-GSSSRKGCLW  346
Mouse--FOXN1    KTGSLPVSEIYNFMTEHFPYFKTAPDGWKNSVRHNLSLNKCFEKVENKS-GSSSRKGCLW  346
Danre--FOXN4    KTGSLPVSEIYSFMKEHFPYFKTAPDGWKNSVRHNLSLNKCFEKVENKM-SGSSRKGCLW  306
HyFOXN1|ORF7    DQGSLPVSEIYKYMQSRFPYFKTAPDGWKNSVRHNLSLNKAFCKLERPD-GTSQRKGCLW  250
HyFOXN1-Wenger  DQGSLPVSEIYKYMQSRFPYFKTAPDGWKNSVRHNLSLNKAFCKLERPD-GTSQRKGCLW  249
HyFOXI1c|ORF6   SCKRLTLSEIYEYMANTFEILKKRGTGWRNCVRHTLSLNECFLKLHRP----ENGRSCNW   65
Drome--fd102C   TDMKLVLSDIYQYILDNYPYFRSRGPGWRNSIRHNLSLNDCFIKSGRS----ANGKGHYW  201
Caeel--fkh-10   PQKKMVLAEVYEWIMNEYPYFRSRGAGWRNSIRHNLSLNDCFVKAGRA----ANGKGHYW  114
Danre--FoxI1    QDKKLTLSQIYQYVADNFPFYKKSKAGWQNSIRHNLSLNDCFKKVARDE--DDPGKGNYW  215
Human-FoxI1-a   PDKRLTLSQIYQYVADNFPFYNKSKAGWQNSIRHNLSLNDCFKKVPRDE--DDPGKGNYW  197
Mouse--FoxI1    PDQRLTLSQIYQYVADNFPFYNKSKAGWQNSIRHNLSLNDCFKKVPRDE--DDPGKGNYW  191
                    6 6  6Y 5    5         W N 6RHnLSL  cF 4          4g  W

HyFOXP1|ORF14   TVDDVE-------FFRRRMTKPG-LPKRDYYM------PGQQEETDSPPEPEDDYVADDL  654
Drome--FOXP-d   TVDEIE-------FYKRRPQRTAGIGNNLT---------GAT------NSPDTNYFVAM-  434
Danre-FOXP1-B   TVDELE-------FQKRRPQKISGSPALVK---------NIH------TT--LGYGPALS  553
Mouse--FOXP-1   TVDEVE-------FQKRRPQKISGNPSLIK---------NMQ------SS--HAYCTPLN  598
Human-FOXP1-a   TVDEVE-------FQKRRPQKISGNPSLIK---------NMQ------SS--HAYCTPLN  570
Drome-Jumeau    AMNPDRINKMDEEVQKWSRKDPAAIRGAMVYP--------QHLESLERGEMK-HGSAD-S  537
Human--FOXN1    ALNPAKIDKMQEELQKWKRKDPIAVRKSMAKP--------EELDSLIGDKRE-KLGSPLL  397
Mouse--FOXN1    ALNPSKIDKMQEELQKWKRKDPIAVRKSMAKP--------EELDSLIGDKRE-KLGSPLL  397
Danre--FOXN4    ALNPAKIDKMEEEMQKWKRKDLPAIRRSMANP--------DELDKLITDRPE-SCRQKSV  357
HyFOXN1|ORF7    SLKPEKKRPT--------------------------------------------------  260
HyFOXN1-Wenger  SLKPEKKDQLRREIRKWKKKHPEAIKASMANP--------DDLSIS----SD-SCDDDSM  296
HyFOXI1c|ORF6   TVHPAYFEAFSKGDYRKRRANRK-KSRSVSWIDDRNLAYPQTIGLPYIREPDIRMREDLS  124
Drome--fd102C   AIHPANMEDFRKGDFRRRKAQRK-VRKHMGLSVDDASTDSPSPPPLDLTTPP--------  252
Caeel--fkh-10   AVHPACVKDFERGDFRRRRAQRK-VRRHMGLQVEDGDSSDEEGSPGSD--PS--------  163
Danre--FoxI1    TLDPNCEKMFDNGNFRRKRKRRA-DGNAMSVKSEDALKLADTSSLMSASQP---------  265
Human-FoxI1-a   TLDPNCEKMFDNGNFRRKRKRKS-DVSSSTAS----LALEKTESSLPVDSPK--------  244
Mouse--FoxI1    TLDPNCEKMFDNGNFRRKRKRKS-DSSSSTSS----LASEKTENGLLASSPK--------  238
                 6
```

```
HyFOXP1|ORF14    P-------SMAMES--NQKHGDSGYSPKSNSEIGYEDTET-------VNIKEEYNDDYQ-  697
Drome--FOXP-d    -------------------------------NIGYVGLK---------------------  442
Danre-FOXP1-B    A-------AF------QASMAE-NIPLYTTASIGSPTLNSLA-----SVIREEMNGAMD-  593
Mouse--FOXP-1    A-------AL------QASMAENSIPLYTTASMGNPTLGSLA-----SAIREELNGAME-  639
Human-FOXP1-a    A-------AL------QASMAENSIPLYTTASMGNPTLGNLA-----SAIREELNGAME-  611
Drome-Jumeau     DV--------ELDSQ-------SEIEESS-----DLEEHEFE---------DTMVDAMLV  568
Human--FOXN1     GCP-----PPGLSGS-------GPIRPLAPPAGLSPPLHSLHPAPGPIPGKNPLQDLLM-  444
Mouse--FOXN1     GCP-----PPGLAGP-------GPIRPMAPSAGLSQPLHPMHPAPGPMPGKNPLQDLLG-  444
Danre--FOXN4     D-------------P-------GMTRLPSCPPGQTLPLA-----AQ-MQPQ-PVVTLSL-  389
HyFOXN1|ORF7     ------------------------------------------------------------    -
HyFOXN1-Wenger   E-------------D-------SKIQSP----EHTVPES-----ALDMPPSTPLLDDTV-  326
HyFOXI1c|ORF6    RVPVNDQHTNTLWQSYASSPAIK--------PEQYHTLNQF-----------PYDNRMT-  164
Drome--fd102C    ----PPSSQSALQ---------------LS-ALGYPYHQHY-------------IGQFF-  278
Caeel--fkh-10    ----PPIFPTALWN--------------FNCAPRAPIKSFT-------------IEAIL-  191
Danre--FoxI1     -------------SLQNSPTSSDPKSSPSPSAEHSPCFSNF-------------IGNMN-  298
Human-FoxI1-a    ----TTEPQDILDGASPGGTTSSPEKRPSPPPSGAPCLNSF-------------LSSMT-  286
Mouse--FoxI1     ----PTEPQEVLDTASPDTTTSSPEKRSSPAPSGTPCLNNF-------------LSTMT-  280


HyFOXP1|ORF14    -------HRSYERN-SDDCLPDEITM---------------------------VTAAPS-  721
Drome--FOXP-d    ------------------------------------------------------------    -
Danre-FOXP1-B    -------HGNS--N-GSDSSPGRSPLPAMHHISVKEEP---MDPEEHEGPLSLVTTANHS  640
Mouse--FOXP-1    -------HTNS--N-ESDSSPGRSPMQAVHPIHVKEEP---LDPEEAEGPLSLVTTANHS  686
Human-FOXP1-a    -------HTNS--N-ESDSSPGRSPMQAVHPVHVKEEP---LDPEEAEGPLSLVTTANHS  658
Drome-Jumeau     EEEDEEEDGDD-------------DEQIINDFDAEDERH--ANGNQ-ANNLPINHPL---  609
Human--FOXN1     GHTPS-CYGQTYLHLSPGLAPPGPPQPLFPQPDGHLELRAQPGTPQ-DSPLPAHTPPSHS  502
Mouse--FOXN1     GHAPS-CYGQTYPHLSPSLAPSGHQQPLFPQPDGHLELQAQPGTPQ-DSPLPAHTPPSHG  502
Danre--FOXN4     QCLPM-------------------------HQHLQLQLQNQSRLAP-SSPAPAQTPPLHT  423
HyFOXN1|ORF7     ------------------------------------------------------------    -
HyFOXN1-Wenger   KDEPV-------------------------DE-----------------------V---A  335
HyFOXI1c|ORF6    ----TLPQGSSNLNTSPFSSHSMHP----------YQQHPET-----Q--YPIIQTHNEG  203
Drome--fd102C    ----NRSSA-------PGMTHYSPPDPA-----LLMQRQEANNLDQTIQPTQLQQPHSHH  322
Caeel--fkh-10    ----EHH-----------------------------------------------------  194
Danre--FoxI1     ----SIMSGNAVR-SRDGSSAHLGD-----------------------------FTQHGMS  325
Human-FoxI1-a    ----AYVSGGSPT-SHPLVTPGLSPEPS-----DKTGQ-NSLTFNS-FSPLTNLSNHSGG  334
Mouse--FoxI1     ----AYVSGTNPI-SRSVATPGLSSEPI-----DKMGQ-NSLNFNS-YTPLTNLSSHGNG  328
```

```
HyFOXP1|ORF14    ------------------------------------------------------------      -
Drome--FOXP-d    ------------------------------------------------------------      -
Danre-FOXP1-B    PD---------------------------------------------------------F    643
Mouse--FOXP-1    PD---------------------------------------------------------F    689
Human-FOXP1-a    PD---------------------------------------------------------F    661
Drome-Jumeau     --------------------------------------------------LGQKSN--DF    617
Human--FOXN1     AK--------------------------------------LLAE----PSPARTMH--DT    518
Mouse--FOXN1     AK--------------------------------------LMAE----PSSARTMH--DT    518
Danre--FOXN4     VP--------------------------------------DMTNSSLPQHPAKQHT--DF    443
HyFOXN1|ORF7     ------------------------------------------------------------      -
HyFOXN1-Wenger   VE--------------------------------------DVAKELF-------GS--DL    348
HyFOXI1c|ORF6    SDAPQTSRYSNTRVKDSRMYQNHMDYLQENPPSQYPLDTHCLTNSAFGTRTYRQD-----    258
Drome--fd102C    QH---------------------FAYINSTT--------TTTIANMFSQTRKRQFDVASL    353
Caeel--fkh-10    ------------------------------------------------------------      -
Danre--FoxI1     GH------EISPPSEPGHLNTNRLNYYSASH-----------NNSGLINSISNHFSVNNL    368
Human-FoxI1-a    GD------WAN------PMPTNMLSYGGSVL-------------SQFSPHFYNSVNTSGV    369
Mouse--FoxI1     GE------WAN------PVATNALGYGGSVF-------------NQFSPHFYNSINTNGI    363


HyFOXP1|ORF14    ------------------------------------------------------------      -
Drome--FOXP-d    ------------------------------------------------------------      -
Danre-FOXP1-B    DHHRDYEDDHGTEDML--------------------------------------------    659
Mouse--FOXP-1    DHDRDYEDEPVNEDME--------------------------------------------    705
Human-FOXP1-a    DHDRDYEDEPVNEDME--------------------------------------------    677
Drome-Jumeau     DIEVG--------DLYDAIDI--------------------------------EDDKESV    637
Human--FOXN1     LLPDG--------DLGTDLDAINPS----------LTDFDFQGNLWE----QLKDDSLAL    556
Mouse--FOXN1     LLPDG--------DLGTDLDAINPS----------LTDFDFQGNLWE----QLKDDSLAL    556
Danre--FOXN4     YTVHT--------DVNSEVDALDPS----------IMDFAWQGNLWE----EMKDDSFNL    481
HyFOXN1|ORF7     ------------------------------------------------------------      -
HyFOXN1-Wenger   LFNHD--------D-------------------------LQADEFWN----ELMDTNMLS    371
HyFOXI1c|ORF6    --------------------------------------------FF--------------    260
Drome--fd102C    LAPDVQIVDIVSEDQESSVTPTTSARTTTQTHHTVITKQTIHREVVLGLEKPVQDADIDA    413
Caeel--fkh-10    ------------------------------------------------------------      -
Danre--FoxI1     IYHRD----------GSEV-----------------------------------------    377
Human-FoxI1-a    LYPRE----------GTEV-----------------------------------------    378
Mouse--FoxI1     LFPRE----------GTEV-----------------------------------------    372
```

```
HyFOXP1|ORF14      ------------------------------------------------------------   -
Drome--FOXP-d      ------------------------------------------------------------   -
Danre-FOXP1-B      ------------------------------------------------------------   -
Mouse--FOXP-1      ------------------------------------------------------------   -
Human-FOXP1-a      ------------------------------------------------------------   -
Drome-Jumeau       ---RRIISNDQHIIELNPADLNATDGYNQQPALKRARVDIN--YAIGPAGE---------   683
Human--FOXN1       DPLVL-VTSSPTSSSMPPPQ-PPPHCFPPGPCLTETGSGAGDLAAPGSGGSG--------   606
Mouse--FOXN1       DPLVL-VTSSPTSSSMLPPP-PAAHCFPPGPCLAETGNEAGELAPPGSGGSG--------   606
Danre--FOXN4       EA-LGTLSNSPLRL-------------------SDCDLDTSSVTPVSSAGGL--------   513
HyFOXN1|ORF7       ------------------------------------------------------------   -
HyFOXN1-Wenger     SHDRKYLSEDIVEL-------------------E-SS---VKIEPLSPSTNF--------   400
HyFOXI1c|ORF6      ------------------------------------------------------------   -
Drome--fd102C      DIDVE-VNVDV----VDDSIIPDTDSYTDGEDRKKTRGTLKQIFSIEDNNSYLIDGRLSS   468
Caeel--fkh-10      ------------------------------------------------------------   -
Danre--FoxI1       ------------------------------------------------------------   -
Human-FoxI1-a      ------------------------------------------------------------   -
Mouse--FoxI1       ------------------------------------------------------------   -
```

```
HyFOXP1|ORF14      ------------------------------------------------------------   -
Drome--FOXP-d      ------------------------------------------------------------   -
Danre-FOXP1-B      ------------------------------------------------------------   -
Mouse--FOXP-1      ------------------------------------------------------------   -
Human-FOXP1-a      ------------------------------------------------------------   -
Drome-Jumeau       -------------------------------------------L----------------   684
Human--FOXN1       ----------------------------ALG-----------DL----------------   611
Mouse--FOXN1       ----------------------------ALG-----------DM----------------   611
Danre--FOXN4       ----------------------------PYP-----------DL----------------   518
HyFOXN1|ORF7       ------------------------------------------------------------   -
HyFOXN1-Wenger     ----------------------------FSS-----------AI----------------   405
HyFOXI1c|ORF6      ------------------------------------------------------------   -
Drome--fd102C      FDADPDHDPDHDPEDLEQHNFSIASSAARSLGSSSSQHEESSSLEECPIQECIVAPQTAL   528
Caeel--fkh-10      ------------------------------------------------------------   -
Danre--FoxI1       ------------------------------------------------------------   -
Human-FoxI1-a      ------------------------------------------------------------   -
Mouse--FoxI1       ------------------------------------------------------------   -
```

```
HyFOXP1|ORF14     ------------------------------------------------------------    -
Drome--FOXP-d     ------------------------------------------------------------    -
Danre-FOXP1-B     ------------------------------------------------------------    -
Mouse--FOXP-1     ------------------------------------------------------------    -
Human-FOXP1-a     ------------------------------------------------------------    -
Drome-Jumeau      -----------------------------E--QQYGQKVKVQQVIQPQQHPPTYNR----  709
Human--FOXN1      -----------------------------HLTTLYSAFMELE------PTPPTAPAGPSV  636
Mouse--FOXN1      -----------------------------HLSTLYSAFVELE------STPSSAAAGPAV  636
Danre--FOXN4      -----------------------------QVTGLYSSYSAID------ALSNQY-----M  538
HyFOXN1|ORF7      ------------------------------------------------------------    -
HyFOXN1-Wenger    -----------------------------HQPT-LASF----------------------  413
HyFOXI1c|ORF6     ------------------------------------------------------------    -
Drome--fd102C     ISSMATAKTSTLSVPISDAYLELNQVDQHMLSRYYGSYIAAAARRA------SIDASNTS  582
Caeel--fkh-10     ------------------------------------------------------------    -
Danre--FoxI1      ------------------------------------------------------------    -
Human-FoxI1-a     ------------------------------------------------------------    -
Mouse--FoxI1      ------------------------------------------------------------    -


HyFOXP1|ORF14     -----------------    -
Drome--FOXP-d     -----------------    -
Danre-FOXP1-B     -----------------    -
Mouse--FOXP-1     -----------------    -
Human-FOXP1-a     -----------------    -
Drome-Jumeau      ----RKMPLVNRVI---  719
Human--FOXN1      YLSPSSKPVALA-----  648
Mouse--FOXN1      YLSPGSKPLALA-----  648
Danre--FOXN4      NTQGGTKPIVLL-----  550
HyFOXN1|ORF7      -----------------    -
HyFOXN1-Wenger    -----------------    -
HyFOXI1c|ORF6     -----------------    -
Drome--fd102C     RTSSITPPPKIEILSQK  599
Caeel--fkh-10     -----------------    -
Danre--FoxI1      -----------------    -
Human-FoxI1-a     -----------------    -
Mouse--FoxI1      -----------------    -
```

Phylogenetic tree of FOX-proteins:

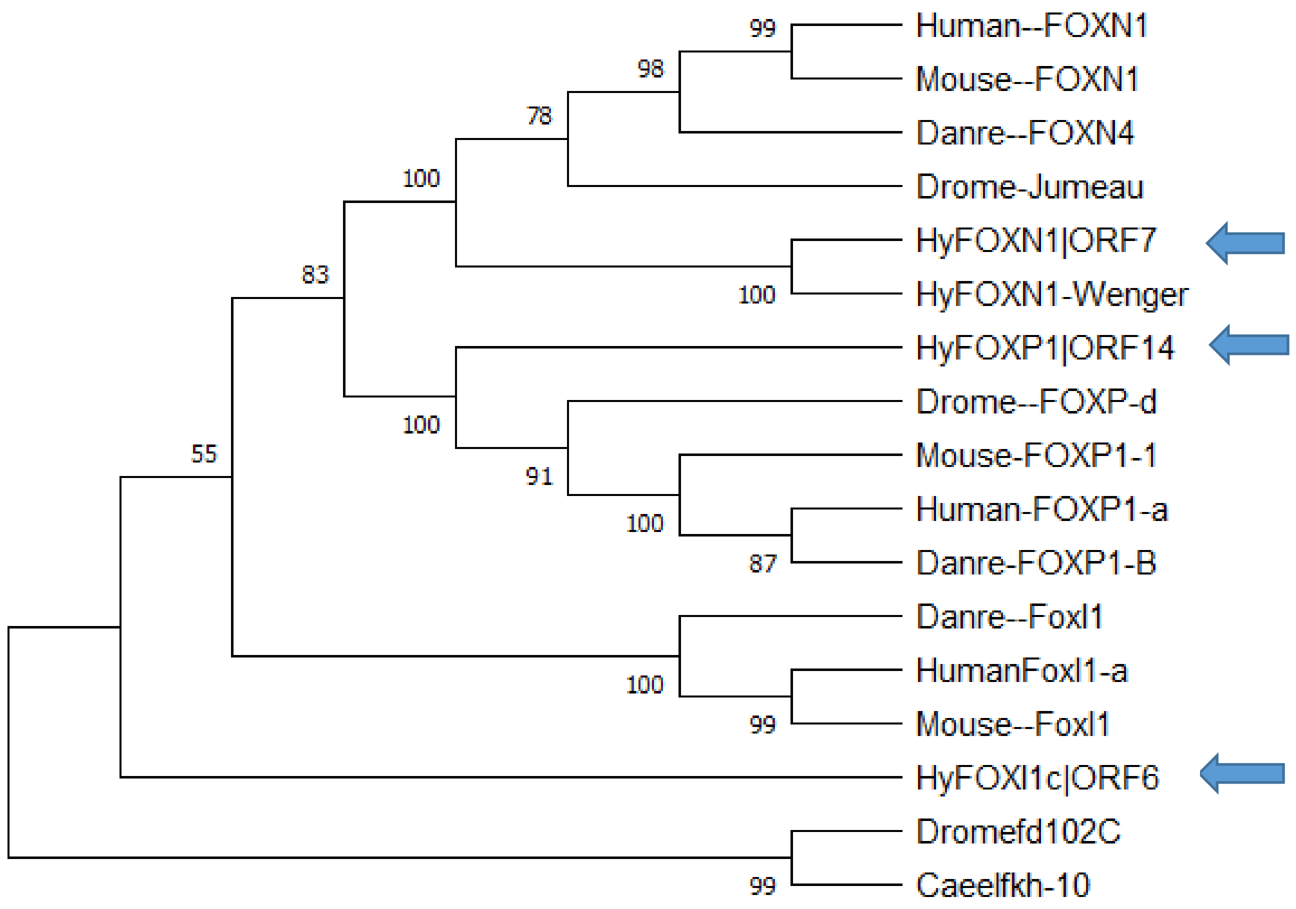


Forkhead domain is conserved in all Hydra-Forkhead proteins. TRINITY_DN1167_c0_g1 (HyFoxN1|ORF7) is identical with CDG72033.1 from Wenger (Wenger and Galliot, 2013)

**8.) APCD**

**t11061aep|APCD1_CHICK**

TRINITY_DN870_c0_g1 (**HyAPCDD1|ORF12**)

CDG68860.1 Hydra vulgaris Protein APCDD1 [Hydra vulgaris] (**HyAPCDD1-Wenger**)
Reference: Punctuated emergences of genetic and phenotypic innovations in eumetazoan, bilaterian, euteleostome, and hominidae ancestors.(Wenger and Galliot, 2013)

Alignment of **HyAPCDD1|ORF12, HyAPCDD1-Wenger** and other APCDD1 proteins from other species:

```
HyAPCDD1|ORF12    ~~~~~~~~~~~~~~MKSHFLLLVLVS---------------------FYSTKAANDKCED   25
HyAPCDD1-Wenger   ~~~~~~~~~~~~~~MKSHFLLLVLVS---------------------FYSTKAANDKCED   25
HumanAPCDD1pre    ~~~~~~MSWPRRLLLRYLFPALLLHGLGEGSALLHPDSRSHPRSLEKSAWRAFKESQCHH   54
MouseAPCDD1pre    ~~~~~~MSRVRRLLLGYLFPALLLHGLGEGSALLHPDSRSHPRSLEKSAWRAFKESQCHH   54
HumanAPCDD1-L1    ~~~~~~~~~MPAA-MLPYACVLVLLG---------AH-TAPA-AGEAGGSCLRWEPHCQQ   39
DanreAPCDD1L-1    MAGERLIHLLRTIWMVFLMQVLSVRG----SKLWEVP-TAPLVPHTNLSTRLFWEPQCQA   55
DanreAPCDD1L-2    MAGERLIHLLRTIWMVFLMQVLSVRG----SKLWEVP-TAPLVPHTNLSTRLFWEPQCQA   55
                                6      L 6                                 C

HyAPCDD1|ORF12    FSKHIDFHNPTFFYAPLSPIGSWTSSSCEVIPGPRYILRKWNIK---ESVFYGTIFHYSD   82
HyAPCDD1-Wenger   FSKHIDFHNPTFFYAPLSPIGSWTSSSCEVIPGPRYILRKWNIK---ESVFYGTIFHYSD   82
HumanAPCDD1pre    MLKHLHNGARITVQMPPTIEGHWVSTGCEVRSGPEFITRSYRFY--HNNTFKAYQFYYGS  112
MouseAPCDD1pre    MLKHLHNGARITVQMPPTIEGHWVSTGCEVRSGPEFMTRSYRFY--NNNTFKAYQFYYGS  112
HumanAPCDD1-L1    PLPD---RVPSTAILPPRLNGPWISTGCEVRPGPEFLTRAYTFY--PSRLFRAHQFYYED   94
DanreAPCDD1L-1    QLRHLQNEGRITATIPPKLEGHWVSSRCEVRPGPEFLTRSYTLYQSPTRLFRALQHYYSD  115
DanreAPCDD1L-2    QLRHLQNEGRITATIPPKLEGHWVSSRCEVRPGPEFLTRSYTLYQSPTRLFRALQHYYSD  115
                     h           P    G W S3 CEV  GP 56 R 5         F      Y

HyAPCDD1|ORF12    SWCTQPLFSVQVSGSYKIHAATATPQAKISKACEFKFEKIFYKTDNVTTLQETLELIKQK  142
HyAPCDD1-Wenger   SWCTQPLFSVQVSGSYKIHAATATPQAKISKACEFKFEKIFYKTDNVTTLQETLELIKQK  142
HumanAPCDD1pre    NRCTNPTYTLIIRGKIRLRQASWIIRGGTEA--DYQLHNVQVICHTEAVAEKLGQQVNRT  170
MouseAPCDD1pre    NRCTNPTYTLIIRGKIRLRQASWIIRGGTEA--DYQLHGVQVICHTEAVAEQLSRLVNRT  170
HumanAPCDD1-L1    PFCGEPAHSLLVKGKVRLRRASWVTRGATEA--DYHLHKVGIVFHSRRALVDVTGRLNQT  152
DanreAPCDD1L-1    SECQVPTYSLVIRGKLRLRQASWITRGATEA--EHHLHKVGMVIHSQKAIHHLATRLPSS  173
DanreAPCDD1L-2    SECHVPTYSLVIRGKLRLRQASWITLGATEA--EHHLHKVGMVIHSQKAIHHLATRLPSS  173
                    C  P  36 6 G  46  A3                 6                6
```

```
HyAPCDD1|ORF12   CIDAIPVLTMVK-ESDSENITWDLFINKNIKIENAAITKKHCRYYFGLHSNSYKKISMIG   201
HyAPCDD1-Wenger  CIDAIPVLTMVK-ESDSENITWDLFINKNIKIENAAITKKHCRYYFGLHSNSYKKISMIG   201
HumanAPCDD1pre   CPGF-----LADGGPWVQDVAYDLWREE---------NGCECTKAVNFA---MHELQLIR   213
MouseAPCDD1pre   CPGF-----LAPGGPWVQDVAYDLWQEE---------SNHECTKAVNFA---MHELQLIR   213
HumanAPCDD1-L1   RAGRDCARRLPPARAWLPGALYELRSAR---------AQGDCLEALGLT---MHELSLVR   200
DanreAPCDD1L-1   CLGLKAE------GHLVLHRFYELFNAK---------AEKDCLGALGFS---MMELELLR   215
DanreAPCDD1L-2   CLGLKAE------GHLVLHRFYELFNAK---------AEKDCLGALGFS---MMELELLR   215
                 c                    5 L                 C             6 66

HyAPCDD1|ORF12   LKKDFS-------NENDVLYFSDIPPKHHEKSKDYIGDGFQYGLF--RVHRPNCPVCELI   252
HyAPCDD1-Wenger  LKKDFS-------NENDVLYFSDIPPKHHEKSKDYIGDGFQYGLF--RVHRPNCPVCELI   252
HumanAPCDD1pre   VEKQYLHHN--LDHLVEELFLGDIHTDATQ-RMFYRPSSYQPPLQNAKNHDHACIACRII   270
MouseAPCDD1pre   VEKQYPHHS--LDHLVEELFLGDIHTDATQ-RVFYRPSSYQPPLQNAKNHNHACIACRII   270
HumanAPCDD1-L1   VQRRLQPQPRASPRLVEELYLGDIHTDPAE-RRHYRPTGYQRPLQSALHHVQPCPACGLI   259
DanreAPCDD1L-1   VETHHHPHG----RPAQELFLGDVHTEWNE-RVHYRPSGYQEPLQNAMHHIHPCPVCGLV   270
DanreAPCDD1L-2   VETHHHPHG----RPAQELFLGDVHTDWNE-RVHYRPSGYQEPLQNAMHHIHPCPVCGLV   270
                 6                 L5  D6     2    Y    5Q  L     H   C  C 66

HyAPCDD1|ORF12   NVSEEQ--PILPPY-DHSQDFIGGWVSSVCEAVNDNTFQTRYFEFYPPNGENFGDVSLQI   309
HyAPCDD1-Wenger  NVSEEQ--PILPPY-DHSQDFIGGWVSSVCEAVNDNTFQTRYFEFYPPNGENFGDVSLQI   309
HumanAPCDD1pre   YRSDEHHPPILPPKADLTIGLHGEWVSQRCEVRPEVLFLTRHFIFHDNNNTWEG----HY   326
MouseAPCDD1pre   FRSDEHHPPILPPKADLTIGLHGEWVSQRCEVRPEVLFLTRHFIFHDNNNTWEG----HY   326
HumanAPCDD1-L1   ARSDVHHPPVLPPPLALPLHLGGWWVSSGCEVRPAVLFLTRLFTFHGHSRSWEG----YY   315
DanreAPCDD1L-1   YRASEQHPPILPPSPSIPLNLNGNWVSQRCESRPAVLFLTRLFVFNEEQHTWEG----TY   326
DanreAPCDD1L-2   YRASEQHPPILPPSPSIPLNLNGNWVSQRCESRPAVLFLTRLFAFNEEQRTWEG----TY   326
                     e   P6LPP         G WVS  CE      F TR F F        G

HyAPCDD1|ORF12   NLYLDQNCKDKRLIIKARGKVDEISPHESVDGILKLSLILKSMLLTGYNGPTLGIMR-QP   368
HyAPCDD1-Wenger  NLYLDQNCKDKRLIIKARGKVDEISPHESVDGILKLSLILKSMLLTGYNGPTLGIMR-QP   368
HumanAPCDD1pre   YHYSDPVCKHPTFSIYARGRYSRGVLSSRVMGGTEFVFKVNHMKVTPMDAATASLLNVFN   386
MouseAPCDD1pre   YHYSDPVCKHPTFTIYARGRYSRGVLSSKVMGGTEFVFKVNHMKVTPMDAATASLLNVFS   386
HumanAPCDD1-L1   HHFSDPACRQPTFTVYAAGRYTRGTPSTRVRGGTELVFEVTRAHVTPMDQVTTAMLNFSE   375
DanreAPCDD1L-1   HHYSDPTCRQPSFTLSASGNYVKVGQSIKVRGATELVFKVTHAKVIVFDRALLRELNSTQ   386
DanreAPCDD1L-2   HHYSDPTCRQPSFTLSASGNYVKVGQSIKVRGATELVFKVTHAKVIVFDRALLRELNSTQ   386
                   5 D  C4     6 A G          V G       6    6   1      6
```

```
HyAPCDD1|ORF12    DNRCGDSEKWKYGFEQDVTHTNGCKMLGLIVPKNS-DMFIRVVSRQGKKELYIQDNDEKG  427
HyAPCDD1-Wenger   DNRCGDSEKWKYGFEQDVTHTNGCKMLGLIVPKNS-DMFIRVVSRQGKKELYIQDNDEKG  427
HumanAPCDD1pre    GNECGAEGSWQVGIQQDVTHTNGCVALGIKLPHTEYEIFKMEQDARGRYLLFNGQRPSDG  446
MouseAPCDD1pre    GNECGAEGSWQVGIQQDVTHTNGCVALGIKLPHTEYEIFKMEQDTRGRYLLFNGQRPSDG  446
HumanAPCDD1-L1    PSSCGGAGAWSMGTERDVTATNGCLPLGIRLPHVEYELFKMEQDPLGQSLLFIGQRPTDG  435
DanreAPCDD1L-1    NGRCGQAGGWEAGIEQDITWTNGCDALGIRLPHKEYELFKMEVDRKGHPLLFNGERPTDG  446
DanreAPCDD1L-2    NGRCGQAGGWEAGIEQDITWTNGCDALGIRLPHKEYELFKMEVDRKGHPLLFNGERPTDG  446
                     CG    W  G 2qD6T TNGC  LG6 6P     6F       G   L5       G

HyAPCDD1|ORF12    TFFTDYLIPCNIASANISKMTTSELPTPRINLIPEAPDFDIMDQDFDSPSST--------  479
HyAPCDD1-Wenger   TFFTDYLIPCNIASANISKMTTSELPTPRINLIPEAPDFDIMDQDFDSPSST--------  479
HumanAPCDD1pre    SS------------PDRPEKRATSYQMPLVQCASSSPRAEDLAED--SGSSLYGRAP---  489
MouseAPCDD1pre    SS------------PDRPEKRATSYQMPLVQCASSSPRAEELLED--SQGHLYGRAA---  489
HumanAPCDD1-L1    SS------------PDTPEKRPTSYQAPLVLCHGEAPDFSRPPQH--RPSLQ-----KHP  476
DanreAPCDD1L-1    SS------------PERPAKRPTSFQTPMVQCSTSVRV---EPHH--SPYSDHSKGPKKS  489
DanreAPCDD1L-2    SS------------PERPAKRPTSFQTPMVQCSTSVRV---EPHH--SPYSDHSKGPKKS  489
                  3                     3    P 6                 s

HyAPCDD1|ORF12    -ASSYRCSLVLLV-------LLKMLMQFFD  501
HyAPCDD1-Wenger   -ASSYRCSLVLLV-------LLKMLMQFFD  501
HumanAPCDD1pre    ---GRHTWSLLLAALACLVPLLHWNIRR~~  514
MouseAPCDD1pre    ---GRTAGSLLLPAFVSLWTLPHWRILR~~  514
HumanAPCDD1-L1    STGGLHIAPFPLLPLVLGLAFLHWL~~~~~  501
DanreAPCDD1L-1    SANALHSSVI-LL-----SLINAWIIHLF~  512
DanreAPCDD1L-2    SANALHSSVI-LL-----SLINAWIIHLF~  512
                             L        l
```

HyAPCDD1|ORF12 is identical with HyAPCDD1-Wenger (Wenger and Galliot, 2013).

9.) **IRX6**

**t16018aep|IRX6_HUMAN**
TRINITY_DN3014_c0_g1 (**HyIRX2|ORF7**)

Alignment of **HyIRX2|ORF7** and IRX proteins from other animals:

```
HyIRX2|ORF7  ------------------------------------------------------------     -
Mouse--IRX2  ------------------------------------------------------------     -
DromeMirror  MTVHSNETGICLVMNAVSTHLATSSPNTPPTNAPQSPPSTLPTLACPAAQPLSAPGIAGP    60
Danre-IRX3b  ------------------------------------------------------------     -
Danre-IRX4b  ------------------------------------------------------------     -
Human--IRX6  ------------------------------------------------------------     -


HyIRX2|ORF7  -------MRLQEGESSSNLSATKAIFPVQFLKNNILFPTMEY------------------    35
Mouse--IRX2  ----------------------------------MSYPQ-GYLYQAP-------------    12
DromeMirror  GHSGMVTSTVPPGARST--------SPCGPSAGLPALSLVGAPPPGPHSSAPGGAPAPGA   112
Danre-IRX3b  ----------------------------------MSLPQLGYKYIRP-------LYSTER    19
Danre-IRX4b  ----------------------------------MSFTQFGYSYPAA-------PQLLMS    19
Human--IRX6  ----------------------------------MSFPHFGHPYRGA-------SQFLAS    19
                                                     g

HyIRX2|ORF7  --NASLCREKKMFCHLKEDS----CTSYCLCRAG-RTYNRIL---------------LPS    73
Mouse--IRX2  ------------------GSL----ALYSCPAYGASALAAPRSEELARSASGSAFSPYPG    50
DromeMirror  GNPPNRCCDTGRTIYTDPVSGQ----TICSCQYDMLNYQRLA------AAGGVPLGVYPE   162
Danre-IRX3b  R---------GGAEISVTGSG---------------------------SLSSALSGVYGA    43
Danre-IRX4b  SNALSTCFESSG-SLLDSGVPTSPQNSLYCPVYESRLVASSR------HELIPT-AVYGN    71
Human--IRX6  ASSSTTCCESTQRSVSDVASGSTPAPALCCAPYDSRLLGSAR------PELGAALGIYGA    73
                                s                                     y
```

```
HyIRX2|ORF7  TVEKKHTICSQNVEDCLSNSWH---------MNSGIKNNYDSFLLN-RLSSSYNKYNNES  123
Mouse--IRX2  SAAFT-AQAATGFGSPLQYSADAAAAAAAGFPSYV-GSPYDTHTTGMTGAISYHPYGS--  106
DromeMirror  GMSAY--L--------SGIAADQPPFY-ANPAGIDLKENLVAGASPWPYPSMYHPYDA--  209
Danre-IRX3b  PFAST----AQGYSAFLPYSSDLSVL-NQLSSQYEFKDSPGLQHAGFP-HAAFYPYGH--   95
Danre-IRX4b  SCSKN-----HVYDSCSTYGPDSAACYPLG--KLNEKDGVQTGHSRVTQSSAYYPYDY--  122
Human--IRX6  PYAAA--AAAQSYPGYLPYSPEPPSLYGALNPQYEFKEAAGSFTSSLAQPGAYYPYER--  129
                                                 k               5 pY

                                                    Homeobox domain
HyIRX2|ORF7  CSILDY--------ALLASARRKNATKETTSVLKSWLNEHRKNPYPSKSEKVMLAIMTKM  175
Mouse--IRX2  -AAYPYQ--------LNDPAYRKNATRDATATLKAWLNEHRKNPYPTKGEKIMLAIITKM  157
DromeMirror  -AFAGYPFNSYGMD--LNGARRKNATRETTSTLKAWLNEHKKNPYPTKGEKIMLAIITKM  266
Danre-IRX3b  -Q---YQ--------FGDPSRPKNATRESTSTLKAWLSEHRKNPYPTKGEKIMLAIITKM  143
Danre-IRX4b  -PIGQYSYDRYGYGSVDVGTRRKNATRETTSTLKAWLQEHKKNPYPTKGEKIMLAIITKM  181
Human--IRX6  -TLGQYQYERYGAVELSGAGRRKNATRETTSTLKAWLNEHRKNPYPTKGEKIMLAIITKM  188
                  Y              rrKNAT4e TstLKaWL EH4KNPYP3KgEK6MLAI6TKM

HyIRX2|ORF7  TLTQVSTWFANARRRLKKENKMTWSPRKKSNSKNSLKEGSSCSSEPIPEEMDI-------  228
Mouse--IRX2  TLTQVSTWFANARRRLKKENKMTWAPRNKSEDEDEDEGDASRSKE---ESSDKAQDGTET  214
DromeMirror  TLTQVSTWFANARRRLKKENKMTWEPRNRVDDDDANIDDDDDKNT---EDNDLLDAKDSG  323
Danre-IRX3b  TLTQVSTWFANARRRLKKENKMTWVPKTRTDEDGNVYTSDNEDAEKRDE--DE-------  194
Danre-IRX4b  TLTQVSTWFANARRRLKKENKMTWSPRNKNSDEKECDDDQEDLDEAQEEPIKTEQDFNDN  241
Human--IRX6  TLTQVSTWFANARRRLKKENKMTWAPKNKGGEERKAEGGEEDSLGCLTADTKEVTASQEA  248
             TLTQVSTWFANARRRLKKENKMTW P4 4                   e

HyIRX2|ORF7  -------DIKEQIFSAQDCNINT-----------------------------NEENSGKN  252
Mouse--IRX2  SAEDEGISLHVDSLTDHSCSAESDGEKLPCRAGDALCESGSECKDKFEDLEDEED-EEDE  273
DromeMirror  VGST-DDKDRSG---------------------------------RLGDMMTDRPGESNN  349
Danre-IRX3b  -----EIDLENI---------------------------------DTEDIED-----KQD  211
Danre-IRX4b  HGKD-DTDQLHS---------------------------------DLDDFDLVES-DGSE  266
Human--IRX6  R------GLRLS---------------------------------DLEDLEEEEE-EEEE  268
                                                             d
```

```
HyIRX2|ORF7   CIETPFSVAQDTKIQRPSVICVLESPTFDQIKNRFSRDPSPV-DHLQKWVDGCSQEKPDD  311
Mouse--IRX2   CERDLA---------PPKP--VTSSPL---------------TGVEAPLLSPAPE-----  302
DromeMirror   SEWSES---------RPGS--PNGSPD-------LYDRPGSMPPGAHPLFHPAAL-----  386
Danre-IRX3b   CDYQDD---------D------KSTPK--------GSDSEEYDD----------ARAEKR  238
Danre-IRX4b   CESKPSFVVH-----VHSE--TSDHPE-THFKDAFHESVTELTRGVHKF-------SEDR  311
Human--IRX6   AEDEE--VVA-----T------------------AGDRLTEFRKGAQSLPGPCAAAREGR  303
                                         p

HyIRX2|ORF7   LV---CI-K----DLTPPLT-PIDNRVTTFTSQKLFEAFDAAKINKNNDTQYIKSNSYAS  362
Mouse--IRX2   --------------A-APRGGSG--------------GKTPLG-------SRTSPGAPPP  326
DromeMirror   -----------HHHFRPPAGSPPDIAAYHHHQQLLQQHQQAQ-------QNSLQTAVGG  428
Danre-IRX3b   IIEDEEQIK------KSPAEEQ----------------------------EP--SNNISP  262
Danre-IRX4b   L-------R-------APAEDHQMAK--------FY-L--QQG------------QKTI  333
Human--IRX6   LERRECGLAAPRFSFNDPSGSE-EAD--------FLSA--ETG-------SPRLTMHYPC  345
                                 P

HyIRX2|ORF7   EPISLIFSSKQ------------PVCSTKECSVEVLMPESLSSSFTPPSPS---------  401
Mouse--IRX2   ASKPKLWSLAEIATSDLKQPSLGPGCGPPGLPA--AAA--PASTGAPPGGSPYS------  376
DromeMirror   TAKPRIWSLADMASKDSKDSSSGAKDNHPELPP--AHPGFYGHPGQQPSPGKILSPL---  483
Danre-IRX3b   ALKPKIWSLAETATTPDSP--------KKTWI----QRNCDAQTVRNPLH----------  300
Danre-IRX4b   ETKPKIWSLAQTATSLNQVDYSSCMHKRTACP----SSNCDSDSE---MTKRKQESP---  383
Human--IRX6   LEKPRIWSLAHTATASAVEGAPPAR-PRPR------SPECRMIPGQPPASARRLSVPRDS  398
                kp 65Sla  a                                 p

HyIRX2|ORF7   ------FTPQPSPTDTTLYKRDFSNKNDK-----------N---------NIKNGCDNQS  435
Mouse--IRX2   ------ASPLLG---RHLY---YTSPF-----------YGNYTNYGN----LNAALQGQG  409
DromeMirror   AARIPNYSPYVR---PDLYR-GFYGPAAAHLGAPTQEFLEHQRTFGASLAAHNGPLGMNP  539
Danre-IRX3b   ---VQNWTKMA-----------LSAHQMAF--------TSH---------YLGLKHQSIT  329
Danre-IRX4b   VATLRNWVDGV-----------FHDPLFRH----------N---------NLNQTFSNST  413
Human--IRX6   ACDESSCIPKA-----------FGNPKFAL----------Q---------GLPLNCAPCP  428
```

```
HyIRX2|ORF7   ------------NSADFRSQGLTT-------SRELEAVLALTAL-------------CKT  463
Mouse--IRX2   LLRYNTAASS--PGETLHAMPKAASDTGKAGSHSLESHYRPPG--GGYEPKKDTSEGCAV  465
DromeMirror   -LLWKAAVSGAANGHHFAPLSLTTSG-GSSGGQ----QVAPPPVASPSASSSSSSMGCDV  593
Danre-IRX3b   -NIH--------VKHAEQR-----TH-SL-------------------------------  343
Danre-IRX4b   -ELWTDAISSQNNYHEHSGTPITSSH-A--------------------------------  439
Human--IRX6   -RRSEPVVQCQYPSGAEAG-----------------------------------------  446


HyIRX2|ORF7   -------------------------------------------------    -
Mouse--IRX2   VGAGVQTYL----------------------------------------  474
DromeMirror   VHIPTSSGQSAAQHMMGPISSNSTASSSSSHSGKISPGVNVTSLSAKP  641
Danre-IRX3b   -------------------------------------------------    -
Danre-IRX4b   -------------------------------------------------    -
Human--IRX6   -------------------------------------------------    -
```

HyIRX2|ORF7 has a conserved homeobox domain.

---

**10.) ARX**

**t21636aep|AL_DROME**
TRINITY_DN2447_c0_g1 (**HyPrdl|ORF7**)

CAA75669.1 prdl-b protein, partial [Hydra vulgaris] (**HyPrdl-Gauchat**)
Reference: prdl-a, a gene marker for hydra apical differentiation related to triploblastic paired-like head-specific genes.(Gauchat et al., 1998)

XP_002168027.1 PREDICTED: aristaless-related homeobox protein-like [Hydra vulgaris] (**HyARX-pred.**)

Alignment of HyPrdl|ORF7, HyPrdl-Gauchat, XP_002168027.1 (HyARX-pred.) and other ARX proteins:

```
CAEEL-Alr         ------------------------------------------------------------     -
Danre-ARX         MSSQYDD-DSRDRSECKSKSPTVLSSYCIDSILGRRSPCKVRQLGA-QSLPAPVRPD---    55
Human-ARX         MSNQYQEEGCSERPECKSKSPTLLSSYCIDSILGRRSPCKMRLLGAAQSLPAPLTSRADP    60
Mouse-ARX         MSNQYQEEGCSERPECKSKSPTLLSSYCIDSILGRRSPCKMRLLGAAQSLPAPLASRADQ    60
DROME-AL          ------------------------------------------------------------     -
HyPrdl|ORF7       ----------------------------------------------------MIVYNQSH     8
HyARX-pred.       ----------------------------------------------------MIVYNQSH     8
HyPrdl-Gauchat    ----------------------------------------------------MIVYNQSH     8


CAEEL-Alr         ------------------------------------------------------------     -
Danre-ARX         -HEMTTEVTSKENSFDSDMHLPPKL------RRLYGPGGKYLDSGRGFHEHLE-------   101
Human-ARX         EKAVQGSPKSSSAPFEAELHLPPKL------RRLYGPGGRLLQGAAAAAAAAAA-AAAAA   113
Mouse-ARX         EKAMQGSPKSSSAPFEAELHLPPKL------RRLYGPGGRLLQGAAAAAAAAAAAAAAAA   114
DROME-AL          ----------------------------------MGISEEIKLE----------------    10
HyPrdl|ORF7       NSEMPELQQEDNAAFDQNMNIDRKKHTSYSIRDILGLKEGIKSNGE--------------    54
HyARX-pred.       NSEMPELQQEDNAAFDQNMNIDRKKHTSYSIRDILGLKEGIKSNGE--------------    54
HyPrdl-Gauchat    NSEMPELQQEDNAAFDQNMNIDRKKHTSYSIRDILGLKEGIKSNGE--------------    54
                                f        k       r   g        g

CAEEL-Alr         ------------------------------------------MPELK-----KEDSSKDE    13
Danre-ARX         ---------------------KGER----------ERLLDQACESLKISQAPQVSISR-S   129
Human-ARX         AATATAGPRGEAPPPPPPTARPGERPDGAGAAAAAAAAAAAAWDTLKISQAPQVSISR-S   172
Mouse-ARX         TATGTAGPRGEVPPPPPPAARPGERQDSAGAVAA--AAAAAAWDTLKISQAPQVSISR-S   171
DROME-AL          -----------------------------------------ELPQ---------------    14
HyPrdl|ORF7       -------------------------------VSLGNSFVD-AIPSTRSPQFPESDSVS-S    81
HyARX-pred.       -------------------------------VSLGNSFVD-AIPSTRSPQFPESDSVS-S    81
HyPrdl-Gauchat    -------------------------------VSLGNSFVD-AIPSTRSPQFPESDSVS-S    81
                                                           a       q p       s
```

```
CAEEL-Alr        KDLDMNCPPLHR----------PNG-----------------------------------   28
Danre-ARX        KSYRENAP-FSQSD----EGQS----------------------------PEHMAQELVE  156
Human-ARX        KSYRENGAPFVPPPPALDELGGPGGVTHPEERLGVAGGPGSAPAAGGGTGTEDDEEELLE  232
Mouse-ARX        KSYRENGAPFVPPPPALDELSGPGGVAHPEERLSAASGPGSAPAAGGGTGAEDDEEELLE  231
DROME-AL         ------------------------------------------------------------    -
HyPrdl|ORF7      DTYRQQALSM--------------------------------------------------   91
HyARX-pred.      ------------------------------------------------------------    -
HyPrdl-Gauchat   ------------------------------------------------------------    -


CAEEL-Alr        --ADLNQYSKSLMEQLQAQLFANPALQFPSFPPAFSIA--------------ALTNNQHE   72
Danre-ARX        LS--------------TLKFEED--------------------------VVKEEACGDNS  176
Human-ARX        DEEDEDEE-EELLEDDEEELLEDDARALLKEPRRCPVAATGAVAAA---AAAAVATEGGE  288
Mouse-ARX        DEEDEEEE-EELLEDDDEELLEDDARALLKEPRRCSVATTGTVAAAAAAAAAAVATEGGE  290
DROME-AL         ----------------EAKLAHPDAVVLVDRAPGSSAASAG------AALTVSMSVSGGA   52
HyPrdl|ORF7      -PSKKRRY-RTTFT--THQLDELERVFNRTHYPESSRSPTS------PSLRVQSLNIGGS  141
HyARX-pred.      --------------------------------PESSRSPTS------PSLRVQSLNIGGS  103
HyPrdl-Gauchat   --------------------------------PESSRSPTS------PSLRVQSLNIGGS  103
                                                    s                     gg


                                                      Homeobox domain
CAEEL-Alr        LKEDDGKKTPTGDNIL-EAASVLDNRENGSPSDGTNSPDDNGKRKQRRYRTTFSAFQLDE  131
Danre-ARX        LSPKDEE------SLH-NDGDVKDGEDSVCLSAGSDSEEGMLKRKQRRYRTTFTSYQLEE  229
Human-ARX        LSPKEEL------LLHPEDAEGKDGEDSVCLSAGSDSEEGLLKRKQRRYRTTFTSYQLEE  342
Mouse-ARX        LSPKEEL------LLHPEDAEGKDGEDSVCLSAGSDSEEGLLKRKQRRYRTTFTSYQLEE  344
DROME-AL         PSGASGASGGTNSPVSDGNSDC-------------EADEYAPKRKQRRYRTTFTSFQLEE   99
HyPrdl|ORF7      ISGDDCE--TLDSPTLDNDGALTSDEVSPTVLDTYRQQALSMPSKKRRYRTTFTTHQLDE  199
HyARX-pred.      ISGDDCE--TLDSPTLDNDGALTSDEVSPTVLDTYRQQALSMPSKKRRYRTTFTTHQLDE  161
HyPrdl-Gauchat   ISGDDCE--TLDSPTLDNDGALTSDEVSPTVLDTYRQQALSMPSKKRRYRTTFTTHQLDE  161
                  s                       e s                K RRYRTTF3  QL E
```

```
CAEEL-Alr          LEKVFARTHYPDVFTREELATRVQLTEARVQVWFQNRRAKYRKQERSST-HHPYQAPMSI  190
Danre-ARX          LERAFQKTHYPDVFTREELAMRLDLTEARVQVWFQNRRAKWRKREKAGVQAHPTGLPFPG  289
Human-ARX          LERAFQKTHYPDVFTREELAMRLDLTEARVQVWFQNRRAKWRKREKAGAQTHPPGLPFPG  402
Mouse-ARX          LERAFQKTHYPDVFTREELAMRLDLTEARVQVWFQNRRAKWRKREKAGAQTHPPGLPFPG  404
DROME-AL           LEKAFSRTHYPDVFTREELAMKIGLTEARIQVWFQNRRAKWRKQEKVGPQSHPYNPYLPG  159
HyPrdl|ORF7        LERVFNRTHYPDIFLREEMAVKLGLTEARIQVWFQNRRAKWRKRNKSTSLSSPTSPPPLP  259
HyARX-pred.        LERVFNRTHYPDIFLREEMAVKLGLTEARIQVWFQNRRAKWRKRNKSTSLSSPTSPPPLP  221
HyPrdl-Gauchat     LERVFNRTHYPDIFLREEMAVKLGLTEARIQVWFQNRRAKWRKRNKSTSLSSPTSPPPLP  221
                   LE4 F 4THYPD6F REE6A 46 LTEAR6QVWFQNRRAK5RKr 4      P   p

CAEEL-Alr          PNSNGDNPYQMMLSQEAIFA--------AINQQAAAHL----LNEQV--RIATSDRRSQS  236
Danre-ARX          PLAAA-HPLSHYLEGGPFPPHPHPALESAWTAAAA-AAA-------------------AF  328
Human-ARX          PLSAT-HPLSPYLDASPFPPH-HPALDSAWTAAAAAAAA-------------------AF  441
Mouse-ARX          PLSAT-HPLSPYLDASPFPPH-HPALDSAWTAAAAAAAA-------------------AF  443
DROME-AL           GAATMQTVVGAALPPNPFTHLG-FQLRKPFDAQHAANLAAFRYPHLSAAPMIPSGYFNQF  218
HyPrdl|ORF7        QLNNYLKPYTPTQPSTPFNN------------SISNSLFLKSLNHPS-----QTEKFNRF  302
HyARX-pred.        QLNNYLKPYTPTQPSTPFNN------------SISNSLFLKSLNHPS-----QTEKFNRF  264
HyPrdl-Gauchat     QLNNYLK-----------------------------------------------------  228
                    l     p        pf                                         f

CAEEL-Alr          PSVPVTTSSPILPTSVTPTFQNADALNMLFGGITPVTQQLLYVQQFSRAMDAFRTQLLGS  296
Danre-ARX          PGL----APP----------------------------------------PN----SSAL  340
Human-ARX          PSL----PPP----------------------------------------PG----SASL  453
Mouse-ARX          PSL----PPP----------------------------------------PG----SASL  455
DROME-AL           QRA----PPHMLPHGMAGMYSPSSS------------------FQ---SLLA----NMTA  249
HyPrdl|ORF7        NGS----LST----KLEPWYENNIRY---------I-HRNIYNYDS-RSINN----GYRL  339
HyARX-pred.        NGS----LST----KLEPWYENNIRY---------I-HRNIYNYDS-RSINN----GYRL  301
HyPrdl-Gauchat     ------------------------------------------------------------    -
```

```
CAEEL-Alr       VPAGATAEVTDVVALKTEDSKSNSRA----------G---------SSSPTPT-----ES  332
Danre-ARX       PP-ATPLGLGTF--LGTAMFRHPA------------------------------------  361
Human-ARX       PPSGAPLGLSTF--LGAAVFRHPA------------------------------------  475
Mouse-ARX       PPSGAPLGLSTF--LGAAVFRHPA------------------------------------  477
DROME-AL        VPRGTPLGKPPALLVGSPDLHSPNHMLASPPTSPASGHASQHQQHPTAHPPPPQAPPQMP  309
HyPrdl|ORF7     IDKDQPLSWWPV--HGQPV-----------------------------------------  356
HyARX-pred.     IDKDQPLSWWPV--HGQPV-----------------------------------------  318
HyPrdl-Gauchat  ------------------------------------------------------------    -
                     pl        g

CAEEL-Alr       SGAAATSTSPTNFAD--MNSLISDVKPKEESS----------------------------  362
Danre-ARX       ------------FIGPTFGRLFSSMGPLTSASTAAALLRQTAPPVESPVQPSAALPEPPS  409
Human-ARX       ------------FISPAFGRLFSTMAPLTSASTAAALLRQPTPAVEGAVASGA-LA----  518
Mouse-ARX       ------------FISPAFGRLFSTMAPLTSASTAAALLRQPTPAVEGAVASGA-LA----  520
DROME-AL        VGVQPAQLSPQHLVGIALTQQASSLSPTQTSPVALTLSHSP---QRQLPPPSHQAPPPPP  366
HyPrdl|ORF7     ------------------------------------------------------------    -
HyARX-pred.     ------------------------------------------------------------    -
HyPrdl-Gauchat  ------------------------------------------------------------    -

CAEEL-Alr       ---------------------------------------------    -
Danre-ARX       SSSSTAADRRASSIAALRLKAKEHSAQLTQLNILPSGTAGKEVC-  453
Human-ARX       DPATAAADRRASSIAALRLKAKEHAAQLTQLNILPGTSTGKEVC-  562
Mouse-ARX       DPATAAADRRASSIAALRLKAKEHAAQLTQLNILPGTSTGKEVC-  564
DROME-AL        RAATPPEDRRTSSIAALRLKAREHELKLELLR---QNGHGNDVVS  408
HyPrdl|ORF7     ---------------------------------------------    -
HyARX-pred.     ---------------------------------------------    -
HyPrdl-Gauchat  ---------------------------------------------    -
```

HyPrdl|ORF7 is identical with HyPrdl-Gauchat (Gauchat et al., 1998). The homeobox is conserved.

Phylogenetic tree of ARX proteins:

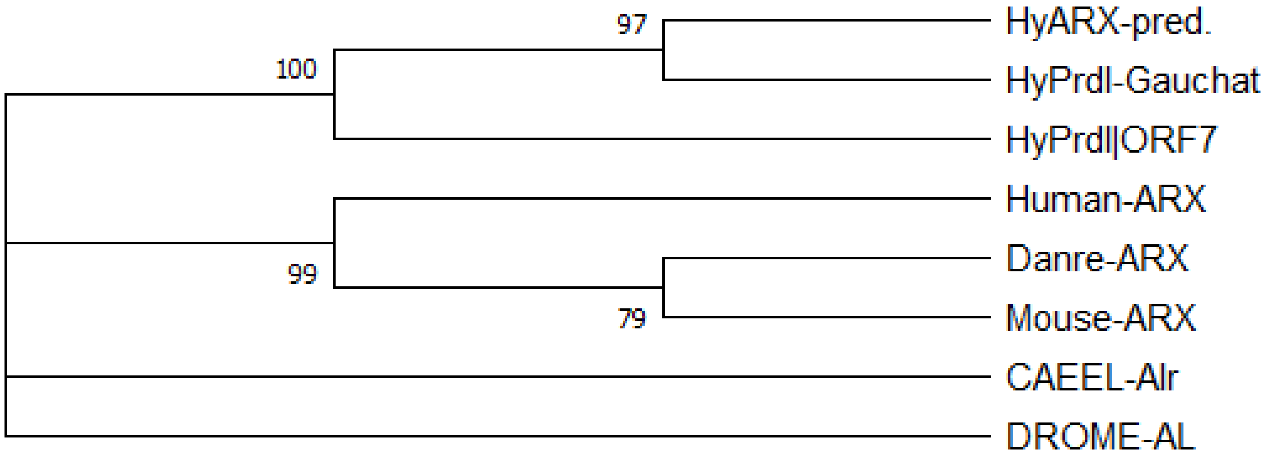


---

**11.) Ptx**

**t5275aep|PITX2_RAT**
TRINITY_DN14675_c0_g1 (**HyPtx|ORF8**)

XP_002164986.2 PREDICTED: pituitary homeobox 1-like [Hydra vulgaris] (**HyPtx1L-pred.**)

Alignment of **HyPtx|ORF8**, **HyPtx1L-pred.** and pituitary homeobox proteins from different animals:

```
HyPtx|ORF8       ------------------------------------------------------------    -
HyPtx1L-pred.    ------------------------------------------------------------    -
CaeelUnc30       ------------------------------------------------------------    -
DromePtx1-C      MDRSSAVGGCGGGGGLGVGVVTSSATALGPGGLTNGGGGVVSGALNGLEAMSAESTGLCL   60
Danre-Pitx1      ------------------------------------------------------------    -
Mouse-Pitx1      ------------------------------------------------------------    -
Human-Pitx1      ------------------------------------------------------------    -


HyPtx|ORF8       ------------------------------------------------------------    -
HyPtx1L-pred.    ------------------------------------------------------------    -
CaeelUnc30       -------------------------------------MDD---NTATLTAIHQQQQSHNR   20
DromePtx1-C      QDLVSAGTANGAGSAGSAESATTTSTALSSGSTGSSTVNGGGSSTSGTEHLHSHHSLHDS  120
Danre-Pitx1      ------------------------------------------------------------    -
Mouse-Pitx1      ------------------------------------------------------------    -
Human-Pitx1      ------------------------------------------------------------    -


HyPtx|ORF8       ------------------------------------------------------------    -
HyPtx1L-pred.    ------------------------------------------------------------    -
CaeelUnc30       F-----S------NPLVC------------VGQLDH----HSLLPEHSISSSLAPLTH--   51
DromePtx1-C      SSSVSISPAISSLMPISSLSHLHHSAGQDLVGGYSQHPH-HTVVPPHTPKHEPLEKLRIW  179
Danre-Pitx1      ------------------------------------------------MNVDMASSF---    9
Mouse-Pitx1      --------------------------MDAFKGGMSLERLPEGLRPPPPPPHDMGPSF---   31
Human-Pitx1      --------------------------MDAFKGGMSLERLPEGLRPPPPPPHDMGPAF---   31


HyPtx|ORF8       ------------------------------------------------------------    -
HyPtx1L-pred.    ------------------ML---TNDKPASSQASNVLKK-----------TESPVQARVK   28
CaeelUnc30       -NPYAFNY---------------SIPLPPTDITTKLPKLELLSLDVKQEQDDNHLDT---   92
DromePtx1-C      AETGDFRDSHSSMTAVANSLDSTHLNNFQTSSTSSISNRSRDRKDGNRSVNETTIKTENI  239
Danre-Pitx1      -----------------------HLPRSASSTRETLENTSSESSDTEAAEKERSVEQRS-   45
Mouse-Pitx1      -----------------------HLARAA-DPREPLENSASESSDADLPDKERGGEAKGP   67
Human-Pitx1      -----------------------HLARPA-DPREPLENSASESSDTELPEKERGGEPKGP   67
```

Homeobox domain

```
HyPtx|ORF8        ---------------------------------------------------MAMREDIAQ    9
HyPtx1L-pred.     QDVE--NKE---DTSGYTAGVDNRIRRQRTHFTVTQLHRLETCFARNRYPDMAMREDIAQ   83
CaeelUnc30        -----SSPTDSTGNGSTNGGKIQKPRRQRTHFTSHQLTELENWFSRNRYPDMACREEIAV  147
DromePtx1-C       SSSGHDEPMTTSGEEPKNDKKNKRQRRQRTHFTSQQLQELEHTFSRNRYPDMSTREEIAM  299
Danre-Pitx1       ------------DDGNADDPKKKKQRRQRTHFTSQQLQELEATFQRNRYPDMSTREEIAV   93
Mouse-Pitx1       EDGG--AGSAGCGGGAEDPAKKKKQRRQRTHFTSQQLQELEATFQRNRYPDMSMREEIAV  125
Human-Pitx1       EDSG--AGGTGCGG-ADDPAKKKKQRRQRTHFTSQQLQELEATFQRNRYPDMSMREEIAV  124
                                          rrqrthft  ql  le  f rnrypdM  RE IA

HyPtx|ORF8        WCSLTESRVRIWFKNRRAKWRKKERHLERPDICN--FAPTYRFESPSPYERRPTFPTTYS   67
HyPtx1L-pred.     WCSLTESRVRIWFKNRRAKWRKKERHLERPDICN--FAPTYRFESPSPYERRPTFPTTYS  141
CaeelUnc30        WISLTEPRVRVWFKNRRAKWRKRERNYVIDNGQGTTKVTAQSLD------PLGSLQNTFP  201
DromePtx1-C       WTNLTEARVRVWFKNRRAKWRKRERNAMNAAVAAADFKS------------------GFG  341
Danre-Pitx1       WTNLTEARVRVWFKNRRAKWRKRERNQQMDLCKN-SYLP------------------QF-  133
Mouse-Pitx1       WTNLTEPRVRVWFKNRRAKWRKRERNQQLDLCKG-GYVP------------------QF-  165
Human-Pitx1       WTNLTEPRVRVWFKNRRAKWRKRERNQQLDLCKG-GYVP------------------QF-  164
                  W  LTE RVR6WFKNRRAKWRK4ER                                5

HyPtx|ORF8        SNVSQPTLPTCSSLNNRSSHASLGHYYSHNTHWYPSDHPVHSLSQRNSPSNSPAGWYGIN  127
HyPtx1L-pred.     SNVSQPTLPTCSSLNNRSSHASLGHYYSHNTHWYPSDHPVHSLSQRNSPSNSPAGWYGIN  201
CaeelUnc30        QTLLQSSSSQ---LDD--SAVTSSSFYGYGGAWQQNPYY-SRNNQ------TTFNWQIKP  249
DromePtx1-C       TQFMQPFA------DD--S-LY--SSYPYN-NWTK---VPSPLGT------KPFPWPV--  378
Danre-Pitx1       SGLMQPYD-------D--V-YP---TYTYN-NWTNKGLTPAPLST------KNFTFFN--  171
Mouse-Pitx1       SGLVQPYE-------D--V-YA--AGYSYN-NWAAKSLAPAPLST------KSFTFFN--  204
Human-Pitx1       SGLVQPYE-------D--V-YA--AGYSYN-NWAAKSLAPAPLST------KSFTFFN--  203
                      Qp          1          Y  n  W          l            5
```

```
HyPtx|ORF8        Y-------------------------NEP-------------VRGYPTCPQSSSSTLSRE   149
HyPtx1L-pred.     Y-------------------------NEP-------------VRGYPTCPQSSSSTLSRE   223
CaeelUnc30        QFQTIPMSPTTAT----------SRFSTAANLAPLPTA----QAAFSTSATSSNDKLKLM   295
DromePtx1-C       -------NPLGSMVAGNHHQNSVNCFNTGASGVAVSM---NNASMLPGSMGSSLSNTS--   426
Danre-Pitx1       -----SMSPLTSQ----------SMFSAPSSISSMSMASGMGHSAVPGMPTTGLNNIGNL   216
Mouse-Pitx1       -----SMSPLSSQ----------SMFSAPSSISSMTMPSSMGPGAVPGMPNSGLNNINNL   249
Human-Pitx1       -----SMSPLSSQ----------SMFSAPSSISSMTMPSSMGPGAVPGMPNSGLNNINNL   248
                                                                p    3

HyPtx|ORF8        DEMMSPISF----SQRTHTFSPGDHSY--------------------------QHSI---   176
HyPtx1L-pred.     DEMMSPISF----SQRTHTFSPGDHSY--------------------------QHSI---   250
CaeelUnc30        DGLSNSLSSSLGQPYQPCQYSGPL------------------------------------   319
DromePtx1-C       ---------NVGAVGAPCPYTTPANPYMYRSAAEPCMSSSMSSSIATLRLKAKQHASAGF   477
Danre-Pitx1       NGIGGS-TINPAMSSSTCPYGPPGSPYSVYR-------DTCNSSLATLRLKSKQHPSFGY   268
Mouse-Pitx1       ---TGS-SLNSAMSPGACPYGTPASPYSVYR-------DTCNSSLASLRLKSKQHSSFGY   298
Human-Pitx1       ---TGS-SLNSAMSPGACPYGTPASPYSVYR-------DTCNSSLASLRLKSKQHSSFGY   297
                                     5      y                          qh

HyPtx|ORF8        ------------------------------------------------------------     -
HyPtx1L-pred.     ------------------------------------------------------------     -
CaeelUnc30        ------------------------------------------------------------     -
DromePtx1-C       GSPYSAPSPVSRSNSAGLSACQYTGVGVTDVVXENALGALLNQHQHHLQHFSGGSVGGSG   537
Danre-Pitx1       SGLQS--------PGSSLNACQYNS-----------------------------------   285
Mouse-Pitx1       GGLQG--------PASGLNACQYNS-----------------------------------   315
Human-Pitx1       GGLQG--------PASGLNACQYNS-----------------------------------   314

HyPtx|ORF8        ------------------------------------------------------------     -
HyPtx1L-pred.     ------------------------------------------------------------     -
CaeelUnc30        ------------------------------------------------------------     -
DromePtx1-C       VPNGMQQHSGNMLHHSNLGLDHSDLGLVGGGPSNHQDEGNHSLEGSNKSPMEAPLANSSN   597
Danre-Pitx1       ------------------------------------------------------------     -
Mouse-Pitx1       ------------------------------------------------------------     -
Human-Pitx1       ------------------------------------------------------------     -
```

```
HyPtx|ORF8      -------------    -
HyPtx1L-pred.   -------------    -
CaeelUnc30      -------------    -
DromePtx1-C     NNNNDNDDEDVID  610
Danre-Pitx1     -------------    -
Mouse-Pitx1     -------------    -
Human-Pitx1     -------------    -
```

Phylogenetic tree of pituitary homeobox proteins:

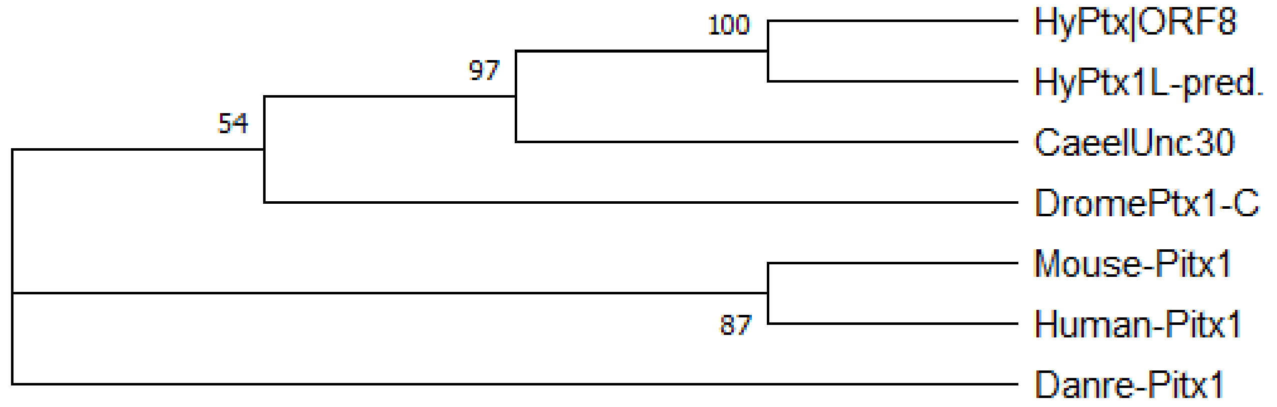


HyPtx is related to Pitx and Unc30. It has a conserved homeobox domain.

**12.) MAD**

**34122aep|MAD1_MOUSE**
TRINITY_DN1402_c1_g1 (**HyMAD1|ORF11**)

CDG70360.1 Hydra vulgaris Max dimerization protein 1 [Hydra vulgaris] (**HyMAD1-Wenger**)
Reference: Punctuated emergences of genetic and phenotypic innovations in eumetazoan, bilaterian, euteleostome, and hominidae ancestors. (Wenger and Galliot, 2013)

XP_002153919.1 PREDICTED: max dimerization protein 1-like [Hydra vulgaris] (**HyMAD1-Pred.**)

Alignment of **HyMAD1|ORF11**, **HyMAD1-Wenger**, **HyMAD1-Pred.** and Max dimerization proteins from other animals:

```
DROME-Mnt-g     MSGIGGIGVLLEAAQFLEQHEKLPQAGGGGGSALGVGVALPQQQQQTVHCKSNSIINNNN  60
CAEEL-MADL1     ------------------------------------------------------------  -
HyMAD1|ORF11    ------------------------------------------------------------  -
HyMAD1-Wenger   ------------------------------------------------------------  -
HyMAD1-Pred.    ------------------------------------------------------------  -
DANRE-MAD1      ------------------------------------------------------------  -
Mouse-MAD1      ------------------------------------------------------------  -
Human-MAD1      ------------------------------------------------------------  -


DROME-Mnt-g     SSQHNRNSSSSSNGIIATPPFTNGNGNGTVGVGNASAIFNVSVNANGNGLSNGHSSPRSQ  120
CAEEL-MADL1     ------------------------------------------------------------  -
HyMAD1|ORF11    ------------------------------------------------------------  -
HyMAD1-Wenger   ------------------------------------------------------------  -
HyMAD1-Pred.    ------------------------------------------------------------  -
DANRE-MAD1      ------------------------------------------------------------  -
Mouse-MAD1      ------------------------------------------------------------  -
Human-MAD1      ------------------------------------------------------------  -
```

```
DROME-Mnt-g     LAAAATAHDYEYSAAGVANGHSEGRPHVSARDQQDHPQHQQHANHMQIQQHYGYMRSNIG  180
CAEEL-MADL1     ------------------------------MEQQLNLGHLLTAARLLDIGALDISSLDLG   30
HyMAD1|ORF11    ----------------------------------MSLETLLEAAKVVEQ-----------   15
HyMAD1-Wenger   ----------------------------------MSLETLLEAAKVVEQ-----------   15
HyMAD1-Pred.    ----------------------------------MSLETLLEAAKVVEQ-----------   15
DANRE-MAD1      ------------------------------MTAIEKLQMLIEAAEYLDR-----------   19
Mouse-MAD1      ----------------------------MATAVGMNIQLLLEAADYLER-----------   21
Human-MAD1      ----------------------------MAAAVRMNIQMLLEAADYLER-----------   21
                                                         l eAa
```

```
DROME-Mnt-g     G-------------VIYGLHKDSADSAASSACGAGAGAAAVGMSIASSA-PAAGSQLSAH  226
CAEEL-MADL1     ALTSTSSSPGSSSPAMFDLSNES--ELRSLFCGKLK----VDKKQSSCASNASTSSQPYC   84
HyMAD1|ORF11    -------------------------------------------R-ENAGKY------TIG   25
HyMAD1-Wenger   -------------------------------------------R-ENAGKY------TIG   25
HyMAD1-Pred.    -------------------------------------------R-ENAGKY------TIG   25
DANRE-MAD1      -------------------------------------------R-EREAEHGYASMLPFT   35
Mouse-MAD1      -------------------------------------------R-EREAEHGYASMLPYS   37
Human-MAD1      -------------------------------------------R-EREAEHGYASMLPYN   37
                                                             e
```

Myc-type, basic helix-loop-helix (bHLH) domain

```
DROME-Mnt-g     HHTGHGGRRRTTSSNSNGAGTREVHNKLEKERRAQLKECYDLLKKVLPMGDEDRKKTSNL  286
CAEEL-MADL1     ---------SSPPARKSSKHSRTAHNELEKTRRANLRGCLETLKMLVPCVSDATRN-TTL  134
HyMAD1|ORF11    NRKD-------RKFAKRSQTYRATHNQLEKNRRAHLRDCLVSLRDLVPNSPDTSKV-TTL   77
HyMAD1-Wenger   NRKD-------RKFAKRSQTYRATHNQLEKNRRAHLRDCLVSLRDLVPNSPDTSKV-TTL   77
HyMAD1-Pred.    NRKD-------RKFAKRSQTYRATHNQLEKNRRAHLRDCLVSLRDLVPNSPDTSKV-TTL   77
DANRE-MAD1      SNKERDGLKRKIKSK-KNQSSRSTHNEMEKNRRAHLRLCLERLKSLVPLGPESNRH-TTL   93
Mouse-MAD1      --KDRDAFKRRNKPKKNSTSSRSTHNEMEKNRRAHLRLCLEKLKGLVPLGPESSRH-TTL   94
Human-MAD1      N-KDRDALKRRNKSKKNNSSSRSTHNEMEKNRRAHLRLCLEKLKGLVPLGPESSRH-TTL   95
                  k         k  k      R tHN 6EKnRRAhL4 Cl  L4 66P  p   4  3tL
```

```
DROME-Mnt-g        TILDTAHKYVNSLSHEVCEQEAKIEKLAKQKIELQKRLKQLSLRRESPPTDQQIIPQADK  346
CAEEL-MADL1        ALLTRARDHIIELQDSNAAQMKKLNDLRDEQDELVAELAQLQADEEVAQATSQ-------  187
HyMAD1|ORF11       SLLQSAKQYIKVLENHDRESQSIKRTLCLEQQHLRKKLAALMNG-FDISSDS--------  128
HyMAD1-Wenger      SLLQSAKQYIKVLENHDRESQSIKRTLCLEQQHLRKKLAALMNG-FDISSDS--------  128
HyMAD1-Pred.       SLLQSAKQYIKVLENHDRESQSIKRTLCLEQQHLRKKLAALMNG-FDISSDS--------  128
DANRE-MAD1         SLLMRAKEHIKRLEDSERKAQHTIDQLQREQRHLRRRLEQLGVERTRMDSMG--------  145
Mouse-MAD1         SLLTKAKLHIKKLEDCDRKAVHQIDQLQREQRHLKRRLEKLGAERTRMDSVG--------  146
Human-MAD1         SLLTKAKLHIKKLEDCDRKAVHQIDQLQREQRHLKRQLEKLGIERIRMDSIG--------  147
                    6L  A   6k L    r        L  2q hL   L  L

DROME-Mnt-g        AVCLATTATTAPPTAASGHNGTTILF------DAGNAC-ATGQLTTVINKTNGGATAAAS  399
CAEEL-MADL1        -ACQTLSQSR-P---ESRASSFTSTS------SRDSPCY----L-----------EYSPS  221
HyMAD1|ORF11       ---SILSPET-S---ISSIEEELIDVENDD-----ETGYGSADD-----------RNSTS  165
HyMAD1-Wenger      ---SILSPET-S---ISSIEEELIDVENDD-----ETGYGSADD-----------RNSTS  165
HyMAD1-Pred.       ---SILSPET-S---ISSIEEELIDVENDD-----ETGYGSADD-----------RNSTS  165
DANRE-MAD1         ---STISSDK-S---DSDQEEVDV--------DVEGTDYLLGDL-----------EWSTS  179
Mouse-MAD1         ---SVVSSER-S---DSDREELDVDVDVDVDVDVEGTDYLNGDL-----------GWSS-  187
Human-MAD1         ---STVSSER-S---DSDREEIDV--------DVESTDYLTGDL-----------DWSSS  181
                      s  3    s    S  ee             t y   d            s s

DROME-Mnt-g        SKHNGNHTGGATTTLTP-AMGIMTIKNGNGSNGNMLAISPTGGHNHQLHQHHNGSPAGSP  458
CAEEL-MADL1        SKPMDSHKPTIID-----LYAEGLIPRGPITFPRPLVY------PHNVFDL---------  261
HyMAD1|ORF11       SSS-----SSLNSVL---------------------------------------------  175
HyMAD1-Wenger      SSS-----SSLNSVL---------------------------------------------  175
HyMAD1-Pred.       SSS-----SSLNSVL---------------------------------------------  175
DANRE-MAD1         SVSDSDERGSIRSSCSDEGYSSASLRRLANSQENSLVL------ACS-------------  220
Mouse-MAD1         SVSDSDERGSMQSLGSDEGYSSATVKRAKLQDGHKAGL------SL--------------  227
Human-MAD1         SVSDSDERGSMQSLGSDEGYSSTSIKRIKLQDSHKACL------GL--------------  221
                   S s
```

```
DROME-Mnt-g     SGIGATTGATITRGSPTPSTPSPSPSSSSSGVSLSTSFMGGSSSSSSSSSSSSSSSSSSS  518
CAEEL-MADL1     -----------MNLPPTP-----FDVSQFLPINLQV------------------------  281
HyMAD1|ORF11    ------------------------------------------------------------    -
HyMAD1-Wenger   ------------------------------------------------------------    -
HyMAD1-Pred.    ------------------------------------------------------------    -
DANRE-MAD1      ---------------------------------L--------------------------  221
Mouse-MAD1      ------------------------------------------------------------    -
Human-MAD1      ------------------------------------------------------------    -

DROME-Mnt-g     SSSPTQQQQIITPTATARSVGLKFHGGAGAAVPAGGGSVNGFHQADKDRNYTFLRLSAGT  578
CAEEL-MADL1     ------------------------------------------------------------    -
HyMAD1|ORF11    ------------------------------------------------------------    -
HyMAD1-Wenger   ------------------------------------------------------------    -
HyMAD1-Pred.    ------------------------------------------------------------    -
DANRE-MAD1      ------------------------------------------------------------    -
Mouse-MAD1      ------------------------------------------------------------    -
Human-MAD1      ------------------------------------------------------------    -

DROME-Mnt-g     TAQ  581
CAEEL-MADL1     ---    -
HyMAD1|ORF11    ---    -
HyMAD1-Wenger   ---    -
HyMAD1-Pred.    ---    -
DANRE-MAD1      ---    -
Mouse-MAD1      ---    -
Human-MAD1      ---    -
```

Myc-type, basic helix-loop-helix (bHLH) domain; HyMAD1|ORF11 is identical with HyMAD-Wenger(Wenger and Galliot, 2013).

13.) **Prickle-like Protein**

**t19041aep|ESN_DROPS**
TRINITY_DN3315_c0_g1 (**HyPrickle|ORF8**)

CDG71924.1 Hydra vulgaris Prickle-like protein 3 [Hydra vulgaris] (**HyPrickle3-Wenger**)
Reference: Punctuated emergences of genetic and phenotypic innovations in eumetazoan, bilaterian, euteleostome, and hominidae ancestors.(Wenger and Galliot, 2013)

XP_012566699.1 PREDICTED: protein prickle-like isoform X1 [Hydra vulgaris] (**HyPrickle-Pred.**)

Alignment of **HyPrickle|ORF8, HyPrickle3-Wenger, HyPrickle-Pred.** and other LIM domain containing proteins:

```
HyPrickle3-Wenger   ------------------------------------------------------------    -
Human-Prickle2      ------------------------------------------------------------    -
DANRE-Prickle3-x1   ------------------------------------------------------------    -
HyPrickle|ORF8      MSTFQPLLHETFNSILPKNTINGNNLDILDEVKNGGFDRKKELNKMSPNEQ--FSKDLLH   58
HyPrickle-Pred.     MSTFQPLLHETFNSILPKNTINGNNLDILDEVKNGGFDRKKELNKMSPNEQ--FSKDLLH   58
CAEEL-LIM-9         -------MRA---------MVRGMVGRGKKKEKNGGVPPSVPSSASQPPPSTTKHAKILP   44
DROME-Limpet        --------------------------------------------------------MLLK    4


HyPrickle3-Wenger   -------MHRSK-----KDKYGVNSKDDLS----NMSLRDHMIHCISCGRSCPGYEPTNW   44
Human-Prickle2      ------------------------------------------------------------    -
DANRE-Prickle3-x1   ------MFTRGS-----KKRR----SNRST----EAEDPDRGQPCMRCGEQCPGFRMHGW   41
HyPrickle|ORF8      -----------NIEQNSINKNGANK--RTPP---PKPPRISRNSSVLSNISSGSTSHDSL  102
HyPrickle-Pred.     -----------NIEQNSINKNGANK--RTPP---PKPPRISRNSSVLSNISSGSTSHDSL  102
CAEEL-LIM-9         SFSCSPPMERKDSRPTLITANTADHSVGSPPARRQ-PPKRVGSGKSASNGPSGVVRTDLA  103
DROME-Limpet        VCPCD----DFQVQPKVVKKQP-----KFPTRERHMAIRRKKSTAMVTTTSENTAKCPEQ   55
```

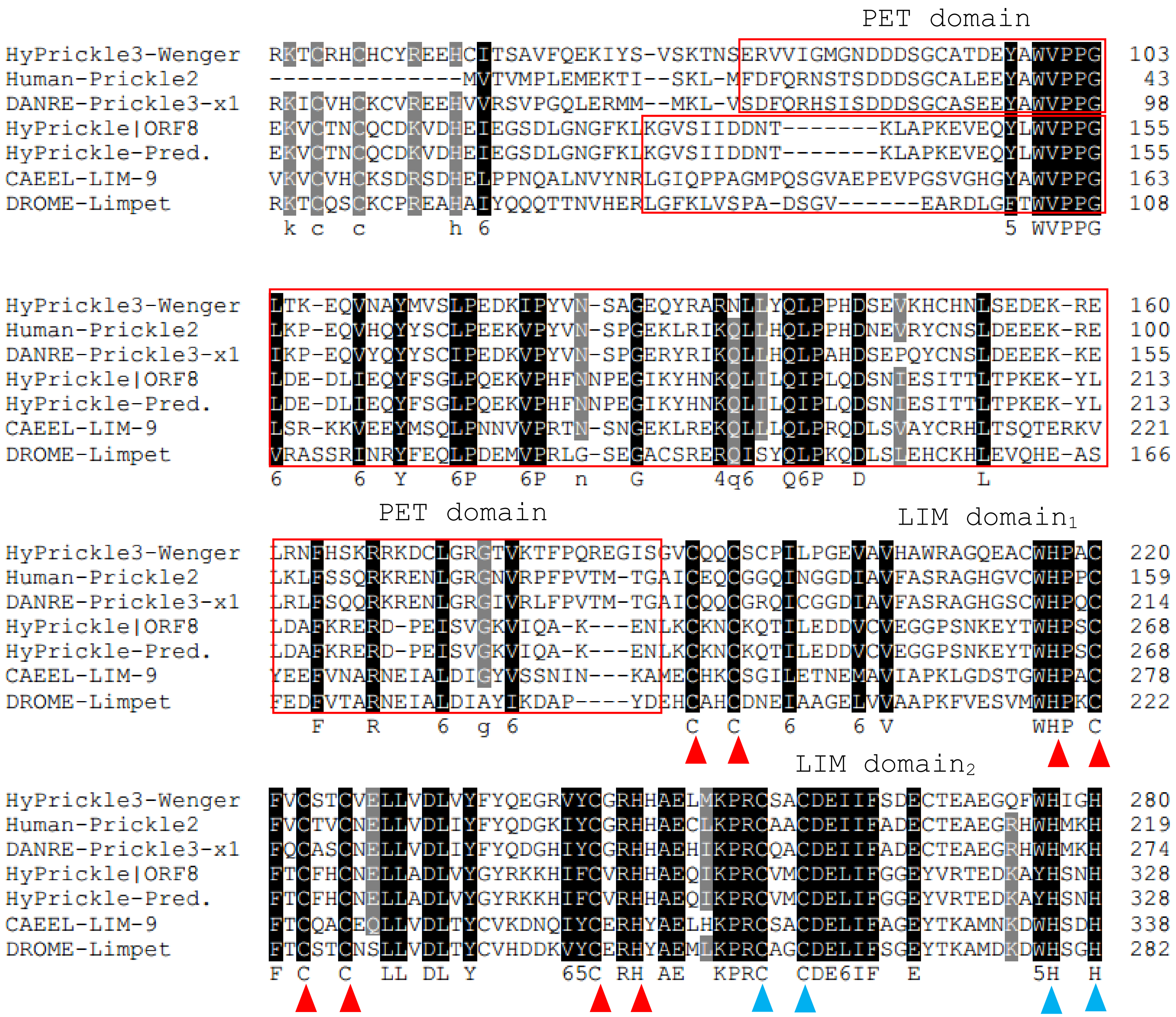
PET domain
HyPrickle3-Wenger RKTCRHCHCYREEHCITSAVFQEKIYS-VSKTNSERVVIGMGNDDDSGCATDEYAWVPPG 103
Human-Prickle2 ---------------MVTVMPLEMEKTI--SKL-MFDFQRNSTSDDDSGCALEEYAWVPPG 43
DANRE-Prickle3-x1 RKICVHCKCVREEHVVRSVPGQLERMM--MKL-VSDFQRHSISDDDSGCASEEYAWVPPG 98
HyPrickle|ORF8 EKVCTNCQCDKVDHEIEGSDLGNGFKLKGVSIIDDNT-------KLAPKEVEQYLWVPPG 155
HyPrickle-Pred. EKVCTNCQCDKVDHEIEGSDLGNGFKLKGVSIIDDNT-------KLAPKEVEQYLWVPPG 155
CAEEL-LIM-9 VKVCVHCKSDRSDHELPPNQALNVYNRLGIQPPAGMPQSGVAEPEVPGSVGHGYAWVPPG 163
DROME-Limpet RKTCQSCKCPREAHAIYQQQTTNVHERLGFKLVSPA-DSGV------EARDLGFTWVPPG 108
k c c h 6 5 WVPPG
HyPrickle3-Wenger LTK-EQVNAYMVSLPEDKIPYVN-SAGEQYRARNLLYQLPPHDSEVKHCHNLSEDEK-RE 160
Human-Prickle2 LKP-EQVHQYYSCLPEEKVPYVN-SPGEKLRIKQLLHQLPPHDNEVRYCNSLDEEEK-RE 100
DANRE-Prickle3-x1 IKP-EQVYQYYSCIPEDKVPYVN-SPGERYRIKQLLHQLPAHDSEPQYCNSLDEEEK-KE 155
HyPrickle|ORF8 LDE-DLIEQYFSGLPQEKVPHFNNPEGIKYHNKQLILQIPLQDSNIESITTLTPKEK-YL 213
HyPrickle-Pred. LDE-DLIEQYFSGLPQEKVPHFNNPEGIKYHNKQLILQIPLQDSNIESITTLTPKEK-YL 213
CAEEL-LIM-9 LSR-KKVEEYMSQLPNNVVPRTN-SNGEKLREKQLLLQLPRQDLSVAYCRHLTSQTERKV 221
DROME-Limpet VRASSRINRYFEQLPDEMVPRLG-SEGACSREREQISYQLPKQDLSLEHCKHLEVQHE-AS 166
6 6 Y 6P 6P n G 4q6 Q6P D L
PET domain
LIM domain1
HyPrickle3-Wenger LRNFHSKRRKDCLGRGTVKTFPQREGISGVCQQCSCPILPGEVAVHAWRAGQEACWHPAC 220
Human-Prickle2 LKLFSSQRKRENLGRGNVRPFPVTM-TGAICEQCGGQINGGDIAVFASRAGHGVCWHPPC 159
DANRE-Prickle3-x1 LRLFSQQRKRENLGRGIVRLFPVTM-TGAICQQCGRQICGGDIAVFASRAGHGSCWHPQC 214
HyPrickle|ORF8 LDAFKRERD-PEISVGKVIQA-K---ENLKCKNCKQTILEDDVCVEGGPSNKEYTWHPSC 268
HyPrickle-Pred. LDAFKRERD-PEISVGKVIQA-K---ENLKCKNCKQTILEDDVCVEGGPSNKEYTWHPSC 268
CAEEL-LIM-9 YEEFVNARNEIALDIGYVSSNIN---KAMECHKCSGILETNEMAVIAPKLGDSTGWHPAC 278
DROME-Limpet FEDFVTARNEIALDIAYIKDAP----YDEHCAHCDNEIAAGELVVAAPKFVESVMWHPKC 222
F R 6 g 6 C C 6 6 V WHP C
LIM domain2
HyPrickle3-Wenger FVCSTCVELLVDLVYFYQEGRVYCGRHHAELMKPRCSACDEIIFSDECTEAEGQFWHIGH 280
Human-Prickle2 FVCTVCNELLVDLIYFYQDGKIYCGRHHAECLKPRCAACDEIIFADECTEAEGRHWHMKH 219
DANRE-Prickle3-x1 FQCASCNELLVDLIYFYQDGHIYCGRHHAEHIKPRCQACDEIIFADECTEAEGRHWHMKH 274
HyPrickle|ORF8 FTCFHCNELLADLVYGYRKKHIFCVRHHAEQIKPRCVMCDELIFGGEYVRTEDKAYHSNH 328
HyPrickle-Pred. FTCFHCNELLADLVYGYRKKHIFCVRHHAEQIKPRCVMCDELIFGGEYVRTEDKAYHSNH 328
CAEEL-LIM-9 FTCQACEQLLVDLTYCVKDNQIYCERHYAELHKPRCSACDELIFAGEYTKAMNKDWHSDH 338
DROME-Limpet FTCSTCNSLLVDLTYCVHDDKVYCERHYAEMLKPRCAGCDELIFSGEYTKAMDKDWHSGH 282
F C C LL DL Y 65C RH AE KPRC CDE6IF E 5H H

```
                                                                           LIM domain3
HyPrickle3-Wenger   FSCFECDGSLGGQRYVMRDNRPICCKCFEKMFAEFCDACGEPIGIDVGQMAHGSQHWHAN   340
Human-Prickle2      FCCFECETVLGGQRYIMKEGRPYCCHCFESLYAEYCDTCAQHIGIDQGQMTYDGQHWHAT   279
DANRE-Prickle3-x1   FCCFECEAALGGQRYIMRESRPYCCRCYESLYAEYCDTCGEHIGIDQGQMTYEGQHWHAS   334
HyPrickle|ORF8      FICSYCERGLTGEQHLVDQGLPICIMCYDEKYASICHMCGKTIGVDEEDVIYDDEHWH--   386
HyPrickle-Pred.     FICSYCERGLTGEQHLVDQGLPICIMCYDEKYASICHMCGKTIGVDEEDVIYDDEHWH--   386
CAEEL-LIM-9         FCCWQCDQTLTGQRYIMRDEQPYCIKCYEDVFANQCDECAKPIGIDSKDLSYKDKHWH--   396
DROME-Limpet        FCCWQCDESLTGQRYVIRDDHPYCIKCYENVFANTCEECNKIIGIDSKDLSYKDKHWH--   340
                    F C  C    L G2  66      P C  C5    5A  C  C   IG6D   6 y   HWH

HyPrickle3-Wenger   EKCFSCFTCGLSLLGQPFLPKNGEIFCSSSCSKGTLPPNPTSTKYPPRSASKNRSL----   396
Human-Prickle2      ETCFCCAHCKKSLLGRPFLPKQGQIFCSRACSAGEDPNGSDSSDSAFQNARAKESRRSAK   339
DANRE-Prickle3-x1   EQCFCCACCRLPLLGRPFLPRGGLIFCSRSCSLGEDPENSDSCDSALQSKSATHKPLQSQ   394
HyPrickle|ORF8      DACLVCTHCHCKLSGTSFVVKDDNFLCSECYQK-TDDKRCKRCMKGFEPGVKRLELKGDF   445
HyPrickle-Pred.     DACLVCTHCHCKLSGTSFVVKDDNFLCSECYQK-TDDKRCKRCMKGFEPGVKRLELKGDF   445
CAEEL-LIM-9         EHCFLCSMCKISLVDMPFGSKNDRIFCSNCYDQ-AFATRCDGCNEIFRAGMKKMEYKGKQ   455
DROME-Limpet        EACFLCFKCHLSLVDKQFGAKADKIYCGNCYDA-QFASRCDGCGEVFRAGTKKMEYKTRQ   399
                      C  C  C    L    F  4     Cs                 LIM domain4

HyPrickle3-Wenger   ---------KPPDSTNSTYIMSSSMSP----EPVRKLIEFRSKTSYRSSLDKYGLAAAEK   443
Human-Prickle2      IGKNKGKTEEPMLNQHSQLQVSSNRLS-ADVDPLSLQMDMLSLSSQTPSLN---------   389
DANRE-Prickle3-x1   QR---SSTPRPQG-GSSD--ISIQRTPAANGGQITLQTKGVH-TSFPPLVN---------   438
HyPrickle|ORF8      WHENC------FVCDSCKKPITSKRFIHHEGKQV-------C----CPC---FDLYFAKR   485
HyPrickle-Pred.     WHENC------FVCDSCKKPITSKRFIHHEGKQV-------C----CPC---FDLYFAKR   485
CAEEL-LIM-9         WHDKC------FCCAHCKLAIGTKSFIPKNDDVF-------C----GPC---YEEKFATR   495
DROME-Limpet        WHENC------FCCCVCKTAIGTKSFIPREQEIY-------C----AGC---YEEKFATR   439
                                        6

HyPrickle3-Wenger   IGDMIRNIAKEGYI--DDSDREDTKSN---------------------------------   468
Human-Prickle2      -RDPIWRSREEPYHYGNKMEQNQTQSPLQLLSQCNIRTSYSPGG-QGAGAQPEMWGKHFS   447
DANRE-Prickle3-x1   -GD--------PPPYGRTS----TPHPLPSAQLCPVANDRAGSLSKDQSETPSADG-PYS   484
HyPrickle|ORF8      CGKCTEVLREGGVACGGNFYHRD----CFICDNCSDPIS--------SQPFQQKDGKRFC   533
HyPrickle-Pred.     CGKCTEVLREGGVACGGNFYHRD----CFICDNCSDPIS--------SQPFQQKDGKRFC   533
CAEEL-LIM-9         CSKCKKVITAGGVTYKNEPWHRE----CFCCTNCNSSLA--------GQRFTSKDEKPYC   543
DROME-Limpet        CIKCNKVITSGGVTYKNEPWHRE----CFTCTHCNITLA--------GQRFTSRDEKPYC   487
                                                      c
                              LIM domain5
```

```
HyPrickle3-Wenger   --SSKVSISSKKELPCFPPSNKHSEKPVLRVPPRHRSKLSTEVWIDMVP-----PKESIT  521
Human-Prickle2      NPKRSSSLAMTGHAGSFIKECREDYYP-----GRLR---SQESYSDMSS-----QSFSET  494
DANRE-Prickle3-x1   ALEFTVELNGYAEFGPFPPSCTS-------------------------------------  507
HyPrickle|ORF8      TPCYKS---------LFAKKCTACGDYII---NGEFYTVDADNWHKNCFRCVTCNEILYR  581
HyPrickle-Pred.     TPCYKS---------LFAKKCTACGDYII---NGEFYTVDADNWHKNCFRCVTCNEILYR  581
CAEEL-LIM-9         ANCYGD---------LFAKRCNACTKPITGIGGAKFISFEDRHWHNDCFICAQCTTSLVG  594
DROME-Limpet        AECFGE---------LFAKRCTACVKPITGIGGTRFISFEDRHWHHDCFVCASCKASLVG  538
                                    F   C
```

LIM domain$_6$

```
HyPrickle3-Wenger   RHEVVQRNQLKINERNKYGNITHINGNASIEKLVQAERQRRRRRKRDDYYSDFEVDKRKQ  581
Human-Prickle2      RGSIQ-VPKYE-EEEEEEGGLSTQQ--------------CRT---RH-PIS---SL----  527
DANRE-Prickle3-x1   RGTST-VK--------DCGNIIESS--------------CSG---ISSQLASFDDL----  537
HyPrickle|ORF8      QSFAQENGVIKLI-----------CENCQDQ-----------------------------  601
HyPrickle-Pred.     QSFAQENGVIKLI-----------CENCQDQ-----------------------------  601
CAEEL-LIM-9         KGFITDGH--EIL-----------CPECAKARLMAANS----------------------  619
DROME-Limpet        RGFITDGP--DIL-----------CPDCAKQKLM--------------------------  559
```

```
HyPrickle3-Wenger   SERRENFSTPQNSSQIAVKKLVDNDFTKTKSTESLDTTK--SRI--VGRNNERENNNKR-  636
Human-Prickle2      -KYTEDMT----P----------TEQTPRGSMESLAL-SNATGLSADGGAKRQEHLSRFS  571
DANRE-Prickle3-x1   -NDSDSFPPPPPP----------VFFDPSDSPDPLASVTEAEDLPAA-------------  573
HyPrickle|ORF8      ------------------------------------------------------------    -
HyPrickle-Pred.     ------------------------------------------------------------    -
CAEEL-LIM-9         ------------------------------------------------------------    -
DROME-Limpet        ------------------------------------------------------------    -
```

```
HyPrickle3-Wenger   ---------------TKSDASLNNTIKYKSK---------------NDAVANLKLGIEVS  666
Human-Prickle2      MPDLSKDSGMNVSEKLSNMGTLNSSMQFRSAESVRSLLSAQQYQEMEGNLHQLSNPIGYR  631
DANRE-Prickle3-x1   -P----------------VRSSTARVSFREPISCSYSVEENEWEDE-------DQP----  605
HyPrickle|ORF8      ------------------------------------------------------------    -
HyPrickle-Pred.     ------------------------------------------------------------    -
CAEEL-LIM-9         ------------------------------------------------------------    -
DROME-Limpet        ------------------------------------------------------------    -
```

```
HyPrickle3-Wenger   TPKRNYQ-----MIDGRRVAGKSIENVENIK-------PRSFGMLSA-------------  701
Human-Prickle2      DLQSHGRMHQSFDFDG-GMAGSKLPGQEGVRIQPMSERTR-------------RR--ATS  675
DANRE-Prickle3-x1   EMQRRDRDEDEQEED-----GEEAEGSLGHRLRICSQVPSRMDLLDGSYHCNFRRGSGGH  660
HyPrickle|ORF8      ------------------------------------------------------------    -
HyPrickle-Pred.     ------------------------------------------------------------    -
CAEEL-LIM-9         ------------------------------------------------------------    -
DROME-Limpet        ------------------------------------------------------------    -


HyPrickle3-Wenger   -EDQKRTRIN-----YVTKDDMAIH------------DRNILKKA---------------  728
Human-Prickle2      RDDNRRFRPHRSRRSRRSRSDNALHLASEREAISRLKDRPPLRAREDYDQFMRQRSFQES  735
DANRE-Prickle3-x1   HSSEKRYRR-----SRGSRSD-----------------RPRLETL----DVERRRSSN--  692
HyPrickle|ORF8      ------------------------------------------------------------    -
HyPrickle-Pred.     ------------------------------------------------------------    -
CAEEL-LIM-9         ------------------------------------------------------------    -
DROME-Limpet        ------------------------------------------------------------    -


HyPrickle3-Wenger   -KKKDKTKNQGQ------------------------------------------------  739
Human-Prickle2      MGHGSRRDLYGQCPRTVSDLALQNAFGDRWGPYFAEYDWCSTCSSSSESDNEGYFLGEPI  795
DANRE-Prickle3-x1   NADGSRN-----------------TSLTLYSSRSRHEDSCSTCSSSSDSEEEGFFLGQRI  735
HyPrickle|ORF8      ------------------------------------------------------------    -
HyPrickle-Pred.     ------------------------------------------------------------    -
CAEEL-LIM-9         ------------------------------------------------------------    -
DROME-Limpet        ------------------------------------------------------------    -


HyPrickle3-Wenger   ----------------------------------------------------CVMS  743
Human-Prickle2      PQPARLRYVTSDELLHKYSSYGLPKSSTLGGRGQLH-------SRKRQKSKNCIIS  844
DANRE-Prickle3-x1   PLPPQLTGQRADEGQTE--------DEEKSQRGSFRRRRRTQSLGQKDKDKNCILS  783
HyPrickle|ORF8      --------------------------------------------------------    -
HyPrickle-Pred.     --------------------------------------------------------    -
CAEEL-LIM-9         --------------------------------------------------------    -
DROME-Limpet        --------------------------------------------------------    -
```

Phylogenetic tree of LIM-domain containing proteins:

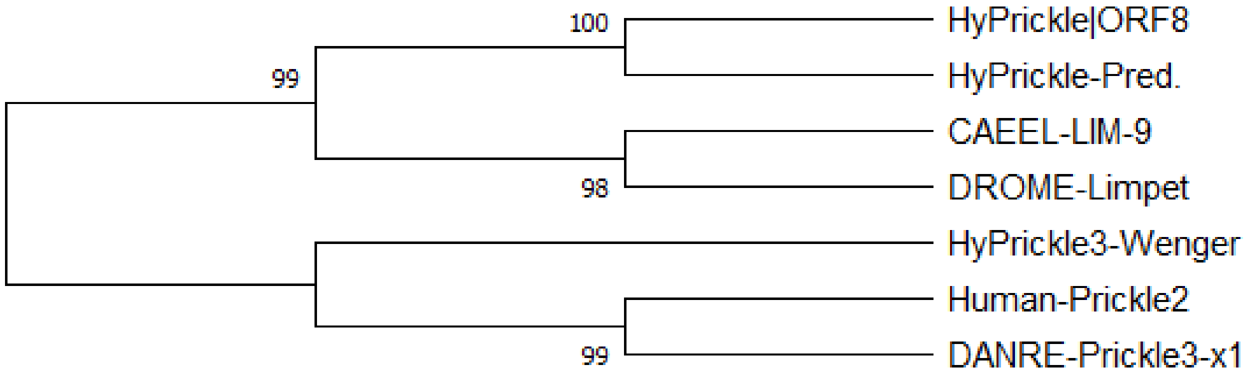


LIM domain proteins include testin, prickle, dyxin and LIMPETin. Structurally, testin and prickle proteins usually contain three LIM domains at C-terminal; LIMPETin has six LIM domains. However, all members of the family contain a PET protein-protein interaction domain. LIM domains share two highly conserved zinc finger motifs which contain eight conserved residues, mostly cysteines (C) and histidines (H), which coordinately bond to two zinc atoms.

Here, HyPrickle|ORF8 has six LIM domains, showing a closer relationship with Limpet proteins. However, HyPrickle3-Wenger has three LIM domains. But they both have a PET domain at the N-terminal.

---

**14.) PAX Proteins**
**Not NR-genes**

**t9974aep|PAX2A_DANRE;**
**t11467aep|PAX6_XENLA;**
**t6559aep|PAX5_HUMAN;**

Alignment of **t9974aep**, **t11467aep**, **t6559aep** and PAX Proteins from different animals:

```
t11467aep-ORF1    ------------------------------------------------------------    -
t9974aep-ORF4     ------------------------------------------------------------    -
XENLA--PAX6       ------------------------------------------------------------    -
Human-PAX6-h      ------------------------------------------------------------    -
t6559aep-ORF1     MLEQWSDDNKMEIFKPQQSGELTCNVDKRSINEDGVGQVICLENPHLLKTLQDHSLAAAT   60
Human-PAX5-s5     ------------------------------------------------------------    -
Human-PAX5-s7     ------------------------------------------------------------    -
Danre--PAX2A      ------------------------------------------------------------    -
Mouse-PAX2-s10    ------------------------------------------------------------    -
```

```
                                            Paired DNA-binding domain  (PS51057)
t11467aep-ORF1    ------------------------------------------------------------    -
t9974aep-ORF4     ------------MPFQDTKFSVNDPGGVNQLGGVFVNGRPLPDYMRHRIIELAQCGVRPS   48
XENLA--PAX6       --------------------MQNSHSGVNQLGGVFVNGRPLPDSTRQKIVELAHSGARPC   40
Human-PAX6-h      --------------------MQNSHSGVNQLGGVFVNGRPLPDSTRQKIVELAHSGARPC   40
t6559aep-ORF1     VSNLNINIKRFNDAKENKNTCRDNHGGINQLGGTFVNGRPLIEPVRRKIVELAHQGVRPC  120
Human-PAX5-s5     -----MDL---EKNYPTPRTSRTGHGGVNQLGGVFVNGRPLPDVVRQRIVELAHQGVRPC   52
Human-PAX5-s7     -----MDL---EKNYPTPRTSRTGHGGVNQLGGVFVNGRPLPDVVRQRIVELAHQGVRPC   52
Danre--PAX2A      -----MDIHCKADPFSAMHLSHIGHGGVNQLGGVFVNGRPLPDVVRQRIVELAHQGVRPC   55
Mouse-PAX2-s10    -----MDMHCKADPFSAMH---PGHGGVNQLGGVFVNGRPLPDVVRQRIVELAHQGVRPC   52
                                          h g nqlggvfvngrplpd  r  i elah g rpc
```

```
t11467aep-ORF1   ----MKKIKNHCV---------------SRVGGSKPKVATPEVVNKIEALKREKHDMFAW   41
t9974aep-ORF4    EISRQLLVSHGCVSKILGRYYETGSVRPGAIGGSKPKVATPKVVCRIVKLKEENPCMFAW  108
XENLA--PAX6      DISRILQVSNGCVSKILGRYYETGSIRPRAIGGSKPRVATPEVVNKIAHYKRECPSIFAW  100
Human-PAX6-h     DISRILQVSNGCVSKILGRYYETGSIRPRAIGGSKPRVATPEVVSKIAQYKRECPSIFAW  100
t6559aep-ORF1    DISRQLRVSHGCVSKILSRFYETGSVRPGVIGGSKPKVATPSVVAKIQEYKQHNPTMFAW  180
Human-PAX5-s5    DISRQLRVSHGCVSKILGRYYETGSIKPGVIGGSKPKVATPKVVEKIAEYKRQNPTMFAW  112
Human-PAX5-s7    DISRQLRVSHGCVSKILGRYYETGSIKPGVIGGSKPKVATPKVVEKIAEYKRQNPTMFAW  112
Danre--PAX2A     DISRQLRVSHGCVSKILGRYYETGSIKPGVIGGSKPKVATPKVVEKIAEYKRQNPTMFAW  115
Mouse-PAX2-s10   DISRQLRVSHGCVSKILGRYYETGSIKPGVIGGSKPKVATPKVVDKIAEYKRQNPTMFAW  112
                 disr l 6s gCVskilgr yetgs  p 6GGSKP4VATP VV 4I  yKr  p 6FAW

t11467aep-ORF1   EIREKLIDSKLCPPSQCPSISSINRILRKRAAERAAERAATDNT-------FHKGKPNLS   94
t9974aep-ORF4    EIRNSLLAEGICDNGNVPSVSSINRILRNHAAEKETKEAKYKQEELSKNNSFGMMNGFT-  167
XENLA--PAX6      EIRDRLLSEGVCTNDNIPSVSSINRVLRNLASDKQQMGSEGMYDKLRM------LNGQTA  154
Human-PAX6-h     EIRDRLLSEGVCTNDNIPSVSSINRVLRNLASEKQQMGADGMYDKLRM------LNGQTG  154
t6559aep-ORF1    EIRDKLLSEQICDSDSVPSVSSINRIVRNRLGSSSHASMADLNSPILK------IDPDMT  234
Human-PAX5-s5    EIRDRLLAERVCDNDTVPSVSSINRIIRTKVQQPPNQP-------VPA----------SS  155
Human-PAX5-s7    EIRDRLLAERVCDNDTVPSVSSINRIIRTKVQQPPNQP-------VPA----------SS  155
Danre--PAX2A     EIRDRLLAEGVCDNDTVPSVSSINRIIRTKVQQPFHPSSDGTGTPLST----------AG  165
Mouse-PAX2-s10   EIRDRLLAEGICDNDTVPSVSSINRIIRTKVQQPFHPTPDGAGTGVTA----------PG  162
                 EIR  L6 e 6C nd  PS6SSINR66R

t11467aep-ORF1   HSQLNYTKDFFLKKELSSLFPVNNHLYSDK-TTKIYPQIYENLKYMHNCYYPHQNN----  149
t9974aep-ORF4    --FGNFGPF-NGLNAITSNQPD-------QNNYGQFP-------FAFNTGFPMLSNPMQQ  210
XENLA--PAX6      -----------TWGSRPGWYPG-----------------------TSVPGQPAQEG----  176
Human-PAX6-h     -----------SWGTRPGWYPG-----------------------TSVPGQPTQDG----  176
t6559aep-ORF1    HYFVHSGLP-IQIGAISTNFPGSIHAHLPRTTSGSYS-------ISGILGMAVPSS----  282
Human-PAX5-s5    HSIVSTGSV-TQ-----------VSSVSTDSAGSSYS-------ISGILGITSPSA----  192
Human-PAX5-s7    HSIVSTGSV-TQ-----------VSSVSTDSAGSSYS-------ISGILGITSPSA----  192
Danre--PAX2A     HTIVPSTAS-PP-----------VSSASNDP-VGSYS-------INGILGIPRSNG----  201
Mouse-PAX2-s10   HTIVPSTAS-PP-----------VSSASNDP-VGSYS-------INGILGIPRSNG----  198
                                                                  g
```

```
t11467aep-ORF1     ------------------------CINCM-----NYNFVNAGICAPPLLPISELRGD---  177
t9974aep-ORF4      QQQQLINLPVNQLSPDRKYKTNNEALQHMANQHFHEQAMQNRLCNDLPPPLEETSMHSHH  270
XENLA--PAX6        -------------------------------------------CQ---------------  178
Human-PAX6-h       -------------------------------------------CQ---------------  178
t6559aep-ORF1      -QGNL--------------GT-----------------LI--MC----------------  292
Human-PAX5-s5      -DTNK--------------RK-----------------RDEGI-----------------  203
Human-PAX5-s7      -DTNK--------------RK-----------------RDEGI-----------------  203
Danre--PAX2A       -EK----------------RK-----------------RDDVLWSGNHLDGRKIGYH---  224
Mouse-PAX2-s10     -EK----------------RK-----------------REEDV-----------------  207


t11467aep-ORF1     -------VNNAQS----KIGDIFQRD-RQIS---KDDLQERTKLVY-S----------CT  211
t9974aep-ORF4      AMQSNDISPNSNIINGDMKDDNYHEDDRLVKLGLSHD----------------NSDSHHH  314
XENLA--PAX6        --PQEGVGENTNSISSNGED----SDEAQMRLQLKRKLQRNRTSFT------------QE  220
Human-PAX6-h       --QQEGGGENTNSISSNGED----SDEAQMRLQLKRKLQRNRTSFT------------QE  220
t6559aep-ORF1      ---SDYSIPNV----------------QQIHSHVLND--EKGNVHIMQEGEIAPSIDHPT  331
Human-PAX5-s5      ---QESPVPNGH--SLPGR--------DFLRKQMRGD------LFT------------QQ  232
Human-PAX5-s7      ---QESPVPNGH--SLPGR--------DFLRKQMRGD------LFT------------QQ  232
Danre--PAX2A       --GSDGSGPNSD--SQGSV--------ESLRKHLRAD------AFT------------QQ  254
Mouse-PAX2-s10     ---SEGSVPNGD--SQSGV--------DSLRKHLRAD------TFT------------QQ  236
                            N                   6      d


t11467aep-ORF1     CLDCIKFRSEKALKLTPLYKNDSEYNMFSVQEFQDNTSYRLGMNERKKAHVAYTGNCFKQ  271
t9974aep-ORF4      HINDVDIHHD---------LSSHNHDILDNRGLK------RHIEES-----VMRGEIG-K  353
XENLA--PAX6        QIEALEKEFE---------RT-HYPDVFARERLA------AKIDL---------PEAR--  253
Human-PAX6-h       QIEALEKEFE---------RT-HYPDVFARERLA------AKIDL---------PEAR--  253
t6559aep-ORF1      QVNANEYVLE---------SSDSEHDSSAASSSS------LSNEED-----KFSNEQI--  369
Human-PAX5-s5      QLEVLDRVFE---------RQ-HYSDIFTT--------------TE-----PIKPEQA--  261
Human-PAX5-s7      QLEVLDRVFE---------RQ-HYSDIFTT--------------TE-----PIKPEQT--  261
Danre--PAX2A       QLEALDRVFE---------RP-SYPDVFPT--------------SE-----HIKPEQA--  283
Mouse-PAX2-s10     QLEALDRVFE---------RP-SYPDVFQA--------------SE-----HIKSEQG--  265
                   q6       e               1 f                           e
```

```
t11467aep-ORF1    LEETYKYFHCSNLNIKDDLQKRNEIRVH-------ANPKNLIPCKNAHV-----------   313
t9974aep-ORF4     EENNNIKSEVRQLSRNR----------------TDSLSPDMKRV----------------   381
XENLA--PAX6       ---IQVWFSNRRAKWRREEKLRNQRRQA------SNTPSHIPISSSFSASVYQPIPQPTT   304
Human-PAX6-h      ---IQVWFSNRRAKWRREEKLRNQRRQA------SNTPSHIPISSSFSTSVYQPIPQPTT   304
t6559aep-ORF1     ---NKPYYMKMDLNFDSENEKSTNKKASMQINGVDDSKFDMSLQSTIHHAFSHPNEQTNH   426
Human-PAX5-s5     ----PPTIIALPP----------------------EEPPHLQPPLPM-------------   282
Human-PAX5-s7     ----TEYSAMASLA-----------------GGLDDMKANLASPTPADIGSSVPGPQS--   298
Danre--PAX2A      ----NEYSLP-ALN-----------------PGLDEVKPSLSTSVSSDLGSSV--SQS--   317
Mouse-PAX2-s10    ----NEYSLP-ALT-----------------PGLDEVKSSLSASANPELGSNVSGTQT--   301
                                                          6

t11467aep-ORF1    ------------------------------------------------------------     -
t9974aep-ORF4     ----------------------------------------------KRYSESPLN-----   390
XENLA--PAX6       PVSSFTSGSMLGRTDT--ALTNSYSALPPM-----PSFTMGNNL--------PM-QPPVP   348
Human-PAX6-h      PVSSFTSGSMLGRTDT--ALTNTYSALPPM-----PSFTMANNL--------PM-QDSFP   348
t6559aep-ORF1     -----TFKKAMRRVRTTFSLEQ-------------------RRALEDAFEKTPYPDAEQR   462
Human-PAX5-s5     ---------TVT------------------------------------------------   285
Human-PAX5-s7     ------YPIVTG------------------------------------------------   304
Danre--PAX2A      ------YPVVTGREMASTTLPGYPPHVPPTGQGSYPTSTLAGMVPGSDFSGNPYSHPQYT   371
Mouse-PAX2-s10    ------YPVVTGRDMTSTTLPGYPPHVPPTGQGSYPTSTLAGMVPGS-------------   342

t11467aep-ORF1    ------------------------------------------------------------     -
t9974aep-ORF4     EQ---W--INQQKISPTWYKYNVFPNSNNIATANSMAAA-LNQINGAINSFQNIAQYSNG   444
XENLA--PAX6       SQTSSYSCMLPTSPSV---NGRSYDTYTPPH------------MQ-THMNSQP-------   385
Human-PAX6-h      LVCQ-FQFKFPEVNLICLNTGQDYSKKKKKK------------KKERKYCVNS-------   388
t6559aep-ORF1     EE-ISIQCDLPEPRVQVWFSNKRAKLRRQDRNEDKQDSSKNKEMISSHLDFQR-------   514
Human-PAX5-s5     ---------------DPWSQAGTKH-----------------------------------   295
Human-PAX5-s7     ---------------SPYYYSAAARGAAPPA---------------AA------------   322
Danre--PAX2A      TYNEAWRFSNPALLSSPYYYSAASRGSGPPT---------------AA------------   404
Mouse-PAX2-s10    ----------------PYYYSAAPRGSAPAA---------------AA------------   359
```

```
t11467aep-ORF1    ------------------------------------------------------------   -
t9974aep-ORF4     SNTLNGLNLNAYTSGLNNVNSLATLNAMNSMVNGNNTMNTLSGLQNLNGQGVSCFTPTFE 504
XENLA--PAX6       -------------------------------------MGTSGTT------STGLISPGVS 402
Human-PAX6-h      -------------------------------------VSDYGDT------T--------- 396
t6559aep-ORF1     ---------------------------------IQNHIKQYDCKNNSEENNQSCFSDEYQ 541
Human-PAX5-s5     ------------------------------------------------------------   -
Human-PAX5-s7     --------------------------------------TAYDRH---------------- 328
Danre--PAX2A      --------------------------------------TAYDRH---------------- 410
Mouse-PAX2-s10    --------------------------------------AAYDRH---------------- 365


t11467aep-ORF1    ------------------------------------------------------------   -
t9974aep-ORF4     SDRNISQNSLNGLGN--------NTDFNWSNNFIGVLSDGTPFQQAVSLTPNGTNSLANS 556
XENLA--PAX6       VPVQVPGSEPD--------------MSQYWPRLQ-------------------------- 422
Human-PAX6-h      --VELSGKKEKWLLE----------PLQFYNCVLYCTTGEGMDLKQGPLYTEGTISVG-- 442
t6559aep-ORF1     DQINVHDDVKNCLLNTELNTSSLPTDHRFTNRVLSEIKTSDSATNSCSLPPVSTLTKS-- 599
Human-PAX5-s5     ------------------------------------------------------------   -
Human-PAX5-s7     ------------------------------------------------------------   -
Danre--PAX2A      ------------------------------------------------------------   -
Mouse-PAX2-s10    ------------------------------------------------------------   -


t11467aep-ORF1    ------------------------------------------------------------   -
t9974aep-ORF4     SSFGLNMTFPSVNKNVMQGPQSPTNSVFAATLSSLQGQQNLFNFHTQLHNGMQTVQYQRP 616
XENLA--PAX6       ------------------------------------------------------------   -
Human-PAX6-h      ---------------------------------------------TNLHFGIQTFIHFGV 457
t6559aep-ORF1     ---------------------------------------------TNLHHDDGWPHQQRK 614
Human-PAX5-s5     ------------------------------------------------------------   -
Human-PAX5-s7     ------------------------------------------------------------   -
Danre--PAX2A      ------------------------------------------------------------   -
Mouse-PAX2-s10    ------------------------------------------------------------   -
```

```
t11467aep-ORF1   ------------------------    -
t9974aep-ORF4    DQLAMKLMGQSMAALRGGHVSHGN  640
XENLA--PAX6      ------------------------    -
Human-PAX6-h     ----LFVNGHLYVIMKKRTM----  473
t6559aep-ORF1    ----LFVLGNY-------------  621
Human-PAX5-s5    ------------------------    -
Human-PAX5-s7    ------------------------    -
Danre--PAX2A     ------------------------    -
Mouse-PAX2-s10   ------------------------    -
```

Phylogenetic tree of PAX proteins:

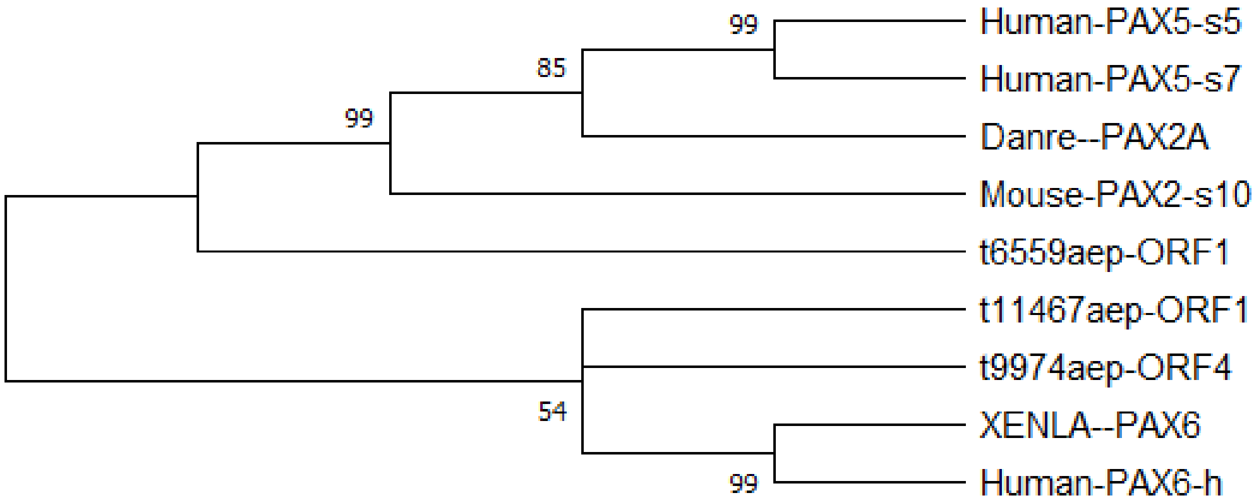


There are three *Hydra* Pax-genes with a conserved paired-DNA binding domain. They group with vertebrate Pax2/5 and Pax6.

---
